# Impact of vector control on effective population sizes; empirical evidence for a control-based genetic bottleneck in the tsetse fly *Glossina fuscipes*

**DOI:** 10.1101/2020.06.25.171678

**Authors:** Allan Muhwezi, Lucas J. Cunningham, Johan Esterhuizen, Inaki Tirados, Enock Matovu, Martin J. Donnelly, Stephen J. Torr

## Abstract

We investigated genetic variation at 37 newly-developed microsatellite loci in populations of the tsetse fly *Glossina fuscipes fuscipes* captured from the upper and lower reaches of a single hydrographical network within an endemic Human African Trypanosomiasis focus. Our primary aim was to assess the impact of vector control using insecticide-treated baits (Tiny Targets) on genetic structure. We initially used *STRUCTURE* to delineate geographical boundaries of two stable ‘ancestral’ reference populations without any history of vector control but marked for either vector control (‘intervention’) or no control (‘non-intervention’). We then used the *ADMIXTURE* model to assess genetic divergence in temporal populations collected after vector control implementation. We applied the Linkage Disequilibrium method to explicitly measure spatial and temporal changes in effective population size (*N_e_*). We observed a significant reduction in *N_e_* coincident with vector control, whereas *N_e_* remained stable in the non-intervention area. Our empirical findings show how classical population genetics approaches detected within a short period of time, a significant genetic bottleneck associated with vector control, and opens up the possibility of using routine genomic surveillance. We have also generated a resource of new genetic markers for studies on the population genetics of tsetse at finer-scale resolution.

**Funding:** This work was funded through a Wellcome Trust Master’s Fellowship in Public Health and Tropical Medicine awarded to Allan Muhwezi (103268/Z/13/Z).

## Introduction

Human African Trypanosomiasis (HAT) or Sleeping sickness is a deadly parasitic disease endemic to tropical Africa. The disease is caused by subspecies of *Trypanosoma brucei* and is transmitted by tsetse flies (*Glossina* spp). There are two aetiological agents: *T. b. gambiense,* accounting for >95% of cases, and *T. b. rhodesiense*, which exhibit separate geographical ranges, ecologies, vectors, rates of disease progression and control strategies. Both diseases are usually fatal if left untreated.

Vector control has recently emerged as an important strategy in efforts to control and eliminate HAT (Lehane et al., 2016). This has greatly been facilitated by the development of eco-friendly tools (Torr et al., 2007), and new methods such as ‘Tiny Targets’, blue-black panels of insecticide treated baits, that are cost-effective and easier to deploy at large-scale in disease endemic foci (Lindh et al., 2009). A recent field trial of this new technology over a 500 Km^2^ area in North-Western Uganda demonstrated a significant reduction in tsetse catches by ≈90% (Tirados et al., 2015). What this and other studies (Courtin et al., 2015, Mahamat et al., 2017) revealed was that monitoring the impact of vector control using sentinel traps is not only expensive but the low numbers of tsetse captured, particularly following the deployment of targets, make robust quantitative estimation of impact difficult. We postulated that genetic diversity methods may be a sensitive and more cost-effective method for estimating intervention impact by maximising the information derived from each collected individual.

In this study we used a panel of microsatellite markers that we recently developed, to assess whether vector control operations using the Tiny Targets were having an underlying effect of on the genetic structure of tsetse populations.

## Methodology

### Study site

The study was conducted in North-West Uganda along the Kochi River, the primary watercourse, in the Koboko HAT focus located between 3.451-3.465°N and 30°58^’^-31°03’E (shown in Figure 1). The river is about 70 km long and drains into the River Nile. The upper Kochi, within the treatment (intervention) block, is characterized by a patchwork of farmland and degraded natural savanna woodland. The lower reaches of the Kochi, outside the treatment (non-intervention) area, consist of dry thickets and wooded sites with fewer farms and a sparse human population.

**Figure 1.**
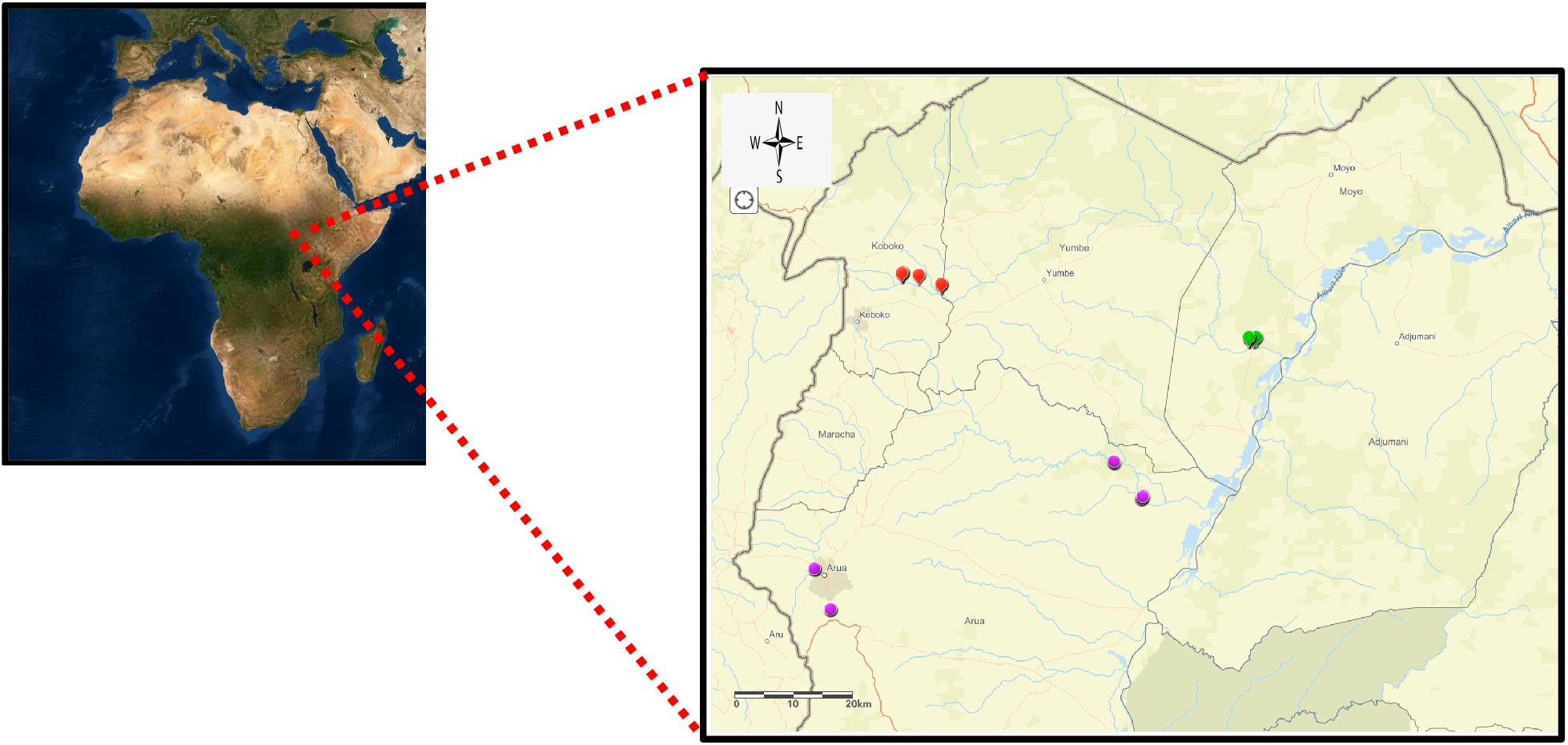
Sampling sites along rivers Kochi and Enyau in North-West Uganda. Red dots show tsetse collection sites that are within the vector control treatment (intervention) block, whereas orange dots show trap sites within the no vector control treatment (non-intervention) block. Purple dots show collection sites along the Enyau river (The map was created using ArcGIS: https://www.arcgis.com/index.html#).

To investigate the degree of inter-connectedness between adjacent river systems and therefore potential for reinvasion, we compared the relatedness of *G. f. fuscipes* collections from River Kochi with collections from the neighbouring River Enyau, pre-intervention (shown in Figure 1). The sampling sites upstream on the River Enyau were in a mixture of grassland and bushland, with dry thickets and wooded sites downstream.

### Tsetse collection

*Glossina f. fuscipes* were collected in March and August 2014, in the upper and lower Kochi areas, before any vector control implementation, as well as along River Enyau. This was then followed by the deployment of the Tiny Targets between January 2016 within the area of Koboko District extending along the upper ends of River Kochi. Tsetse were then collected during March 2016 in both the intervention and non-intervention blocks. Details of the collections are summarized in Table 1.

**Table 1.** Details of tsetse specimen sampled in both the intervention and non-intervention blocks along River Kochi

| Collection site | Collection date | Gender | Sample size |
| --- | --- | --- | --- |
| Lower Kochi | August 2014 | Female = 30<br>Male = 18 | 48 |
|  | March 2016 | Female = 34<br>Male = 14 | 48 |
| Upper Kochi | March 2014 | Female = 32<br>Male = 16 | 48 |
|  | March 2016 | Female = 31<br>Male = 17 | 48 |

Samples of *G. f. fuscipes* were collected using pyramidal traps deployed along the river bank. GPS coordinates for every trapping location were recorded. Traps were placed at least 100 m apart and emptied at 24 h intervals over a three-day period per sampling site. Captured tsetse were individually preserved in 95% ethanol and stored at −20°C for subsequent analyses.

### DNA extraction and microsatellite genotyping

We initially developed a panel of 41 novel microsatellite markers using *in silico* and experimental approaches, spanning 96.9% of the ~374 Mb tsetse genome. DNA was then extracted for each *G. f. fuscipes* specimen using the Qiagen DNeasy blood and tissue kit (Qiagen, Inc., Valencia, CA). The tsetse legs, head and thorax were used. A total of 48 *G. f. fuscipes* specimen per treatment group, comprising both males and females, were then genotyped using 41 pairs of microsatellite primers grouped into eight panels (summarized in Table 2). The sample size choice was based on a simulation study that predicted precise estimations of *N_e_* for organisms with relatively low *N_e_* < 200 could be obtained with 10-20 loci, 10 alleles and 50 samples (Waples and Do, 2010).

**Table 2.**
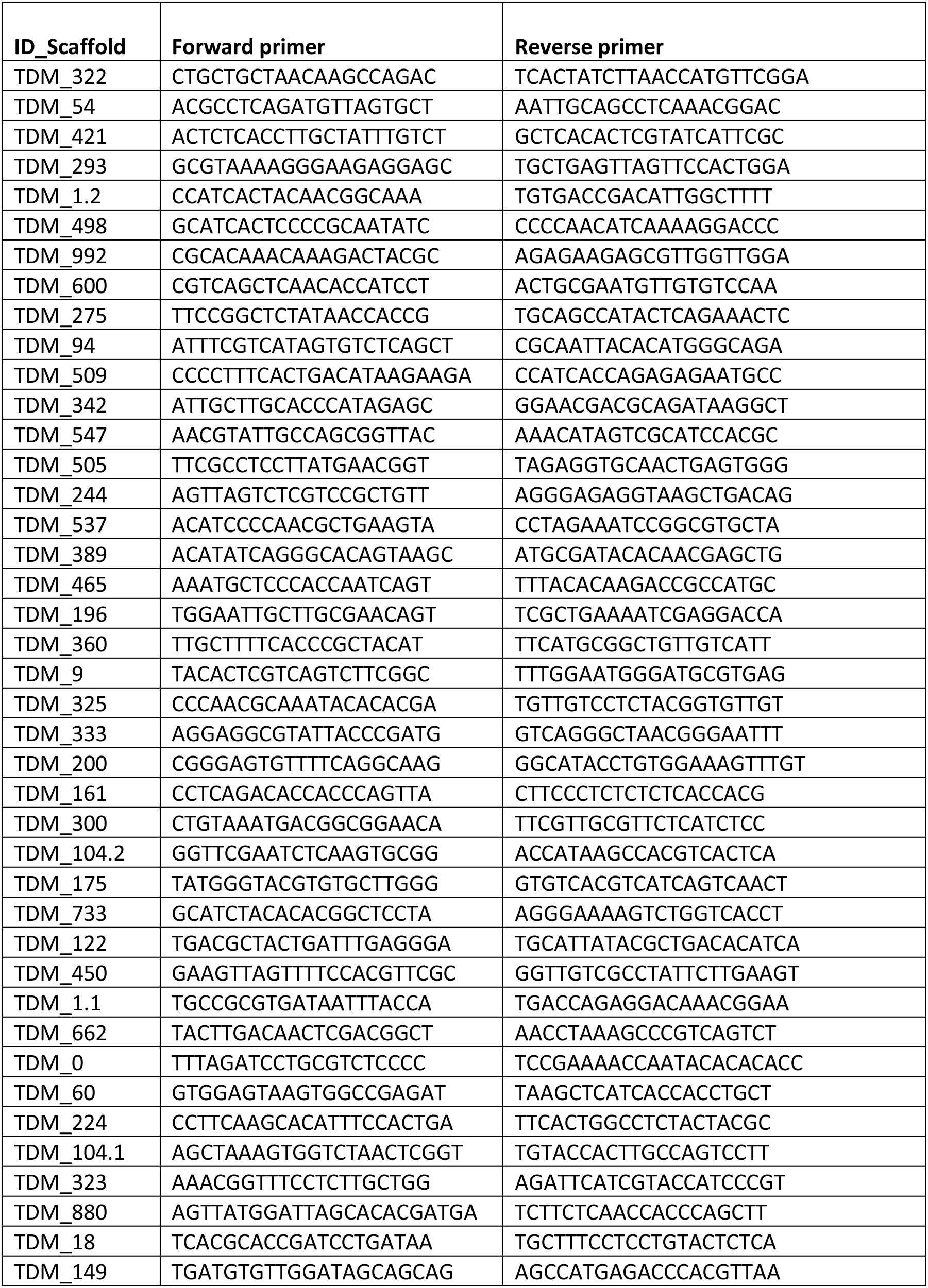
Tsetse microsatellite primer sequences developed in this study

To run the PCR reactions, we initially prepared a mix in water containing 2 μM of each of the primers per panel. For the reaction mix, we added 6.25 μl of a 2x Type-it Multiplex PCR master mix, 1.25 μL of the 10x primer mix and 3 μL of RNase-free water. 10.5 μL of this reaction mix was then dispensed into separate wells. 2 μL template DNA was then added to the individual PCR wells containing the reaction mix making a total volume of 12.5 μl. For the PCR program, we started with an initial heat-activation step at 95°C for 5 min to activate the HotStarTaq Plus DNA Polymerase, followed by 28 three-step cycles of denaturation at 95°C for 30 s, annealing at 60°C for 90 s, and extension at 72°C for 30 s. This was then followed by a final extension at 60°C for 30 min. Agarose gels for each product were then run to confirm success of the PCR. 1:10 dilutions of each of the PCR product were then prepared and the samples shipped to Macrogen, South Korea for genotyping.

Electropherograms were scored using GeneMarker (SoftGenetics). Fragment sizes were exported into Excel and then analysed using R software package MsatAllele (Alberto, 2009), to obtain the final allele classifications.

### Data analysis

We tested hypotheses of presence of population structure in the tsetse collections, due to either geographical isolation and/ or drift associated with vector control intervention.

### Microsatellite characterization

We explicitly checked the loci for patterns of sex-linked inheritance. We constructed two genotype files, one containing males and the other containing females and then ran allele frequency analyses on each data set. Polymorphic X-linked loci will have observed heterozygosity of zero in males and non-zero in females.

We then determined the allelic richness in the entire dataset and the polymorphic information content (PIC) of only 40 loci since one locus characteristically had many stutter peaks, that made it very difficult to generate a score that reflects the true genotype. We calculated both the expected and observed heterozygosity in the population and the frequency of null alleles using two independent methods based on either an iterative expectation and maximization (EM) approach implemented in GENEPOP (Raymond and Rousset, 1995), or a likelihood approach implemented in CERVUS (Kalinowski et al., 2007).

### Contemporary *N_e_* and Linkage Disequilibrium (LD)

We estimated *N_e_* using a single-sample LD method (Waples and Do, 2010). An effect of low population size is that the small number of parents usually produce offspring with increased levels of random LD, which can be measured as a product of the squared correlation (*r^2^*) of alleles at different loci. If the loci are unlinked, then the magnitude of LD can be estimated as:

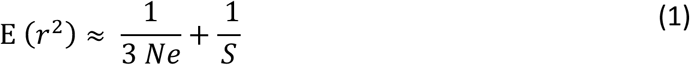

Where *S* in the number of individuals in the sample. There are many allele combinations when many polymorphic markers are used and this substantially increases the power of the LD method to detect drift.

Two estimates of *N_e_* were calculated based on either a randomly mating population without selfing or a population that randomly mates and then undergoes lifetime monogamy. All calculations were performed using NeEstimator V2.1 (Do et al., 2014). We excluded alleles at a frequency of <5% as described by (Waples and Do, 2010) since the LD method has little or no bias at *P_crit_* ≥ 0.05, and the empirical correction method was developed for such data that excluded alleles at a frequency of <5%.

The LD method also assumes a closed population with limited migration as well as loci that are under random recombination. We tested for deviations from Linkage Equilibrium using GENEPOP 4.5 (Raymond and Rousset, 1995).

### Bayesian clustering analyses

We employed a Bayesian clustering approach using *STRUCTURE* 2.0 (Pritchard et al., 2000), to delineate population boundaries and identify groups with distinctive allele frequencies. This method calculates the likelihood of data being grouped into a given number of clusters that maximize Hardy Weinberg Equilibrium and minimise LD. To perform this analysis, we set most parameters to their default values as recommended in the user’s manual (Pritchard et al., 2003). We chose the admixture model and correlated allele frequencies between populations, as this configuration has been considered optimal in subtle population structure (Falush et al., 2003, Rodríguez-Ramilo et al., 2009), and inferred the degree of admixture alpha from the data. Lambda, the parameter of the distribution of allelic frequencies, was set to one. The lengths of the burn-in and MCMC (Markov chain Monte Carlo) calculation were each set to 50000.

## Results

### Microsatellite characterization

We genotyped 41 microsatellite loci. Electropherograms for one locus had multiple peaks and this marker was dropped from the dataset. There was no evidence for patterns of sex-linked inheritance with any of the 40 microsatellite loci. Estimates of observed heterozygosity for all loci, in either female or the male stratifications were above 5% except for one locus that was monomorphic in both populations (shown in Supplementary 1). The average number of alleles per locus across the 192 samples was 7.2, ranging from 2 – 15 alleles. The Polymorphic Information Content (PIC) values ranged from 0.11 – 0.8 (shown in Supplementary 2). We restricted our analyses to 37 microsatellite loci where data were available from all temporal and spatial samples.

A possible draw back with microsatellite markers is the likelihood of null alleles, which could spuriously increase observed homozygosity estimates. Given that, the impact of these purported null alleles on downstream analyses in natural populations is not well known, we further filtered the genotyped panel down to 29 loci, with which 96.6% of the markers have null allele frequency estimates of ≤ 0.4 and 75.9% of markers have null allele frequency of ≤ 0.2. The latter is considered rare and uncommon with probably minimal effect on the exclusion probability of a locus (Dakin and Avise, 2004). Results using a panel of 37 loci were compared with those using a panel of 29 loci.

### Linkage Disequilibrium tests

Of the 2664 population-by-locus tests of LD after applying a Bonferroni correction, only 51 (1.9%.) showed significant departures. Similarly, for the panel of 29 loci, only 18 (1.1 %) tests showed significant (p<0.05) evidence of LD after applying a Bonferroni correction (shown in Supplementary 3).

### *STRUCTURE* analysis

We used the coefficient *Q-matrix* to deduce the proportion of membership initially for only the two pre-intervention reference populations (results summarized in Figure 2).

**Figure 2.**
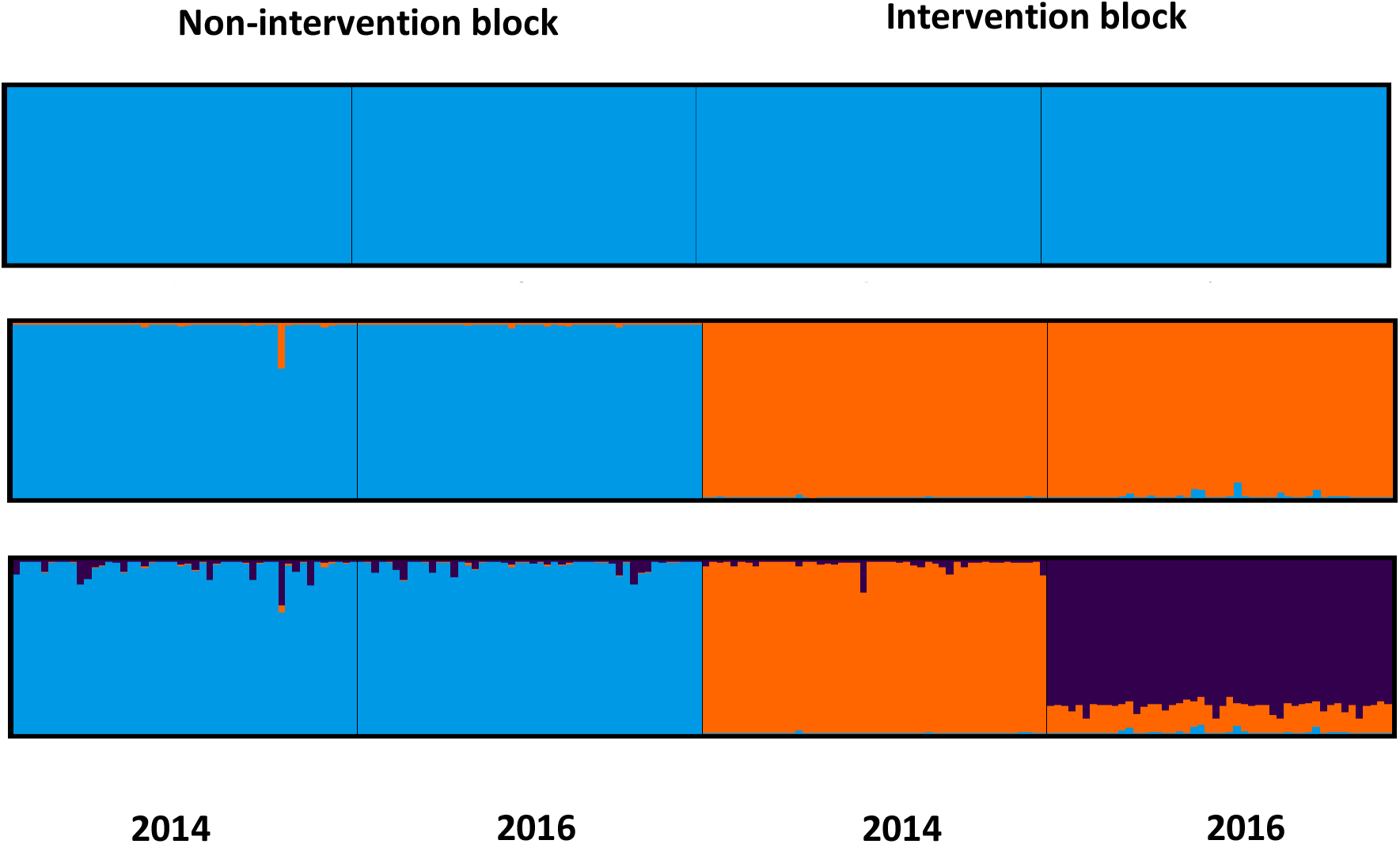
Estimated population structure of *G. f. fuscipes* along River Kochi. Each individual is represented by a thin vertical line, which is partitioned into *K* coloured segments that represent the individual’s estimated membership fractions in *K* clusters. Black lines separate individuals of different populations. Twenty-five *structure* runs at each *K* produced individual membership coefficients, having pairwise similarity coefficients above 0.97. The figure shown for *K* is based on the highest probability run at that

At *K=*2, there was a clear separation of populations upstream and downstream of the river with no traces of admixture or recent migrants, and membership of >99.6% to either of the two clusters inferred. This clearly suggests that the habitats upstream and downstream are discrete.

We then included the post-intervention collections into the analysis. At *K=*2, there was a clear spatial separation of populations based on geographical collection either upstream and downstream, with membership of >99% in either of the two clusters.

At *K*=3, there was clear genetic divergence in temporal samples in the intervention block, with membership of >98.4% in either of the two inferred clusters. Samples in the non-intervention block maintained admixture levels of 98.7% and 99.3%. This result suggests that temporal samples in the non-intervention arm share a more recent ancestor and are probably the same population, however for the temporal samples in the intervention, the results clearly suggest occurrence of a strong recent drift event, which has consequently resulted in divergence of allele frequencies.

### Contemporary *N_e_* estimates

We estimated *N_e_* using two models that assumed either random mating between individuals without selfing, or random mating followed by lifelong monogamy (summarized in Figure 3 and Table 3). In both models, *N_e_* estimates were temporally stable in the non-intervention area, and all had overlapping confidence intervals and thus suggestive of a non-significant difference.

**Table 3.** *N_e_* estimates in *G. f. fuscipes* collections along river Kochi

| Population | $N_e$ | 95% Parametric CI | | loci | Mating model |
| --- | --- | --- | --- | --- | --- |
|  |  | Lower | Upper |  |  |
| Downstream 2014 | 57.6 | 46.1 | 74.5 | 37 | Random mating |
| Downstream 2016 | 54.0 | 43.6 | 69.1 | 37 | Random mating |
| Upstream 2014 | 117.5 | 90 | 165 | 37 | Random mating |
| Upstream 2016 | 61.2 | 51 | 75.1 | 37 | Random mating |
| Downstream 2014 | 115.9 | 93.3 | 149.2 | 37 | Monogamy |
| Downstream 2016 | 109.1 | 88.5 | 139.1 | 37 | Monogamy |
| Upstream 2014 | 236.3 | 181.4 | 331.2 | 37 | Monogamy |
| Upstream 2016 | 123.8 | 103.6 | 151.7 | 37 | Monogamy |
| Downstream 2014 | 78.0 | 58 | 113.8 | 29 | Random mating |
| Downstream 2016 | 82.2 | 60 | 123.8 | 29 | Random mating |
| Upstream 2014 | 223.0 | 139 | 511.7 | 29 | Random mating |
| Upstream 2016 | 92.1 | 70.8 | 127.7 | 29 | Random mating |
| Downstream 2014 | 157.1 | 117.3 | 228.3 | 29 | Monogamy |
| Downstream 2016 | 165.9 | 121.6 | 249.0 | 29 | Monogamy |
| Upstream 2014 | 447.4 | 279.8 | 1024.0 | 29 | Monogamy |
| Upstream 2016 | 185.6 | 143.1 | 256.9 | 29 | Monogamy |

**Figure 3.**
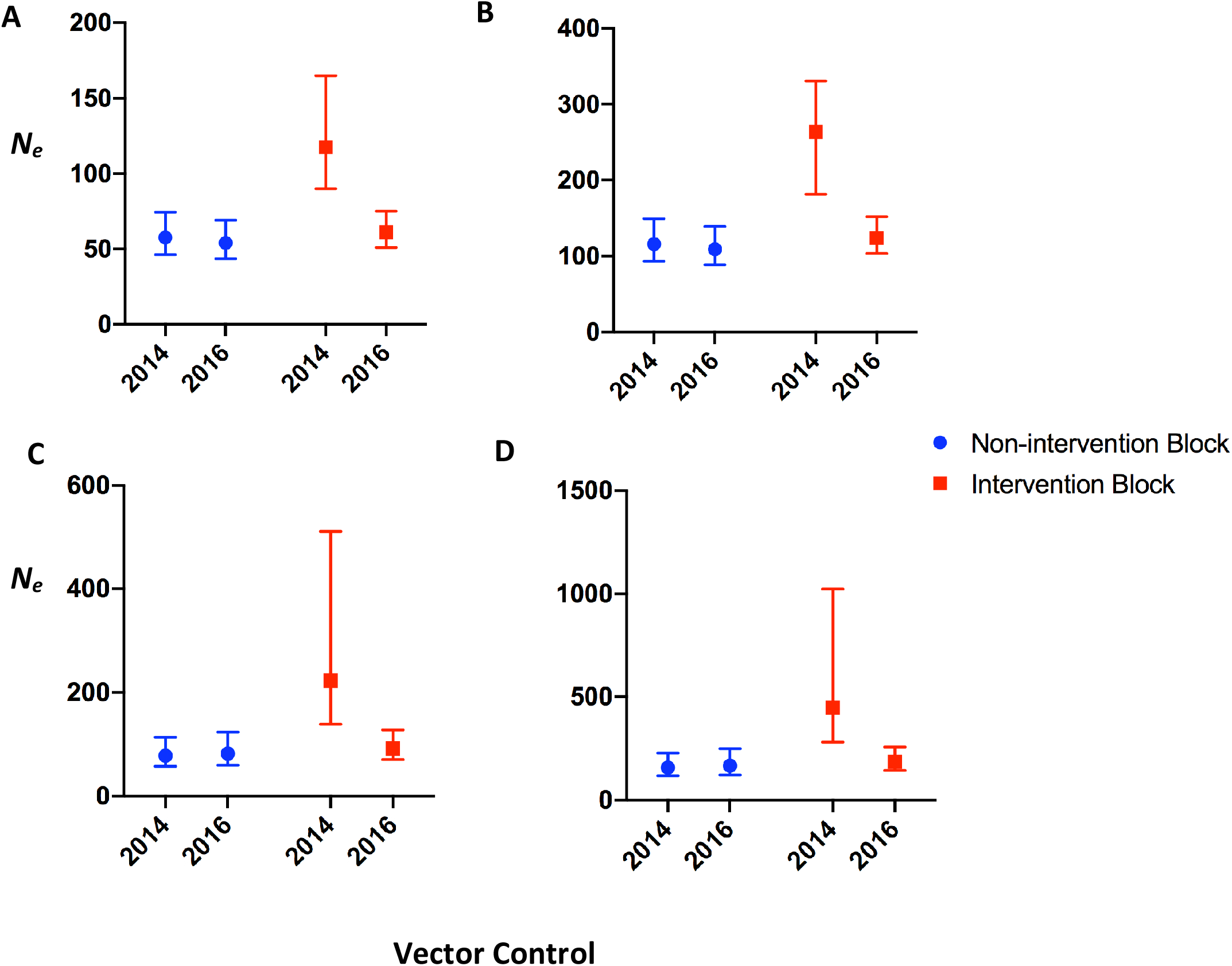
Temporal changes in effective population sizes in areas without vector control (blue dots) and areas with anti-vector Tiny Target tools (red dots), using either a panel of 37 microsatellite markers (A, B), or 29 microsatellite markers (C, D), and a model of a randomly mating population without selfing (A, C) or a population that randomly mates and then undergoes lifelong monogamy (B and D).

In contrast, within the intervention area, the introduction of the Tiny Targets was followed with a drastic reduction in *N_e_* of >50% from baseline levels.

Given that we did not know the impact of purported null allele frequencies on *N_e_* we also used a filtered data set of 29 microsatellite markers. There were no qualitative differences in the trend observed using either sample set. The random mating model followed with lifelong monogamy, arguably more relevant to tsetse flies, gave consistently higher estimates of *N_e_* in all the estimates.

### Movement of tsetse between rivers

We used *STRUCTURE* to assess if there were any recent migrants between rivers Kochi and Enyau, either upstream or downstream (shown in Figure 4). Populations show segregation both between and within rivers. At *K* = 3, there was clear genetic divergence in spatial samples upstream and downstream river Kochi, and between rivers Kochi and Enyau, with a mean similarity score of 99.7%. Spatial resolution in samples collected along river Enyau was not evident but this could possibly be as a result of the fewer samples genotyped from this area.

**Figure 4.**
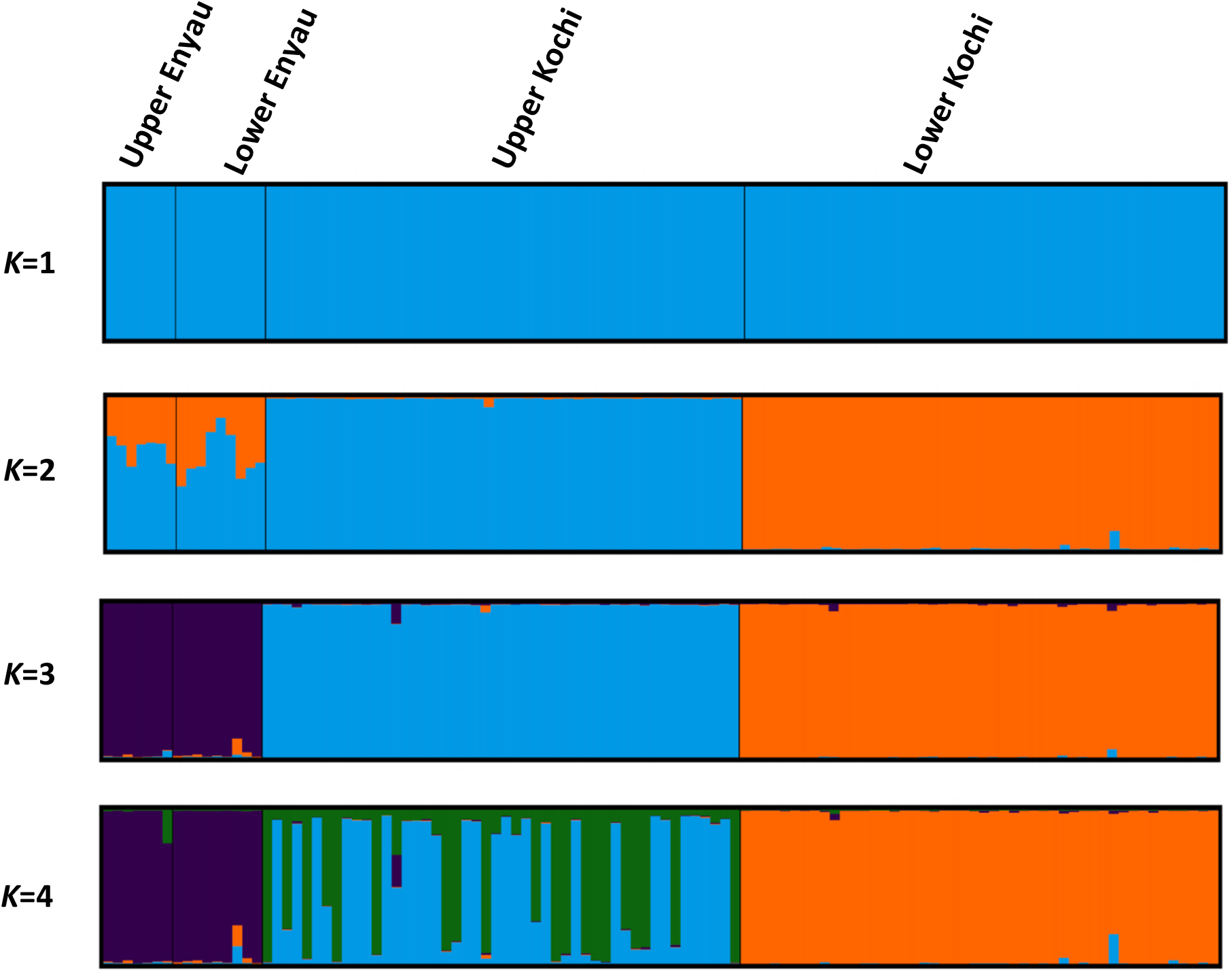
Estimated population structure of *G. f. fuscipes* between Rivers Kochi and Enyau. The figure shown for *K* is based on the highest probability run at that

## Discussion

We investigated genetic variation at 37 microsatellite loci in populations of *G. f. fuscipes* from the upper and lower reaches of a primary river tributary within an endemic HAT focus, in an attempt to assess the impact of vector control using Tiny Targets.

### Geospatial effects

The presence of genetically discrete populations of *G. f. fuscipes* along the single hydrographical network was assessed using both *a posteriori* and *a priori* knowledge about the origins of the individuals. This was necessary for our study in order to determine the independence of the two populations in estimating changes in *N_e_*. Methods for estimating *N_e_* are based on the assumption that the only factor responsible for changes in the genetic properties of a population is genetic drift and that systematic forces of mutation, selection and migration are absent.

Mutation rates are often very low and the gamete frequency changes caused by mutation are inversely proportional to population size, so that mutations usually make negligible contribution to overall levels of gametic disequilibrium (Hamilton, 2011).

Effects of selection could also be ignored given the neutrality of our markers, and direct selection on most markers are unlikely to be strong enough to cause substantial changes in allele frequencies. However, the effects of migration are not negligible and can substantially bias the estimates of *N_e_* either upwards or downwards (Wang and Whitlock, 2003). Our results reveal strong genetic structuring between tsetse samples from the upper and lower reaches of the river and within the neighbouring river system.

### Seasonal effects

Estimates of *N_e_* can be sensitive to strong reductions in population size. If population size varies between generations, *N_e_* will be the harmonic mean of the single generation effective sizes and thus will approximate the lowest size (Nei, 1987, Waples, 1991). Temporal variations in the *N_e_* of *G. f. fuscipes* in lower Kochi were 57.6 and 54, and the 95% CIs overlapped. Thus, extreme bottlenecks are unlikely to have occurred in these populations over the 1.5-year sampling interval. These *N_e_* results of temporal stability were consistent with other studies (Echodu et al., 2011, Hyseni et al., 2012, Opiro et al., 2016).

These results are also consistent with entomological data showing that in areas without any tsetse control, *G. fuscipes* numbers are relatively stable with mean daily catches ranging between 0.7 and 3.9 tsetse/trap (mean = 1.9) (Tirados et al., 2015). This interseasonal range in catches however greatly varies for savannah tsetse, ranging from 35- and 13-fold differences between the lowest and highest for *G. morsitans* and *G. pallidipes* (Hargrove and Vale, 1980), and has also been shown to vary for other insect disease vectors such as mosquitoes with lower catches during the dry season (Lehmann et al., 2014).

### Impact of vector control using the Tiny Targets

Our empirical findings conclusively show a severe genetic bottleneck seemingly caused by the deployment of Tiny Targets. The baseline *N_e_* in the intervention block was reduced by nearly 50% from 117.5 (95% CI; 90-165) to 61.2 (95% CI; 51-75.1), and these results are qualitatively consistent with field data showing a reduction in tsetse catches by up to 90% after implementation of Tiny Targets tools (Tirados et al., 2015). This was also supported by evidence of distinct population structure in populations before and after vector control implementation using Bayesian approaches.

The mean generation time of tsetse is approximately 73 days, and this would generally account for the time of deposition of the mother tsetse to the time of deposition of all her own offspring. In our study, effects of drift were evident throughout the 2-y sampling period equivalent to five generations. These results might probably hint to the presence of selective sweeps in genomic regions of these tsetse populations after sustained application of vector control.

### Implications of these results in efforts to control vector-borne diseases

The efficiency of vector control may be improved using knowledge on the population genetics of the target species. In this study, we have shown that there is large and significant genetic differentiation, and restricted gene flow within tsetse populations, and that spatially limited interventions can be effective.

Tsetse are highly mobile, dispersing diffusively at up to 1 km a day; studies of *G. f. fuscipes* in Uganda have shown that this species disperses at ~350 m/day (Rogers, 1977, Vale et al., 1984). A common long-term threat to tsetse control programmes is that tsetse from neighbouring areas will re-invade and/or survivors within the intervention area will form the basis of a new population once control is relaxed. Estimates of *N_e_* and population genetic structure indices, could help in assessing the suitability of the operational units selected for vector control and hence assist in the design of sustainable interventions.

## Acknowledgement

We would like to thank Jessica Lingley and Emily Rippon who helped through the lab work and the field team based in Uganda who assisted in the fieldwork collections including: Victor Drapari, Edward Aziku, and Henry Ombanya.

