## Supplemental Table 1 for "Impact of vector control on effective population sizes; empirical evidence for a control-based genetic bottleneck in the tsetse fly *Glossina fuscipes*"

**Table 1, S1**. Heterozygosity estimates in female *G. f. fuscipes* (n = 30)

| **Locus** | **k** | **N** | **HObs** | **HExp** |
|---|---|---|---|---|
| TDM_1.2 | 5 | 29 | 0.310 | 0.520 |
| TDM_293 | 3 | 29 | 0.103 | 0.164 |
| TDM_322 | 2 | 30 | 0.100 | 0.155 |
| TDM_421 | 5 | 29 | 0.414 | 0.458 |
| TDM_498 | 4 | 30 | 0.633 | 0.553 |
| TDM_54 | 2 | 30 | 0.067 | 0.427 |
| TDM_275 | 3 | 30 | 0.333 | 0.389 |
| TDM_509 | 6 | 30 | 0.567 | 0.676 |
| TDM_600 | 2 | 30 | 0.467 | 0.452 |
| TDM_04 | 2 | 30 | 0.533 | 0.499 |
| TDM_092 | 2 | 21 | 0.048 | 0.048 |
| TDM_244 | 2 | 29 | 0.207 | 0.407 |
| TDM_342 | 3 | 28 | 0.536 | 0.605 |
| TDM_389 | 3 | 29 | 0.483 | 0.417 |
| TDM_505 | 3 | 29 | 0.138 | 0.192 |
| TDM_537 | 3 | 28 | 0.214 | 0.231 |
| TDM_547 | 3 | 29 | 0.483 | 0.596 |
| TDM_196 | 6 | 30 | 0.633 | 0.738 |
| TDM_360 | 2 | 30 | 0.167 | 0.155 |
| TDM_465 | 6 | 30 | 0.867 | 0.824 |
| TDM_0 | 4 | 30 | 0.700 | 0.690 |
| TDM_161 | 5 | 30 | 0.533 | 0.546 |
| TDM_200 | 3 | 28 | 0.500 | 0.600 |
| TDM_325 | 3 | 30 | 0.733 | 0.666 |
| TDM_333 | 4 | 30 | 0.667 | 0.578 |
| TDM_104.2 | 3 | 27 | 0.444 | 0.616 |
| TDM_122 | 3 | 30 | 0.400 | 0.520 |
| TDM_175 | 3 | 30 | 0.267 | 0.320 |
| TDM_300 | 9 | 28 | 0.321 | 0.798 |
| TDM_733 | 2 | 30 | 0.167 | 0.155 |
| TDM_0 | 4 | 29 | 0.207 | 0.431 |
| TDM_1.1 | 3 | 29 | 0.517 | 0.642 |
| TDM_450 | 3 | 29 | 0.483 | 0.482 |
| TDM_60 | 4 | 30 | 0.367 | 0.376 |
| TDM_662 | 2 | 30 | 0.100 | 0.155 |
| TDM_104.1 | 1 | 30 | 0.000 | 0.000 |
| TDM_149 | 4 | 30 | 0.400 | 0.393 |
| TDM_224 | 5 | 30 | 0.333 | 0.377 |
| TDM_323 | 8 | 30 | 0.733 | 0.814 |
| TDM_880 | 3 | 28 | 0.500 | 0.408 |

**K** = number of alleles
**N** = number of individuals scored for a particular locus
**Hobs** = observed heterozygosity
**HExp** = expected heterozygosity

**Table 2, S1**. Heterozygosity estimates in male *G. f. fuscipes* (n = 18)

| **Locus** | **k** | **N** | **HObs** | **HExp** |
|---|---|---|---|---|
| TDM_1.2 | 5 | 18 | 0.222 | 0.735 |
| TDM_293 | 2 | 17 | 0.176 | 0.258 |
| TDM_322 | 2 | 18 | 0.167 | 0.386 |
| TDM_421 | 5 | 18 | 0.556 | 0.470 |
| TDM_498 | 3 | 18 | 0.556 | 0.589 |
| TDM_54 | 2 | 18 | 0.222 | 0.413 |
| TDM_275 | 3 | 18 | 0.222 | 0.552 |
| TDM_509 | 5 | 18 | 0.611 | 0.630 |
| TDM_600 | 2 | 18 | 0.333 | 0.457 |
| TDM_04 | 2 | 18 | 0.389 | 0.475 |
| TDM_092 | 2 | 11 | 0.273 | 0.247 |
| TDM_244 | 3 | 18 | 0.556 | 0.465 |
| TDM_342 | 3 | 18 | 0.611 | 0.586 |
| TDM_389 | 3 | 17 | 0.529 | 0.522 |
| TDM_505 | 3 | 18 | 0.222 | 0.427 |
| TDM_537 | 3 | 18 | 0.278 | 0.332 |
| TDM_547 | 3 | 18 | 0.611 | 0.595 |
| TDM_196 | 5 | 17 | 0.706 | 0.702 |
| TDM_360 | 2 | 18 | 0.111 | 0.108 |
| TDM_465 | 6 | 18 | 0.778 | 0.787 |
| TDM_0 | 4 | 17 | 0.941 | 0.742 |
| TDM_161 | 5 | 18 | 0.611 | 0.657 |
| TDM_200 | 3 | 17 | 0.412 | 0.561 |
| TDM_325 | 4 | 18 | 0.722 | 0.687 |
| TDM_333 | 4 | 18 | 0.444 | 0.538 |
| TDM_104.2 | 3 | 16 | 0.313 | 0.542 |
| TDM_122 | 3 | 18 | 0.222 | 0.608 |
| TDM_175 | 3 | 18 | 0.222 | 0.427 |
| TDM_300 | 4 | 15 | 0.133 | 0.717 |
| TDM_733 | 2 | 18 | 0.278 | 0.246 |
| TDM_0 | 3 | 16 | 0.125 | 0.234 |
| TDM_1.1 | 4 | 18 | 0.278 | 0.687 |
| TDM_450 | 3 | 18 | 0.333 | 0.427 |
| TDM_60 | 4 | 18 | 0.389 | 0.732 |
| TDM_662 | 2 | 18 | 0.167 | 0.386 |
| TDM_104.1 | 1 | 18 | 0.000 | 0.000 |
| TDM_149 | 3 | 18 | 0.500 | 0.567 |
| TDM_224 | 3 | 18 | 0.111 | 0.603 |
| TDM_323 | 8 | 18 | 0.667 | 0.806 |
| TDM_880 | 2 | 18 | 0.389 | 0.322 |

**K** = number of alleles
**N** = number of individuals scored for a particular locus
**Hobs** = observed heterozygosity
**HExp** = expected heterozygosity
