## Supplemental Table 2 for "Impact of vector control on effective population sizes; empirical evidence for a control-based genetic bottleneck in the tsetse fly *Glossina fuscipes*"

**Table 1, S2**. Characterization of 40 microsatellite markers based on a sample of 192 individuals

| Locus | k | N | HObs | HExp | PIC | F(Null)_1 | F(Null)_2 |
|---|---|---|---|---|---|---|---|
| TDM_1.2 | 8 | 190 | 0.295 | 0.782 | 0.748 | 0.4543 | 0.2755 |
| TDM_293 | 6 | 190 | 0.184 | 0.354 | 0.324 | 0.3192 | 0.2020 |
| TDM_322 | 2 | 192 | 0.135 | 0.242 | 0.212 | 0.2815 | 0.8898 |
| TDM_421 | 9 | 191 | 0.618 | 0.579 | 0.557 | -0.0466 | 0.0523 |
| TDM_498 | 10 | 192 | 0.620 | 0.670 | 0.611 | 0.0391 | 0.0379 |
| TDM_54 | 6 | 191 | 0.251 | 0.557 | 0.460 | 0.3780 | 0.2017 |
| TDM_275 | 9 | 192 | 0.292 | 0.501 | 0.466 | 0.2488 | 0.1655 |
| TDM_509 | 8 | 192 | 0.651 | 0.698 | 0.654 | 0.0362 | 0.0232 |
| TDM_600 | 3 | 192 | 0.552 | 0.502 | 0.411 | -0.0534 | 0.6005 |
| TDM_04 | 5 | 192 | 0.396 | 0.537 | 0.431 | 0.1390 | 0.1138 |
| TDM_092 | 2 | 141 | 0.099 | 0.254 | 0.221 | 0.4353 | 0.8952 |
| TDM_244 | 7 | 190 | 0.458 | 0.679 | 0.614 | 0.1932 | 0.1407 |
| TDM_342 | 4 | 189 | 0.519 | 0.539 | 0.486 | 0.0172 | 0.0493 |
| TDM_389 | 3 | 189 | 0.556 | 0.504 | 0.439 | -0.0525 | 0.0565 |
| TDM_505 | 9 | 143 | 0.378 | 0.552 | 0.524 | 0.1862 | 0.1583 |
| TDM_537 | 6 | 142 | 0.415 | 0.461 | 0.424 | 0.0691 | 0.1753 |
| TDM_547 | 8 | 190 | 0.611 | 0.790 | 0.755 | 0.1297 | 0.0960 |
| TDM_196 | 12 | 191 | 0.707 | 0.799 | 0.770 | 0.0585 | 0.0667 |
| TDM_360 | 6 | 192 | 0.193 | 0.193 | 0.186 | -0.0017 | 0.0672 |
| TDM_465 | 9 | 192 | 0.813 | 0.850 | 0.829 | 0.0226 | 0.0173 |
| TDM_0 | 4 | 143 | 0.531 | 0.586 | 0.538 | 0.0571 | 0.6060 |
| TDM_161 | 9 | 192 | 0.635 | 0.687 | 0.632 | 0.0388 | 0.0611 |
| TDM_200 | 10 | 188 | 0.452 | 0.720 | 0.684 | 0.2221 | 0.1687 |
| TDM_325 | 7 | 192 | 0.677 | 0.671 | 0.606 | -0.0043 | 0.0000 |
| TDM_333 | 8 | 192 | 0.635 | 0.756 | 0.714 | 0.0854 | 0.3740 |
| TDM_104.2 | 4 | 173 | 0.289 | 0.677 | 0.607 | 0.4009 | 0.4987 |
| TDM_122 | 5 | 191 | 0.435 | 0.651 | 0.604 | 0.1964 | 0.5983 |
| TDM_175 | 7 | 191 | 0.382 | 0.427 | 0.395 | 0.0632 | 0.0618 |
| TDM_300 | 15 | 181 | 0.525 | 0.862 | 0.845 | 0.2487 | 0.1771 |
| TDM_733 | 4 | 191 | 0.115 | 0.119 | 0.113 | 0.0114 | 0.9382 |
| TDM_0 | 9 | 174 | 0.213 | 0.804 | 0.774 | 0.5844 | 0.3794 |
| TDM_1.1 | 7 | 191 | 0.429 | 0.754 | 0.708 | 0.2770 | 0.1860 |
| TDM_450 | 12 | 191 | 0.524 | 0.541 | 0.506 | 0.0015 | 0.0441 |
| TDM_60 | 9 | 192 | 0.427 | 0.787 | 0.753 | 0.3013 | 0.2070 |
| TDM_662 | 2 | 192 | 0.146 | 0.257 | 0.224 | 0.2748 | 0.8809 |
| TDM_104.1 | 6 | 192 | 0.042 | 0.451 | 0.368 | 0.8332 | 0.2936 |
| TDM_149 | 7 | 192 | 0.500 | 0.651 | 0.609 | 0.1454 | 0.0862 |
| TDM_224 | 9 | 192 | 0.411 | 0.607 | 0.581 | 0.1867 | 0.1492 |
| TDM_323 | 13 | 192 | 0.740 | 0.858 | 0.841 | 0.0745 | 0.1396 |
| TDM_880 | 9 | 189 | 0.450 | 0.613 | 0.575 | 0.1658 | 0.2533 |

| | |
|---|---|
| Number of individuals: | 192 |
| Number of loci: | 40 |
| Mean number of alleles per locus: | 7.200 |
| Mean proportion of loci typed: | 0.9641 |
| Mean expected heterozygosity: | 0.5881 |
| Mean polymorphic information content (PIC): | 0.5449 |
| Combined non-exclusion probability (first parent): | 0.00001195 |
| Combined non-exclusion probability (second parent): | 9.004E-0010 |
| Combined non-exclusion probability (parent pair): | 3.989E-0016 |
| Combined non-exclusion probability (identity): | 8.959E-0030 |
| Combined non-exclusion probability (sib identity): | 1.034E-0012 |

K= number of alleles

N= number of individuals

Hobs= observed heterozygosity

HExp = expected heterozygosity

PIC= polymorphic information content

F(Null)_1 = purported null frequency based on a likelihood estimate

F(Null)_2 = purported null frequency based on expectation and maximization approach
