## Supplemental Table 3 for "Impact of vector control on effective population sizes; empirical evidence for a control-based genetic bottleneck in the tsetse fly *Glossina fuscipes*"

**Table 1, S3**. Linkage Disequilibrium tests for the panel of 37 loci

| | POP | Locus#1 | Locus#2 | P-Value | S.E. |
|---|---|---|---|---|---|
| 1 | LK2014 | A-1.2 | A-293 | 0.99942 | 0.00039 |
| 2 | LK2014 | A-1.2 | A-322 | 0.48476 | 0.01176 |
| 3 | LK2014 | A-293 | A-322 | 0.43883 | 0.00980 |
| 4 | LK2014 | A-1.2 | A-421 | 0.68698 | 0.01788 |
| 5 | LK2014 | A-293 | A-421 | 0.98989 | 0.00220 |
| 6 | LK2014 | A-322 | A-421 | 0.07164 | 0.00498 |
| 7 | LK2014 | A-1.2 | A-498 | 0.66753 | 0.01549 |
| 8 | LK2014 | A-293 | A-498 | 0.59981 | 0.01352 |
| 9 | LK2014 | A-322 | A-498 | 0.74380 | 0.00582 |
| 10 | LK2014 | A-421 | A-498 | 0.21877 | 0.01094 |
| 11 | LK2014 | A-1.2 | A-54 | 0.21447 | 0.00872 |
| 12 | LK2014 | A-293 | A-54 | 0.46071 | 0.00600 |
| 13 | LK2014 | A-322 | A-54 | 0.34287 | 0.00485 |
| 14 | LK2014 | A-421 | A-54 | 0.77585 | 0.00556 |
| 15 | LK2014 | A-498 | A-54 | 0.18868 | 0.00671 |
| 16 | LK2014 | A-1.2 | A-275 | 0.24972 | 0.01522 |
| 17 | LK2014 | A-293 | A-275 | 0.07832 | 0.00798 |
| 18 | LK2014 | A-322 | A-275 | 0.00000 | 0.00000 |
| 19 | LK2014 | A-421 | A-275 | 0.23023 | 0.01110 |
| 20 | LK2014 | A-498 | A-275 | 0.60473 | 0.01262 |
| 21 | LK2014 | A-54 | A-275 | 0.02542 | 0.00207 |
| 22 | LK2014 | A-1.2 | A-509 | 0.22022 | 0.02433 |
| 23 | LK2014 | A-293 | A-509 | 0.61518 | 0.02518 |
| 24 | LK2014 | A-322 | A-509 | 0.29323 | 0.01384 |
| 25 | LK2014 | A-421 | A-509 | 0.42348 | 0.02198 |
| 26 | LK2014 | A-498 | A-509 | 0.20537 | 0.01463 |
| 27 | LK2014 | A-54 | A-509 | 0.46905 | 0.01104 |
| 28 | LK2014 | A-275 | A-509 | 0.08608 | 0.01178 |
| 29 | LK2014 | A-1.2 | A-600 | 0.42118 | 0.01056 |
| 30 | LK2014 | A-293 | A-600 | 0.73453 | 0.00666 |
| 31 | LK2014 | A-322 | A-600 | 0.55643 | 0.00540 |
| 32 | LK2014 | A-421 | A-600 | 0.58008 | 0.00854 |
| 33 | LK2014 | A-498 | A-600 | 0.62091 | 0.00772 |
| 34 | LK2014 | A-54 | A-600 | 0.93016 | 0.00160 |
| 35 | LK2014 | A-275 | A-600 | 0.78501 | 0.00531 |
| 36 | LK2014 | A-509 | A-600 | 0.53389 | 0.01168 |
| 37 | LK2014 | A-1.2 | A-04 | 0.47537 | 0.01104 |
| 38 | LK2014 | A-293 | A-04 | 0.01641 | 0.00162 |
| 39 | LK2014 | A-322 | A-04 | 0.66095 | 0.00383 |
| 40 | LK2014 | A-421 | A-04 | 0.03252 | 0.00244 |
| 41 | LK2014 | A-498 | A-04 | 0.44272 | 0.00665 |
| 42 | LK2014 | A-54 | A-04 | 0.80738 | 0.00331 |
| 43 | LK2014 | A-275 | A-04 | 0.42121 | 0.00605 |
| 44 | LK2014 | A-509 | A-04 | 0.43937 | 0.00963 |
| 45 | LK2014 | A-600 | A-04 | 0.51810 | 0.00476 |

| | | | | | |
|---|---|---|---|---|---|
| 46 | LK2014 | A-1.2 | A-092 | 0.10845 | 0.00709 |
| 47 | LK2014 | A-293 | A-092 | 1.00000 | 0.00000 |
| 48 | LK2014 | A-322 | A-092 | 0.17027 | 0.00464 |
| 49 | LK2014 | A-421 | A-092 | 0.87614 | 0.00523 |
| 50 | LK2014 | A-498 | A-092 | 1.00000 | 0.00000 |
| 51 | LK2014 | A-54 | A-092 | 1.00000 | 0.00000 |
| 52 | LK2014 | A-275 | A-092 | 0.38456 | 0.00854 |
| 53 | LK2014 | A-509 | A-092 | 0.55022 | 0.01597 |
| 54 | LK2014 | A-600 | A-092 | 0.00013 | 0.00009 |
| 55 | LK2014 | A-04 | A-092 | 0.83132 | 0.00290 |
| 56 | LK2014 | A-1.2 | A-244 | 0.54970 | 0.01231 |
| 57 | LK2014 | A-293 | A-244 | 0.72536 | 0.00715 |
| 58 | LK2014 | A-322 | A-244 | 0.33659 | 0.00581 |
| 59 | LK2014 | A-421 | A-244 | 0.08861 | 0.00502 |
| 60 | LK2014 | A-498 | A-244 | 0.33673 | 0.00701 |
| 61 | LK2014 | A-54 | A-244 | 0.54634 | 0.00602 |
| 62 | LK2014 | A-275 | A-244 | 0.62878 | 0.00745 |
| 63 | LK2014 | A-509 | A-244 | 0.54222 | 0.01136 |
| 64 | LK2014 | A-600 | A-244 | 0.19525 | 0.00524 |
| 65 | LK2014 | A-04 | A-244 | 0.38140 | 0.00557 |
| 66 | LK2014 | A-092 | A-244 | 0.33387 | 0.00543 |
| 67 | LK2014 | A-1.2 | A-342 | 0.47521 | 0.02203 |
| 68 | LK2014 | A-293 | A-342 | 0.07926 | 0.00990 |
| 69 | LK2014 | A-322 | A-342 | 0.34894 | 0.00832 |
| 70 | LK2014 | A-421 | A-342 | 0.67532 | 0.01379 |
| 71 | LK2014 | A-498 | A-342 | 0.46485 | 0.01369 |
| 72 | LK2014 | A-54 | A-342 | 0.72004 | 0.00804 |
| 73 | LK2014 | A-275 | A-342 | 0.33642 | 0.01385 |
| 74 | LK2014 | A-509 | A-342 | 0.76727 | 0.01904 |
| 75 | LK2014 | A-600 | A-342 | 0.31135 | 0.00939 |
| 76 | LK2014 | A-04 | A-342 | 0.68648 | 0.00660 |
| 77 | LK2014 | A-092 | A-342 | 1.00000 | 0.00000 |
| 78 | LK2014 | A-244 | A-342 | 0.98834 | 0.00125 |
| 79 | LK2014 | A-1.2 | A-389 | 0.54166 | 0.01803 |
| 80 | LK2014 | A-293 | A-389 | 0.15076 | 0.00971 |
| 81 | LK2014 | A-322 | A-389 | 0.96227 | 0.00166 |
| 82 | LK2014 | A-421 | A-389 | 0.60064 | 0.01087 |
| 83 | LK2014 | A-498 | A-389 | 0.16228 | 0.00753 |
| 84 | LK2014 | A-54 | A-389 | 0.23972 | 0.00615 |
| 85 | LK2014 | A-275 | A-389 | 0.92941 | 0.00401 |
| 86 | LK2014 | A-509 | A-389 | 0.55193 | 0.01854 |
| 87 | LK2014 | A-600 | A-389 | 0.34109 | 0.00699 |
| 88 | LK2014 | A-04 | A-389 | 0.22359 | 0.00716 |
| 89 | LK2014 | A-092 | A-389 | 0.72631 | 0.00709 |
| 90 | LK2014 | A-244 | A-389 | 0.02021 | 0.00210 |
| 91 | LK2014 | A-342 | A-389 | 0.71503 | 0.01133 |
| 92 | LK2014 | A-1.2 | A-547 | 0.53590 | 0.01364 |
| 93 | LK2014 | A-293 | A-547 | 0.56931 | 0.01474 |

| | | | | | |
|---|---|---|---|---|---|
| 94 | LK2014 | A-322 | A-547 | 0.31939 | 0.00635 |
| 95 | LK2014 | A-421 | A-547 | 0.73735 | 0.00849 |
| 96 | LK2014 | A-498 | A-547 | 0.02407 | 0.00293 |
| 97 | LK2014 | A-54 | A-547 | 0.97569 | 0.00115 |
| 98 | LK2014 | A-275 | A-547 | 0.30719 | 0.01056 |
| 99 | LK2014 | A-509 | A-547 | 0.15260 | 0.01399 |
| 100 | LK2014 | A-600 | A-547 | 0.65350 | 0.00721 |
| 101 | LK2014 | A-04 | A-547 | 0.15518 | 0.00524 |
| 102 | LK2014 | A-092 | A-547 | 1.00000 | 0.00000 |
| 103 | LK2014 | A-244 | A-547 | 0.54007 | 0.00745 |
| 104 | LK2014 | A-342 | A-547 | 0.34838 | 0.01151 |
| 105 | LK2014 | A-389 | A-547 | 0.23158 | 0.00773 |
| 106 | LK2014 | A-1.2 | A-196 | 0.87338 | 0.01741 |
| 107 | LK2014 | A-293 | A-196 | 0.50988 | 0.02792 |
| 108 | LK2014 | A-322 | A-196 | 0.59225 | 0.01702 |
| 109 | LK2014 | A-421 | A-196 | 0.84403 | 0.01602 |
| 110 | LK2014 | A-498 | A-196 | 0.27953 | 0.02119 |
| 111 | LK2014 | A-54 | A-196 | 0.87010 | 0.00750 |
| 112 | LK2014 | A-275 | A-196 | 0.72983 | 0.02332 |
| 113 | LK2014 | A-509 | A-196 | 0.56971 | 0.03343 |
| 114 | LK2014 | A-600 | A-196 | 0.04390 | 0.00476 |
| 115 | LK2014 | A-04 | A-196 | 0.56800 | 0.01269 |
| 116 | LK2014 | A-092 | A-196 | 0.26580 | 0.01450 |
| 117 | LK2014 | A-244 | A-196 | 0.49469 | 0.01407 |
| 118 | LK2014 | A-342 | A-196 | 0.25612 | 0.02207 |
| 119 | LK2014 | A-389 | A-196 | 0.21874 | 0.01822 |
| 120 | LK2014 | A-547 | A-196 | 0.88528 | 0.01121 |
| 121 | LK2014 | A-1.2 | A-360 | 0.60703 | 0.01548 |
| 122 | LK2014 | A-293 | A-360 | 0.14541 | 0.01050 |
| 123 | LK2014 | A-322 | A-360 | 0.58828 | 0.00714 |
| 124 | LK2014 | A-421 | A-360 | 0.57725 | 0.01278 |
| 125 | LK2014 | A-498 | A-360 | 0.28977 | 0.01176 |
| 126 | LK2014 | A-54 | A-360 | 0.72906 | 0.00529 |
| 127 | LK2014 | A-275 | A-360 | 0.71471 | 0.01057 |
| 128 | LK2014 | A-509 | A-360 | 0.19793 | 0.01615 |
| 129 | LK2014 | A-600 | A-360 | 0.73483 | 0.00493 |
| 130 | LK2014 | A-04 | A-360 | 0.28689 | 0.00792 |
| 131 | LK2014 | A-092 | A-360 | 1.00000 | 0.00000 |
| 132 | LK2014 | A-244 | A-360 | 0.63003 | 0.00753 |
| 133 | LK2014 | A-342 | A-360 | 0.86730 | 0.00894 |
| 134 | LK2014 | A-389 | A-360 | 0.61788 | 0.00963 |
| 135 | LK2014 | A-547 | A-360 | 0.66786 | 0.00911 |
| 136 | LK2014 | A-196 | A-360 | 0.80469 | 0.01601 |
| 137 | LK2014 | A-1.2 | A-465 | 0.88489 | 0.01725 |
| 138 | LK2014 | A-293 | A-465 | 0.05832 | 0.01059 |
| 139 | LK2014 | A-322 | A-465 | 0.03830 | 0.00473 |
| 140 | LK2014 | A-421 | A-465 | 0.59387 | 0.02290 |
| 141 | LK2014 | A-498 | A-465 | 0.67973 | 0.01997 |

| | | | | | |
|---|---|---|---|---|---|
| 142 | LK2014 | A-54 | A-465 | 0.38889 | 0.01338 |
| 143 | LK2014 | A-275 | A-465 | 0.44256 | 0.02555 |
| 144 | LK2014 | A-509 | A-465 | 0.87626 | 0.01996 |
| 145 | LK2014 | A-600 | A-465 | 0.07330 | 0.00772 |
| 146 | LK2014 | A-04 | A-465 | 0.15632 | 0.00905 |
| 147 | LK2014 | A-092 | A-465 | 0.81091 | 0.01139 |
| 148 | LK2014 | A-244 | A-465 | 0.53106 | 0.01459 |
| 149 | LK2014 | A-342 | A-465 | 0.23940 | 0.02367 |
| 150 | LK2014 | A-389 | A-465 | 0.55210 | 0.02233 |
| 151 | LK2014 | A-547 | A-465 | 0.51596 | 0.02109 |
| 152 | LK2014 | A-196 | A-465 | 0.99822 | 0.00118 |
| 153 | LK2014 | A-360 | A-465 | 0.50141 | 0.02254 |
| 154 | LK2014 | A-1.2 | A-161 | 0.68393 | 0.01966 |
| 155 | LK2014 | A-293 | A-161 | 0.03697 | 0.00683 |
| 156 | LK2014 | A-322 | A-161 | 0.88948 | 0.00523 |
| 157 | LK2014 | A-421 | A-161 | 0.24359 | 0.01501 |
| 158 | LK2014 | A-498 | A-161 | 0.18479 | 0.01145 |
| 159 | LK2014 | A-54 | A-161 | 0.57682 | 0.01064 |
| 160 | LK2014 | A-275 | A-161 | 0.94457 | 0.00610 |
| 161 | LK2014 | A-509 | A-161 | 0.65097 | 0.02560 |
| 162 | LK2014 | A-600 | A-161 | 0.61144 | 0.01063 |
| 163 | LK2014 | A-04 | A-161 | 0.31711 | 0.00970 |
| 164 | LK2014 | A-092 | A-161 | 0.84570 | 0.00789 |
| 165 | LK2014 | A-244 | A-161 | 0.88964 | 0.00577 |
| 166 | LK2014 | A-342 | A-161 | 0.35235 | 0.01932 |
| 167 | LK2014 | A-389 | A-161 | 0.66266 | 0.01466 |
| 168 | LK2014 | A-547 | A-161 | 0.68542 | 0.01291 |
| 169 | LK2014 | A-196 | A-161 | 0.72136 | 0.02346 |
| 170 | LK2014 | A-360 | A-161 | 0.09029 | 0.00904 |
| 171 | LK2014 | A-465 | A-161 | 0.20759 | 0.02262 |
| 172 | LK2014 | A-1.2 | A-200 | 0.01180 | 0.00401 |
| 173 | LK2014 | A-293 | A-200 | 0.26637 | 0.01798 |
| 174 | LK2014 | A-322 | A-200 | 0.00000 | 0.00000 |
| 175 | LK2014 | A-421 | A-200 | 0.49250 | 0.01695 |
| 176 | LK2014 | A-498 | A-200 | 0.06807 | 0.00702 |
| 177 | LK2014 | A-54 | A-200 | 0.65246 | 0.00813 |
| 178 | LK2014 | A-275 | A-200 | 0.00388 | 0.00227 |
| 179 | LK2014 | A-509 | A-200 | 0.62611 | 0.02445 |
| 180 | LK2014 | A-600 | A-200 | 0.92880 | 0.00385 |
| 181 | LK2014 | A-04 | A-200 | 0.45971 | 0.01024 |
| 182 | LK2014 | A-092 | A-200 | 0.25291 | 0.00843 |
| 183 | LK2014 | A-244 | A-200 | 0.24260 | 0.01006 |
| 184 | LK2014 | A-342 | A-200 | 0.54947 | 0.01637 |
| 185 | LK2014 | A-389 | A-200 | 0.20552 | 0.01367 |
| 186 | LK2014 | A-547 | A-200 | 0.18738 | 0.01162 |
| 187 | LK2014 | A-196 | A-200 | 0.01105 | 0.00443 |
| 188 | LK2014 | A-360 | A-200 | 0.36210 | 0.01464 |
| 189 | LK2014 | A-465 | A-200 | 0.23190 | 0.02669 |

| | | | | | |
|---|---|---|---|---|---|
| 190 | LK2014 | A-161 | A-200 | 0.03120 | 0.00804 |
| 191 | LK2014 | A-1.2 | A-325 | 0.09967 | 0.01250 |
| 192 | LK2014 | A-293 | A-325 | 0.30423 | 0.01383 |
| 193 | LK2014 | A-322 | A-325 | 0.74959 | 0.00791 |
| 194 | LK2014 | A-421 | A-325 | 0.28718 | 0.01256 |
| 195 | LK2014 | A-498 | A-325 | 0.03552 | 0.00471 |
| 196 | LK2014 | A-54 | A-325 | 0.10021 | 0.00593 |
| 197 | LK2014 | A-275 | A-325 | 0.59226 | 0.01440 |
| 198 | LK2014 | A-509 | A-325 | 0.27378 | 0.02054 |
| 199 | LK2014 | A-600 | A-325 | 0.15246 | 0.00704 |
| 200 | LK2014 | A-04 | A-325 | 0.57888 | 0.00829 |
| 201 | LK2014 | A-092 | A-325 | 0.36027 | 0.00841 |
| 202 | LK2014 | A-244 | A-325 | 0.10401 | 0.00610 |
| 203 | LK2014 | A-342 | A-325 | 0.30517 | 0.01518 |
| 204 | LK2014 | A-389 | A-325 | 0.62447 | 0.01319 |
| 205 | LK2014 | A-547 | A-325 | 0.49127 | 0.01240 |
| 206 | LK2014 | A-196 | A-325 | 0.34049 | 0.02362 |
| 207 | LK2014 | A-360 | A-325 | 0.22871 | 0.01065 |
| 208 | LK2014 | A-465 | A-325 | 0.04878 | 0.00972 |
| 209 | LK2014 | A-161 | A-325 | 0.57384 | 0.01783 |
| 210 | LK2014 | A-200 | A-325 | 0.32686 | 0.01626 |
| 211 | LK2014 | A-1.2 | A-333 | 0.70265 | 0.02475 |
| 212 | LK2014 | A-293 | A-333 | 0.09512 | 0.01144 |
| 213 | LK2014 | A-322 | A-333 | 0.01990 | 0.00272 |
| 214 | LK2014 | A-421 | A-333 | 0.15496 | 0.01608 |
| 215 | LK2014 | A-498 | A-333 | 0.04182 | 0.00624 |
| 216 | LK2014 | A-54 | A-333 | 0.62679 | 0.00923 |
| 217 | LK2014 | A-275 | A-333 | 0.02980 | 0.00586 |
| 218 | LK2014 | A-509 | A-333 | 0.26932 | 0.02592 |
| 219 | LK2014 | A-600 | A-333 | 0.24912 | 0.01015 |
| 220 | LK2014 | A-04 | A-333 | 0.03310 | 0.00353 |
| 221 | LK2014 | A-092 | A-333 | 0.02802 | 0.00386 |
| 222 | LK2014 | A-244 | A-333 | 0.26842 | 0.01197 |
| 223 | LK2014 | A-342 | A-333 | 0.47283 | 0.02339 |
| 224 | LK2014 | A-389 | A-333 | 0.16034 | 0.01393 |
| 225 | LK2014 | A-547 | A-333 | 0.87678 | 0.00855 |
| 226 | LK2014 | A-196 | A-333 | 0.67199 | 0.03224 |
| 227 | LK2014 | A-360 | A-333 | 0.10576 | 0.01197 |
| 228 | LK2014 | A-465 | A-333 | 0.33244 | 0.03117 |
| 229 | LK2014 | A-161 | A-333 | 0.15897 | 0.01892 |
| 230 | LK2014 | A-200 | A-333 | 0.48207 | 0.02528 |
| 231 | LK2014 | A-325 | A-333 | 0.08018 | 0.01052 |
| 232 | LK2014 | A-1.2 | A-104.2 | 0.92601 | 0.00801 |
| 233 | LK2014 | A-293 | A-104.2 | 0.53446 | 0.01285 |
| 234 | LK2014 | A-322 | A-104.2 | 0.51358 | 0.00835 |
| 235 | LK2014 | A-421 | A-104.2 | 0.48997 | 0.01454 |
| 236 | LK2014 | A-498 | A-104.2 | 0.94103 | 0.00464 |
| 237 | LK2014 | A-54 | A-104.2 | 0.28768 | 0.00785 |

| | | | | | |
|---|---|---|---|---|---|
| 238 | LK2014 | A-275 | A-104.2 | 0.88059 | 0.00668 |
| 239 | LK2014 | A-509 | A-104.2 | 0.06154 | 0.00750 |
| 240 | LK2014 | A-600 | A-104.2 | 0.40695 | 0.00763 |
| 241 | LK2014 | A-04 | A-104.2 | 0.10624 | 0.00481 |
| 242 | LK2014 | A-092 | A-104.2 | 0.27047 | 0.00800 |
| 243 | LK2014 | A-244 | A-104.2 | 0.48675 | 0.00807 |
| 244 | LK2014 | A-342 | A-104.2 | 0.85622 | 0.00939 |
| 245 | LK2014 | A-389 | A-104.2 | 0.27157 | 0.01211 |
| 246 | LK2014 | A-547 | A-104.2 | 0.71645 | 0.00947 |
| 247 | LK2014 | A-196 | A-104.2 | 0.87486 | 0.01348 |
| 248 | LK2014 | A-360 | A-104.2 | 0.20163 | 0.00887 |
| 249 | LK2014 | A-465 | A-104.2 | 0.72755 | 0.02218 |
| 250 | LK2014 | A-161 | A-104.2 | 0.66896 | 0.01371 |
| 251 | LK2014 | A-200 | A-104.2 | 0.11858 | 0.01158 |
| 252 | LK2014 | A-325 | A-104.2 | 0.23968 | 0.01199 |
| 253 | LK2014 | A-333 | A-104.2 | 0.00619 | 0.00178 |
| 254 | LK2014 | A-1.2 | A-122 | 0.29460 | 0.01185 |
| 255 | LK2014 | A-293 | A-122 | 0.09035 | 0.00681 |
| 256 | LK2014 | A-322 | A-122 | 0.00000 | 0.00000 |
| 257 | LK2014 | A-421 | A-122 | 0.13486 | 0.00857 |
| 258 | LK2014 | A-498 | A-122 | 0.81926 | 0.00655 |
| 259 | LK2014 | A-54 | A-122 | 0.41970 | 0.00748 |
| 260 | LK2014 | A-275 | A-122 | 0.00000 | 0.00000 |
| 261 | LK2014 | A-509 | A-122 | 0.12599 | 0.01366 |
| 262 | LK2014 | A-600 | A-122 | 0.79156 | 0.00466 |
| 263 | LK2014 | A-04 | A-122 | 0.84867 | 0.00394 |
| 264 | LK2014 | A-092 | A-122 | 0.14140 | 0.00604 |
| 265 | LK2014 | A-244 | A-122 | 0.94883 | 0.00250 |
| 266 | LK2014 | A-342 | A-122 | 0.58082 | 0.01320 |
| 267 | LK2014 | A-389 | A-122 | 0.05970 | 0.00382 |
| 268 | LK2014 | A-547 | A-122 | 0.13099 | 0.00562 |
| 269 | LK2014 | A-196 | A-122 | 0.11061 | 0.01210 |
| 270 | LK2014 | A-360 | A-122 | 0.22026 | 0.00837 |
| 271 | LK2014 | A-465 | A-122 | 0.42757 | 0.02248 |
| 272 | LK2014 | A-161 | A-122 | 0.23053 | 0.01224 |
| 273 | LK2014 | A-200 | A-122 | 0.00546 | 0.00157 |
| 274 | LK2014 | A-325 | A-122 | 0.52227 | 0.01317 |
| 275 | LK2014 | A-333 | A-122 | 0.52791 | 0.01780 |
| 276 | LK2014 | A-104.2 | A-122 | 0.30635 | 0.01155 |
| 277 | LK2014 | A-1.2 | A-175 | 0.00479 | 0.00108 |
| 278 | LK2014 | A-293 | A-175 | 0.75305 | 0.00849 |
| 279 | LK2014 | A-322 | A-175 | 0.97773 | 0.00127 |
| 280 | LK2014 | A-421 | A-175 | 0.19430 | 0.00867 |
| 281 | LK2014 | A-498 | A-175 | 0.71146 | 0.00922 |
| 282 | LK2014 | A-54 | A-175 | 0.24007 | 0.00549 |
| 283 | LK2014 | A-275 | A-175 | 0.36391 | 0.01037 |
| 284 | LK2014 | A-509 | A-175 | 0.43367 | 0.01488 |
| 285 | LK2014 | A-600 | A-175 | 0.79156 | 0.00476 |

| | | | | | |
|---|---|---|---|---|---|
| 286 | LK2014 | A-04 | A-175 | 0.18701 | 0.00638 |
| 287 | LK2014 | A-092 | A-175 | 0.89623 | 0.00460 |
| 288 | LK2014 | A-244 | A-175 | 0.45022 | 0.00709 |
| 289 | LK2014 | A-342 | A-175 | 0.29250 | 0.01122 |
| 290 | LK2014 | A-389 | A-175 | 0.03245 | 0.00347 |
| 291 | LK2014 | A-547 | A-175 | 0.47612 | 0.01047 |
| 292 | LK2014 | A-196 | A-175 | 0.72893 | 0.01545 |
| 293 | LK2014 | A-360 | A-175 | 0.73263 | 0.00641 |
| 294 | LK2014 | A-465 | A-175 | 0.45362 | 0.02047 |
| 295 | LK2014 | A-161 | A-175 | 0.74762 | 0.01150 |
| 296 | LK2014 | A-200 | A-175 | 0.62471 | 0.01284 |
| 297 | LK2014 | A-325 | A-175 | 0.05698 | 0.00510 |
| 298 | LK2014 | A-333 | A-175 | 0.38941 | 0.01384 |
| 299 | LK2014 | A-104.2 | A-175 | 0.76041 | 0.00777 |
| 300 | LK2014 | A-122 | A-175 | 0.81426 | 0.00655 |
| 301 | LK2014 | A-1.2 | A-300 | 0.22006 | 0.02982 |
| 302 | LK2014 | A-293 | A-300 | 0.83006 | 0.01877 |
| 303 | LK2014 | A-322 | A-300 | 0.53731 | 0.01731 |
| 304 | LK2014 | A-421 | A-300 | 0.98251 | 0.00379 |
| 305 | LK2014 | A-498 | A-300 | 0.42144 | 0.02223 |
| 306 | LK2014 | A-54 | A-300 | 0.93112 | 0.00514 |
| 307 | LK2014 | A-275 | A-300 | 0.39968 | 0.02641 |
| 308 | LK2014 | A-509 | A-300 | 0.21424 | 0.02757 |
| 309 | LK2014 | A-600 | A-300 | 0.32110 | 0.01484 |
| 310 | LK2014 | A-04 | A-300 | 0.45577 | 0.01428 |
| 311 | LK2014 | A-092 | A-300 | 0.25009 | 0.01183 |
| 312 | LK2014 | A-244 | A-300 | 0.32571 | 0.01456 |
| 313 | LK2014 | A-342 | A-300 | 0.37746 | 0.02785 |
| 314 | LK2014 | A-389 | A-300 | 0.08198 | 0.01098 |
| 315 | LK2014 | A-547 | A-300 | 0.04288 | 0.00795 |
| 316 | LK2014 | A-196 | A-300 | 0.36984 | 0.04045 |
| 317 | LK2014 | A-360 | A-300 | 0.11479 | 0.01405 |
| 318 | LK2014 | A-465 | A-300 | 0.03261 | 0.01103 |
| 319 | LK2014 | A-161 | A-300 | 0.46312 | 0.02801 |
| 320 | LK2014 | A-200 | A-300 | 0.87985 | 0.01600 |
| 321 | LK2014 | A-325 | A-300 | 0.32386 | 0.02766 |
| 322 | LK2014 | A-333 | A-300 | 0.32727 | 0.03436 |
| 323 | LK2014 | A-104.2 | A-300 | 0.15879 | 0.02029 |
| 324 | LK2014 | A-122 | A-300 | 0.00875 | 0.00288 |
| 325 | LK2014 | A-175 | A-300 | 0.66519 | 0.01740 |
| 326 | LK2014 | A-1.2 | A-733 | 0.90081 | 0.00621 |
| 327 | LK2014 | A-293 | A-733 | 1.00000 | 0.00000 |
| 328 | LK2014 | A-322 | A-733 | 0.69581 | 0.00428 |
| 329 | LK2014 | A-421 | A-733 | 0.39454 | 0.00995 |
| 330 | LK2014 | A-498 | A-733 | 0.61108 | 0.00774 |
| 331 | LK2014 | A-54 | A-733 | 0.46382 | 0.00538 |
| 332 | LK2014 | A-275 | A-733 | 0.80959 | 0.00542 |
| 333 | LK2014 | A-509 | A-733 | 0.49205 | 0.01418 |

| | | | | | |
|---|---|---|---|---|---|
| 334 | LK2014 | A-600 | A-733 | 0.50798 | 0.00496 |
| 335 | LK2014 | A-04 | A-733 | 0.18804 | 0.00462 |
| 336 | LK2014 | A-092 | A-733 | 1.00000 | 0.00000 |
| 337 | LK2014 | A-244 | A-733 | 0.21621 | 0.00502 |
| 338 | LK2014 | A-342 | A-733 | 0.03351 | 0.00302 |
| 339 | LK2014 | A-389 | A-733 | 0.88854 | 0.00393 |
| 340 | LK2014 | A-547 | A-733 | 1.00000 | 0.00000 |
| 341 | LK2014 | A-196 | A-733 | 0.76109 | 0.01322 |
| 342 | LK2014 | A-360 | A-733 | 0.56240 | 0.00844 |
| 343 | LK2014 | A-465 | A-733 | 0.36539 | 0.01676 |
| 344 | LK2014 | A-161 | A-733 | 0.30929 | 0.01168 |
| 345 | LK2014 | A-200 | A-733 | 0.15403 | 0.00946 |
| 346 | LK2014 | A-325 | A-733 | 0.51829 | 0.01104 |
| 347 | LK2014 | A-333 | A-733 | 0.99315 | 0.00136 |
| 348 | LK2014 | A-104.2 | A-733 | 0.36162 | 0.00915 |
| 349 | LK2014 | A-122 | A-733 | 0.93027 | 0.00304 |
| 350 | LK2014 | A-175 | A-733 | 0.69963 | 0.00633 |
| 351 | LK2014 | A-300 | A-733 | 0.22576 | 0.01588 |
| 352 | LK2014 | A-1.2 | A-0 | 0.10775 | 0.00886 |
| 353 | LK2014 | A-293 | A-0 | 1.00000 | 0.00000 |
| 354 | LK2014 | A-322 | A-0 | 0.42115 | 0.00413 |
| 355 | LK2014 | A-421 | A-0 | 0.47227 | 0.01174 |
| 356 | LK2014 | A-498 | A-0 | 0.04545 | 0.00416 |
| 357 | LK2014 | A-54 | A-0 | 0.48713 | 0.00617 |
| 358 | LK2014 | A-275 | A-0 | 0.52143 | 0.00969 |
| 359 | LK2014 | A-509 | A-0 | 0.89865 | 0.00934 |
| 360 | LK2014 | A-600 | A-0 | 0.44732 | 0.00643 |
| 361 | LK2014 | A-04 | A-0 | 0.29967 | 0.00555 |
| 362 | LK2014 | A-092 | A-0 | 0.80611 | 0.00557 |
| 363 | LK2014 | A-244 | A-0 | 0.74649 | 0.00617 |
| 364 | LK2014 | A-342 | A-0 | 0.40514 | 0.01299 |
| 365 | LK2014 | A-389 | A-0 | 0.16419 | 0.00893 |
| 366 | LK2014 | A-547 | A-0 | 0.35820 | 0.00960 |
| 367 | LK2014 | A-196 | A-0 | 0.86493 | 0.01083 |
| 368 | LK2014 | A-360 | A-0 | 0.03144 | 0.00295 |
| 369 | LK2014 | A-465 | A-0 | 0.35962 | 0.02135 |
| 370 | LK2014 | A-161 | A-0 | 0.42732 | 0.01336 |
| 371 | LK2014 | A-200 | A-0 | 0.90429 | 0.00796 |
| 372 | LK2014 | A-325 | A-0 | 0.45342 | 0.01243 |
| 373 | LK2014 | A-333 | A-0 | 0.21011 | 0.01316 |
| 374 | LK2014 | A-104.2 | A-0 | 0.49924 | 0.01067 |
| 375 | LK2014 | A-122 | A-0 | 0.81378 | 0.00652 |
| 376 | LK2014 | A-175 | A-0 | 0.21323 | 0.00739 |
| 377 | LK2014 | A-300 | A-0 | 0.68518 | 0.01999 |
| 378 | LK2014 | A-733 | A-0 | 1.00000 | 0.00000 |
| 379 | LK2014 | A-1.2 | A-1.1 | 0.00000 | 0.00000 |
| 380 | LK2014 | A-293 | A-1.1 | 0.42911 | 0.01390 |
| 381 | LK2014 | A-322 | A-1.1 | 0.06803 | 0.00386 |

| | | | | | |
|---|---|---|---|---|---|
| 382 | LK2014 | A-421 | A-1.1 | 0.20513 | 0.01134 |
| 383 | LK2014 | A-498 | A-1.1 | 0.26170 | 0.01147 |
| 384 | LK2014 | A-54 | A-1.1 | 0.42070 | 0.00808 |
| 385 | LK2014 | A-275 | A-1.1 | 0.03967 | 0.00456 |
| 386 | LK2014 | A-509 | A-1.1 | 0.98397 | 0.00289 |
| 387 | LK2014 | A-600 | A-1.1 | 0.10597 | 0.00476 |
| 388 | LK2014 | A-04 | A-1.1 | 0.68109 | 0.00783 |
| 389 | LK2014 | A-092 | A-1.1 | 0.19149 | 0.00609 |
| 390 | LK2014 | A-244 | A-1.1 | 0.49487 | 0.00875 |
| 391 | LK2014 | A-342 | A-1.1 | 0.42087 | 0.01394 |
| 392 | LK2014 | A-389 | A-1.1 | 0.89285 | 0.00536 |
| 393 | LK2014 | A-547 | A-1.1 | 0.62254 | 0.01085 |
| 394 | LK2014 | A-196 | A-1.1 | 0.06798 | 0.00874 |
| 395 | LK2014 | A-360 | A-1.1 | 0.67193 | 0.00999 |
| 396 | LK2014 | A-465 | A-1.1 | 0.45415 | 0.02311 |
| 397 | LK2014 | A-161 | A-1.1 | 0.59014 | 0.01466 |
| 398 | LK2014 | A-200 | A-1.1 | 0.00001 | 0.00001 |
| 399 | LK2014 | A-325 | A-1.1 | 0.45800 | 0.01428 |
| 400 | LK2014 | A-333 | A-1.1 | 0.99216 | 0.00156 |
| 401 | LK2014 | A-104.2 | A-1.1 | 0.41152 | 0.01213 |
| 402 | LK2014 | A-122 | A-1.1 | 0.09384 | 0.00651 |
| 403 | LK2014 | A-175 | A-1.1 | 0.14825 | 0.00723 |
| 404 | LK2014 | A-300 | A-1.1 | 0.65015 | 0.02405 |
| 405 | LK2014 | A-733 | A-1.1 | 0.16433 | 0.00691 |
| 406 | LK2014 | A-0 | A-1.1 | 0.63801 | 0.00878 |
| 407 | LK2014 | A-1.2 | A-450 | 0.65958 | 0.01940 |
| 408 | LK2014 | A-293 | A-450 | 0.03703 | 0.00534 |
| 409 | LK2014 | A-322 | A-450 | 0.77788 | 0.00797 |
| 410 | LK2014 | A-421 | A-450 | 0.21515 | 0.01313 |
| 411 | LK2014 | A-498 | A-450 | 0.32897 | 0.01278 |
| 412 | LK2014 | A-54 | A-450 | 0.63935 | 0.00997 |
| 413 | LK2014 | A-275 | A-450 | 0.66103 | 0.01255 |
| 414 | LK2014 | A-509 | A-450 | 0.54214 | 0.02536 |
| 415 | LK2014 | A-600 | A-450 | 0.37238 | 0.00940 |
| 416 | LK2014 | A-04 | A-450 | 0.70401 | 0.00743 |
| 417 | LK2014 | A-092 | A-450 | 0.42966 | 0.00934 |
| 418 | LK2014 | A-244 | A-450 | 0.05376 | 0.00397 |
| 419 | LK2014 | A-342 | A-450 | 0.00205 | 0.00076 |
| 420 | LK2014 | A-389 | A-450 | 0.66855 | 0.01325 |
| 421 | LK2014 | A-547 | A-450 | 0.43695 | 0.01245 |
| 422 | LK2014 | A-196 | A-450 | 0.79425 | 0.01992 |
| 423 | LK2014 | A-360 | A-450 | 0.21444 | 0.01183 |
| 424 | LK2014 | A-465 | A-450 | 0.52474 | 0.02797 |
| 425 | LK2014 | A-161 | A-450 | 0.06788 | 0.00837 |
| 426 | LK2014 | A-200 | A-450 | 0.04467 | 0.00913 |
| 427 | LK2014 | A-325 | A-450 | 0.81767 | 0.01064 |
| 428 | LK2014 | A-333 | A-450 | 0.40541 | 0.02206 |
| 429 | LK2014 | A-104.2 | A-450 | 0.68443 | 0.01475 |

| | | | | | |
|---|---|---|---|---|---|
| 430 | LK2014 | A-122 | A-450 | 0.12091 | 0.00711 |
| 431 | LK2014 | A-175 | A-450 | 0.96901 | 0.00319 |
| 432 | LK2014 | A-300 | A-450 | 0.23555 | 0.02333 |
| 433 | LK2014 | A-733 | A-450 | 0.72595 | 0.00866 |
| 434 | LK2014 | A-0 | A-450 | 0.75763 | 0.01174 |
| 435 | LK2014 | A-1.1 | A-450 | 0.34833 | 0.01279 |
| 436 | LK2014 | A-1.2 | A-60 | 0.17292 | 0.01888 |
| 437 | LK2014 | A-293 | A-60 | 0.43880 | 0.02825 |
| 438 | LK2014 | A-322 | A-60 | 0.02925 | 0.00362 |
| 439 | LK2014 | A-421 | A-60 | 0.10399 | 0.01051 |
| 440 | LK2014 | A-498 | A-60 | 0.38511 | 0.01560 |
| 441 | LK2014 | A-54 | A-60 | 0.09798 | 0.00658 |
| 442 | LK2014 | A-275 | A-60 | 0.19328 | 0.01554 |
| 443 | LK2014 | A-509 | A-60 | 0.46854 | 0.03132 |
| 444 | LK2014 | A-600 | A-60 | 0.83357 | 0.00627 |
| 445 | LK2014 | A-04 | A-60 | 0.82468 | 0.00574 |
| 446 | LK2014 | A-092 | A-60 | 0.45081 | 0.01435 |
| 447 | LK2014 | A-244 | A-60 | 0.76413 | 0.00864 |
| 448 | LK2014 | A-342 | A-60 | 0.75785 | 0.01620 |
| 449 | LK2014 | A-389 | A-60 | 0.30498 | 0.01583 |
| 450 | LK2014 | A-547 | A-60 | 0.31195 | 0.01302 |
| 451 | LK2014 | A-196 | A-60 | 0.27360 | 0.02923 |
| 452 | LK2014 | A-360 | A-60 | 0.08054 | 0.00718 |
| 453 | LK2014 | A-465 | A-60 | 0.55413 | 0.03083 |
| 454 | LK2014 | A-161 | A-60 | 0.04843 | 0.00881 |
| 455 | LK2014 | A-200 | A-60 | 0.02895 | 0.00581 |
| 456 | LK2014 | A-325 | A-60 | 0.10062 | 0.01085 |
| 457 | LK2014 | A-333 | A-60 | 0.91854 | 0.01034 |
| 458 | LK2014 | A-104.2 | A-60 | 0.68156 | 0.01560 |
| 459 | LK2014 | A-122 | A-60 | 0.00270 | 0.00084 |
| 460 | LK2014 | A-175 | A-60 | 0.10472 | 0.00832 |
| 461 | LK2014 | A-300 | A-60 | 0.04545 | 0.00941 |
| 462 | LK2014 | A-733 | A-60 | 0.00375 | 0.00122 |
| 463 | LK2014 | A-0 | A-60 | 0.21987 | 0.01342 |
| 464 | LK2014 | A-1.1 | A-60 | 0.01003 | 0.00230 |
| 465 | LK2014 | A-450 | A-60 | 0.66203 | 0.02017 |
| 466 | LK2014 | A-1.2 | A-662 | 0.54068 | 0.01197 |
| 467 | LK2014 | A-293 | A-662 | 0.46434 | 0.00969 |
| 468 | LK2014 | A-322 | A-662 | 0.00000 | 0.00000 |
| 469 | LK2014 | A-421 | A-662 | 0.05280 | 0.00366 |
| 470 | LK2014 | A-498 | A-662 | 0.67367 | 0.00710 |
| 471 | LK2014 | A-54 | A-662 | 0.27208 | 0.00436 |
| 472 | LK2014 | A-275 | A-662 | 0.00000 | 0.00000 |
| 473 | LK2014 | A-509 | A-662 | 0.13363 | 0.01008 |
| 474 | LK2014 | A-600 | A-662 | 0.57599 | 0.00394 |
| 475 | LK2014 | A-04 | A-662 | 0.64244 | 0.00378 |
| 476 | LK2014 | A-092 | A-662 | 0.28667 | 0.00611 |
| 477 | LK2014 | A-244 | A-662 | 0.26312 | 0.00556 |

| | | | | | |
|---|---|---|---|---|---|
| 478 | LK2014 | A-342 | A-662 | 0.30833 | 0.00844 |
| 479 | LK2014 | A-389 | A-662 | 0.93985 | 0.00247 |
| 480 | LK2014 | A-547 | A-662 | 0.40777 | 0.00693 |
| 481 | LK2014 | A-196 | A-662 | 0.71689 | 0.01407 |
| 482 | LK2014 | A-360 | A-662 | 0.62251 | 0.00748 |
| 483 | LK2014 | A-465 | A-662 | 0.09771 | 0.00804 |
| 484 | LK2014 | A-161 | A-662 | 0.92454 | 0.00496 |
| 485 | LK2014 | A-200 | A-662 | 0.00025 | 0.00018 |
| 486 | LK2014 | A-325 | A-662 | 0.62745 | 0.00819 |
| 487 | LK2014 | A-333 | A-662 | 0.06263 | 0.00466 |
| 488 | LK2014 | A-104.2 | A-662 | 0.82131 | 0.00509 |
| 489 | LK2014 | A-122 | A-662 | 0.00000 | 0.00000 |
| 490 | LK2014 | A-175 | A-662 | 0.97879 | 0.00112 |
| 491 | LK2014 | A-300 | A-662 | 0.54293 | 0.01619 |
| 492 | LK2014 | A-733 | A-662 | 0.63823 | 0.00455 |
| 493 | LK2014 | A-0 | A-662 | 0.37871 | 0.00469 |
| 494 | LK2014 | A-1.1 | A-662 | 0.07509 | 0.00401 |
| 495 | LK2014 | A-450 | A-662 | 0.61876 | 0.00943 |
| 496 | LK2014 | A-60 | A-662 | 0.04691 | 0.00582 |
| 497 | LK2014 | A-1.2 | A-104.1 | 0.72782 | 0.00972 |
| 498 | LK2014 | A-293 | A-104.1 | 0.03897 | 0.00238 |
| 499 | LK2014 | A-322 | A-104.1 | 1.00000 | 0.00000 |
| 500 | LK2014 | A-421 | A-104.1 | 0.58506 | 0.00655 |
| 501 | LK2014 | A-498 | A-104.1 | 0.18695 | 0.00521 |
| 502 | LK2014 | A-54 | A-104.1 | 0.39720 | 0.00367 |
| 503 | LK2014 | A-275 | A-104.1 | 1.00000 | 0.00000 |
| 504 | LK2014 | A-509 | A-104.1 | 1.00000 | 0.00000 |
| 505 | LK2014 | A-600 | A-104.1 | 0.39185 | 0.00359 |
| 506 | LK2014 | A-04 | A-104.1 | 0.26635 | 0.00221 |
| 507 | LK2014 | A-092 | A-104.1 | 1.00000 | 0.00000 |
| 508 | LK2014 | A-244 | A-104.1 | 1.00000 | 0.00000 |
| 509 | LK2014 | A-342 | A-104.1 | 0.04178 | 0.00252 |
| 510 | LK2014 | A-389 | A-104.1 | 0.57439 | 0.00584 |
| 511 | LK2014 | A-547 | A-104.1 | 0.61534 | 0.00600 |
| 512 | LK2014 | A-196 | A-104.1 | 0.68165 | 0.01526 |
| 513 | LK2014 | A-360 | A-104.1 | 1.00000 | 0.00000 |
| 514 | LK2014 | A-465 | A-104.1 | 0.14289 | 0.00752 |
| 515 | LK2014 | A-161 | A-104.1 | 0.06491 | 0.00380 |
| 516 | LK2014 | A-200 | A-104.1 | 1.00000 | 0.00000 |
| 517 | LK2014 | A-325 | A-104.1 | 1.00000 | 0.00000 |
| 518 | LK2014 | A-333 | A-104.1 | 0.75933 | 0.00867 |
| 519 | LK2014 | A-104.2 | A-104.1 | 0.60325 | 0.00736 |
| 520 | LK2014 | A-122 | A-104.1 | 1.00000 | 0.00000 |
| 521 | LK2014 | A-175 | A-104.1 | 0.48331 | 0.00506 |
| 522 | LK2014 | A-300 | A-104.1 | 1.00000 | 0.00000 |
| 523 | LK2014 | A-733 | A-104.1 | 1.00000 | 0.00000 |
| 524 | LK2014 | A-0 | A-104.1 | 1.00000 | 0.00000 |
| 525 | LK2014 | A-1.1 | A-104.1 | 0.16468 | 0.00365 |

| | | | | | |
|---|---|---|---|---|---|
| 526 | LK2014 | A-450 | A-104.1 | 0.08371 | 0.00363 |
| 527 | LK2014 | A-60 | A-104.1 | 0.16474 | 0.00638 |
| 528 | LK2014 | A-662 | A-104.1 | 1.00000 | 0.00000 |
| 529 | LK2014 | A-1.2 | A-149 | 0.00726 | 0.00209 |
| 530 | LK2014 | A-293 | A-149 | 0.60457 | 0.01555 |
| 531 | LK2014 | A-322 | A-149 | 0.54717 | 0.00728 |
| 532 | LK2014 | A-421 | A-149 | 0.05063 | 0.00557 |
| 533 | LK2014 | A-498 | A-149 | 0.40866 | 0.01257 |
| 534 | LK2014 | A-54 | A-149 | 0.11757 | 0.00463 |
| 535 | LK2014 | A-275 | A-149 | 0.23911 | 0.01033 |
| 536 | LK2014 | A-509 | A-149 | 0.15791 | 0.01358 |
| 537 | LK2014 | A-600 | A-149 | 0.38946 | 0.00792 |
| 538 | LK2014 | A-04 | A-149 | 0.36644 | 0.00740 |
| 539 | LK2014 | A-092 | A-149 | 0.44736 | 0.01027 |
| 540 | LK2014 | A-244 | A-149 | 0.65069 | 0.00700 |
| 541 | LK2014 | A-342 | A-149 | 0.99596 | 0.00096 |
| 542 | LK2014 | A-389 | A-149 | 0.80404 | 0.00754 |
| 543 | LK2014 | A-547 | A-149 | 0.56501 | 0.01057 |
| 544 | LK2014 | A-196 | A-149 | 0.88228 | 0.01347 |
| 545 | LK2014 | A-360 | A-149 | 0.64277 | 0.01147 |
| 546 | LK2014 | A-465 | A-149 | 0.06058 | 0.00960 |
| 547 | LK2014 | A-161 | A-149 | 0.10028 | 0.01005 |
| 548 | LK2014 | A-200 | A-149 | 0.65062 | 0.01515 |
| 549 | LK2014 | A-325 | A-149 | 0.00234 | 0.00112 |
| 550 | LK2014 | A-333 | A-149 | 0.36605 | 0.02020 |
| 551 | LK2014 | A-104.2 | A-149 | 0.72120 | 0.01024 |
| 552 | LK2014 | A-122 | A-149 | 0.10892 | 0.00706 |
| 553 | LK2014 | A-175 | A-149 | 0.05249 | 0.00385 |
| 554 | LK2014 | A-300 | A-149 | 0.63035 | 0.02750 |
| 555 | LK2014 | A-733 | A-149 | 0.26841 | 0.00880 |
| 556 | LK2014 | A-0 | A-149 | 0.81096 | 0.00722 |
| 557 | LK2014 | A-1.1 | A-149 | 0.39319 | 0.01080 |
| 558 | LK2014 | A-450 | A-149 | 0.34618 | 0.01431 |
| 559 | LK2014 | A-60 | A-149 | 0.70996 | 0.01557 |
| 560 | LK2014 | A-662 | A-149 | 0.44833 | 0.00829 |
| 561 | LK2014 | A-104.1 | A-149 | 1.00000 | 0.00000 |
| 562 | LK2014 | A-1.2 | A-224 | 0.57962 | 0.02804 |
| 563 | LK2014 | A-293 | A-224 | 0.15092 | 0.01626 |
| 564 | LK2014 | A-322 | A-224 | 0.09499 | 0.00780 |
| 565 | LK2014 | A-421 | A-224 | 0.58968 | 0.01959 |
| 566 | LK2014 | A-498 | A-224 | 0.05875 | 0.00819 |
| 567 | LK2014 | A-54 | A-224 | 0.04964 | 0.00438 |
| 568 | LK2014 | A-275 | A-224 | 0.35722 | 0.01854 |
| 569 | LK2014 | A-509 | A-224 | 0.06046 | 0.01457 |
| 570 | LK2014 | A-600 | A-224 | 0.86011 | 0.00594 |
| 571 | LK2014 | A-04 | A-224 | 0.18778 | 0.00727 |
| 572 | LK2014 | A-092 | A-224 | 0.35020 | 0.01354 |
| 573 | LK2014 | A-244 | A-224 | 0.85413 | 0.00685 |

| | | | | | |
|---|---|---|---|---|---|
| 574 | LK2014 | A-342 | A-224 | 0.39567 | 0.02257 |
| 575 | LK2014 | A-389 | A-224 | 0.23285 | 0.01594 |
| 576 | LK2014 | A-547 | A-224 | 0.42797 | 0.01530 |
| 577 | LK2014 | A-196 | A-224 | 0.46697 | 0.03477 |
| 578 | LK2014 | A-360 | A-224 | 0.09530 | 0.00903 |
| 579 | LK2014 | A-465 | A-224 | 0.72845 | 0.02919 |
| 580 | LK2014 | A-161 | A-224 | 0.25654 | 0.02281 |
| 581 | LK2014 | A-200 | A-224 | 0.55811 | 0.02263 |
| 582 | LK2014 | A-325 | A-224 | 0.00094 | 0.00081 |
| 583 | LK2014 | A-333 | A-224 | 0.34669 | 0.02546 |
| 584 | LK2014 | A-104.2 | A-224 | 0.61842 | 0.01624 |
| 585 | LK2014 | A-122 | A-224 | 0.54531 | 0.01723 |
| 586 | LK2014 | A-175 | A-224 | 0.43322 | 0.01274 |
| 587 | LK2014 | A-300 | A-224 | 0.54718 | 0.03443 |
| 588 | LK2014 | A-733 | A-224 | 0.84678 | 0.00787 |
| 589 | LK2014 | A-0 | A-224 | 0.96763 | 0.00428 |
| 590 | LK2014 | A-1.1 | A-224 | 0.81387 | 0.01097 |
| 591 | LK2014 | A-450 | A-224 | 0.32985 | 0.02052 |
| 592 | LK2014 | A-60 | A-224 | 0.19200 | 0.02194 |
| 593 | LK2014 | A-662 | A-224 | 0.14532 | 0.00893 |
| 594 | LK2014 | A-104.1 | A-224 | 0.05823 | 0.00393 |
| 595 | LK2014 | A-149 | A-224 | 0.26039 | 0.01996 |
| 596 | LK2014 | A-1.2 | A-323 | 0.62711 | 0.03104 |
| 597 | LK2014 | A-293 | A-323 | 0.42776 | 0.02717 |
| 598 | LK2014 | A-322 | A-323 | 0.72755 | 0.01370 |
| 599 | LK2014 | A-421 | A-323 | 0.62159 | 0.02498 |
| 600 | LK2014 | A-498 | A-323 | 0.33550 | 0.01932 |
| 601 | LK2014 | A-54 | A-323 | 0.38478 | 0.01378 |
| 602 | LK2014 | A-275 | A-323 | 0.42904 | 0.02263 |
| 603 | LK2014 | A-509 | A-323 | 0.28868 | 0.03302 |
| 604 | LK2014 | A-600 | A-323 | 0.51807 | 0.01374 |
| 605 | LK2014 | A-04 | A-323 | 0.92743 | 0.00547 |
| 606 | LK2014 | A-092 | A-323 | 0.97928 | 0.00337 |
| 607 | LK2014 | A-244 | A-323 | 0.15448 | 0.01012 |
| 608 | LK2014 | A-342 | A-323 | 0.51534 | 0.02867 |
| 609 | LK2014 | A-389 | A-323 | 0.54713 | 0.02095 |
| 610 | LK2014 | A-547 | A-323 | 0.30210 | 0.01961 |
| 611 | LK2014 | A-196 | A-323 | 0.04219 | 0.01273 |
| 612 | LK2014 | A-360 | A-323 | 0.55237 | 0.02309 |
| 613 | LK2014 | A-465 | A-323 | 0.95050 | 0.01431 |
| 614 | LK2014 | A-161 | A-323 | 0.54916 | 0.02722 |
| 615 | LK2014 | A-200 | A-323 | 0.98190 | 0.00667 |
| 616 | LK2014 | A-325 | A-323 | 0.22649 | 0.02290 |
| 617 | LK2014 | A-333 | A-323 | 0.68756 | 0.03265 |
| 618 | LK2014 | A-104.2 | A-323 | 0.67533 | 0.02063 |
| 619 | LK2014 | A-122 | A-323 | 0.54110 | 0.02225 |
| 620 | LK2014 | A-175 | A-323 | 0.34008 | 0.01583 |
| 621 | LK2014 | A-300 | A-323 | 0.89814 | 0.02111 |

| | | | | | |
|---|---|---|---|---|---|
| 622 | LK2014 | A-733 | A-323 | 0.11397 | 0.00851 |
| 623 | LK2014 | A-0 | A-323 | 0.23107 | 0.01638 |
| 624 | LK2014 | A-1.1 | A-323 | 0.54511 | 0.02206 |
| 625 | LK2014 | A-450 | A-323 | 0.33455 | 0.02569 |
| 626 | LK2014 | A-60 | A-323 | 0.22721 | 0.02617 |
| 627 | LK2014 | A-662 | A-323 | 0.66099 | 0.01298 |
| 628 | LK2014 | A-104.1 | A-323 | 0.37315 | 0.01056 |
| 629 | LK2014 | A-149 | A-323 | 0.88648 | 0.01273 |
| 630 | LK2014 | A-224 | A-323 | 0.50677 | 0.03238 |
| 631 | LK2014 | A-1.2 | A-880 | 0.08207 | 0.00962 |
| 632 | LK2014 | A-293 | A-880 | 0.95737 | 0.00490 |
| 633 | LK2014 | A-322 | A-880 | 0.37600 | 0.01023 |
| 634 | LK2014 | A-421 | A-880 | 0.00814 | 0.00199 |
| 635 | LK2014 | A-498 | A-880 | 0.54131 | 0.01254 |
| 636 | LK2014 | A-54 | A-880 | 0.54726 | 0.00819 |
| 637 | LK2014 | A-275 | A-880 | 0.40436 | 0.01281 |
| 638 | LK2014 | A-509 | A-880 | 0.41365 | 0.02081 |
| 639 | LK2014 | A-600 | A-880 | 0.33503 | 0.00810 |
| 640 | LK2014 | A-04 | A-880 | 0.07838 | 0.00541 |
| 641 | LK2014 | A-092 | A-880 | 0.83863 | 0.00516 |
| 642 | LK2014 | A-244 | A-880 | 0.13000 | 0.00655 |
| 643 | LK2014 | A-342 | A-880 | 0.41510 | 0.01482 |
| 644 | LK2014 | A-389 | A-880 | 0.39500 | 0.01240 |
| 645 | LK2014 | A-547 | A-880 | 0.81046 | 0.00820 |
| 646 | LK2014 | A-196 | A-880 | 0.95474 | 0.00655 |
| 647 | LK2014 | A-360 | A-880 | 0.47181 | 0.01326 |
| 648 | LK2014 | A-465 | A-880 | 0.39430 | 0.02359 |
| 649 | LK2014 | A-161 | A-880 | 0.32863 | 0.01763 |
| 650 | LK2014 | A-200 | A-880 | 0.81367 | 0.01355 |
| 651 | LK2014 | A-325 | A-880 | 0.63254 | 0.01421 |
| 652 | LK2014 | A-333 | A-880 | 0.54736 | 0.02055 |
| 653 | LK2014 | A-104.2 | A-880 | 0.51778 | 0.01450 |
| 654 | LK2014 | A-122 | A-880 | 0.82223 | 0.00995 |
| 655 | LK2014 | A-175 | A-880 | 0.69410 | 0.00910 |
| 656 | LK2014 | A-300 | A-880 | 0.92333 | 0.01124 |
| 657 | LK2014 | A-733 | A-880 | 0.21338 | 0.00999 |
| 658 | LK2014 | A-0 | A-880 | 0.26485 | 0.01045 |
| 659 | LK2014 | A-1.1 | A-880 | 0.84371 | 0.00758 |
| 660 | LK2014 | A-450 | A-880 | 0.23492 | 0.01277 |
| 661 | LK2014 | A-60 | A-880 | 0.28629 | 0.01763 |
| 662 | LK2014 | A-662 | A-880 | 0.45053 | 0.00958 |
| 663 | LK2014 | A-104.1 | A-880 | 0.40115 | 0.00710 |
| 664 | LK2014 | A-149 | A-880 | 0.54623 | 0.01579 |
| 665 | LK2014 | A-224 | A-880 | 0.40667 | 0.02010 |
| 666 | LK2014 | A-323 | A-880 | 0.77597 | 0.01834 |
| 667 | LK2016 | A-1.2 | A-293 | 0.04321 | 0.00695 |
| 668 | LK2016 | A-1.2 | A-322 | 0.02704 | 0.00507 |
| 669 | LK2016 | A-293 | A-322 | 0.61039 | 0.00803 |

| | | | | | |
|---|---|---|---|---|---|
| 670 | LK2016 | A-1.2 | A-421 | 0.63254 | 0.02741 |
| 671 | LK2016 | A-293 | A-421 | 0.10364 | 0.01018 |
| 672 | LK2016 | A-322 | A-421 | 0.33075 | 0.01233 |
| 673 | LK2016 | A-1.2 | A-498 | 0.75459 | 0.01958 |
| 674 | LK2016 | A-293 | A-498 | 0.65464 | 0.01092 |
| 675 | LK2016 | A-322 | A-498 | 0.33533 | 0.00936 |
| 676 | LK2016 | A-421 | A-498 | 0.62669 | 0.01569 |
| 677 | LK2016 | A-1.2 | A-54 | 0.30200 | 0.01334 |
| 678 | LK2016 | A-293 | A-54 | 0.85267 | 0.00356 |
| 679 | LK2016 | A-322 | A-54 | 0.71548 | 0.00393 |
| 680 | LK2016 | A-421 | A-54 | 0.76533 | 0.00854 |
| 681 | LK2016 | A-498 | A-54 | 0.45866 | 0.00877 |
| 682 | LK2016 | A-1.2 | A-275 | 0.16835 | 0.01817 |
| 683 | LK2016 | A-293 | A-275 | 0.80096 | 0.00767 |
| 684 | LK2016 | A-322 | A-275 | 0.00000 | 0.00000 |
| 685 | LK2016 | A-421 | A-275 | 0.13543 | 0.01124 |
| 686 | LK2016 | A-498 | A-275 | 0.21830 | 0.00997 |
| 687 | LK2016 | A-54 | A-275 | 0.74976 | 0.00523 |
| 688 | LK2016 | A-1.2 | A-509 | 0.17025 | 0.02550 |
| 689 | LK2016 | A-293 | A-509 | 0.68091 | 0.01774 |
| 690 | LK2016 | A-322 | A-509 | 0.22698 | 0.01112 |
| 691 | LK2016 | A-421 | A-509 | 0.51660 | 0.02784 |
| 692 | LK2016 | A-498 | A-509 | 0.28319 | 0.02036 |
| 693 | LK2016 | A-54 | A-509 | 0.91437 | 0.00619 |
| 694 | LK2016 | A-275 | A-509 | 0.58928 | 0.01791 |
| 695 | LK2016 | A-1.2 | A-600 | 0.53615 | 0.01222 |
| 696 | LK2016 | A-293 | A-600 | 0.50512 | 0.00606 |
| 697 | LK2016 | A-322 | A-600 | 0.75600 | 0.00352 |
| 698 | LK2016 | A-421 | A-600 | 0.89331 | 0.00475 |
| 699 | LK2016 | A-498 | A-600 | 0.16053 | 0.00670 |
| 700 | LK2016 | A-54 | A-600 | 0.17984 | 0.00397 |
| 701 | LK2016 | A-275 | A-600 | 0.90975 | 0.00287 |
| 702 | LK2016 | A-509 | A-600 | 0.35552 | 0.01236 |
| 703 | LK2016 | A-1.2 | A-04 | 0.76495 | 0.00988 |
| 704 | LK2016 | A-293 | A-04 | 0.50944 | 0.00543 |
| 705 | LK2016 | A-322 | A-04 | 0.43853 | 0.00476 |
| 706 | LK2016 | A-421 | A-04 | 0.77190 | 0.00691 |
| 707 | LK2016 | A-498 | A-04 | 0.86463 | 0.00435 |
| 708 | LK2016 | A-54 | A-04 | 0.39414 | 0.00549 |
| 709 | LK2016 | A-275 | A-04 | 0.08314 | 0.00397 |
| 710 | LK2016 | A-509 | A-04 | 0.18001 | 0.00956 |
| 711 | LK2016 | A-600 | A-04 | 0.52833 | 0.00604 |
| 712 | LK2016 | A-1.2 | A-092 | 0.16670 | 0.00740 |
| 713 | LK2016 | A-293 | A-092 | 0.78090 | 0.00333 |
| 714 | LK2016 | A-322 | A-092 | 1.00000 | 0.00000 |
| 715 | LK2016 | A-421 | A-092 | 0.16140 | 0.00531 |
| 716 | LK2016 | A-498 | A-092 | 0.95162 | 0.00164 |
| 717 | LK2016 | A-54 | A-092 | 0.45128 | 0.00340 |

| | | | | | |
|---|---|---|---|---|---|
| 718 | LK2016 | A-275 | A-092 | 0.64953 | 0.00404 |
| 719 | LK2016 | A-509 | A-092 | 0.76245 | 0.00492 |
| 720 | LK2016 | A-600 | A-092 | 0.15775 | 0.00222 |
| 721 | LK2016 | A-04 | A-092 | 0.09531 | 0.00220 |
| 722 | LK2016 | A-1.2 | A-244 | 0.65653 | 0.02319 |
| 723 | LK2016 | A-293 | A-244 | 0.60693 | 0.01025 |
| 724 | LK2016 | A-322 | A-244 | 0.38618 | 0.00786 |
| 725 | LK2016 | A-421 | A-244 | 0.51252 | 0.01866 |
| 726 | LK2016 | A-498 | A-244 | 0.48323 | 0.01339 |
| 727 | LK2016 | A-54 | A-244 | 0.00059 | 0.00044 |
| 728 | LK2016 | A-275 | A-244 | 0.56214 | 0.01056 |
| 729 | LK2016 | A-509 | A-244 | 0.83525 | 0.01449 |
| 730 | LK2016 | A-600 | A-244 | 0.11715 | 0.00503 |
| 731 | LK2016 | A-04 | A-244 | 0.64670 | 0.00646 |
| 732 | LK2016 | A-092 | A-244 | 0.14826 | 0.00317 |
| 733 | LK2016 | A-1.2 | A-342 | 0.51986 | 0.02327 |
| 734 | LK2016 | A-293 | A-342 | 0.09883 | 0.00699 |
| 735 | LK2016 | A-322 | A-342 | 0.99318 | 0.00095 |
| 736 | LK2016 | A-421 | A-342 | 0.05626 | 0.00882 |
| 737 | LK2016 | A-498 | A-342 | 0.38869 | 0.01389 |
| 738 | LK2016 | A-54 | A-342 | 0.37499 | 0.00799 |
| 739 | LK2016 | A-275 | A-342 | 0.94785 | 0.00365 |
| 740 | LK2016 | A-509 | A-342 | 0.46008 | 0.01980 |
| 741 | LK2016 | A-600 | A-342 | 0.33560 | 0.00726 |
| 742 | LK2016 | A-04 | A-342 | 0.53375 | 0.00841 |
| 743 | LK2016 | A-092 | A-342 | 0.73817 | 0.00436 |
| 744 | LK2016 | A-244 | A-342 | 0.68209 | 0.01099 |
| 745 | LK2016 | A-1.2 | A-389 | 0.96242 | 0.00716 |
| 746 | LK2016 | A-293 | A-389 | 0.58183 | 0.01117 |
| 747 | LK2016 | A-322 | A-389 | 0.75259 | 0.00639 |
| 748 | LK2016 | A-421 | A-389 | 0.55945 | 0.01704 |
| 749 | LK2016 | A-498 | A-389 | 0.28097 | 0.01338 |
| 750 | LK2016 | A-54 | A-389 | 0.95988 | 0.00191 |
| 751 | LK2016 | A-275 | A-389 | 0.86734 | 0.00639 |
| 752 | LK2016 | A-509 | A-389 | 0.96992 | 0.00432 |
| 753 | LK2016 | A-600 | A-389 | 0.75401 | 0.00496 |
| 754 | LK2016 | A-04 | A-389 | 0.18459 | 0.00661 |
| 755 | LK2016 | A-092 | A-389 | 1.00000 | 0.00000 |
| 756 | LK2016 | A-244 | A-389 | 0.88102 | 0.00602 |
| 757 | LK2016 | A-342 | A-389 | 0.38685 | 0.01280 |
| 758 | LK2016 | A-1.2 | A-547 | 0.64848 | 0.01880 |
| 759 | LK2016 | A-293 | A-547 | 0.08578 | 0.00512 |
| 760 | LK2016 | A-322 | A-547 | 0.64195 | 0.00681 |
| 761 | LK2016 | A-421 | A-547 | 0.91675 | 0.00656 |
| 762 | LK2016 | A-498 | A-547 | 0.25312 | 0.01039 |
| 763 | LK2016 | A-54 | A-547 | 0.25344 | 0.00674 |
| 764 | LK2016 | A-275 | A-547 | 0.55647 | 0.00901 |
| 765 | LK2016 | A-509 | A-547 | 0.95175 | 0.00558 |

| | | | | | |
|---|---|---|---|---|---|
| 766 | LK2016 | A-600 | A-547 | 0.87492 | 0.00350 |
| 767 | LK2016 | A-04 | A-547 | 0.75505 | 0.00517 |
| 768 | LK2016 | A-092 | A-547 | 0.79968 | 0.00291 |
| 769 | LK2016 | A-244 | A-547 | 0.24421 | 0.00958 |
| 770 | LK2016 | A-342 | A-547 | 0.69927 | 0.00916 |
| 771 | LK2016 | A-389 | A-547 | 0.47569 | 0.01117 |
| 772 | LK2016 | A-1.2 | A-196 | 0.90743 | 0.01829 |
| 773 | LK2016 | A-293 | A-196 | 0.17714 | 0.01480 |
| 774 | LK2016 | A-322 | A-196 | 0.72330 | 0.01174 |
| 775 | LK2016 | A-421 | A-196 | 0.45627 | 0.02883 |
| 776 | LK2016 | A-498 | A-196 | 0.83865 | 0.01373 |
| 777 | LK2016 | A-54 | A-196 | 0.99789 | 0.00051 |
| 778 | LK2016 | A-275 | A-196 | 0.90282 | 0.00985 |
| 779 | LK2016 | A-509 | A-196 | 0.50963 | 0.03165 |
| 780 | LK2016 | A-600 | A-196 | 0.07711 | 0.00587 |
| 781 | LK2016 | A-04 | A-196 | 0.48481 | 0.01083 |
| 782 | LK2016 | A-092 | A-196 | 0.85474 | 0.00461 |
| 783 | LK2016 | A-244 | A-196 | 0.60249 | 0.01703 |
| 784 | LK2016 | A-342 | A-196 | 0.34642 | 0.02039 |
| 785 | LK2016 | A-389 | A-196 | 0.50266 | 0.01991 |
| 786 | LK2016 | A-547 | A-196 | 0.70668 | 0.01428 |
| 787 | LK2016 | A-1.2 | A-360 | 0.50788 | 0.01129 |
| 788 | LK2016 | A-293 | A-360 | 0.02110 | 0.00138 |
| 789 | LK2016 | A-322 | A-360 | 0.83525 | 0.00196 |
| 790 | LK2016 | A-421 | A-360 | 0.20178 | 0.00600 |
| 791 | LK2016 | A-498 | A-360 | 0.76236 | 0.00359 |
| 792 | LK2016 | A-54 | A-360 | 0.21959 | 0.00317 |
| 793 | LK2016 | A-275 | A-360 | 0.80903 | 0.00315 |
| 794 | LK2016 | A-509 | A-360 | 0.40832 | 0.00830 |
| 795 | LK2016 | A-600 | A-360 | 0.85912 | 0.00161 |
| 796 | LK2016 | A-04 | A-360 | 0.86661 | 0.00176 |
| 797 | LK2016 | A-092 | A-360 | 0.59246 | 0.00153 |
| 798 | LK2016 | A-244 | A-360 | 0.79051 | 0.00393 |
| 799 | LK2016 | A-342 | A-360 | 0.02938 | 0.00167 |
| 800 | LK2016 | A-389 | A-360 | 0.77376 | 0.00411 |
| 801 | LK2016 | A-547 | A-360 | 0.37217 | 0.00501 |
| 802 | LK2016 | A-196 | A-360 | 0.09278 | 0.00481 |
| 803 | LK2016 | A-1.2 | A-465 | 0.85812 | 0.02505 |
| 804 | LK2016 | A-293 | A-465 | 0.56710 | 0.02408 |
| 805 | LK2016 | A-322 | A-465 | 0.50257 | 0.01437 |
| 806 | LK2016 | A-421 | A-465 | 0.04955 | 0.01072 |
| 807 | LK2016 | A-498 | A-465 | 0.78109 | 0.01928 |
| 808 | LK2016 | A-54 | A-465 | 0.10290 | 0.00804 |
| 809 | LK2016 | A-275 | A-465 | 0.01675 | 0.00460 |
| 810 | LK2016 | A-509 | A-465 | 0.10054 | 0.02269 |
| 811 | LK2016 | A-600 | A-465 | 0.62842 | 0.01207 |
| 812 | LK2016 | A-04 | A-465 | 0.61558 | 0.01274 |
| 813 | LK2016 | A-092 | A-465 | 0.76357 | 0.00736 |

| | | | | | |
|---|---|---|---|---|---|
| 814 | LK2016 | A-244 | A-465 | 0.92303 | 0.00966 |
| 815 | LK2016 | A-342 | A-465 | 0.40418 | 0.02399 |
| 816 | LK2016 | A-389 | A-465 | 0.54093 | 0.02256 |
| 817 | LK2016 | A-547 | A-465 | 0.20667 | 0.01697 |
| 818 | LK2016 | A-196 | A-465 | 0.80019 | 0.02419 |
| 819 | LK2016 | A-360 | A-465 | 0.16460 | 0.00594 |
| 820 | LK2016 | A-1.2 | A-161 | 0.64422 | 0.03168 |
| 821 | LK2016 | A-293 | A-161 | 0.72616 | 0.01664 |
| 822 | LK2016 | A-322 | A-161 | 0.16469 | 0.00855 |
| 823 | LK2016 | A-421 | A-161 | 0.81624 | 0.01655 |
| 824 | LK2016 | A-498 | A-161 | 0.53377 | 0.01650 |
| 825 | LK2016 | A-54 | A-161 | 0.27122 | 0.00966 |
| 826 | LK2016 | A-275 | A-161 | 0.80301 | 0.01165 |
| 827 | LK2016 | A-509 | A-161 | 0.91101 | 0.01491 |
| 828 | LK2016 | A-600 | A-161 | 0.09277 | 0.00609 |
| 829 | LK2016 | A-04 | A-161 | 0.41199 | 0.01121 |
| 830 | LK2016 | A-092 | A-161 | 0.69536 | 0.00693 |
| 831 | LK2016 | A-244 | A-161 | 0.58674 | 0.01654 |
| 832 | LK2016 | A-342 | A-161 | 0.50085 | 0.01919 |
| 833 | LK2016 | A-389 | A-161 | 0.69939 | 0.01643 |
| 834 | LK2016 | A-547 | A-161 | 0.71217 | 0.01215 |
| 835 | LK2016 | A-196 | A-161 | 0.47939 | 0.02608 |
| 836 | LK2016 | A-360 | A-161 | 0.77096 | 0.00590 |
| 837 | LK2016 | A-465 | A-161 | 0.52555 | 0.03310 |
| 838 | LK2016 | A-1.2 | A-200 | 0.00245 | 0.00095 |
| 839 | LK2016 | A-293 | A-200 | 0.46551 | 0.01199 |
| 840 | LK2016 | A-322 | A-200 | 0.00043 | 0.00022 |
| 841 | LK2016 | A-421 | A-200 | 0.01805 | 0.00382 |
| 842 | LK2016 | A-498 | A-200 | 0.25118 | 0.01279 |
| 843 | LK2016 | A-54 | A-200 | 0.96898 | 0.00158 |
| 844 | LK2016 | A-275 | A-200 | 0.01090 | 0.00176 |
| 845 | LK2016 | A-509 | A-200 | 0.93032 | 0.00677 |
| 846 | LK2016 | A-600 | A-200 | 0.70268 | 0.00676 |
| 847 | LK2016 | A-04 | A-200 | 0.74904 | 0.00587 |
| 848 | LK2016 | A-092 | A-200 | 1.00000 | 0.00000 |
| 849 | LK2016 | A-244 | A-200 | 0.06249 | 0.00576 |
| 850 | LK2016 | A-342 | A-200 | 0.42382 | 0.01551 |
| 851 | LK2016 | A-389 | A-200 | 0.54577 | 0.01162 |
| 852 | LK2016 | A-547 | A-200 | 0.34089 | 0.01049 |
| 853 | LK2016 | A-196 | A-200 | 0.08865 | 0.01076 |
| 854 | LK2016 | A-360 | A-200 | 0.37943 | 0.00572 |
| 855 | LK2016 | A-465 | A-200 | 0.38564 | 0.02174 |
| 856 | LK2016 | A-161 | A-200 | 0.26870 | 0.01481 |
| 857 | LK2016 | A-1.2 | A-325 | 0.78770 | 0.01800 |
| 858 | LK2016 | A-293 | A-325 | 0.42710 | 0.01204 |
| 859 | LK2016 | A-322 | A-325 | 0.96885 | 0.00215 |
| 860 | LK2016 | A-421 | A-325 | 0.95277 | 0.00603 |
| 861 | LK2016 | A-498 | A-325 | 0.48902 | 0.01582 |

| | | | | | |
|---|---|---|---|---|---|
| 862 | LK2016 | A-54 | A-325 | 0.71373 | 0.00751 |
| 863 | LK2016 | A-275 | A-325 | 0.54282 | 0.01236 |
| 864 | LK2016 | A-509 | A-325 | 0.71666 | 0.01718 |
| 865 | LK2016 | A-600 | A-325 | 0.55877 | 0.00956 |
| 866 | LK2016 | A-04 | A-325 | 0.01318 | 0.00177 |
| 867 | LK2016 | A-092 | A-325 | 0.03026 | 0.00229 |
| 868 | LK2016 | A-244 | A-325 | 0.30340 | 0.01459 |
| 869 | LK2016 | A-342 | A-325 | 0.03224 | 0.00473 |
| 870 | LK2016 | A-389 | A-325 | 0.53899 | 0.01476 |
| 871 | LK2016 | A-547 | A-325 | 0.48265 | 0.01264 |
| 872 | LK2016 | A-196 | A-325 | 0.25450 | 0.01891 |
| 873 | LK2016 | A-360 | A-325 | 0.14859 | 0.00351 |
| 874 | LK2016 | A-465 | A-325 | 0.92448 | 0.01108 |
| 875 | LK2016 | A-161 | A-325 | 0.37312 | 0.01953 |
| 876 | LK2016 | A-200 | A-325 | 0.95230 | 0.00440 |
| 877 | LK2016 | A-1.2 | A-333 | 0.02605 | 0.00802 |
| 878 | LK2016 | A-293 | A-333 | 0.43724 | 0.01348 |
| 879 | LK2016 | A-322 | A-333 | 0.16641 | 0.00663 |
| 880 | LK2016 | A-421 | A-333 | 0.30946 | 0.02072 |
| 881 | LK2016 | A-498 | A-333 | 0.93714 | 0.00505 |
| 882 | LK2016 | A-54 | A-333 | 0.47406 | 0.00911 |
| 883 | LK2016 | A-275 | A-333 | 0.40086 | 0.01443 |
| 884 | LK2016 | A-509 | A-333 | 0.24491 | 0.02256 |
| 885 | LK2016 | A-600 | A-333 | 0.18156 | 0.00728 |
| 886 | LK2016 | A-04 | A-333 | 0.42791 | 0.00826 |
| 887 | LK2016 | A-092 | A-333 | 0.68715 | 0.00628 |
| 888 | LK2016 | A-244 | A-333 | 0.32795 | 0.01671 |
| 889 | LK2016 | A-342 | A-333 | 0.35405 | 0.01296 |
| 890 | LK2016 | A-389 | A-333 | 0.55863 | 0.01525 |
| 891 | LK2016 | A-547 | A-333 | 0.38855 | 0.01297 |
| 892 | LK2016 | A-196 | A-333 | 0.20598 | 0.01887 |
| 893 | LK2016 | A-360 | A-333 | 0.51859 | 0.00622 |
| 894 | LK2016 | A-465 | A-333 | 0.57034 | 0.02534 |
| 895 | LK2016 | A-161 | A-333 | 0.45192 | 0.02175 |
| 896 | LK2016 | A-200 | A-333 | 0.51317 | 0.01459 |
| 897 | LK2016 | A-325 | A-333 | 0.84366 | 0.01150 |
| 898 | LK2016 | A-1.2 | A-104.2 | 0.83005 | 0.01386 |
| 899 | LK2016 | A-293 | A-104.2 | 0.45519 | 0.01239 |
| 900 | LK2016 | A-322 | A-104.2 | 0.14567 | 0.00557 |
| 901 | LK2016 | A-421 | A-104.2 | 0.06068 | 0.00762 |
| 902 | LK2016 | A-498 | A-104.2 | 0.67417 | 0.01299 |
| 903 | LK2016 | A-54 | A-104.2 | 0.64842 | 0.00693 |
| 904 | LK2016 | A-275 | A-104.2 | 0.21766 | 0.00977 |
| 905 | LK2016 | A-509 | A-104.2 | 0.03122 | 0.00803 |
| 906 | LK2016 | A-600 | A-104.2 | 0.84752 | 0.00503 |
| 907 | LK2016 | A-04 | A-104.2 | 0.40014 | 0.00743 |
| 908 | LK2016 | A-092 | A-104.2 | 0.37091 | 0.00469 |
| 909 | LK2016 | A-244 | A-104.2 | 0.88332 | 0.00596 |

| | | | | | |
|---|---|---|---|---|---|
| 910 | LK2016 | A-342 | A-104.2 | 0.59605 | 0.01053 |
| 911 | LK2016 | A-389 | A-104.2 | 0.58459 | 0.01241 |
| 912 | LK2016 | A-547 | A-104.2 | 0.67315 | 0.01009 |
| 913 | LK2016 | A-196 | A-104.2 | 0.85386 | 0.01270 |
| 914 | LK2016 | A-360 | A-104.2 | 0.10820 | 0.00396 |
| 915 | LK2016 | A-465 | A-104.2 | 0.46613 | 0.02306 |
| 916 | LK2016 | A-161 | A-104.2 | 0.83819 | 0.01151 |
| 917 | LK2016 | A-200 | A-104.2 | 0.22915 | 0.00921 |
| 918 | LK2016 | A-325 | A-104.2 | 0.34045 | 0.01448 |
| 919 | LK2016 | A-333 | A-104.2 | 0.51730 | 0.01625 |
| 920 | LK2016 | A-1.2 | A-122 | 0.05204 | 0.01208 |
| 921 | LK2016 | A-293 | A-122 | 0.33518 | 0.01286 |
| 922 | LK2016 | A-322 | A-122 | 0.00000 | 0.00000 |
| 923 | LK2016 | A-421 | A-122 | 0.14109 | 0.01312 |
| 924 | LK2016 | A-498 | A-122 | 0.44688 | 0.01423 |
| 925 | LK2016 | A-54 | A-122 | 0.88143 | 0.00432 |
| 926 | LK2016 | A-275 | A-122 | 0.00000 | 0.00000 |
| 927 | LK2016 | A-509 | A-122 | 0.59136 | 0.02245 |
| 928 | LK2016 | A-600 | A-122 | 0.12411 | 0.00559 |
| 929 | LK2016 | A-04 | A-122 | 0.25214 | 0.00702 |
| 930 | LK2016 | A-092 | A-122 | 0.90706 | 0.00251 |
| 931 | LK2016 | A-244 | A-122 | 0.65084 | 0.01109 |
| 932 | LK2016 | A-342 | A-122 | 0.01626 | 0.00284 |
| 933 | LK2016 | A-389 | A-122 | 0.88118 | 0.00766 |
| 934 | LK2016 | A-547 | A-122 | 0.26858 | 0.00875 |
| 935 | LK2016 | A-196 | A-122 | 0.99450 | 0.00169 |
| 936 | LK2016 | A-360 | A-122 | 0.63297 | 0.00517 |
| 937 | LK2016 | A-465 | A-122 | 0.15548 | 0.01802 |
| 938 | LK2016 | A-161 | A-122 | 0.83455 | 0.01084 |
| 939 | LK2016 | A-200 | A-122 | 0.01848 | 0.00259 |
| 940 | LK2016 | A-325 | A-122 | 0.75491 | 0.01296 |
| 941 | LK2016 | A-333 | A-122 | 0.09428 | 0.00844 |
| 942 | LK2016 | A-104.2 | A-122 | 0.41935 | 0.01288 |
| 943 | LK2016 | A-1.2 | A-175 | 0.69414 | 0.02205 |
| 944 | LK2016 | A-293 | A-175 | 0.66269 | 0.01293 |
| 945 | LK2016 | A-322 | A-175 | 0.69381 | 0.00691 |
| 946 | LK2016 | A-421 | A-175 | 0.25925 | 0.01619 |
| 947 | LK2016 | A-498 | A-175 | 0.45579 | 0.01331 |
| 948 | LK2016 | A-54 | A-175 | 0.94697 | 0.00210 |
| 949 | LK2016 | A-275 | A-175 | 0.24670 | 0.01114 |
| 950 | LK2016 | A-509 | A-175 | 0.76568 | 0.01807 |
| 951 | LK2016 | A-600 | A-175 | 0.81929 | 0.00543 |
| 952 | LK2016 | A-04 | A-175 | 0.46206 | 0.00713 |
| 953 | LK2016 | A-092 | A-175 | 0.19247 | 0.00391 |
| 954 | LK2016 | A-244 | A-175 | 0.24405 | 0.01170 |
| 955 | LK2016 | A-342 | A-175 | 0.01226 | 0.00209 |
| 956 | LK2016 | A-389 | A-175 | 0.64899 | 0.01306 |
| 957 | LK2016 | A-547 | A-175 | 0.04491 | 0.00406 |

| | | | | | |
|---|---|---|---|---|---|
| 958 | LK2016 | A-196 | A-175 | 0.90817 | 0.01034 |
| 959 | LK2016 | A-360 | A-175 | 0.22767 | 0.00387 |
| 960 | LK2016 | A-465 | A-175 | 0.54701 | 0.02336 |
| 961 | LK2016 | A-161 | A-175 | 0.02390 | 0.00386 |
| 962 | LK2016 | A-200 | A-175 | 0.40624 | 0.01498 |
| 963 | LK2016 | A-325 | A-175 | 0.31118 | 0.01304 |
| 964 | LK2016 | A-333 | A-175 | 0.11811 | 0.01014 |
| 965 | LK2016 | A-104.2 | A-175 | 0.79678 | 0.00910 |
| 966 | LK2016 | A-122 | A-175 | 0.18566 | 0.00991 |
| 967 | LK2016 | A-1.2 | A-300 | 0.27610 | 0.03030 |
| 968 | LK2016 | A-293 | A-300 | 0.07541 | 0.00935 |
| 969 | LK2016 | A-322 | A-300 | 0.90547 | 0.00893 |
| 970 | LK2016 | A-421 | A-300 | 0.42546 | 0.02737 |
| 971 | LK2016 | A-498 | A-300 | 0.86692 | 0.01323 |
| 972 | LK2016 | A-54 | A-300 | 0.95177 | 0.00423 |
| 973 | LK2016 | A-275 | A-300 | 0.66119 | 0.02035 |
| 974 | LK2016 | A-509 | A-300 | 0.02196 | 0.00854 |
| 975 | LK2016 | A-600 | A-300 | 0.51518 | 0.01511 |
| 976 | LK2016 | A-04 | A-300 | 0.39123 | 0.01188 |
| 977 | LK2016 | A-092 | A-300 | 0.33716 | 0.00893 |
| 978 | LK2016 | A-244 | A-300 | 0.87707 | 0.00977 |
| 979 | LK2016 | A-342 | A-300 | 0.10901 | 0.01476 |
| 980 | LK2016 | A-389 | A-300 | 0.21100 | 0.01588 |
| 981 | LK2016 | A-547 | A-300 | 0.08894 | 0.01063 |
| 982 | LK2016 | A-196 | A-300 | 0.54412 | 0.03306 |
| 983 | LK2016 | A-360 | A-300 | 0.56796 | 0.01085 |
| 984 | LK2016 | A-465 | A-300 | 0.80523 | 0.02639 |
| 985 | LK2016 | A-161 | A-300 | 0.65718 | 0.02924 |
| 986 | LK2016 | A-200 | A-300 | 0.70781 | 0.02076 |
| 987 | LK2016 | A-325 | A-300 | 0.00913 | 0.00396 |
| 988 | LK2016 | A-333 | A-300 | 0.56695 | 0.02786 |
| 989 | LK2016 | A-104.2 | A-300 | 0.99698 | 0.00125 |
| 990 | LK2016 | A-122 | A-300 | 0.62264 | 0.02461 |
| 991 | LK2016 | A-175 | A-300 | 0.00510 | 0.00197 |
| 992 | LK2016 | A-1.2 | A-733 | 0.35618 | 0.00980 |
| 993 | LK2016 | A-293 | A-733 | 0.30411 | 0.00373 |
| 994 | LK2016 | A-322 | A-733 | 1.00000 | 0.00000 |
| 995 | LK2016 | A-421 | A-733 | 0.77887 | 0.00595 |
| 996 | LK2016 | A-498 | A-733 | 0.28018 | 0.00476 |
| 997 | LK2016 | A-54 | A-733 | 0.37631 | 0.00342 |
| 998 | LK2016 | A-275 | A-733 | 0.97784 | 0.00084 |
| 999 | LK2016 | A-509 | A-733 | 0.97463 | 0.00197 |
| 1000 | LK2016 | A-600 | A-733 | 0.55230 | 0.00342 |
| 1001 | LK2016 | A-04 | A-733 | 0.90398 | 0.00132 |
| 1002 | LK2016 | A-092 | A-733 | 0.55349 | 0.00154 |
| 1003 | LK2016 | A-244 | A-733 | 0.45821 | 0.00582 |
| 1004 | LK2016 | A-342 | A-733 | 0.80189 | 0.00293 |
| 1005 | LK2016 | A-389 | A-733 | 0.13811 | 0.00486 |

| | | | | | |
|---|---|---|---|---|---|
| 1006 | LK2016 | A-547 | A-733 | 0.74971 | 0.00339 |
| 1007 | LK2016 | A-196 | A-733 | 0.10772 | 0.00478 |
| 1008 | LK2016 | A-360 | A-733 | 1.00000 | 0.00000 |
| 1009 | LK2016 | A-465 | A-733 | 0.99015 | 0.00126 |
| 1010 | LK2016 | A-161 | A-733 | 0.07087 | 0.00396 |
| 1011 | LK2016 | A-200 | A-733 | 0.81276 | 0.00323 |
| 1012 | LK2016 | A-325 | A-733 | 0.51959 | 0.00593 |
| 1013 | LK2016 | A-333 | A-733 | 0.22834 | 0.00560 |
| 1014 | LK2016 | A-104.2 | A-733 | 0.19021 | 0.00407 |
| 1015 | LK2016 | A-122 | A-733 | 1.00000 | 0.00000 |
| 1016 | LK2016 | A-175 | A-733 | 0.62888 | 0.00429 |
| 1017 | LK2016 | A-300 | A-733 | 0.35256 | 0.00963 |
| 1018 | LK2016 | A-1.2 | A-0 | 0.38249 | 0.02974 |
| 1019 | LK2016 | A-293 | A-0 | 0.88388 | 0.00623 |
| 1020 | LK2016 | A-322 | A-0 | 0.56965 | 0.01264 |
| 1021 | LK2016 | A-421 | A-0 | 0.35281 | 0.02068 |
| 1022 | LK2016 | A-498 | A-0 | 0.44594 | 0.01681 |
| 1023 | LK2016 | A-54 | A-0 | 0.19685 | 0.00819 |
| 1024 | LK2016 | A-275 | A-0 | 0.79934 | 0.01195 |
| 1025 | LK2016 | A-509 | A-0 | 0.19904 | 0.02041 |
| 1026 | LK2016 | A-600 | A-0 | 0.38188 | 0.00880 |
| 1027 | LK2016 | A-04 | A-0 | 0.52491 | 0.00884 |
| 1028 | LK2016 | A-092 | A-0 | 0.22234 | 0.00463 |
| 1029 | LK2016 | A-244 | A-0 | 0.83567 | 0.00853 |
| 1030 | LK2016 | A-342 | A-0 | 0.09853 | 0.00804 |
| 1031 | LK2016 | A-389 | A-0 | 0.85927 | 0.00874 |
| 1032 | LK2016 | A-547 | A-0 | 0.16965 | 0.00845 |
| 1033 | LK2016 | A-196 | A-0 | 0.29768 | 0.02401 |
| 1034 | LK2016 | A-360 | A-0 | 0.03326 | 0.00213 |
| 1035 | LK2016 | A-465 | A-0 | 0.93920 | 0.01364 |
| 1036 | LK2016 | A-161 | A-0 | 0.23445 | 0.01903 |
| 1037 | LK2016 | A-200 | A-0 | 0.03149 | 0.00443 |
| 1038 | LK2016 | A-325 | A-0 | 0.21673 | 0.01464 |
| 1039 | LK2016 | A-333 | A-0 | 0.54775 | 0.02104 |
| 1040 | LK2016 | A-104.2 | A-0 | 0.73500 | 0.01203 |
| 1041 | LK2016 | A-122 | A-0 | 0.36115 | 0.01761 |
| 1042 | LK2016 | A-175 | A-0 | 0.39369 | 0.01845 |
| 1043 | LK2016 | A-300 | A-0 | 0.93017 | 0.01335 |
| 1044 | LK2016 | A-733 | A-0 | 0.21923 | 0.00580 |
| 1045 | LK2016 | A-1.2 | A-1.1 | 0.00000 | 0.00000 |
| 1046 | LK2016 | A-293 | A-1.1 | 0.06066 | 0.00576 |
| 1047 | LK2016 | A-322 | A-1.1 | 0.21342 | 0.00668 |
| 1048 | LK2016 | A-421 | A-1.1 | 0.37334 | 0.01687 |
| 1049 | LK2016 | A-498 | A-1.1 | 0.49457 | 0.01366 |
| 1050 | LK2016 | A-54 | A-1.1 | 0.67419 | 0.00767 |
| 1051 | LK2016 | A-275 | A-1.1 | 0.03107 | 0.00442 |
| 1052 | LK2016 | A-509 | A-1.1 | 0.78160 | 0.01699 |
| 1053 | LK2016 | A-600 | A-1.1 | 0.52975 | 0.00875 |

| | | | | | |
|---|---|---|---|---|---|
| 1054 | LK2016 | A-04 | A-1.1 | 0.04567 | 0.00316 |
| 1055 | LK2016 | A-092 | A-1.1 | 0.84863 | 0.00364 |
| 1056 | LK2016 | A-244 | A-1.1 | 0.23878 | 0.01368 |
| 1057 | LK2016 | A-342 | A-1.1 | 0.73330 | 0.01125 |
| 1058 | LK2016 | A-389 | A-1.1 | 0.26144 | 0.01190 |
| 1059 | LK2016 | A-547 | A-1.1 | 0.35451 | 0.01122 |
| 1060 | LK2016 | A-196 | A-1.1 | 0.76492 | 0.01862 |
| 1061 | LK2016 | A-360 | A-1.1 | 0.02870 | 0.00187 |
| 1062 | LK2016 | A-465 | A-1.1 | 0.18508 | 0.01790 |
| 1063 | LK2016 | A-161 | A-1.1 | 0.65658 | 0.01753 |
| 1064 | LK2016 | A-200 | A-1.1 | 0.00036 | 0.00032 |
| 1065 | LK2016 | A-325 | A-1.1 | 0.52575 | 0.01556 |
| 1066 | LK2016 | A-333 | A-1.1 | 0.21451 | 0.01342 |
| 1067 | LK2016 | A-104.2 | A-1.1 | 0.42306 | 0.01347 |
| 1068 | LK2016 | A-122 | A-1.1 | 0.62402 | 0.01414 |
| 1069 | LK2016 | A-175 | A-1.1 | 0.63406 | 0.01344 |
| 1070 | LK2016 | A-300 | A-1.1 | 0.39271 | 0.02347 |
| 1071 | LK2016 | A-733 | A-1.1 | 0.29157 | 0.00563 |
| 1072 | LK2016 | A-0 | A-1.1 | 0.59428 | 0.01691 |
| 1073 | LK2016 | A-1.2 | A-450 | 0.23050 | 0.01466 |
| 1074 | LK2016 | A-293 | A-450 | 0.08581 | 0.00521 |
| 1075 | LK2016 | A-322 | A-450 | 0.15276 | 0.00474 |
| 1076 | LK2016 | A-421 | A-450 | 0.08316 | 0.00800 |
| 1077 | LK2016 | A-498 | A-450 | 0.17589 | 0.00732 |
| 1078 | LK2016 | A-54 | A-450 | 0.36220 | 0.00642 |
| 1079 | LK2016 | A-275 | A-450 | 0.54292 | 0.00987 |
| 1080 | LK2016 | A-509 | A-450 | 0.60586 | 0.01367 |
| 1081 | LK2016 | A-600 | A-450 | 0.09186 | 0.00354 |
| 1082 | LK2016 | A-04 | A-450 | 0.01564 | 0.00140 |
| 1083 | LK2016 | A-092 | A-450 | 0.87247 | 0.00214 |
| 1084 | LK2016 | A-244 | A-450 | 0.49497 | 0.01006 |
| 1085 | LK2016 | A-342 | A-450 | 0.00308 | 0.00087 |
| 1086 | LK2016 | A-389 | A-450 | 0.76922 | 0.00868 |
| 1087 | LK2016 | A-547 | A-450 | 0.60952 | 0.00862 |
| 1088 | LK2016 | A-196 | A-450 | 0.58995 | 0.01590 |
| 1089 | LK2016 | A-360 | A-450 | 0.50105 | 0.00428 |
| 1090 | LK2016 | A-465 | A-450 | 0.91839 | 0.00824 |
| 1091 | LK2016 | A-161 | A-450 | 0.25732 | 0.01355 |
| 1092 | LK2016 | A-200 | A-450 | 0.43212 | 0.00929 |
| 1093 | LK2016 | A-325 | A-450 | 0.79106 | 0.00794 |
| 1094 | LK2016 | A-333 | A-450 | 0.71925 | 0.00951 |
| 1095 | LK2016 | A-104.2 | A-450 | 0.21305 | 0.00725 |
| 1096 | LK2016 | A-122 | A-450 | 0.31877 | 0.01031 |
| 1097 | LK2016 | A-175 | A-450 | 0.18377 | 0.00789 |
| 1098 | LK2016 | A-300 | A-450 | 0.30651 | 0.01711 |
| 1099 | LK2016 | A-733 | A-450 | 0.26044 | 0.00476 |
| 1100 | LK2016 | A-0 | A-450 | 0.31430 | 0.01171 |
| 1101 | LK2016 | A-1.1 | A-450 | 0.97878 | 0.00182 |

| | | | | | |
|---|---|---|---|---|---|
| 1102 | LK2016 | A-1.2 | A-60 | 0.61861 | 0.03399 |
| 1103 | LK2016 | A-293 | A-60 | 0.93916 | 0.00698 |
| 1104 | LK2016 | A-322 | A-60 | 0.01706 | 0.00270 |
| 1105 | LK2016 | A-421 | A-60 | 0.16583 | 0.01700 |
| 1106 | LK2016 | A-498 | A-60 | 0.98678 | 0.00342 |
| 1107 | LK2016 | A-54 | A-60 | 0.65913 | 0.00936 |
| 1108 | LK2016 | A-275 | A-60 | 0.10936 | 0.01261 |
| 1109 | LK2016 | A-509 | A-60 | 0.99987 | 0.00013 |
| 1110 | LK2016 | A-600 | A-60 | 0.78325 | 0.00785 |
| 1111 | LK2016 | A-04 | A-60 | 0.14902 | 0.00751 |
| 1112 | LK2016 | A-092 | A-60 | 0.03761 | 0.00275 |
| 1113 | LK2016 | A-244 | A-60 | 0.56439 | 0.02085 |
| 1114 | LK2016 | A-342 | A-60 | 0.88480 | 0.01171 |
| 1115 | LK2016 | A-389 | A-60 | 0.36928 | 0.02003 |
| 1116 | LK2016 | A-547 | A-60 | 0.98007 | 0.00291 |
| 1117 | LK2016 | A-196 | A-60 | 0.20989 | 0.02802 |
| 1118 | LK2016 | A-360 | A-60 | 0.96178 | 0.00238 |
| 1119 | LK2016 | A-465 | A-60 | 0.47736 | 0.03504 |
| 1120 | LK2016 | A-161 | A-60 | 0.35804 | 0.02517 |
| 1121 | LK2016 | A-200 | A-60 | 0.00033 | 0.00033 |
| 1122 | LK2016 | A-325 | A-60 | 0.08093 | 0.01231 |
| 1123 | LK2016 | A-333 | A-60 | 0.87927 | 0.01293 |
| 1124 | LK2016 | A-104.2 | A-60 | 0.84860 | 0.01249 |
| 1125 | LK2016 | A-122 | A-60 | 0.53820 | 0.02198 |
| 1126 | LK2016 | A-175 | A-60 | 0.66780 | 0.02062 |
| 1127 | LK2016 | A-300 | A-60 | 0.19597 | 0.02851 |
| 1128 | LK2016 | A-733 | A-60 | 0.95828 | 0.00219 |
| 1129 | LK2016 | A-0 | A-60 | 0.82836 | 0.01547 |
| 1130 | LK2016 | A-1.1 | A-60 | 0.01140 | 0.00366 |
| 1131 | LK2016 | A-450 | A-60 | 0.63299 | 0.01537 |
| 1132 | LK2016 | A-1.2 | A-662 | 0.03898 | 0.00587 |
| 1133 | LK2016 | A-293 | A-662 | 0.61730 | 0.00696 |
| 1134 | LK2016 | A-322 | A-662 | 0.00000 | 0.00000 |
| 1135 | LK2016 | A-421 | A-662 | 0.34878 | 0.01124 |
| 1136 | LK2016 | A-498 | A-662 | 0.33762 | 0.00965 |
| 1137 | LK2016 | A-54 | A-662 | 0.71176 | 0.00405 |
| 1138 | LK2016 | A-275 | A-662 | 0.00000 | 0.00000 |
| 1139 | LK2016 | A-509 | A-662 | 0.26152 | 0.01216 |
| 1140 | LK2016 | A-600 | A-662 | 0.75897 | 0.00339 |
| 1141 | LK2016 | A-04 | A-662 | 0.42406 | 0.00449 |
| 1142 | LK2016 | A-092 | A-662 | 1.00000 | 0.00000 |
| 1143 | LK2016 | A-244 | A-662 | 0.36501 | 0.00785 |
| 1144 | LK2016 | A-342 | A-662 | 0.99272 | 0.00089 |
| 1145 | LK2016 | A-389 | A-662 | 0.75166 | 0.00693 |
| 1146 | LK2016 | A-547 | A-662 | 0.62691 | 0.00546 |
| 1147 | LK2016 | A-196 | A-662 | 0.73540 | 0.01126 |
| 1148 | LK2016 | A-360 | A-662 | 0.83690 | 0.00183 |
| 1149 | LK2016 | A-465 | A-662 | 0.49587 | 0.01522 |

| | | | | | |
|---|---|---|---|---|---|
| 1150 | LK2016 | A-161 | A-662 | 0.17438 | 0.00852 |
| 1151 | LK2016 | A-200 | A-662 | 0.00018 | 0.00014 |
| 1152 | LK2016 | A-325 | A-662 | 0.96788 | 0.00251 |
| 1153 | LK2016 | A-333 | A-662 | 0.15229 | 0.00681 |
| 1154 | LK2016 | A-104.2 | A-662 | 0.15285 | 0.00648 |
| 1155 | LK2016 | A-122 | A-662 | 0.00000 | 0.00000 |
| 1156 | LK2016 | A-175 | A-662 | 0.68550 | 0.00640 |
| 1157 | LK2016 | A-300 | A-662 | 0.92926 | 0.00602 |
| 1158 | LK2016 | A-733 | A-662 | 1.00000 | 0.00000 |
| 1159 | LK2016 | A-0 | A-662 | 0.55909 | 0.01225 |
| 1160 | LK2016 | A-1.1 | A-662 | 0.21477 | 0.00664 |
| 1161 | LK2016 | A-450 | A-662 | 0.16737 | 0.00481 |
| 1162 | LK2016 | A-60 | A-662 | 0.01167 | 0.00207 |
| 1163 | LK2016 | A-1.2 | A-104.1 | No | contingency |
| 1164 | LK2016 | A-293 | A-104.1 | No | contingency |
| 1165 | LK2016 | A-322 | A-104.1 | No | contingency |
| 1166 | LK2016 | A-421 | A-104.1 | No | contingency |
| 1167 | LK2016 | A-498 | A-104.1 | No | contingency |
| 1168 | LK2016 | A-54 | A-104.1 | No | contingency |
| 1169 | LK2016 | A-275 | A-104.1 | No | contingency |
| 1170 | LK2016 | A-509 | A-104.1 | No | contingency |
| 1171 | LK2016 | A-600 | A-104.1 | No | contingency |
| 1172 | LK2016 | A-04 | A-104.1 | No | contingency |
| 1173 | LK2016 | A-092 | A-104.1 | No | contingency |
| 1174 | LK2016 | A-244 | A-104.1 | No | contingency |
| 1175 | LK2016 | A-342 | A-104.1 | No | contingency |
| 1176 | LK2016 | A-389 | A-104.1 | No | contingency |
| 1177 | LK2016 | A-547 | A-104.1 | No | contingency |
| 1178 | LK2016 | A-196 | A-104.1 | No | contingency |
| 1179 | LK2016 | A-360 | A-104.1 | No | contingency |
| 1180 | LK2016 | A-465 | A-104.1 | No | contingency |
| 1181 | LK2016 | A-161 | A-104.1 | No | contingency |
| 1182 | LK2016 | A-200 | A-104.1 | No | contingency |
| 1183 | LK2016 | A-325 | A-104.1 | No | contingency |
| 1184 | LK2016 | A-333 | A-104.1 | No | contingency |
| 1185 | LK2016 | A-104.2 | A-104.1 | No | contingency |
| 1186 | LK2016 | A-122 | A-104.1 | No | contingency |
| 1187 | LK2016 | A-175 | A-104.1 | No | contingency |
| 1188 | LK2016 | A-300 | A-104.1 | No | contingency |
| 1189 | LK2016 | A-733 | A-104.1 | No | contingency |
| 1190 | LK2016 | A-0 | A-104.1 | No | contingency |
| 1191 | LK2016 | A-1.1 | A-104.1 | No | contingency |
| 1192 | LK2016 | A-450 | A-104.1 | No | contingency |
| 1193 | LK2016 | A-60 | A-104.1 | No | contingency |
| 1194 | LK2016 | A-662 | A-104.1 | No | contingency |
| 1195 | LK2016 | A-1.2 | A-149 | 0.13425 | 0.01863 |
| 1196 | LK2016 | A-293 | A-149 | 0.01497 | 0.00227 |
| 1197 | LK2016 | A-322 | A-149 | 0.09692 | 0.00549 |

| | | | | | |
|---|---|---|---|---|---|
| 1198 | LK2016 | A-421 | A-149 | 0.48122 | 0.01922 |
| 1199 | LK2016 | A-498 | A-149 | 0.54394 | 0.01535 |
| 1200 | LK2016 | A-54 | A-149 | 0.83304 | 0.00526 |
| 1201 | LK2016 | A-275 | A-149 | 0.55460 | 0.01391 |
| 1202 | LK2016 | A-509 | A-149 | 0.68168 | 0.02211 |
| 1203 | LK2016 | A-600 | A-149 | 0.33058 | 0.00788 |
| 1204 | LK2016 | A-04 | A-149 | 0.89369 | 0.00375 |
| 1205 | LK2016 | A-092 | A-149 | 0.27575 | 0.00524 |
| 1206 | LK2016 | A-244 | A-149 | 0.98066 | 0.00243 |
| 1207 | LK2016 | A-342 | A-149 | 0.56539 | 0.01451 |
| 1208 | LK2016 | A-389 | A-149 | 0.03684 | 0.00463 |
| 1209 | LK2016 | A-547 | A-149 | 0.00885 | 0.00169 |
| 1210 | LK2016 | A-196 | A-149 | 0.13367 | 0.01345 |
| 1211 | LK2016 | A-360 | A-149 | 0.28735 | 0.00616 |
| 1212 | LK2016 | A-465 | A-149 | 0.06919 | 0.00938 |
| 1213 | LK2016 | A-161 | A-149 | 0.00948 | 0.00220 |
| 1214 | LK2016 | A-200 | A-149 | 0.17836 | 0.01122 |
| 1215 | LK2016 | A-325 | A-149 | 0.50748 | 0.01733 |
| 1216 | LK2016 | A-333 | A-149 | 0.93467 | 0.00663 |
| 1217 | LK2016 | A-104.2 | A-149 | 0.82179 | 0.01015 |
| 1218 | LK2016 | A-122 | A-149 | 0.31103 | 0.01386 |
| 1219 | LK2016 | A-175 | A-149 | 0.14000 | 0.00988 |
| 1220 | LK2016 | A-300 | A-149 | 0.56365 | 0.02460 |
| 1221 | LK2016 | A-733 | A-149 | 0.89874 | 0.00225 |
| 1222 | LK2016 | A-0 | A-149 | 0.25875 | 0.01638 |
| 1223 | LK2016 | A-1.1 | A-149 | 0.46034 | 0.01485 |
| 1224 | LK2016 | A-450 | A-149 | 0.42791 | 0.01033 |
| 1225 | LK2016 | A-60 | A-149 | 0.68163 | 0.02296 |
| 1226 | LK2016 | A-662 | A-149 | 0.11639 | 0.00620 |
| 1227 | LK2016 | A-104.1 | A-149 | No | contingency |
| 1228 | LK2016 | A-1.2 | A-224 | 0.55450 | 0.03394 |
| 1229 | LK2016 | A-293 | A-224 | 0.44054 | 0.01872 |
| 1230 | LK2016 | A-322 | A-224 | 0.00000 | 0.00000 |
| 1231 | LK2016 | A-421 | A-224 | 0.64394 | 0.02522 |
| 1232 | LK2016 | A-498 | A-224 | 0.40414 | 0.01778 |
| 1233 | LK2016 | A-54 | A-224 | 0.45447 | 0.00968 |
| 1234 | LK2016 | A-275 | A-224 | 0.00000 | 0.00000 |
| 1235 | LK2016 | A-509 | A-224 | 0.90920 | 0.01487 |
| 1236 | LK2016 | A-600 | A-224 | 0.98206 | 0.00144 |
| 1237 | LK2016 | A-04 | A-224 | 0.86100 | 0.00609 |
| 1238 | LK2016 | A-092 | A-224 | 1.00000 | 0.00000 |
| 1239 | LK2016 | A-244 | A-224 | 0.07568 | 0.00828 |
| 1240 | LK2016 | A-342 | A-224 | 0.87494 | 0.00889 |
| 1241 | LK2016 | A-389 | A-224 | 0.57696 | 0.01767 |
| 1242 | LK2016 | A-547 | A-224 | 0.01976 | 0.00328 |
| 1243 | LK2016 | A-196 | A-224 | 0.72166 | 0.02293 |
| 1244 | LK2016 | A-360 | A-224 | 0.75678 | 0.00604 |
| 1245 | LK2016 | A-465 | A-224 | 0.40806 | 0.03453 |

| | | | | | |
|---|---|---|---|---|---|
| 1246 | LK2016 | A-161 | A-224 | 0.27716 | 0.02398 |
| 1247 | LK2016 | A-200 | A-224 | 0.00427 | 0.00150 |
| 1248 | LK2016 | A-325 | A-224 | 0.74936 | 0.01717 |
| 1249 | LK2016 | A-333 | A-224 | 0.90954 | 0.00956 |
| 1250 | LK2016 | A-104.2 | A-224 | 0.22202 | 0.01384 |
| 1251 | LK2016 | A-122 | A-224 | 0.00000 | 0.00000 |
| 1252 | LK2016 | A-175 | A-224 | 0.80878 | 0.01295 |
| 1253 | LK2016 | A-300 | A-224 | 0.74314 | 0.02489 |
| 1254 | LK2016 | A-733 | A-224 | 0.96578 | 0.00181 |
| 1255 | LK2016 | A-0 | A-224 | 0.98058 | 0.00485 |
| 1256 | LK2016 | A-1.1 | A-224 | 0.29127 | 0.01756 |
| 1257 | LK2016 | A-450 | A-224 | 0.53224 | 0.01469 |
| 1258 | LK2016 | A-60 | A-224 | 0.01391 | 0.00504 |
| 1259 | LK2016 | A-662 | A-224 | 0.00000 | 0.00000 |
| 1260 | LK2016 | A-104.1 | A-224 | No | contingency |
| 1261 | LK2016 | A-149 | A-224 | 0.00981 | 0.00305 |
| 1262 | LK2016 | A-1.2 | A-323 | 0.80268 | 0.03174 |
| 1263 | LK2016 | A-293 | A-323 | 0.34151 | 0.02272 |
| 1264 | LK2016 | A-322 | A-323 | 0.89472 | 0.01014 |
| 1265 | LK2016 | A-421 | A-323 | 0.75999 | 0.02809 |
| 1266 | LK2016 | A-498 | A-323 | 0.60459 | 0.02745 |
| 1267 | LK2016 | A-54 | A-323 | 0.15571 | 0.01128 |
| 1268 | LK2016 | A-275 | A-323 | 0.97242 | 0.00718 |
| 1269 | LK2016 | A-509 | A-323 | 0.88798 | 0.02428 |
| 1270 | LK2016 | A-600 | A-323 | 0.51063 | 0.01477 |
| 1271 | LK2016 | A-04 | A-323 | 0.85247 | 0.00806 |
| 1272 | LK2016 | A-092 | A-323 | 0.21451 | 0.00752 |
| 1273 | LK2016 | A-244 | A-323 | 0.28389 | 0.02370 |
| 1274 | LK2016 | A-342 | A-323 | 0.37408 | 0.02543 |
| 1275 | LK2016 | A-389 | A-323 | 0.44176 | 0.02433 |
| 1276 | LK2016 | A-547 | A-323 | 0.94161 | 0.00825 |
| 1277 | LK2016 | A-196 | A-323 | 0.99986 | 0.00014 |
| 1278 | LK2016 | A-360 | A-323 | 0.26781 | 0.01066 |
| 1279 | LK2016 | A-465 | A-323 | 0.89906 | 0.02367 |
| 1280 | LK2016 | A-161 | A-323 | 0.63852 | 0.03223 |
| 1281 | LK2016 | A-200 | A-323 | 0.88432 | 0.01482 |
| 1282 | LK2016 | A-325 | A-323 | 0.08618 | 0.01474 |
| 1283 | LK2016 | A-333 | A-323 | 0.61599 | 0.02839 |
| 1284 | LK2016 | A-104.2 | A-323 | 0.32836 | 0.02195 |
| 1285 | LK2016 | A-122 | A-323 | 0.48815 | 0.02745 |
| 1286 | LK2016 | A-175 | A-323 | 0.09158 | 0.01476 |
| 1287 | LK2016 | A-300 | A-323 | 0.76929 | 0.03062 |
| 1288 | LK2016 | A-733 | A-323 | 0.10383 | 0.00616 |
| 1289 | LK2016 | A-0 | A-323 | 0.75509 | 0.02512 |
| 1290 | LK2016 | A-1.1 | A-323 | 0.04368 | 0.00811 |
| 1291 | LK2016 | A-450 | A-323 | 0.99988 | 0.00007 |
| 1292 | LK2016 | A-60 | A-323 | 0.71560 | 0.03238 |
| 1293 | LK2016 | A-662 | A-323 | 0.88372 | 0.00918 |

| | | | | | |
|---|---|---|---|---|---|
| 1294 | LK2016 | A-104.1 | A-323 | No | contingency |
| 1295 | LK2016 | A-149 | A-323 | 0.11199 | 0.01654 |
| 1296 | LK2016 | A-224 | A-323 | 0.41029 | 0.03241 |
| 1297 | LK2016 | A-1.2 | A-880 | 0.53570 | 0.01268 |
| 1298 | LK2016 | A-293 | A-880 | 0.44416 | 0.00676 |
| 1299 | LK2016 | A-322 | A-880 | 0.75208 | 0.00396 |
| 1300 | LK2016 | A-421 | A-880 | 0.71945 | 0.00887 |
| 1301 | LK2016 | A-498 | A-880 | 0.77727 | 0.00650 |
| 1302 | LK2016 | A-54 | A-880 | 0.18842 | 0.00462 |
| 1303 | LK2016 | A-275 | A-880 | 0.34393 | 0.00692 |
| 1304 | LK2016 | A-509 | A-880 | 0.67922 | 0.00938 |
| 1305 | LK2016 | A-600 | A-880 | 0.46387 | 0.00496 |
| 1306 | LK2016 | A-04 | A-880 | 0.51590 | 0.00548 |
| 1307 | LK2016 | A-092 | A-880 | 0.72080 | 0.00229 |
| 1308 | LK2016 | A-244 | A-880 | 0.01542 | 0.00164 |
| 1309 | LK2016 | A-342 | A-880 | 0.81671 | 0.00577 |
| 1310 | LK2016 | A-389 | A-880 | 0.83755 | 0.00467 |
| 1311 | LK2016 | A-547 | A-880 | 0.81837 | 0.00417 |
| 1312 | LK2016 | A-196 | A-880 | 0.33846 | 0.01218 |
| 1313 | LK2016 | A-360 | A-880 | 0.42425 | 0.00376 |
| 1314 | LK2016 | A-465 | A-880 | 0.12611 | 0.00844 |
| 1315 | LK2016 | A-161 | A-880 | 0.51380 | 0.00895 |
| 1316 | LK2016 | A-200 | A-880 | 0.50247 | 0.00787 |
| 1317 | LK2016 | A-325 | A-880 | 0.01092 | 0.00162 |
| 1318 | LK2016 | A-333 | A-880 | 0.62430 | 0.00831 |
| 1319 | LK2016 | A-104.2 | A-880 | 0.46348 | 0.00766 |
| 1320 | LK2016 | A-122 | A-880 | 0.56493 | 0.00842 |
| 1321 | LK2016 | A-175 | A-880 | 0.49195 | 0.00785 |
| 1322 | LK2016 | A-300 | A-880 | 0.16582 | 0.01172 |
| 1323 | LK2016 | A-733 | A-880 | 0.04621 | 0.00206 |
| 1324 | LK2016 | A-0 | A-880 | 0.41067 | 0.00836 |
| 1325 | LK2016 | A-1.1 | A-880 | 0.19514 | 0.00592 |
| 1326 | LK2016 | A-450 | A-880 | 0.68790 | 0.00608 |
| 1327 | LK2016 | A-60 | A-880 | 0.88629 | 0.00538 |
| 1328 | LK2016 | A-662 | A-880 | 0.75630 | 0.00370 |
| 1329 | LK2016 | A-104.1 | A-880 | No | contingency |
| 1330 | LK2016 | A-149 | A-880 | 0.37851 | 0.00831 |
| 1331 | LK2016 | A-224 | A-880 | 0.93941 | 0.00356 |
| 1332 | LK2016 | A-323 | A-880 | 0.71619 | 0.01435 |
| 1333 | UK2014 | A-1.2 | A-293 | 0.42330 | 0.02799 |
| 1334 | UK2014 | A-1.2 | A-322 | 0.39274 | 0.01310 |
| 1335 | UK2014 | A-293 | A-322 | 0.19523 | 0.01089 |
| 1336 | UK2014 | A-1.2 | A-421 | 0.52024 | 0.02771 |
| 1337 | UK2014 | A-293 | A-421 | 0.12095 | 0.01776 |
| 1338 | UK2014 | A-322 | A-421 | 0.85291 | 0.00734 |
| 1339 | UK2014 | A-1.2 | A-498 | 0.66310 | 0.02709 |
| 1340 | UK2014 | A-293 | A-498 | 0.04968 | 0.01276 |
| 1341 | UK2014 | A-322 | A-498 | 0.28847 | 0.01493 |

| | | | | | |
|---|---|---|---|---|---|
| 1342 | UK2014 | A-421 | A-498 | 0.09838 | 0.01738 |
| 1343 | UK2014 | A-1.2 | A-54 | 0.07558 | 0.01184 |
| 1344 | UK2014 | A-293 | A-54 | 0.09956 | 0.01529 |
| 1345 | UK2014 | A-322 | A-54 | 0.58269 | 0.00989 |
| 1346 | UK2014 | A-421 | A-54 | 0.24549 | 0.02135 |
| 1347 | UK2014 | A-498 | A-54 | 0.54525 | 0.02869 |
| 1348 | UK2014 | A-1.2 | A-275 | 0.03009 | 0.00616 |
| 1349 | UK2014 | A-293 | A-275 | 0.13848 | 0.01458 |
| 1350 | UK2014 | A-322 | A-275 | 0.00053 | 0.00050 |
| 1351 | UK2014 | A-421 | A-275 | 0.40997 | 0.02356 |
| 1352 | UK2014 | A-498 | A-275 | 0.27389 | 0.02338 |
| 1353 | UK2014 | A-54 | A-275 | 0.01155 | 0.00308 |
| 1354 | UK2014 | A-1.2 | A-509 | 0.54184 | 0.02863 |
| 1355 | UK2014 | A-293 | A-509 | 0.72553 | 0.02478 |
| 1356 | UK2014 | A-322 | A-509 | 0.80307 | 0.01082 |
| 1357 | UK2014 | A-421 | A-509 | 0.79107 | 0.02602 |
| 1358 | UK2014 | A-498 | A-509 | 0.62422 | 0.03178 |
| 1359 | UK2014 | A-54 | A-509 | 0.08076 | 0.01178 |
| 1360 | UK2014 | A-275 | A-509 | 0.65372 | 0.02406 |
| 1361 | UK2014 | A-1.2 | A-600 | 0.21350 | 0.01101 |
| 1362 | UK2014 | A-293 | A-600 | 0.69974 | 0.01097 |
| 1363 | UK2014 | A-322 | A-600 | 0.49409 | 0.00607 |
| 1364 | UK2014 | A-421 | A-600 | 0.95163 | 0.00440 |
| 1365 | UK2014 | A-498 | A-600 | 0.52525 | 0.01643 |
| 1366 | UK2014 | A-54 | A-600 | 0.18284 | 0.00918 |
| 1367 | UK2014 | A-275 | A-600 | 0.81077 | 0.00745 |
| 1368 | UK2014 | A-509 | A-600 | 0.65079 | 0.01470 |
| 1369 | UK2014 | A-1.2 | A-04 | 0.09724 | 0.01014 |
| 1370 | UK2014 | A-293 | A-04 | 0.82548 | 0.01456 |
| 1371 | UK2014 | A-322 | A-04 | 0.26427 | 0.00968 |
| 1372 | UK2014 | A-421 | A-04 | 0.96774 | 0.00578 |
| 1373 | UK2014 | A-498 | A-04 | 0.55611 | 0.02405 |
| 1374 | UK2014 | A-54 | A-04 | 0.21997 | 0.01589 |
| 1375 | UK2014 | A-275 | A-04 | 0.57139 | 0.01645 |
| 1376 | UK2014 | A-509 | A-04 | 0.41781 | 0.02476 |
| 1377 | UK2014 | A-600 | A-04 | 0.52795 | 0.01144 |
| 1378 | UK2014 | A-1.2 | A-092 | 0.12974 | 0.00714 |
| 1379 | UK2014 | A-293 | A-092 | 0.98394 | 0.00206 |
| 1380 | UK2014 | A-322 | A-092 | 0.68003 | 0.00401 |
| 1381 | UK2014 | A-421 | A-092 | 0.54861 | 0.01093 |
| 1382 | UK2014 | A-498 | A-092 | 0.19088 | 0.01143 |
| 1383 | UK2014 | A-54 | A-092 | 0.18125 | 0.00767 |
| 1384 | UK2014 | A-275 | A-092 | 0.16633 | 0.00564 |
| 1385 | UK2014 | A-509 | A-092 | 0.21936 | 0.01075 |
| 1386 | UK2014 | A-600 | A-092 | 0.54733 | 0.00582 |
| 1387 | UK2014 | A-04 | A-092 | 0.39985 | 0.00856 |
| 1388 | UK2014 | A-1.2 | A-244 | 0.36467 | 0.02773 |
| 1389 | UK2014 | A-293 | A-244 | 0.42801 | 0.02712 |

| | | | | | |
|---|---|---|---|---|---|
| 1390 | UK2014 | A-322 | A-244 | 0.13367 | 0.00761 |
| 1391 | UK2014 | A-421 | A-244 | 0.18399 | 0.01952 |
| 1392 | UK2014 | A-498 | A-244 | 0.02917 | 0.00857 |
| 1393 | UK2014 | A-54 | A-244 | 0.62206 | 0.02178 |
| 1394 | UK2014 | A-275 | A-244 | 0.10831 | 0.01164 |
| 1395 | UK2014 | A-509 | A-244 | 0.45384 | 0.02788 |
| 1396 | UK2014 | A-600 | A-244 | 0.62338 | 0.01139 |
| 1397 | UK2014 | A-04 | A-244 | 0.10281 | 0.01163 |
| 1398 | UK2014 | A-092 | A-244 | 0.43777 | 0.01124 |
| 1399 | UK2014 | A-1.2 | A-342 | 0.86395 | 0.01322 |
| 1400 | UK2014 | A-293 | A-342 | 0.82972 | 0.01716 |
| 1401 | UK2014 | A-322 | A-342 | 0.50990 | 0.01091 |
| 1402 | UK2014 | A-421 | A-342 | 0.76383 | 0.02018 |
| 1403 | UK2014 | A-498 | A-342 | 0.31493 | 0.02612 |
| 1404 | UK2014 | A-54 | A-342 | 0.68364 | 0.01844 |
| 1405 | UK2014 | A-275 | A-342 | 0.57498 | 0.01862 |
| 1406 | UK2014 | A-509 | A-342 | 0.49451 | 0.02675 |
| 1407 | UK2014 | A-600 | A-342 | 0.25991 | 0.01110 |
| 1408 | UK2014 | A-04 | A-342 | 0.10355 | 0.01043 |
| 1409 | UK2014 | A-092 | A-342 | 0.28178 | 0.00962 |
| 1410 | UK2014 | A-244 | A-342 | 0.50830 | 0.02442 |
| 1411 | UK2014 | A-1.2 | A-389 | 0.47376 | 0.02327 |
| 1412 | UK2014 | A-293 | A-389 | 0.20141 | 0.01713 |
| 1413 | UK2014 | A-322 | A-389 | 0.59456 | 0.00905 |
| 1414 | UK2014 | A-421 | A-389 | 0.17884 | 0.01702 |
| 1415 | UK2014 | A-498 | A-389 | 0.48371 | 0.02131 |
| 1416 | UK2014 | A-54 | A-389 | 0.54883 | 0.01955 |
| 1417 | UK2014 | A-275 | A-389 | 0.94396 | 0.00584 |
| 1418 | UK2014 | A-509 | A-389 | 0.75801 | 0.01858 |
| 1419 | UK2014 | A-600 | A-389 | 0.50769 | 0.01140 |
| 1420 | UK2014 | A-04 | A-389 | 0.98804 | 0.00213 |
| 1421 | UK2014 | A-092 | A-389 | 0.57599 | 0.00864 |
| 1422 | UK2014 | A-244 | A-389 | 0.14921 | 0.01250 |
| 1423 | UK2014 | A-342 | A-389 | 0.15935 | 0.01182 |
| 1424 | UK2014 | A-1.2 | A-547 | 0.26540 | 0.02870 |
| 1425 | UK2014 | A-293 | A-547 | 0.30883 | 0.02685 |
| 1426 | UK2014 | A-322 | A-547 | 0.36803 | 0.01570 |
| 1427 | UK2014 | A-421 | A-547 | 0.62115 | 0.03310 |
| 1428 | UK2014 | A-498 | A-547 | 0.46051 | 0.03615 |
| 1429 | UK2014 | A-54 | A-547 | 0.60822 | 0.02654 |
| 1430 | UK2014 | A-275 | A-547 | 0.42128 | 0.02272 |
| 1431 | UK2014 | A-509 | A-547 | 0.09047 | 0.01811 |
| 1432 | UK2014 | A-600 | A-547 | 0.55756 | 0.01598 |
| 1433 | UK2014 | A-04 | A-547 | 0.60071 | 0.02256 |
| 1434 | UK2014 | A-092 | A-547 | 0.72710 | 0.00940 |
| 1435 | UK2014 | A-244 | A-547 | 0.47195 | 0.02719 |
| 1436 | UK2014 | A-342 | A-547 | 0.15293 | 0.02023 |
| 1437 | UK2014 | A-389 | A-547 | 0.09209 | 0.01599 |

| | | | | | |
|---|---|---|---|---|---|
| 1438 | UK2014 | A-1.2 | A-196 | 0.12424 | 0.02496 |
| 1439 | UK2014 | A-293 | A-196 | 0.55275 | 0.03596 |
| 1440 | UK2014 | A-322 | A-196 | 0.19384 | 0.01532 |
| 1441 | UK2014 | A-421 | A-196 | 0.37562 | 0.03547 |
| 1442 | UK2014 | A-498 | A-196 | 0.18761 | 0.03020 |
| 1443 | UK2014 | A-54 | A-196 | 0.49113 | 0.03278 |
| 1444 | UK2014 | A-275 | A-196 | 0.31630 | 0.02787 |
| 1445 | UK2014 | A-509 | A-196 | 0.46802 | 0.03645 |
| 1446 | UK2014 | A-600 | A-196 | 0.97877 | 0.00374 |
| 1447 | UK2014 | A-04 | A-196 | 0.79651 | 0.02173 |
| 1448 | UK2014 | A-092 | A-196 | 0.48996 | 0.01990 |
| 1449 | UK2014 | A-244 | A-196 | 0.32731 | 0.03035 |
| 1450 | UK2014 | A-342 | A-196 | 0.47567 | 0.03773 |
| 1451 | UK2014 | A-389 | A-196 | 0.42618 | 0.02996 |
| 1452 | UK2014 | A-547 | A-196 | 0.94423 | 0.01605 |
| 1453 | UK2014 | A-1.2 | A-360 | 0.90043 | 0.01240 |
| 1454 | UK2014 | A-293 | A-360 | 0.79854 | 0.02132 |
| 1455 | UK2014 | A-322 | A-360 | 0.29045 | 0.01066 |
| 1456 | UK2014 | A-421 | A-360 | 0.95729 | 0.00867 |
| 1457 | UK2014 | A-498 | A-360 | 0.53187 | 0.02915 |
| 1458 | UK2014 | A-54 | A-360 | 0.86861 | 0.01349 |
| 1459 | UK2014 | A-275 | A-360 | 0.91403 | 0.00949 |
| 1460 | UK2014 | A-509 | A-360 | 0.66603 | 0.02329 |
| 1461 | UK2014 | A-600 | A-360 | 0.57797 | 0.01053 |
| 1462 | UK2014 | A-04 | A-360 | 0.29479 | 0.01640 |
| 1463 | UK2014 | A-092 | A-360 | 0.95745 | 0.00288 |
| 1464 | UK2014 | A-244 | A-360 | 0.51183 | 0.02377 |
| 1465 | UK2014 | A-342 | A-360 | 0.49531 | 0.01992 |
| 1466 | UK2014 | A-389 | A-360 | 0.94215 | 0.00749 |
| 1467 | UK2014 | A-547 | A-360 | 0.29953 | 0.02567 |
| 1468 | UK2014 | A-196 | A-360 | 0.08918 | 0.01859 |
| 1469 | UK2014 | A-1.2 | A-465 | 0.51309 | 0.03441 |
| 1470 | UK2014 | A-293 | A-465 | 0.08929 | 0.01692 |
| 1471 | UK2014 | A-322 | A-465 | 0.41692 | 0.01533 |
| 1472 | UK2014 | A-421 | A-465 | 0.89489 | 0.01950 |
| 1473 | UK2014 | A-498 | A-465 | 0.28283 | 0.03299 |
| 1474 | UK2014 | A-54 | A-465 | 0.88072 | 0.01870 |
| 1475 | UK2014 | A-275 | A-465 | 0.44130 | 0.02300 |
| 1476 | UK2014 | A-509 | A-465 | 0.63533 | 0.03032 |
| 1477 | UK2014 | A-600 | A-465 | 0.54801 | 0.01649 |
| 1478 | UK2014 | A-04 | A-465 | 0.37832 | 0.02289 |
| 1479 | UK2014 | A-092 | A-465 | 0.18159 | 0.00903 |
| 1480 | UK2014 | A-244 | A-465 | 0.26658 | 0.02427 |
| 1481 | UK2014 | A-342 | A-465 | 0.47847 | 0.02852 |
| 1482 | UK2014 | A-389 | A-465 | 0.46296 | 0.02501 |
| 1483 | UK2014 | A-547 | A-465 | 0.79455 | 0.03199 |
| 1484 | UK2014 | A-196 | A-465 | 0.25525 | 0.03431 |
| 1485 | UK2014 | A-360 | A-465 | 0.84918 | 0.01845 |

| | | | | | |
|---|---|---|---|---|---|
| 1486 | UK2014 | A-1.2 | A-161 | 0.41025 | 0.03305 |
| 1487 | UK2014 | A-293 | A-161 | 0.84842 | 0.02340 |
| 1488 | UK2014 | A-322 | A-161 | 0.70012 | 0.01343 |
| 1489 | UK2014 | A-421 | A-161 | 0.96567 | 0.01006 |
| 1490 | UK2014 | A-498 | A-161 | 0.97784 | 0.00722 |
| 1491 | UK2014 | A-54 | A-161 | 0.64499 | 0.02752 |
| 1492 | UK2014 | A-275 | A-161 | 0.59750 | 0.02679 |
| 1493 | UK2014 | A-509 | A-161 | 0.88197 | 0.02027 |
| 1494 | UK2014 | A-600 | A-161 | 0.96899 | 0.00435 |
| 1495 | UK2014 | A-04 | A-161 | 0.71099 | 0.02297 |
| 1496 | UK2014 | A-092 | A-161 | 0.71794 | 0.01134 |
| 1497 | UK2014 | A-244 | A-161 | 0.43914 | 0.03180 |
| 1498 | UK2014 | A-342 | A-161 | 0.57919 | 0.02857 |
| 1499 | UK2014 | A-389 | A-161 | 0.29734 | 0.02422 |
| 1500 | UK2014 | A-547 | A-161 | 0.91423 | 0.01789 |
| 1501 | UK2014 | A-196 | A-161 | 0.17084 | 0.02963 |
| 1502 | UK2014 | A-360 | A-161 | 0.80911 | 0.02314 |
| 1503 | UK2014 | A-465 | A-161 | 0.37815 | 0.03591 |
| 1504 | UK2014 | A-1.2 | A-200 | 0.82923 | 0.02588 |
| 1505 | UK2014 | A-293 | A-200 | 0.45056 | 0.03621 |
| 1506 | UK2014 | A-322 | A-200 | 0.04220 | 0.00580 |
| 1507 | UK2014 | A-421 | A-200 | 0.02868 | 0.01298 |
| 1508 | UK2014 | A-498 | A-200 | 0.89366 | 0.02365 |
| 1509 | UK2014 | A-54 | A-200 | 0.63101 | 0.03175 |
| 1510 | UK2014 | A-275 | A-200 | 0.00117 | 0.00117 |
| 1511 | UK2014 | A-509 | A-200 | 0.68196 | 0.03144 |
| 1512 | UK2014 | A-600 | A-200 | 0.35040 | 0.01924 |
| 1513 | UK2014 | A-04 | A-200 | 0.26043 | 0.02480 |
| 1514 | UK2014 | A-092 | A-200 | 0.90291 | 0.00670 |
| 1515 | UK2014 | A-244 | A-200 | 0.31449 | 0.02954 |
| 1516 | UK2014 | A-342 | A-200 | 0.15810 | 0.02032 |
| 1517 | UK2014 | A-389 | A-200 | 0.55489 | 0.02767 |
| 1518 | UK2014 | A-547 | A-200 | 0.07168 | 0.01800 |
| 1519 | UK2014 | A-196 | A-200 | 0.66108 | 0.04064 |
| 1520 | UK2014 | A-360 | A-200 | 0.03652 | 0.00891 |
| 1521 | UK2014 | A-465 | A-200 | 0.44700 | 0.03684 |
| 1522 | UK2014 | A-161 | A-200 | 0.81496 | 0.03017 |
| 1523 | UK2014 | A-1.2 | A-325 | 0.13277 | 0.01885 |
| 1524 | UK2014 | A-293 | A-325 | 0.18673 | 0.02137 |
| 1525 | UK2014 | A-322 | A-325 | 0.73142 | 0.01154 |
| 1526 | UK2014 | A-421 | A-325 | 0.83062 | 0.02225 |
| 1527 | UK2014 | A-498 | A-325 | 0.39115 | 0.03035 |
| 1528 | UK2014 | A-54 | A-325 | 0.00612 | 0.00207 |
| 1529 | UK2014 | A-275 | A-325 | 0.25649 | 0.01915 |
| 1530 | UK2014 | A-509 | A-325 | 0.99023 | 0.00397 |
| 1531 | UK2014 | A-600 | A-325 | 0.58968 | 0.01334 |
| 1532 | UK2014 | A-04 | A-325 | 0.12246 | 0.01234 |
| 1533 | UK2014 | A-092 | A-325 | 0.51868 | 0.01173 |

| | | | | | |
|---|---|---|---|---|---|
| 1534 | UK2014 | A-244 | A-325 | 0.88828 | 0.01393 |
| 1535 | UK2014 | A-342 | A-325 | 0.82828 | 0.01711 |
| 1536 | UK2014 | A-389 | A-325 | 0.30058 | 0.02042 |
| 1537 | UK2014 | A-547 | A-325 | 0.73567 | 0.02440 |
| 1538 | UK2014 | A-196 | A-325 | 0.85425 | 0.02477 |
| 1539 | UK2014 | A-360 | A-325 | 0.72684 | 0.02344 |
| 1540 | UK2014 | A-465 | A-325 | 0.54992 | 0.03327 |
| 1541 | UK2014 | A-161 | A-325 | 0.06473 | 0.01436 |
| 1542 | UK2014 | A-200 | A-325 | 0.12929 | 0.02243 |
| 1543 | UK2014 | A-1.2 | A-333 | 0.16597 | 0.01844 |
| 1544 | UK2014 | A-293 | A-333 | 0.57559 | 0.02660 |
| 1545 | UK2014 | A-322 | A-333 | 0.06332 | 0.00545 |
| 1546 | UK2014 | A-421 | A-333 | 0.86626 | 0.01662 |
| 1547 | UK2014 | A-498 | A-333 | 0.15187 | 0.02000 |
| 1548 | UK2014 | A-54 | A-333 | 0.75829 | 0.01721 |
| 1549 | UK2014 | A-275 | A-333 | 0.36147 | 0.01810 |
| 1550 | UK2014 | A-509 | A-333 | 0.66747 | 0.02329 |
| 1551 | UK2014 | A-600 | A-333 | 0.68783 | 0.01178 |
| 1552 | UK2014 | A-04 | A-333 | 0.03305 | 0.00702 |
| 1553 | UK2014 | A-092 | A-333 | 0.55430 | 0.01079 |
| 1554 | UK2014 | A-244 | A-333 | 0.46182 | 0.02469 |
| 1555 | UK2014 | A-342 | A-333 | 0.43936 | 0.02095 |
| 1556 | UK2014 | A-389 | A-333 | 0.96143 | 0.00508 |
| 1557 | UK2014 | A-547 | A-333 | 0.72729 | 0.02316 |
| 1558 | UK2014 | A-196 | A-333 | 0.78683 | 0.02751 |
| 1559 | UK2014 | A-360 | A-333 | 0.81217 | 0.01540 |
| 1560 | UK2014 | A-465 | A-333 | 0.60228 | 0.02754 |
| 1561 | UK2014 | A-161 | A-333 | 0.67047 | 0.02859 |
| 1562 | UK2014 | A-200 | A-333 | 0.32070 | 0.02821 |
| 1563 | UK2014 | A-325 | A-333 | 0.65329 | 0.02301 |
| 1564 | UK2014 | A-1.2 | A-104.2 | 0.88143 | 0.01038 |
| 1565 | UK2014 | A-293 | A-104.2 | 0.91179 | 0.00989 |
| 1566 | UK2014 | A-322 | A-104.2 | 0.83943 | 0.00531 |
| 1567 | UK2014 | A-421 | A-104.2 | 0.26661 | 0.01687 |
| 1568 | UK2014 | A-498 | A-104.2 | 0.77655 | 0.01808 |
| 1569 | UK2014 | A-54 | A-104.2 | 0.43564 | 0.02057 |
| 1570 | UK2014 | A-275 | A-104.2 | 0.13323 | 0.01094 |
| 1571 | UK2014 | A-509 | A-104.2 | 0.06552 | 0.00903 |
| 1572 | UK2014 | A-600 | A-104.2 | 0.51375 | 0.01129 |
| 1573 | UK2014 | A-04 | A-104.2 | 0.18816 | 0.01205 |
| 1574 | UK2014 | A-092 | A-104.2 | 0.19929 | 0.00663 |
| 1575 | UK2014 | A-244 | A-104.2 | 0.72658 | 0.01247 |
| 1576 | UK2014 | A-342 | A-104.2 | 0.44117 | 0.01854 |
| 1577 | UK2014 | A-389 | A-104.2 | 0.60851 | 0.01515 |
| 1578 | UK2014 | A-547 | A-104.2 | 0.05362 | 0.00805 |
| 1579 | UK2014 | A-196 | A-104.2 | 0.79493 | 0.02273 |
| 1580 | UK2014 | A-360 | A-104.2 | 0.60807 | 0.01495 |
| 1581 | UK2014 | A-465 | A-104.2 | 0.21925 | 0.01849 |

| | | | | | |
|---|---|---|---|---|---|
| 1582 | UK2014 | A-161 | A-104.2 | 0.22523 | 0.02061 |
| 1583 | UK2014 | A-200 | A-104.2 | 0.41350 | 0.02560 |
| 1584 | UK2014 | A-325 | A-104.2 | 0.66428 | 0.02003 |
| 1585 | UK2014 | A-333 | A-104.2 | 0.44531 | 0.01679 |
| 1586 | UK2014 | A-1.2 | A-122 | 0.03480 | 0.01123 |
| 1587 | UK2014 | A-293 | A-122 | 0.95215 | 0.01149 |
| 1588 | UK2014 | A-322 | A-122 | 0.00000 | 0.00000 |
| 1589 | UK2014 | A-421 | A-122 | 0.44152 | 0.03400 |
| 1590 | UK2014 | A-498 | A-122 | 0.77278 | 0.02751 |
| 1591 | UK2014 | A-54 | A-122 | 0.33400 | 0.02378 |
| 1592 | UK2014 | A-275 | A-122 | 0.00000 | 0.00000 |
| 1593 | UK2014 | A-509 | A-122 | 0.68806 | 0.02965 |
| 1594 | UK2014 | A-600 | A-122 | 0.25777 | 0.01365 |
| 1595 | UK2014 | A-04 | A-122 | 0.02246 | 0.00582 |
| 1596 | UK2014 | A-092 | A-122 | 0.63917 | 0.01242 |
| 1597 | UK2014 | A-244 | A-122 | 0.38085 | 0.02794 |
| 1598 | UK2014 | A-342 | A-122 | 0.30818 | 0.02605 |
| 1599 | UK2014 | A-389 | A-122 | 0.57683 | 0.02527 |
| 1600 | UK2014 | A-547 | A-122 | 0.68762 | 0.03118 |
| 1601 | UK2014 | A-196 | A-122 | 0.63834 | 0.03587 |
| 1602 | UK2014 | A-360 | A-122 | 0.36431 | 0.02832 |
| 1603 | UK2014 | A-465 | A-122 | 0.34989 | 0.03154 |
| 1604 | UK2014 | A-161 | A-122 | 0.99407 | 0.00273 |
| 1605 | UK2014 | A-200 | A-122 | 0.00000 | 0.00000 |
| 1606 | UK2014 | A-325 | A-122 | 0.42095 | 0.03494 |
| 1607 | UK2014 | A-333 | A-122 | 0.59321 | 0.02720 |
| 1608 | UK2014 | A-104.2 | A-122 | 0.68179 | 0.01961 |
| 1609 | UK2014 | A-1.2 | A-175 | 0.81674 | 0.01531 |
| 1610 | UK2014 | A-293 | A-175 | 0.54264 | 0.02355 |
| 1611 | UK2014 | A-322 | A-175 | 0.96077 | 0.00250 |
| 1612 | UK2014 | A-421 | A-175 | 0.65299 | 0.01967 |
| 1613 | UK2014 | A-498 | A-175 | 0.44486 | 0.02443 |
| 1614 | UK2014 | A-54 | A-175 | 0.01426 | 0.00366 |
| 1615 | UK2014 | A-275 | A-175 | 0.91230 | 0.00860 |
| 1616 | UK2014 | A-509 | A-175 | 0.05603 | 0.00969 |
| 1617 | UK2014 | A-600 | A-175 | 0.07130 | 0.00529 |
| 1618 | UK2014 | A-04 | A-175 | 0.36848 | 0.01416 |
| 1619 | UK2014 | A-092 | A-175 | 0.49260 | 0.00839 |
| 1620 | UK2014 | A-244 | A-175 | 0.88066 | 0.01170 |
| 1621 | UK2014 | A-342 | A-175 | 0.61294 | 0.01854 |
| 1622 | UK2014 | A-389 | A-175 | 0.39441 | 0.01474 |
| 1623 | UK2014 | A-547 | A-175 | 0.77247 | 0.01879 |
| 1624 | UK2014 | A-196 | A-175 | 0.34576 | 0.03005 |
| 1625 | UK2014 | A-360 | A-175 | 0.15071 | 0.01373 |
| 1626 | UK2014 | A-465 | A-175 | 0.59082 | 0.02544 |
| 1627 | UK2014 | A-161 | A-175 | 0.55508 | 0.02379 |
| 1628 | UK2014 | A-200 | A-175 | 0.07987 | 0.01502 |
| 1629 | UK2014 | A-325 | A-175 | 0.69877 | 0.01950 |

| | | | | | |
|---|---|---|---|---|---|
| 1630 | UK2014 | A-333 | A-175 | 0.62349 | 0.01669 |
| 1631 | UK2014 | A-104.2 | A-175 | 0.55514 | 0.01570 |
| 1632 | UK2014 | A-122 | A-175 | 0.32154 | 0.02111 |
| 1633 | UK2014 | A-1.2 | A-300 | 0.89537 | 0.01798 |
| 1634 | UK2014 | A-293 | A-300 | 0.77205 | 0.02723 |
| 1635 | UK2014 | A-322 | A-300 | 0.20419 | 0.01295 |
| 1636 | UK2014 | A-421 | A-300 | 0.92590 | 0.01657 |
| 1637 | UK2014 | A-498 | A-300 | 0.01442 | 0.00686 |
| 1638 | UK2014 | A-54 | A-300 | 0.07294 | 0.01789 |
| 1639 | UK2014 | A-275 | A-300 | 0.90540 | 0.01572 |
| 1640 | UK2014 | A-509 | A-300 | 0.57033 | 0.03561 |
| 1641 | UK2014 | A-600 | A-300 | 0.94992 | 0.00669 |
| 1642 | UK2014 | A-04 | A-300 | 0.59928 | 0.02543 |
| 1643 | UK2014 | A-092 | A-300 | 0.94359 | 0.00526 |
| 1644 | UK2014 | A-244 | A-300 | 0.91614 | 0.01584 |
| 1645 | UK2014 | A-342 | A-300 | 0.24889 | 0.02923 |
| 1646 | UK2014 | A-389 | A-300 | 0.73543 | 0.02102 |
| 1647 | UK2014 | A-547 | A-300 | 0.92327 | 0.01922 |
| 1648 | UK2014 | A-196 | A-300 | 0.37252 | 0.04054 |
| 1649 | UK2014 | A-360 | A-300 | 0.49652 | 0.03449 |
| 1650 | UK2014 | A-465 | A-300 | 0.64590 | 0.03606 |
| 1651 | UK2014 | A-161 | A-300 | 0.47355 | 0.04073 |
| 1652 | UK2014 | A-200 | A-300 | 0.85327 | 0.02808 |
| 1653 | UK2014 | A-325 | A-300 | 0.20428 | 0.02768 |
| 1654 | UK2014 | A-333 | A-300 | 0.53856 | 0.03219 |
| 1655 | UK2014 | A-104.2 | A-300 | 0.73833 | 0.02062 |
| 1656 | UK2014 | A-122 | A-300 | 0.26240 | 0.03275 |
| 1657 | UK2014 | A-175 | A-300 | 0.79952 | 0.02292 |
| 1658 | UK2014 | A-1.2 | A-733 | 0.54939 | 0.00773 |
| 1659 | UK2014 | A-293 | A-733 | 0.77585 | 0.00736 |
| 1660 | UK2014 | A-322 | A-733 | 0.34700 | 0.00391 |
| 1661 | UK2014 | A-421 | A-733 | 1.00000 | 0.00000 |
| 1662 | UK2014 | A-498 | A-733 | 0.50674 | 0.01237 |
| 1663 | UK2014 | A-54 | A-733 | 0.08533 | 0.00403 |
| 1664 | UK2014 | A-275 | A-733 | 1.00000 | 0.00000 |
| 1665 | UK2014 | A-509 | A-733 | 1.00000 | 0.00000 |
| 1666 | UK2014 | A-600 | A-733 | 0.28744 | 0.00380 |
| 1667 | UK2014 | A-04 | A-733 | 1.00000 | 0.00000 |
| 1668 | UK2014 | A-092 | A-733 | 1.00000 | 0.00000 |
| 1669 | UK2014 | A-244 | A-733 | 1.00000 | 0.00000 |
| 1670 | UK2014 | A-342 | A-733 | 0.77977 | 0.00612 |
| 1671 | UK2014 | A-389 | A-733 | 0.46933 | 0.00621 |
| 1672 | UK2014 | A-547 | A-733 | 0.73181 | 0.00960 |
| 1673 | UK2014 | A-196 | A-733 | 0.11931 | 0.00748 |
| 1674 | UK2014 | A-360 | A-733 | 0.43518 | 0.01059 |
| 1675 | UK2014 | A-465 | A-733 | 0.08256 | 0.00580 |
| 1676 | UK2014 | A-161 | A-733 | 0.01053 | 0.00148 |
| 1677 | UK2014 | A-200 | A-733 | 0.08358 | 0.00588 |

| | | | | | |
|---|---|---|---|---|---|
| 1678 | UK2014 | A-325 | A-733 | 0.00388 | 0.00075 |
| 1679 | UK2014 | A-333 | A-733 | 0.55021 | 0.00844 |
| 1680 | UK2014 | A-104.2 | A-733 | 1.00000 | 0.00000 |
| 1681 | UK2014 | A-122 | A-733 | 1.00000 | 0.00000 |
| 1682 | UK2014 | A-175 | A-733 | 0.24121 | 0.00624 |
| 1683 | UK2014 | A-300 | A-733 | 0.17308 | 0.00935 |
| 1684 | UK2014 | A-1.2 | A-0 | 0.59675 | 0.02873 |
| 1685 | UK2014 | A-293 | A-0 | 0.97114 | 0.00538 |
| 1686 | UK2014 | A-322 | A-0 | 0.53640 | 0.01315 |
| 1687 | UK2014 | A-421 | A-0 | 0.81840 | 0.02203 |
| 1688 | UK2014 | A-498 | A-0 | 0.41879 | 0.02851 |
| 1689 | UK2014 | A-54 | A-0 | 0.40693 | 0.02312 |
| 1690 | UK2014 | A-275 | A-0 | 0.00010 | 0.00010 |
| 1691 | UK2014 | A-509 | A-0 | 0.72887 | 0.02487 |
| 1692 | UK2014 | A-600 | A-0 | 0.89284 | 0.00693 |
| 1693 | UK2014 | A-04 | A-0 | 0.37417 | 0.02133 |
| 1694 | UK2014 | A-092 | A-0 | 0.10435 | 0.00698 |
| 1695 | UK2014 | A-244 | A-0 | 0.77549 | 0.01980 |
| 1696 | UK2014 | A-342 | A-0 | 0.21551 | 0.02202 |
| 1697 | UK2014 | A-389 | A-0 | 0.92071 | 0.00919 |
| 1698 | UK2014 | A-547 | A-0 | 0.31941 | 0.02871 |
| 1699 | UK2014 | A-196 | A-0 | 0.48996 | 0.03915 |
| 1700 | UK2014 | A-360 | A-0 | 0.03272 | 0.00724 |
| 1701 | UK2014 | A-465 | A-0 | 0.90103 | 0.01457 |
| 1702 | UK2014 | A-161 | A-0 | 0.61257 | 0.03088 |
| 1703 | UK2014 | A-200 | A-0 | 0.01331 | 0.00644 |
| 1704 | UK2014 | A-325 | A-0 | 0.17872 | 0.02474 |
| 1705 | UK2014 | A-333 | A-0 | 0.42622 | 0.02836 |
| 1706 | UK2014 | A-104.2 | A-0 | 0.67595 | 0.01902 |
| 1707 | UK2014 | A-122 | A-0 | 0.00000 | 0.00000 |
| 1708 | UK2014 | A-175 | A-0 | 0.10814 | 0.01515 |
| 1709 | UK2014 | A-300 | A-0 | 0.07920 | 0.01952 |
| 1710 | UK2014 | A-733 | A-0 | 0.29046 | 0.00906 |
| 1711 | UK2014 | A-1.2 | A-1.1 | 0.31326 | 0.02618 |
| 1712 | UK2014 | A-293 | A-1.1 | 0.19194 | 0.02156 |
| 1713 | UK2014 | A-322 | A-1.1 | 0.68318 | 0.00954 |
| 1714 | UK2014 | A-421 | A-1.1 | 0.41757 | 0.02768 |
| 1715 | UK2014 | A-498 | A-1.1 | 0.62845 | 0.02743 |
| 1716 | UK2014 | A-54 | A-1.1 | 0.48600 | 0.02311 |
| 1717 | UK2014 | A-275 | A-1.1 | 0.03983 | 0.00754 |
| 1718 | UK2014 | A-509 | A-1.1 | 0.19887 | 0.02034 |
| 1719 | UK2014 | A-600 | A-1.1 | 0.53095 | 0.01228 |
| 1720 | UK2014 | A-04 | A-1.1 | 0.36887 | 0.01715 |
| 1721 | UK2014 | A-092 | A-1.1 | 0.38294 | 0.00931 |
| 1722 | UK2014 | A-244 | A-1.1 | 0.22299 | 0.01861 |
| 1723 | UK2014 | A-342 | A-1.1 | 0.28714 | 0.02297 |
| 1724 | UK2014 | A-389 | A-1.1 | 0.01412 | 0.00366 |
| 1725 | UK2014 | A-547 | A-1.1 | 0.59176 | 0.02506 |

| | | | | | |
|---|---|---|---|---|---|
| 1726 | UK2014 | A-196 | A-1.1 | 0.66978 | 0.03193 |
| 1727 | UK2014 | A-360 | A-1.1 | 0.55234 | 0.02231 |
| 1728 | UK2014 | A-465 | A-1.1 | 0.01680 | 0.00535 |
| 1729 | UK2014 | A-161 | A-1.1 | 0.78469 | 0.02478 |
| 1730 | UK2014 | A-200 | A-1.1 | 0.66988 | 0.02853 |
| 1731 | UK2014 | A-325 | A-1.1 | 0.25508 | 0.02606 |
| 1732 | UK2014 | A-333 | A-1.1 | 0.37999 | 0.02101 |
| 1733 | UK2014 | A-104.2 | A-1.1 | 0.12029 | 0.01235 |
| 1734 | UK2014 | A-122 | A-1.1 | 0.91789 | 0.01505 |
| 1735 | UK2014 | A-175 | A-1.1 | 0.10639 | 0.01165 |
| 1736 | UK2014 | A-300 | A-1.1 | 0.92024 | 0.01651 |
| 1737 | UK2014 | A-733 | A-1.1 | 0.84457 | 0.00534 |
| 1738 | UK2014 | A-0 | A-1.1 | 0.28494 | 0.02233 |
| 1739 | UK2014 | A-1.2 | A-450 | 0.83924 | 0.02380 |
| 1740 | UK2014 | A-293 | A-450 | 0.89255 | 0.01934 |
| 1741 | UK2014 | A-322 | A-450 | 0.70913 | 0.01514 |
| 1742 | UK2014 | A-421 | A-450 | 0.93284 | 0.01693 |
| 1743 | UK2014 | A-498 | A-450 | 0.18374 | 0.02791 |
| 1744 | UK2014 | A-54 | A-450 | 0.22756 | 0.02402 |
| 1745 | UK2014 | A-275 | A-450 | 0.47759 | 0.02614 |
| 1746 | UK2014 | A-509 | A-450 | 0.51407 | 0.03228 |
| 1747 | UK2014 | A-600 | A-450 | 0.72259 | 0.01441 |
| 1748 | UK2014 | A-04 | A-450 | 0.43038 | 0.02399 |
| 1749 | UK2014 | A-092 | A-450 | 0.52244 | 0.01284 |
| 1750 | UK2014 | A-244 | A-450 | 0.80263 | 0.02210 |
| 1751 | UK2014 | A-342 | A-450 | 0.76627 | 0.02491 |
| 1752 | UK2014 | A-389 | A-450 | 0.17142 | 0.01698 |
| 1753 | UK2014 | A-547 | A-450 | 0.01612 | 0.00589 |
| 1754 | UK2014 | A-196 | A-450 | 0.70067 | 0.03554 |
| 1755 | UK2014 | A-360 | A-450 | 0.27246 | 0.02787 |
| 1756 | UK2014 | A-465 | A-450 | 0.66148 | 0.03363 |
| 1757 | UK2014 | A-161 | A-450 | 0.26972 | 0.03003 |
| 1758 | UK2014 | A-200 | A-450 | 0.67733 | 0.03578 |
| 1759 | UK2014 | A-325 | A-450 | 0.72186 | 0.02820 |
| 1760 | UK2014 | A-333 | A-450 | 0.47862 | 0.02565 |
| 1761 | UK2014 | A-104.2 | A-450 | 0.10699 | 0.01335 |
| 1762 | UK2014 | A-122 | A-450 | 0.04030 | 0.01091 |
| 1763 | UK2014 | A-175 | A-450 | 0.67831 | 0.02481 |
| 1764 | UK2014 | A-300 | A-450 | 0.13700 | 0.02383 |
| 1765 | UK2014 | A-733 | A-450 | 0.47065 | 0.01218 |
| 1766 | UK2014 | A-0 | A-450 | 0.01030 | 0.00589 |
| 1767 | UK2014 | A-1.1 | A-450 | 0.29367 | 0.02846 |
| 1768 | UK2014 | A-1.2 | A-60 | 0.77785 | 0.02400 |
| 1769 | UK2014 | A-293 | A-60 | 0.31597 | 0.02789 |
| 1770 | UK2014 | A-322 | A-60 | 0.25076 | 0.01267 |
| 1771 | UK2014 | A-421 | A-60 | 0.49181 | 0.03063 |
| 1772 | UK2014 | A-498 | A-60 | 0.85117 | 0.02432 |
| 1773 | UK2014 | A-54 | A-60 | 0.01452 | 0.00705 |

| | | | | | |
|---|---|---|---|---|---|
| 1774 | UK2014 | A-275 | A-60 | 0.00607 | 0.00249 |
| 1775 | UK2014 | A-509 | A-60 | 0.99156 | 0.00440 |
| 1776 | UK2014 | A-600 | A-60 | 0.09377 | 0.00903 |
| 1777 | UK2014 | A-04 | A-60 | 0.45876 | 0.02208 |
| 1778 | UK2014 | A-092 | A-60 | 0.79550 | 0.00969 |
| 1779 | UK2014 | A-244 | A-60 | 0.29777 | 0.02576 |
| 1780 | UK2014 | A-342 | A-60 | 0.11019 | 0.01727 |
| 1781 | UK2014 | A-389 | A-60 | 0.18808 | 0.01937 |
| 1782 | UK2014 | A-547 | A-60 | 0.12658 | 0.02070 |
| 1783 | UK2014 | A-196 | A-60 | 0.36133 | 0.03858 |
| 1784 | UK2014 | A-360 | A-60 | 0.88435 | 0.01600 |
| 1785 | UK2014 | A-465 | A-60 | 0.25522 | 0.02921 |
| 1786 | UK2014 | A-161 | A-60 | 0.74549 | 0.03011 |
| 1787 | UK2014 | A-200 | A-60 | 0.01451 | 0.00946 |
| 1788 | UK2014 | A-325 | A-60 | 0.41414 | 0.02974 |
| 1789 | UK2014 | A-333 | A-60 | 0.32109 | 0.02457 |
| 1790 | UK2014 | A-104.2 | A-60 | 0.60262 | 0.01907 |
| 1791 | UK2014 | A-122 | A-60 | 0.10248 | 0.01840 |
| 1792 | UK2014 | A-175 | A-60 | 0.63570 | 0.02445 |
| 1793 | UK2014 | A-300 | A-60 | 0.84606 | 0.02668 |
| 1794 | UK2014 | A-733 | A-60 | 0.57440 | 0.00997 |
| 1795 | UK2014 | A-0 | A-60 | 0.07783 | 0.01688 |
| 1796 | UK2014 | A-1.1 | A-60 | 0.00000 | 0.00000 |
| 1797 | UK2014 | A-450 | A-60 | 0.93909 | 0.01364 |
| 1798 | UK2014 | A-1.2 | A-662 | 0.35620 | 0.01235 |
| 1799 | UK2014 | A-293 | A-662 | 0.10075 | 0.00777 |
| 1800 | UK2014 | A-322 | A-662 | 0.00000 | 0.00000 |
| 1801 | UK2014 | A-421 | A-662 | 0.54455 | 0.01608 |
| 1802 | UK2014 | A-498 | A-662 | 0.26717 | 0.01211 |
| 1803 | UK2014 | A-54 | A-662 | 0.78246 | 0.00811 |
| 1804 | UK2014 | A-275 | A-662 | 0.00013 | 0.00008 |
| 1805 | UK2014 | A-509 | A-662 | 0.94125 | 0.00441 |
| 1806 | UK2014 | A-600 | A-662 | 0.63300 | 0.00528 |
| 1807 | UK2014 | A-04 | A-662 | 0.28296 | 0.00851 |
| 1808 | UK2014 | A-092 | A-662 | 0.26566 | 0.00375 |
| 1809 | UK2014 | A-244 | A-662 | 0.08989 | 0.00600 |
| 1810 | UK2014 | A-342 | A-662 | 0.55605 | 0.01041 |
| 1811 | UK2014 | A-389 | A-662 | 0.57933 | 0.00956 |
| 1812 | UK2014 | A-547 | A-662 | 0.54911 | 0.01469 |
| 1813 | UK2014 | A-196 | A-662 | 0.36828 | 0.01872 |
| 1814 | UK2014 | A-360 | A-662 | 0.77097 | 0.00774 |
| 1815 | UK2014 | A-465 | A-662 | 0.22619 | 0.01178 |
| 1816 | UK2014 | A-161 | A-662 | 0.18644 | 0.01157 |
| 1817 | UK2014 | A-200 | A-662 | 0.29363 | 0.01571 |
| 1818 | UK2014 | A-325 | A-662 | 0.92834 | 0.00490 |
| 1819 | UK2014 | A-333 | A-662 | 0.51590 | 0.01011 |
| 1820 | UK2014 | A-104.2 | A-662 | 0.84471 | 0.00535 |
| 1821 | UK2014 | A-122 | A-662 | 0.00000 | 0.00000 |

| | | | | | |
|---|---|---|---|---|---|
| 1822 | UK2014 | A-175 | A-662 | 0.88556 | 0.00474 |
| 1823 | UK2014 | A-300 | A-662 | 0.37359 | 0.01608 |
| 1824 | UK2014 | A-733 | A-662 | 0.40593 | 0.00438 |
| 1825 | UK2014 | A-0 | A-662 | 0.62187 | 0.01232 |
| 1826 | UK2014 | A-1.1 | A-662 | 0.72136 | 0.00924 |
| 1827 | UK2014 | A-450 | A-662 | 0.74186 | 0.01263 |
| 1828 | UK2014 | A-60 | A-662 | 0.21374 | 0.01201 |
| 1829 | UK2014 | A-1.2 | A-104.1 | 0.34347 | 0.02518 |
| 1830 | UK2014 | A-293 | A-104.1 | 0.25430 | 0.02384 |
| 1831 | UK2014 | A-322 | A-104.1 | 0.83438 | 0.00893 |
| 1832 | UK2014 | A-421 | A-104.1 | 0.39460 | 0.02728 |
| 1833 | UK2014 | A-498 | A-104.1 | 0.21161 | 0.02206 |
| 1834 | UK2014 | A-54 | A-104.1 | 0.22787 | 0.01883 |
| 1835 | UK2014 | A-275 | A-104.1 | 0.32943 | 0.01797 |
| 1836 | UK2014 | A-509 | A-104.1 | 0.22441 | 0.02157 |
| 1837 | UK2014 | A-600 | A-104.1 | 0.99272 | 0.00114 |
| 1838 | UK2014 | A-04 | A-104.1 | 0.79847 | 0.01310 |
| 1839 | UK2014 | A-092 | A-104.1 | 0.53101 | 0.00911 |
| 1840 | UK2014 | A-244 | A-104.1 | 0.29389 | 0.02268 |
| 1841 | UK2014 | A-342 | A-104.1 | 0.70581 | 0.02010 |
| 1842 | UK2014 | A-389 | A-104.1 | 0.34656 | 0.01798 |
| 1843 | UK2014 | A-547 | A-104.1 | 0.15547 | 0.01913 |
| 1844 | UK2014 | A-196 | A-104.1 | 0.00000 | 0.00000 |
| 1845 | UK2014 | A-360 | A-104.1 | 0.75204 | 0.01872 |
| 1846 | UK2014 | A-465 | A-104.1 | 0.50228 | 0.02766 |
| 1847 | UK2014 | A-161 | A-104.1 | 0.07123 | 0.01400 |
| 1848 | UK2014 | A-200 | A-104.1 | 0.95420 | 0.01199 |
| 1849 | UK2014 | A-325 | A-104.1 | 0.66302 | 0.02768 |
| 1850 | UK2014 | A-333 | A-104.1 | 0.75426 | 0.01748 |
| 1851 | UK2014 | A-104.2 | A-104.1 | 0.42274 | 0.01761 |
| 1852 | UK2014 | A-122 | A-104.1 | 0.84974 | 0.01709 |
| 1853 | UK2014 | A-175 | A-104.1 | 0.33932 | 0.01808 |
| 1854 | UK2014 | A-300 | A-104.1 | 0.65415 | 0.02965 |
| 1855 | UK2014 | A-733 | A-104.1 | 0.42068 | 0.00883 |
| 1856 | UK2014 | A-0 | A-104.1 | 0.82512 | 0.01849 |
| 1857 | UK2014 | A-1.1 | A-104.1 | 0.03546 | 0.00629 |
| 1858 | UK2014 | A-450 | A-104.1 | 0.63965 | 0.02771 |
| 1859 | UK2014 | A-60 | A-104.1 | 0.56196 | 0.02958 |
| 1860 | UK2014 | A-662 | A-104.1 | 0.92355 | 0.00482 |
| 1861 | UK2014 | A-1.2 | A-149 | 0.79833 | 0.02292 |
| 1862 | UK2014 | A-293 | A-149 | 0.76342 | 0.02503 |
| 1863 | UK2014 | A-322 | A-149 | 0.23621 | 0.01159 |
| 1864 | UK2014 | A-421 | A-149 | 0.30478 | 0.02769 |
| 1865 | UK2014 | A-498 | A-149 | 0.55142 | 0.02851 |
| 1866 | UK2014 | A-54 | A-149 | 0.69081 | 0.02235 |
| 1867 | UK2014 | A-275 | A-149 | 0.34994 | 0.02135 |
| 1868 | UK2014 | A-509 | A-149 | 0.93868 | 0.01170 |
| 1869 | UK2014 | A-600 | A-149 | 0.98291 | 0.00231 |

| | | | | | |
|---|---|---|---|---|---|
| 1870 | UK2014 | A-04 | A-149 | 0.24945 | 0.01918 |
| 1871 | UK2014 | A-092 | A-149 | 0.37250 | 0.01481 |
| 1872 | UK2014 | A-244 | A-149 | 0.63047 | 0.02476 |
| 1873 | UK2014 | A-342 | A-149 | 0.88226 | 0.01590 |
| 1874 | UK2014 | A-389 | A-149 | 0.70410 | 0.01988 |
| 1875 | UK2014 | A-547 | A-149 | 0.38876 | 0.02948 |
| 1876 | UK2014 | A-196 | A-149 | 0.45177 | 0.03706 |
| 1877 | UK2014 | A-360 | A-149 | 0.31045 | 0.02407 |
| 1878 | UK2014 | A-465 | A-149 | 0.96316 | 0.01013 |
| 1879 | UK2014 | A-161 | A-149 | 0.37317 | 0.03193 |
| 1880 | UK2014 | A-200 | A-149 | 0.39578 | 0.03395 |
| 1881 | UK2014 | A-325 | A-149 | 0.09412 | 0.01182 |
| 1882 | UK2014 | A-333 | A-149 | 0.73382 | 0.02297 |
| 1883 | UK2014 | A-104.2 | A-149 | 0.32670 | 0.02062 |
| 1884 | UK2014 | A-122 | A-149 | 0.11278 | 0.01780 |
| 1885 | UK2014 | A-175 | A-149 | 0.38083 | 0.02424 |
| 1886 | UK2014 | A-300 | A-149 | 0.29308 | 0.03154 |
| 1887 | UK2014 | A-733 | A-149 | 1.00000 | 0.00000 |
| 1888 | UK2014 | A-0 | A-149 | 0.00000 | 0.00000 |
| 1889 | UK2014 | A-1.1 | A-149 | 0.15211 | 0.01829 |
| 1890 | UK2014 | A-450 | A-149 | 0.02422 | 0.00878 |
| 1891 | UK2014 | A-60 | A-149 | 0.68699 | 0.02818 |
| 1892 | UK2014 | A-662 | A-149 | 0.20145 | 0.01066 |
| 1893 | UK2014 | A-104.1 | A-149 | 0.47384 | 0.02705 |
| 1894 | UK2014 | A-1.2 | A-224 | 0.00782 | 0.00512 |
| 1895 | UK2014 | A-293 | A-224 | 0.87928 | 0.02012 |
| 1896 | UK2014 | A-322 | A-224 | 0.00000 | 0.00000 |
| 1897 | UK2014 | A-421 | A-224 | 0.96949 | 0.00962 |
| 1898 | UK2014 | A-498 | A-224 | 0.74130 | 0.02938 |
| 1899 | UK2014 | A-54 | A-224 | 0.43656 | 0.02904 |
| 1900 | UK2014 | A-275 | A-224 | 0.00000 | 0.00000 |
| 1901 | UK2014 | A-509 | A-224 | 0.59024 | 0.03204 |
| 1902 | UK2014 | A-600 | A-224 | 0.08280 | 0.00853 |
| 1903 | UK2014 | A-04 | A-224 | 0.49685 | 0.02807 |
| 1904 | UK2014 | A-092 | A-224 | 0.57459 | 0.01540 |
| 1905 | UK2014 | A-244 | A-224 | 0.38907 | 0.02945 |
| 1906 | UK2014 | A-342 | A-224 | 0.41896 | 0.03397 |
| 1907 | UK2014 | A-389 | A-224 | 0.04747 | 0.01129 |
| 1908 | UK2014 | A-547 | A-224 | 0.44565 | 0.03497 |
| 1909 | UK2014 | A-196 | A-224 | 0.13944 | 0.02752 |
| 1910 | UK2014 | A-360 | A-224 | 0.69633 | 0.02905 |
| 1911 | UK2014 | A-465 | A-224 | 0.65816 | 0.03478 |
| 1912 | UK2014 | A-161 | A-224 | 0.96209 | 0.01003 |
| 1913 | UK2014 | A-200 | A-224 | 0.03734 | 0.01490 |
| 1914 | UK2014 | A-325 | A-224 | 0.34692 | 0.03081 |
| 1915 | UK2014 | A-333 | A-224 | 0.56683 | 0.02961 |
| 1916 | UK2014 | A-104.2 | A-224 | 0.51651 | 0.02514 |
| 1917 | UK2014 | A-122 | A-224 | 0.00000 | 0.00000 |

| | | | | | |
|---|---|---|---|---|---|
| 1918 | UK2014 | A-175 | A-224 | 0.94670 | 0.00857 |
| 1919 | UK2014 | A-300 | A-224 | 0.78656 | 0.03138 |
| 1920 | UK2014 | A-733 | A-224 | 0.81317 | 0.00992 |
| 1921 | UK2014 | A-0 | A-224 | 0.00407 | 0.00263 |
| 1922 | UK2014 | A-1.1 | A-224 | 0.42845 | 0.03092 |
| 1923 | UK2014 | A-450 | A-224 | 0.53669 | 0.03689 |
| 1924 | UK2014 | A-60 | A-224 | 0.00047 | 0.00047 |
| 1925 | UK2014 | A-662 | A-224 | 0.00000 | 0.00000 |
| 1926 | UK2014 | A-104.1 | A-224 | 0.69507 | 0.02805 |
| 1927 | UK2014 | A-149 | A-224 | 0.37550 | 0.03346 |
| 1928 | UK2014 | A-1.2 | A-323 | 0.25083 | 0.03201 |
| 1929 | UK2014 | A-293 | A-323 | 0.36041 | 0.03513 |
| 1930 | UK2014 | A-322 | A-323 | 0.17263 | 0.01317 |
| 1931 | UK2014 | A-421 | A-323 | 0.85404 | 0.02694 |
| 1932 | UK2014 | A-498 | A-323 | 0.52400 | 0.04186 |
| 1933 | UK2014 | A-54 | A-323 | 0.27177 | 0.03237 |
| 1934 | UK2014 | A-275 | A-323 | 0.79711 | 0.02363 |
| 1935 | UK2014 | A-509 | A-323 | 0.24444 | 0.03393 |
| 1936 | UK2014 | A-600 | A-323 | 0.76930 | 0.01576 |
| 1937 | UK2014 | A-04 | A-323 | 0.23788 | 0.02337 |
| 1938 | UK2014 | A-092 | A-323 | 0.84491 | 0.01004 |
| 1939 | UK2014 | A-244 | A-323 | 0.14170 | 0.02161 |
| 1940 | UK2014 | A-342 | A-323 | 0.24471 | 0.02687 |
| 1941 | UK2014 | A-389 | A-323 | 0.93066 | 0.01121 |
| 1942 | UK2014 | A-547 | A-323 | 0.23222 | 0.03749 |
| 1943 | UK2014 | A-196 | A-323 | 0.01254 | 0.00731 |
| 1944 | UK2014 | A-360 | A-323 | 0.10314 | 0.01844 |
| 1945 | UK2014 | A-465 | A-323 | 0.67272 | 0.04127 |
| 1946 | UK2014 | A-161 | A-323 | 0.48010 | 0.03964 |
| 1947 | UK2014 | A-200 | A-323 | 0.00964 | 0.00513 |
| 1948 | UK2014 | A-325 | A-323 | 0.30415 | 0.03281 |
| 1949 | UK2014 | A-333 | A-323 | 0.70879 | 0.02939 |
| 1950 | UK2014 | A-104.2 | A-323 | 0.55698 | 0.02685 |
| 1951 | UK2014 | A-122 | A-323 | 0.81137 | 0.02986 |
| 1952 | UK2014 | A-175 | A-323 | 0.25181 | 0.02681 |
| 1953 | UK2014 | A-300 | A-323 | 0.40784 | 0.04166 |
| 1954 | UK2014 | A-733 | A-323 | 0.17719 | 0.00901 |
| 1955 | UK2014 | A-0 | A-323 | 0.40580 | 0.03688 |
| 1956 | UK2014 | A-1.1 | A-323 | 0.78523 | 0.02739 |
| 1957 | UK2014 | A-450 | A-323 | 0.13482 | 0.02305 |
| 1958 | UK2014 | A-60 | A-323 | 0.08287 | 0.02200 |
| 1959 | UK2014 | A-662 | A-323 | 0.14387 | 0.01218 |
| 1960 | UK2014 | A-104.1 | A-323 | 0.52978 | 0.03233 |
| 1961 | UK2014 | A-149 | A-323 | 0.23358 | 0.03022 |
| 1962 | UK2014 | A-224 | A-323 | 0.56144 | 0.04089 |
| 1963 | UK2014 | A-1.2 | A-880 | 0.44528 | 0.03284 |
| 1964 | UK2014 | A-293 | A-880 | 0.09703 | 0.01775 |
| 1965 | UK2014 | A-322 | A-880 | 0.87297 | 0.00873 |

| | | | | | |
|---|---|---|---|---|---|
| 1966 | UK2014 | A-421 | A-880 | 0.23868 | 0.02942 |
| 1967 | UK2014 | A-498 | A-880 | 0.70570 | 0.02906 |
| 1968 | UK2014 | A-54 | A-880 | 0.49764 | 0.02882 |
| 1969 | UK2014 | A-275 | A-880 | 0.50084 | 0.02585 |
| 1970 | UK2014 | A-509 | A-880 | 0.10386 | 0.02209 |
| 1971 | UK2014 | A-600 | A-880 | 0.79784 | 0.01220 |
| 1972 | UK2014 | A-04 | A-880 | 0.93173 | 0.01158 |
| 1973 | UK2014 | A-092 | A-880 | 0.14291 | 0.00981 |
| 1974 | UK2014 | A-244 | A-880 | 0.10654 | 0.01601 |
| 1975 | UK2014 | A-342 | A-880 | 0.17970 | 0.02239 |
| 1976 | UK2014 | A-389 | A-880 | 0.86247 | 0.01401 |
| 1977 | UK2014 | A-547 | A-880 | 0.02615 | 0.00776 |
| 1978 | UK2014 | A-196 | A-880 | 0.84572 | 0.03046 |
| 1979 | UK2014 | A-360 | A-880 | 0.62767 | 0.02856 |
| 1980 | UK2014 | A-465 | A-880 | 0.55393 | 0.03952 |
| 1981 | UK2014 | A-161 | A-880 | 0.31715 | 0.03535 |
| 1982 | UK2014 | A-200 | A-880 | 0.94444 | 0.01573 |
| 1983 | UK2014 | A-325 | A-880 | 0.51828 | 0.03437 |
| 1984 | UK2014 | A-333 | A-880 | 0.03449 | 0.01009 |
| 1985 | UK2014 | A-104.2 | A-880 | 0.98313 | 0.00370 |
| 1986 | UK2014 | A-122 | A-880 | 0.98890 | 0.00669 |
| 1987 | UK2014 | A-175 | A-880 | 0.13940 | 0.01926 |
| 1988 | UK2014 | A-300 | A-880 | 0.97914 | 0.00813 |
| 1989 | UK2014 | A-733 | A-880 | 0.61754 | 0.01164 |
| 1990 | UK2014 | A-0 | A-880 | 0.11990 | 0.02066 |
| 1991 | UK2014 | A-1.1 | A-880 | 0.42118 | 0.03171 |
| 1992 | UK2014 | A-450 | A-880 | 0.97716 | 0.00645 |
| 1993 | UK2014 | A-60 | A-880 | 0.32112 | 0.03287 |
| 1994 | UK2014 | A-662 | A-880 | 0.91878 | 0.00734 |
| 1995 | UK2014 | A-104.1 | A-880 | 0.00000 | 0.00000 |
| 1996 | UK2014 | A-149 | A-880 | 0.40383 | 0.03394 |
| 1997 | UK2014 | A-224 | A-880 | 0.54677 | 0.04018 |
| 1998 | UK2014 | A-323 | A-880 | 0.40080 | 0.03829 |
| 1999 | UK2016 | A-1.2 | A-293 | 0.73337 | 0.01898 |
| 2000 | UK2016 | A-1.2 | A-322 | 0.00064 | 0.00045 |
| 2001 | UK2016 | A-293 | A-322 | 0.09733 | 0.00615 |
| 2002 | UK2016 | A-1.2 | A-421 | 0.78196 | 0.02313 |
| 2003 | UK2016 | A-293 | A-421 | 0.85245 | 0.01536 |
| 2004 | UK2016 | A-322 | A-421 | 0.71265 | 0.01254 |
| 2005 | UK2016 | A-1.2 | A-498 | 0.62923 | 0.02578 |
| 2006 | UK2016 | A-293 | A-498 | 0.70892 | 0.02361 |
| 2007 | UK2016 | A-322 | A-498 | 0.45973 | 0.01216 |
| 2008 | UK2016 | A-421 | A-498 | 0.72202 | 0.02820 |
| 2009 | UK2016 | A-1.2 | A-54 | 0.64021 | 0.02381 |
| 2010 | UK2016 | A-293 | A-54 | 0.90091 | 0.01068 |
| 2011 | UK2016 | A-322 | A-54 | 0.38298 | 0.01173 |
| 2012 | UK2016 | A-421 | A-54 | 0.24652 | 0.02458 |
| 2013 | UK2016 | A-498 | A-54 | 0.26524 | 0.02385 |

| | | | | | |
|---|---|---|---|---|---|
| 2014 | UK2016 | A-1.2 | A-275 | 0.05195 | 0.00763 |
| 2015 | UK2016 | A-293 | A-275 | 0.18005 | 0.01289 |
| 2016 | UK2016 | A-322 | A-275 | 0.00144 | 0.00039 |
| 2017 | UK2016 | A-421 | A-275 | 0.27632 | 0.01970 |
| 2018 | UK2016 | A-498 | A-275 | 0.09974 | 0.01142 |
| 2019 | UK2016 | A-54 | A-275 | 0.36068 | 0.01584 |
| 2020 | UK2016 | A-1.2 | A-509 | 0.06660 | 0.01422 |
| 2021 | UK2016 | A-293 | A-509 | 0.94864 | 0.00809 |
| 2022 | UK2016 | A-322 | A-509 | 0.75845 | 0.01050 |
| 2023 | UK2016 | A-421 | A-509 | 0.33499 | 0.02978 |
| 2024 | UK2016 | A-498 | A-509 | 0.12886 | 0.01975 |
| 2025 | UK2016 | A-54 | A-509 | 0.59377 | 0.02786 |
| 2026 | UK2016 | A-275 | A-509 | 0.86853 | 0.01142 |
| 2027 | UK2016 | A-1.2 | A-600 | 0.07843 | 0.00876 |
| 2028 | UK2016 | A-293 | A-600 | 0.19063 | 0.01166 |
| 2029 | UK2016 | A-322 | A-600 | 0.49554 | 0.00720 |
| 2030 | UK2016 | A-421 | A-600 | 0.96627 | 0.00539 |
| 2031 | UK2016 | A-498 | A-600 | 0.30105 | 0.01615 |
| 2032 | UK2016 | A-54 | A-600 | 0.67149 | 0.01327 |
| 2033 | UK2016 | A-275 | A-600 | 0.49210 | 0.01146 |
| 2034 | UK2016 | A-509 | A-600 | 0.17126 | 0.01484 |
| 2035 | UK2016 | A-1.2 | A-04 | 0.35422 | 0.01941 |
| 2036 | UK2016 | A-293 | A-04 | 0.06456 | 0.00819 |
| 2037 | UK2016 | A-322 | A-04 | 0.35226 | 0.00853 |
| 2038 | UK2016 | A-421 | A-04 | 0.61365 | 0.02203 |
| 2039 | UK2016 | A-498 | A-04 | 0.78549 | 0.01881 |
| 2040 | UK2016 | A-54 | A-04 | 0.31432 | 0.01809 |
| 2041 | UK2016 | A-275 | A-04 | 0.02796 | 0.00396 |
| 2042 | UK2016 | A-509 | A-04 | 0.66996 | 0.02211 |
| 2043 | UK2016 | A-600 | A-04 | 0.44268 | 0.01335 |
| 2044 | UK2016 | A-1.2 | A-092 | 0.81517 | 0.00770 |
| 2045 | UK2016 | A-293 | A-092 | 0.27773 | 0.00855 |
| 2046 | UK2016 | A-322 | A-092 | 0.43745 | 0.00548 |
| 2047 | UK2016 | A-421 | A-092 | 0.82193 | 0.00801 |
| 2048 | UK2016 | A-498 | A-092 | 0.92369 | 0.00442 |
| 2049 | UK2016 | A-54 | A-092 | 0.27846 | 0.01008 |
| 2050 | UK2016 | A-275 | A-092 | 0.74368 | 0.00669 |
| 2051 | UK2016 | A-509 | A-092 | 0.20156 | 0.01054 |
| 2052 | UK2016 | A-600 | A-092 | 0.64275 | 0.00713 |
| 2053 | UK2016 | A-04 | A-092 | 0.98953 | 0.00138 |
| 2054 | UK2016 | A-1.2 | A-244 | 0.08666 | 0.01288 |
| 2055 | UK2016 | A-293 | A-244 | 0.62562 | 0.01767 |
| 2056 | UK2016 | A-322 | A-244 | 0.47914 | 0.00948 |
| 2057 | UK2016 | A-421 | A-244 | 0.70539 | 0.02246 |
| 2058 | UK2016 | A-498 | A-244 | 0.98403 | 0.00426 |
| 2059 | UK2016 | A-54 | A-244 | 0.48163 | 0.02398 |
| 2060 | UK2016 | A-275 | A-244 | 0.52601 | 0.01369 |
| 2061 | UK2016 | A-509 | A-244 | 0.47625 | 0.02556 |

| | | | | | |
|---|---|---|---|---|---|
| 2062 | UK2016 | A-600 | A-244 | 0.81324 | 0.01032 |
| 2063 | UK2016 | A-04 | A-244 | 0.00532 | 0.00167 |
| 2064 | UK2016 | A-092 | A-244 | 0.39233 | 0.00942 |
| 2065 | UK2016 | A-1.2 | A-342 | 0.53321 | 0.01888 |
| 2066 | UK2016 | A-293 | A-342 | 0.19691 | 0.01333 |
| 2067 | UK2016 | A-322 | A-342 | 0.85707 | 0.00518 |
| 2068 | UK2016 | A-421 | A-342 | 0.44394 | 0.02549 |
| 2069 | UK2016 | A-498 | A-342 | 0.45784 | 0.01995 |
| 2070 | UK2016 | A-54 | A-342 | 0.87974 | 0.01104 |
| 2071 | UK2016 | A-275 | A-342 | 0.13422 | 0.00842 |
| 2072 | UK2016 | A-509 | A-342 | 0.90793 | 0.01083 |
| 2073 | UK2016 | A-600 | A-342 | 0.80043 | 0.00872 |
| 2074 | UK2016 | A-04 | A-342 | 0.94557 | 0.00461 |
| 2075 | UK2016 | A-092 | A-342 | 0.92006 | 0.00379 |
| 2076 | UK2016 | A-244 | A-342 | 0.88825 | 0.00890 |
| 2077 | UK2016 | A-1.2 | A-389 | 0.38975 | 0.01374 |
| 2078 | UK2016 | A-293 | A-389 | 0.83068 | 0.00666 |
| 2079 | UK2016 | A-322 | A-389 | 0.50562 | 0.00654 |
| 2080 | UK2016 | A-421 | A-389 | 0.04667 | 0.00588 |
| 2081 | UK2016 | A-498 | A-389 | 0.35723 | 0.01286 |
| 2082 | UK2016 | A-54 | A-389 | 0.64672 | 0.01285 |
| 2083 | UK2016 | A-275 | A-389 | 0.39326 | 0.00984 |
| 2084 | UK2016 | A-509 | A-389 | 0.78909 | 0.00934 |
| 2085 | UK2016 | A-600 | A-389 | 0.44103 | 0.00908 |
| 2086 | UK2016 | A-04 | A-389 | 0.03308 | 0.00376 |
| 2087 | UK2016 | A-092 | A-389 | 0.80901 | 0.00385 |
| 2088 | UK2016 | A-244 | A-389 | 0.73962 | 0.01025 |
| 2089 | UK2016 | A-342 | A-389 | 0.09726 | 0.00660 |
| 2090 | UK2016 | A-1.2 | A-547 | 0.34707 | 0.02913 |
| 2091 | UK2016 | A-293 | A-547 | 0.05929 | 0.00997 |
| 2092 | UK2016 | A-322 | A-547 | 0.35472 | 0.01182 |
| 2093 | UK2016 | A-421 | A-547 | 0.14773 | 0.02067 |
| 2094 | UK2016 | A-498 | A-547 | 0.94495 | 0.01296 |
| 2095 | UK2016 | A-54 | A-547 | 0.48004 | 0.02536 |
| 2096 | UK2016 | A-275 | A-547 | 0.62461 | 0.01683 |
| 2097 | UK2016 | A-509 | A-547 | 0.92244 | 0.01555 |
| 2098 | UK2016 | A-600 | A-547 | 0.37415 | 0.01889 |
| 2099 | UK2016 | A-04 | A-547 | 0.81562 | 0.01440 |
| 2100 | UK2016 | A-092 | A-547 | 0.35328 | 0.01045 |
| 2101 | UK2016 | A-244 | A-547 | 0.73217 | 0.01714 |
| 2102 | UK2016 | A-342 | A-547 | 0.57273 | 0.02000 |
| 2103 | UK2016 | A-389 | A-547 | 0.74458 | 0.01201 |
| 2104 | UK2016 | A-1.2 | A-196 | 0.54193 | 0.02916 |
| 2105 | UK2016 | A-293 | A-196 | 0.84254 | 0.02006 |
| 2106 | UK2016 | A-322 | A-196 | 0.51056 | 0.01476 |
| 2107 | UK2016 | A-421 | A-196 | 0.67379 | 0.02797 |
| 2108 | UK2016 | A-498 | A-196 | 0.50255 | 0.02799 |
| 2109 | UK2016 | A-54 | A-196 | 0.19181 | 0.02097 |

| | | | | | |
|---|---|---|---|---|---|
| 2110 | UK2016 | A-275 | A-196 | 0.57327 | 0.02060 |
| 2111 | UK2016 | A-509 | A-196 | 0.52150 | 0.03054 |
| 2112 | UK2016 | A-600 | A-196 | 0.90819 | 0.00893 |
| 2113 | UK2016 | A-04 | A-196 | 0.58887 | 0.02170 |
| 2114 | UK2016 | A-092 | A-196 | 0.47271 | 0.01229 |
| 2115 | UK2016 | A-244 | A-196 | 0.84721 | 0.01445 |
| 2116 | UK2016 | A-342 | A-196 | 0.12993 | 0.01433 |
| 2117 | UK2016 | A-389 | A-196 | 0.97249 | 0.00402 |
| 2118 | UK2016 | A-547 | A-196 | 0.73715 | 0.02700 |
| 2119 | UK2016 | A-1.2 | A-360 | 0.25498 | 0.01514 |
| 2120 | UK2016 | A-293 | A-360 | 0.38076 | 0.01614 |
| 2121 | UK2016 | A-322 | A-360 | 0.66411 | 0.00596 |
| 2122 | UK2016 | A-421 | A-360 | 0.58076 | 0.01955 |
| 2123 | UK2016 | A-498 | A-360 | 0.57160 | 0.01765 |
| 2124 | UK2016 | A-54 | A-360 | 0.77267 | 0.01286 |
| 2125 | UK2016 | A-275 | A-360 | 0.57043 | 0.01043 |
| 2126 | UK2016 | A-509 | A-360 | 0.57080 | 0.01997 |
| 2127 | UK2016 | A-600 | A-360 | 0.71123 | 0.00914 |
| 2128 | UK2016 | A-04 | A-360 | 0.94673 | 0.00428 |
| 2129 | UK2016 | A-092 | A-360 | 0.05557 | 0.00297 |
| 2130 | UK2016 | A-244 | A-360 | 0.54301 | 0.01455 |
| 2131 | UK2016 | A-342 | A-360 | 0.62450 | 0.01153 |
| 2132 | UK2016 | A-389 | A-360 | 0.69901 | 0.00734 |
| 2133 | UK2016 | A-547 | A-360 | 0.95770 | 0.00530 |
| 2134 | UK2016 | A-196 | A-360 | 0.18917 | 0.01345 |
| 2135 | UK2016 | A-1.2 | A-465 | 0.14677 | 0.01835 |
| 2136 | UK2016 | A-293 | A-465 | 0.27077 | 0.01942 |
| 2137 | UK2016 | A-322 | A-465 | 0.96271 | 0.00320 |
| 2138 | UK2016 | A-421 | A-465 | 0.22750 | 0.02806 |
| 2139 | UK2016 | A-498 | A-465 | 0.48893 | 0.03268 |
| 2140 | UK2016 | A-54 | A-465 | 0.64716 | 0.02268 |
| 2141 | UK2016 | A-275 | A-465 | 0.68120 | 0.01642 |
| 2142 | UK2016 | A-509 | A-465 | 0.25423 | 0.02728 |
| 2143 | UK2016 | A-600 | A-465 | 0.58653 | 0.01549 |
| 2144 | UK2016 | A-04 | A-465 | 0.87654 | 0.01210 |
| 2145 | UK2016 | A-092 | A-465 | 0.90876 | 0.00633 |
| 2146 | UK2016 | A-244 | A-465 | 0.80615 | 0.01518 |
| 2147 | UK2016 | A-342 | A-465 | 0.57539 | 0.02025 |
| 2148 | UK2016 | A-389 | A-465 | 0.79652 | 0.00971 |
| 2149 | UK2016 | A-547 | A-465 | 0.06216 | 0.01517 |
| 2150 | UK2016 | A-196 | A-465 | 0.72143 | 0.02602 |
| 2151 | UK2016 | A-360 | A-465 | 0.36137 | 0.01792 |
| 2152 | UK2016 | A-1.2 | A-161 | 0.07541 | 0.01522 |
| 2153 | UK2016 | A-293 | A-161 | 0.20359 | 0.02119 |
| 2154 | UK2016 | A-322 | A-161 | 0.10758 | 0.00872 |
| 2155 | UK2016 | A-421 | A-161 | 0.55441 | 0.03402 |
| 2156 | UK2016 | A-498 | A-161 | 0.65197 | 0.03024 |
| 2157 | UK2016 | A-54 | A-161 | 0.02551 | 0.00780 |

| | | | | | |
|---|---|---|---|---|---|
| 2158 | UK2016 | A-275 | A-161 | 0.69180 | 0.01645 |
| 2159 | UK2016 | A-509 | A-161 | 0.17981 | 0.02358 |
| 2160 | UK2016 | A-600 | A-161 | 0.98146 | 0.00376 |
| 2161 | UK2016 | A-04 | A-161 | 0.22437 | 0.01974 |
| 2162 | UK2016 | A-092 | A-161 | 0.76363 | 0.01000 |
| 2163 | UK2016 | A-244 | A-161 | 0.46738 | 0.02727 |
| 2164 | UK2016 | A-342 | A-161 | 0.25425 | 0.01810 |
| 2165 | UK2016 | A-389 | A-161 | 0.16001 | 0.01096 |
| 2166 | UK2016 | A-547 | A-161 | 0.69993 | 0.03223 |
| 2167 | UK2016 | A-196 | A-161 | 0.31422 | 0.02793 |
| 2168 | UK2016 | A-360 | A-161 | 0.23744 | 0.01807 |
| 2169 | UK2016 | A-465 | A-161 | 0.82498 | 0.02251 |
| 2170 | UK2016 | A-1.2 | A-200 | 0.00000 | 0.00000 |
| 2171 | UK2016 | A-293 | A-200 | 0.31454 | 0.02592 |
| 2172 | UK2016 | A-322 | A-200 | 0.00012 | 0.00009 |
| 2173 | UK2016 | A-421 | A-200 | 0.36425 | 0.03296 |
| 2174 | UK2016 | A-498 | A-200 | 0.62390 | 0.03266 |
| 2175 | UK2016 | A-54 | A-200 | 0.99481 | 0.00264 |
| 2176 | UK2016 | A-275 | A-200 | 0.00000 | 0.00000 |
| 2177 | UK2016 | A-509 | A-200 | 0.38201 | 0.03385 |
| 2178 | UK2016 | A-600 | A-200 | 0.67323 | 0.01874 |
| 2179 | UK2016 | A-04 | A-200 | 0.22721 | 0.02030 |
| 2180 | UK2016 | A-092 | A-200 | 0.55146 | 0.01433 |
| 2181 | UK2016 | A-244 | A-200 | 0.20579 | 0.02074 |
| 2182 | UK2016 | A-342 | A-200 | 0.47644 | 0.02393 |
| 2183 | UK2016 | A-389 | A-200 | 0.10537 | 0.00926 |
| 2184 | UK2016 | A-547 | A-200 | 0.68601 | 0.03103 |
| 2185 | UK2016 | A-196 | A-200 | 0.76537 | 0.02653 |
| 2186 | UK2016 | A-360 | A-200 | 0.07916 | 0.01154 |
| 2187 | UK2016 | A-465 | A-200 | 0.99115 | 0.00457 |
| 2188 | UK2016 | A-161 | A-200 | 0.61041 | 0.03236 |
| 2189 | UK2016 | A-1.2 | A-325 | 0.65962 | 0.02301 |
| 2190 | UK2016 | A-293 | A-325 | 0.30181 | 0.02115 |
| 2191 | UK2016 | A-322 | A-325 | 0.43496 | 0.01128 |
| 2192 | UK2016 | A-421 | A-325 | 0.60379 | 0.02727 |
| 2193 | UK2016 | A-498 | A-325 | 0.93930 | 0.01170 |
| 2194 | UK2016 | A-54 | A-325 | 0.11213 | 0.01271 |
| 2195 | UK2016 | A-275 | A-325 | 0.52285 | 0.01778 |
| 2196 | UK2016 | A-509 | A-325 | 0.55531 | 0.02618 |
| 2197 | UK2016 | A-600 | A-325 | 0.51601 | 0.01495 |
| 2198 | UK2016 | A-04 | A-325 | 0.31586 | 0.01763 |
| 2199 | UK2016 | A-092 | A-325 | 0.86339 | 0.00737 |
| 2200 | UK2016 | A-244 | A-325 | 0.50289 | 0.02211 |
| 2201 | UK2016 | A-342 | A-325 | 0.63324 | 0.01765 |
| 2202 | UK2016 | A-389 | A-325 | 0.33919 | 0.01239 |
| 2203 | UK2016 | A-547 | A-325 | 0.83877 | 0.01921 |
| 2204 | UK2016 | A-196 | A-325 | 0.86658 | 0.01777 |
| 2205 | UK2016 | A-360 | A-325 | 0.37380 | 0.01531 |

| | | | | | |
|---|---|---|---|---|---|
| 2206 | UK2016 | A-465 | A-325 | 0.43181 | 0.02794 |
| 2207 | UK2016 | A-161 | A-325 | 0.33803 | 0.02814 |
| 2208 | UK2016 | A-200 | A-325 | 0.42590 | 0.02704 |
| 2209 | UK2016 | A-1.2 | A-333 | 0.81140 | 0.02016 |
| 2210 | UK2016 | A-293 | A-333 | 0.23231 | 0.02151 |
| 2211 | UK2016 | A-322 | A-333 | 0.32638 | 0.01312 |
| 2212 | UK2016 | A-421 | A-333 | 0.28084 | 0.03087 |
| 2213 | UK2016 | A-498 | A-333 | 0.94203 | 0.01092 |
| 2214 | UK2016 | A-54 | A-333 | 0.15235 | 0.01783 |
| 2215 | UK2016 | A-275 | A-333 | 0.71971 | 0.01663 |
| 2216 | UK2016 | A-509 | A-333 | 0.94308 | 0.01224 |
| 2217 | UK2016 | A-600 | A-333 | 0.09060 | 0.01002 |
| 2218 | UK2016 | A-04 | A-333 | 0.16246 | 0.01608 |
| 2219 | UK2016 | A-092 | A-333 | 0.90690 | 0.00541 |
| 2220 | UK2016 | A-244 | A-333 | 0.12904 | 0.01505 |
| 2221 | UK2016 | A-342 | A-333 | 0.16987 | 0.01571 |
| 2222 | UK2016 | A-389 | A-333 | 0.24126 | 0.01370 |
| 2223 | UK2016 | A-547 | A-333 | 0.99613 | 0.00185 |
| 2224 | UK2016 | A-196 | A-333 | 0.87123 | 0.01769 |
| 2225 | UK2016 | A-360 | A-333 | 0.18561 | 0.01379 |
| 2226 | UK2016 | A-465 | A-333 | 0.23161 | 0.02510 |
| 2227 | UK2016 | A-161 | A-333 | 0.62737 | 0.03118 |
| 2228 | UK2016 | A-200 | A-333 | 0.76978 | 0.02613 |
| 2229 | UK2016 | A-325 | A-333 | 0.69100 | 0.02543 |
| 2230 | UK2016 | A-1.2 | A-104.2 | 0.65919 | 0.02243 |
| 2231 | UK2016 | A-293 | A-104.2 | 0.92595 | 0.00812 |
| 2232 | UK2016 | A-322 | A-104.2 | 0.89782 | 0.00518 |
| 2233 | UK2016 | A-421 | A-104.2 | 0.79448 | 0.02029 |
| 2234 | UK2016 | A-498 | A-104.2 | 0.49969 | 0.02638 |
| 2235 | UK2016 | A-54 | A-104.2 | 0.31188 | 0.01497 |
| 2236 | UK2016 | A-275 | A-104.2 | 0.60413 | 0.01513 |
| 2237 | UK2016 | A-509 | A-104.2 | 0.04756 | 0.00862 |
| 2238 | UK2016 | A-600 | A-104.2 | 0.25985 | 0.01274 |
| 2239 | UK2016 | A-04 | A-104.2 | 0.31264 | 0.01756 |
| 2240 | UK2016 | A-092 | A-104.2 | 0.30878 | 0.01148 |
| 2241 | UK2016 | A-244 | A-104.2 | 0.36692 | 0.01836 |
| 2242 | UK2016 | A-342 | A-104.2 | 0.14171 | 0.01205 |
| 2243 | UK2016 | A-389 | A-104.2 | 0.22489 | 0.00907 |
| 2244 | UK2016 | A-547 | A-104.2 | 0.93925 | 0.00915 |
| 2245 | UK2016 | A-196 | A-104.2 | 0.28501 | 0.02443 |
| 2246 | UK2016 | A-360 | A-104.2 | 0.95171 | 0.00334 |
| 2247 | UK2016 | A-465 | A-104.2 | 0.46367 | 0.02471 |
| 2248 | UK2016 | A-161 | A-104.2 | 0.45854 | 0.02373 |
| 2249 | UK2016 | A-200 | A-104.2 | 0.72435 | 0.02252 |
| 2250 | UK2016 | A-325 | A-104.2 | 0.60069 | 0.02307 |
| 2251 | UK2016 | A-333 | A-104.2 | 0.49688 | 0.02554 |
| 2252 | UK2016 | A-1.2 | A-122 | 0.00240 | 0.00128 |
| 2253 | UK2016 | A-293 | A-122 | 0.85667 | 0.01370 |

| | | | | | |
|---|---|---|---|---|---|
| 2254 | UK2016 | A-322 | A-122 | 0.00000 | 0.00000 |
| 2255 | UK2016 | A-421 | A-122 | 0.05679 | 0.01297 |
| 2256 | UK2016 | A-498 | A-122 | 0.14660 | 0.01981 |
| 2257 | UK2016 | A-54 | A-122 | 0.31210 | 0.02468 |
| 2258 | UK2016 | A-275 | A-122 | 0.00000 | 0.00000 |
| 2259 | UK2016 | A-509 | A-122 | 0.34553 | 0.02936 |
| 2260 | UK2016 | A-600 | A-122 | 0.23380 | 0.01518 |
| 2261 | UK2016 | A-04 | A-122 | 0.01765 | 0.00373 |
| 2262 | UK2016 | A-092 | A-122 | 0.89436 | 0.00678 |
| 2263 | UK2016 | A-244 | A-122 | 0.71754 | 0.02091 |
| 2264 | UK2016 | A-342 | A-122 | 0.64317 | 0.01753 |
| 2265 | UK2016 | A-389 | A-122 | 0.79647 | 0.01076 |
| 2266 | UK2016 | A-547 | A-122 | 0.40573 | 0.02912 |
| 2267 | UK2016 | A-196 | A-122 | 0.81875 | 0.01952 |
| 2268 | UK2016 | A-360 | A-122 | 0.95098 | 0.00599 |
| 2269 | UK2016 | A-465 | A-122 | 0.75132 | 0.02260 |
| 2270 | UK2016 | A-161 | A-122 | 0.33889 | 0.03063 |
| 2271 | UK2016 | A-200 | A-122 | 0.00000 | 0.00000 |
| 2272 | UK2016 | A-325 | A-122 | 0.74053 | 0.02021 |
| 2273 | UK2016 | A-333 | A-122 | 0.57587 | 0.02840 |
| 2274 | UK2016 | A-104.2 | A-122 | 0.64621 | 0.02512 |
| 2275 | UK2016 | A-1.2 | A-175 | 0.02922 | 0.00804 |
| 2276 | UK2016 | A-293 | A-175 | 0.07754 | 0.01585 |
| 2277 | UK2016 | A-322 | A-175 | 0.70037 | 0.01198 |
| 2278 | UK2016 | A-421 | A-175 | 0.19053 | 0.02315 |
| 2279 | UK2016 | A-498 | A-175 | 0.68247 | 0.03007 |
| 2280 | UK2016 | A-54 | A-175 | 0.18309 | 0.02197 |
| 2281 | UK2016 | A-275 | A-175 | 0.26045 | 0.01845 |
| 2282 | UK2016 | A-509 | A-175 | 0.38898 | 0.03279 |
| 2283 | UK2016 | A-600 | A-175 | 0.22560 | 0.01970 |
| 2284 | UK2016 | A-04 | A-175 | 0.76810 | 0.02174 |
| 2285 | UK2016 | A-092 | A-175 | 0.13217 | 0.00900 |
| 2286 | UK2016 | A-244 | A-175 | 0.85569 | 0.01932 |
| 2287 | UK2016 | A-342 | A-175 | 0.31123 | 0.02126 |
| 2288 | UK2016 | A-389 | A-175 | 0.22871 | 0.01526 |
| 2289 | UK2016 | A-547 | A-175 | 0.60436 | 0.03440 |
| 2290 | UK2016 | A-196 | A-175 | 0.12065 | 0.01963 |
| 2291 | UK2016 | A-360 | A-175 | 0.36414 | 0.02185 |
| 2292 | UK2016 | A-465 | A-175 | 0.01806 | 0.00601 |
| 2293 | UK2016 | A-161 | A-175 | 0.98850 | 0.00501 |
| 2294 | UK2016 | A-200 | A-175 | 0.51075 | 0.03489 |
| 2295 | UK2016 | A-325 | A-175 | 0.89595 | 0.01484 |
| 2296 | UK2016 | A-333 | A-175 | 0.94997 | 0.01200 |
| 2297 | UK2016 | A-104.2 | A-175 | 0.32592 | 0.02451 |
| 2298 | UK2016 | A-122 | A-175 | 0.49325 | 0.03154 |
| 2299 | UK2016 | A-1.2 | A-300 | 0.70788 | 0.03638 |
| 2300 | UK2016 | A-293 | A-300 | 0.17932 | 0.02983 |
| 2301 | UK2016 | A-322 | A-300 | 0.28997 | 0.02097 |

| | | | | | |
|---|---|---|---|---|---|
| 2302 | UK2016 | A-421 | A-300 | 0.27647 | 0.04002 |
| 2303 | UK2016 | A-498 | A-300 | 0.29358 | 0.03883 |
| 2304 | UK2016 | A-54 | A-300 | 0.70319 | 0.03366 |
| 2305 | UK2016 | A-275 | A-300 | 0.21578 | 0.02441 |
| 2306 | UK2016 | A-509 | A-300 | 0.17216 | 0.03123 |
| 2307 | UK2016 | A-600 | A-300 | 0.14040 | 0.01871 |
| 2308 | UK2016 | A-04 | A-300 | 0.73163 | 0.02757 |
| 2309 | UK2016 | A-092 | A-300 | 0.29594 | 0.01616 |
| 2310 | UK2016 | A-244 | A-300 | 0.92955 | 0.01630 |
| 2311 | UK2016 | A-342 | A-300 | 0.49239 | 0.03163 |
| 2312 | UK2016 | A-389 | A-300 | 0.91066 | 0.01156 |
| 2313 | UK2016 | A-547 | A-300 | 0.44099 | 0.04173 |
| 2314 | UK2016 | A-196 | A-300 | 0.08652 | 0.02147 |
| 2315 | UK2016 | A-360 | A-300 | 0.46345 | 0.03023 |
| 2316 | UK2016 | A-465 | A-300 | 0.77252 | 0.03465 |
| 2317 | UK2016 | A-161 | A-300 | 0.98305 | 0.00687 |
| 2318 | UK2016 | A-200 | A-300 | 0.27682 | 0.03513 |
| 2319 | UK2016 | A-325 | A-300 | 0.98006 | 0.00967 |
| 2320 | UK2016 | A-333 | A-300 | 0.78815 | 0.03343 |
| 2321 | UK2016 | A-104.2 | A-300 | 0.24245 | 0.02977 |
| 2322 | UK2016 | A-122 | A-300 | 0.12939 | 0.02530 |
| 2323 | UK2016 | A-175 | A-300 | 0.11029 | 0.02682 |
| 2324 | UK2016 | A-1.2 | A-733 | 0.10516 | 0.01100 |
| 2325 | UK2016 | A-293 | A-733 | 1.00000 | 0.00000 |
| 2326 | UK2016 | A-322 | A-733 | 0.58357 | 0.01029 |
| 2327 | UK2016 | A-421 | A-733 | 0.68447 | 0.02024 |
| 2328 | UK2016 | A-498 | A-733 | 0.65727 | 0.02000 |
| 2329 | UK2016 | A-54 | A-733 | 0.76198 | 0.01690 |
| 2330 | UK2016 | A-275 | A-733 | 0.54030 | 0.01215 |
| 2331 | UK2016 | A-509 | A-733 | 0.68746 | 0.01962 |
| 2332 | UK2016 | A-600 | A-733 | 0.47268 | 0.01310 |
| 2333 | UK2016 | A-04 | A-733 | 0.65535 | 0.01843 |
| 2334 | UK2016 | A-092 | A-733 | 1.00000 | 0.00000 |
| 2335 | UK2016 | A-244 | A-733 | 0.46213 | 0.01747 |
| 2336 | UK2016 | A-342 | A-733 | 0.04688 | 0.00595 |
| 2337 | UK2016 | A-389 | A-733 | 0.84243 | 0.00571 |
| 2338 | UK2016 | A-547 | A-733 | 0.30881 | 0.01830 |
| 2339 | UK2016 | A-196 | A-733 | 0.89600 | 0.01295 |
| 2340 | UK2016 | A-360 | A-733 | 0.25104 | 0.01296 |
| 2341 | UK2016 | A-465 | A-733 | 0.43418 | 0.02094 |
| 2342 | UK2016 | A-161 | A-733 | 0.66778 | 0.02360 |
| 2343 | UK2016 | A-200 | A-733 | 0.59677 | 0.02271 |
| 2344 | UK2016 | A-325 | A-733 | 0.34693 | 0.02025 |
| 2345 | UK2016 | A-333 | A-733 | 0.55416 | 0.02236 |
| 2346 | UK2016 | A-104.2 | A-733 | 0.26856 | 0.01435 |
| 2347 | UK2016 | A-122 | A-733 | 0.70308 | 0.02018 |
| 2348 | UK2016 | A-175 | A-733 | 0.87215 | 0.01874 |
| 2349 | UK2016 | A-300 | A-733 | 0.31538 | 0.03104 |

| | | | | | |
|---|---|---|---|---|---|
| 2350 | UK2016 | A-1.2 | A-0 | 0.02737 | 0.00463 |
| 2351 | UK2016 | A-293 | A-0 | 0.37896 | 0.01368 |
| 2352 | UK2016 | A-322 | A-0 | 0.09419 | 0.00522 |
| 2353 | UK2016 | A-421 | A-0 | 0.64507 | 0.01904 |
| 2354 | UK2016 | A-498 | A-0 | 0.91176 | 0.00853 |
| 2355 | UK2016 | A-54 | A-0 | 0.62645 | 0.01580 |
| 2356 | UK2016 | A-275 | A-0 | 0.00000 | 0.00000 |
| 2357 | UK2016 | A-509 | A-0 | 0.81261 | 0.01330 |
| 2358 | UK2016 | A-600 | A-0 | 0.92079 | 0.00503 |
| 2359 | UK2016 | A-04 | A-0 | 0.42328 | 0.01581 |
| 2360 | UK2016 | A-092 | A-0 | 0.82735 | 0.00428 |
| 2361 | UK2016 | A-244 | A-0 | 0.93351 | 0.00674 |
| 2362 | UK2016 | A-342 | A-0 | 0.73775 | 0.01088 |
| 2363 | UK2016 | A-389 | A-0 | 0.49228 | 0.01007 |
| 2364 | UK2016 | A-547 | A-0 | 0.54642 | 0.02191 |
| 2365 | UK2016 | A-196 | A-0 | 0.77858 | 0.01444 |
| 2366 | UK2016 | A-360 | A-0 | 0.03351 | 0.00347 |
| 2367 | UK2016 | A-465 | A-0 | 0.84129 | 0.01341 |
| 2368 | UK2016 | A-161 | A-0 | 0.74132 | 0.01737 |
| 2369 | UK2016 | A-200 | A-0 | 0.00000 | 0.00000 |
| 2370 | UK2016 | A-325 | A-0 | 0.28368 | 0.01641 |
| 2371 | UK2016 | A-333 | A-0 | 0.49973 | 0.01833 |
| 2372 | UK2016 | A-104.2 | A-0 | 0.69361 | 0.01289 |
| 2373 | UK2016 | A-122 | A-0 | 0.00000 | 0.00000 |
| 2374 | UK2016 | A-175 | A-0 | 0.41039 | 0.02257 |
| 2375 | UK2016 | A-300 | A-0 | 0.06323 | 0.01415 |
| 2376 | UK2016 | A-733 | A-0 | 0.04168 | 0.00415 |
| 2377 | UK2016 | A-1.2 | A-1.1 | 0.15876 | 0.01601 |
| 2378 | UK2016 | A-293 | A-1.1 | 0.46912 | 0.02367 |
| 2379 | UK2016 | A-322 | A-1.1 | 0.68316 | 0.01135 |
| 2380 | UK2016 | A-421 | A-1.1 | 0.77295 | 0.02473 |
| 2381 | UK2016 | A-498 | A-1.1 | 0.05001 | 0.00921 |
| 2382 | UK2016 | A-54 | A-1.1 | 0.63820 | 0.02193 |
| 2383 | UK2016 | A-275 | A-1.1 | 0.00000 | 0.00000 |
| 2384 | UK2016 | A-509 | A-1.1 | 0.47037 | 0.02845 |
| 2385 | UK2016 | A-600 | A-1.1 | 0.03357 | 0.00551 |
| 2386 | UK2016 | A-04 | A-1.1 | 0.30673 | 0.02054 |
| 2387 | UK2016 | A-092 | A-1.1 | 0.61914 | 0.01119 |
| 2388 | UK2016 | A-244 | A-1.1 | 0.09001 | 0.01012 |
| 2389 | UK2016 | A-342 | A-1.1 | 0.01117 | 0.00298 |
| 2390 | UK2016 | A-389 | A-1.1 | 0.48611 | 0.01281 |
| 2391 | UK2016 | A-547 | A-1.1 | 0.22805 | 0.02169 |
| 2392 | UK2016 | A-196 | A-1.1 | 0.73060 | 0.02351 |
| 2393 | UK2016 | A-360 | A-1.1 | 0.08135 | 0.00780 |
| 2394 | UK2016 | A-465 | A-1.1 | 0.93475 | 0.01098 |
| 2395 | UK2016 | A-161 | A-1.1 | 0.33364 | 0.02616 |
| 2396 | UK2016 | A-200 | A-1.1 | 0.00000 | 0.00000 |
| 2397 | UK2016 | A-325 | A-1.1 | 0.34665 | 0.02577 |

| | | | | | |
|---|---|---|---|---|---|
| 2398 | UK2016 | A-333 | A-1.1 | 0.97564 | 0.00467 |
| 2399 | UK2016 | A-104.2 | A-1.1 | 0.98901 | 0.00295 |
| 2400 | UK2016 | A-122 | A-1.1 | 0.33876 | 0.02309 |
| 2401 | UK2016 | A-175 | A-1.1 | 0.52540 | 0.03201 |
| 2402 | UK2016 | A-300 | A-1.1 | 0.64089 | 0.03274 |
| 2403 | UK2016 | A-733 | A-1.1 | 0.41885 | 0.02025 |
| 2404 | UK2016 | A-0 | A-1.1 | 0.12824 | 0.01165 |
| 2405 | UK2016 | A-1.2 | A-450 | 0.16622 | 0.01881 |
| 2406 | UK2016 | A-293 | A-450 | 0.23711 | 0.01848 |
| 2407 | UK2016 | A-322 | A-450 | 0.08905 | 0.00703 |
| 2408 | UK2016 | A-421 | A-450 | 0.75801 | 0.02129 |
| 2409 | UK2016 | A-498 | A-450 | 0.16602 | 0.02029 |
| 2410 | UK2016 | A-54 | A-450 | 0.90726 | 0.01253 |
| 2411 | UK2016 | A-275 | A-450 | 0.39797 | 0.01691 |
| 2412 | UK2016 | A-509 | A-450 | 0.31904 | 0.02636 |
| 2413 | UK2016 | A-600 | A-450 | 0.48218 | 0.01607 |
| 2414 | UK2016 | A-04 | A-450 | 0.21564 | 0.01638 |
| 2415 | UK2016 | A-092 | A-450 | 0.19543 | 0.00783 |
| 2416 | UK2016 | A-244 | A-450 | 0.32353 | 0.02091 |
| 2417 | UK2016 | A-342 | A-450 | 0.41098 | 0.01940 |
| 2418 | UK2016 | A-389 | A-450 | 0.30709 | 0.01321 |
| 2419 | UK2016 | A-547 | A-450 | 0.40282 | 0.02676 |
| 2420 | UK2016 | A-196 | A-450 | 0.65501 | 0.02499 |
| 2421 | UK2016 | A-360 | A-450 | 0.02239 | 0.00342 |
| 2422 | UK2016 | A-465 | A-450 | 0.58044 | 0.02688 |
| 2423 | UK2016 | A-161 | A-450 | 0.28911 | 0.02562 |
| 2424 | UK2016 | A-200 | A-450 | 0.02482 | 0.00614 |
| 2425 | UK2016 | A-325 | A-450 | 0.76504 | 0.01825 |
| 2426 | UK2016 | A-333 | A-450 | 0.24181 | 0.02254 |
| 2427 | UK2016 | A-104.2 | A-450 | 0.04558 | 0.00797 |
| 2428 | UK2016 | A-122 | A-450 | 0.19022 | 0.02006 |
| 2429 | UK2016 | A-175 | A-450 | 0.45383 | 0.02947 |
| 2430 | UK2016 | A-300 | A-450 | 0.03537 | 0.01326 |
| 2431 | UK2016 | A-733 | A-450 | 0.20493 | 0.01473 |
| 2432 | UK2016 | A-0 | A-450 | 0.05196 | 0.00808 |
| 2433 | UK2016 | A-1.1 | A-450 | 0.56134 | 0.02518 |
| 2434 | UK2016 | A-1.2 | A-60 | 0.04099 | 0.01091 |
| 2435 | UK2016 | A-293 | A-60 | 0.79988 | 0.02145 |
| 2436 | UK2016 | A-322 | A-60 | 0.71710 | 0.01219 |
| 2437 | UK2016 | A-421 | A-60 | 0.61248 | 0.03001 |
| 2438 | UK2016 | A-498 | A-60 | 0.84354 | 0.02178 |
| 2439 | UK2016 | A-54 | A-60 | 0.59487 | 0.02920 |
| 2440 | UK2016 | A-275 | A-60 | 0.00413 | 0.00189 |
| 2441 | UK2016 | A-509 | A-60 | 0.17366 | 0.02443 |
| 2442 | UK2016 | A-600 | A-60 | 0.58387 | 0.02122 |
| 2443 | UK2016 | A-04 | A-60 | 0.25127 | 0.01991 |
| 2444 | UK2016 | A-092 | A-60 | 0.50589 | 0.01207 |
| 2445 | UK2016 | A-244 | A-60 | 0.99173 | 0.00390 |

| | | | | | |
|---|---|---|---|---|---|
| 2446 | UK2016 | A-342 | A-60 | 0.32994 | 0.02382 |
| 2447 | UK2016 | A-389 | A-60 | 0.14035 | 0.01180 |
| 2448 | UK2016 | A-547 | A-60 | 0.78452 | 0.02613 |
| 2449 | UK2016 | A-196 | A-60 | 0.00304 | 0.00242 |
| 2450 | UK2016 | A-360 | A-60 | 0.15909 | 0.01383 |
| 2451 | UK2016 | A-465 | A-60 | 0.99945 | 0.00055 |
| 2452 | UK2016 | A-161 | A-60 | 0.88053 | 0.01976 |
| 2453 | UK2016 | A-200 | A-60 | 0.00162 | 0.00118 |
| 2454 | UK2016 | A-325 | A-60 | 0.70227 | 0.02809 |
| 2455 | UK2016 | A-333 | A-60 | 0.56744 | 0.03289 |
| 2456 | UK2016 | A-104.2 | A-60 | 0.29633 | 0.02325 |
| 2457 | UK2016 | A-122 | A-60 | 0.01109 | 0.00755 |
| 2458 | UK2016 | A-175 | A-60 | 0.21145 | 0.02686 |
| 2459 | UK2016 | A-300 | A-60 | 0.24144 | 0.03709 |
| 2460 | UK2016 | A-733 | A-60 | 0.97841 | 0.00477 |
| 2461 | UK2016 | A-0 | A-60 | 0.03509 | 0.00694 |
| 2462 | UK2016 | A-1.1 | A-60 | 0.03141 | 0.00945 |
| 2463 | UK2016 | A-450 | A-60 | 0.36150 | 0.02849 |
| 2464 | UK2016 | A-1.2 | A-662 | 0.00006 | 0.00006 |
| 2465 | UK2016 | A-293 | A-662 | 0.10376 | 0.00707 |
| 2466 | UK2016 | A-322 | A-662 | 0.00000 | 0.00000 |
| 2467 | UK2016 | A-421 | A-662 | 0.69330 | 0.01230 |
| 2468 | UK2016 | A-498 | A-662 | 0.48010 | 0.01190 |
| 2469 | UK2016 | A-54 | A-662 | 0.34377 | 0.01012 |
| 2470 | UK2016 | A-275 | A-662 | 0.00106 | 0.00039 |
| 2471 | UK2016 | A-509 | A-662 | 0.75572 | 0.00954 |
| 2472 | UK2016 | A-600 | A-662 | 0.49247 | 0.00776 |
| 2473 | UK2016 | A-04 | A-662 | 0.36419 | 0.00859 |
| 2474 | UK2016 | A-092 | A-662 | 0.43094 | 0.00523 |
| 2475 | UK2016 | A-244 | A-662 | 0.50520 | 0.01138 |
| 2476 | UK2016 | A-342 | A-662 | 0.86042 | 0.00521 |
| 2477 | UK2016 | A-389 | A-662 | 0.50519 | 0.00606 |
| 2478 | UK2016 | A-547 | A-662 | 0.34810 | 0.01225 |
| 2479 | UK2016 | A-196 | A-662 | 0.51555 | 0.01296 |
| 2480 | UK2016 | A-360 | A-662 | 0.65276 | 0.00558 |
| 2481 | UK2016 | A-465 | A-662 | 0.95904 | 0.00382 |
| 2482 | UK2016 | A-161 | A-662 | 0.10781 | 0.00811 |
| 2483 | UK2016 | A-200 | A-662 | 0.00001 | 0.00001 |
| 2484 | UK2016 | A-325 | A-662 | 0.43486 | 0.01076 |
| 2485 | UK2016 | A-333 | A-662 | 0.32899 | 0.01318 |
| 2486 | UK2016 | A-104.2 | A-662 | 0.90854 | 0.00460 |
| 2487 | UK2016 | A-122 | A-662 | 0.00000 | 0.00000 |
| 2488 | UK2016 | A-175 | A-662 | 0.69559 | 0.01263 |
| 2489 | UK2016 | A-300 | A-662 | 0.28790 | 0.01976 |
| 2490 | UK2016 | A-733 | A-662 | 0.55115 | 0.01028 |
| 2491 | UK2016 | A-0 | A-662 | 0.09847 | 0.00660 |
| 2492 | UK2016 | A-1.1 | A-662 | 0.70177 | 0.01126 |
| 2493 | UK2016 | A-450 | A-662 | 0.08582 | 0.00648 |

| | | | | | |
|---|---|---|---|---|---|
| 2494 | UK2016 | A-60 | A-662 | 0.70372 | 0.01348 |
| 2495 | UK2016 | A-1.2 | A-104.1 | 0.66608 | 0.01991 |
| 2496 | UK2016 | A-293 | A-104.1 | 0.48697 | 0.01883 |
| 2497 | UK2016 | A-322 | A-104.1 | 1.00000 | 0.00000 |
| 2498 | UK2016 | A-421 | A-104.1 | 0.98204 | 0.00437 |
| 2499 | UK2016 | A-498 | A-104.1 | 0.32443 | 0.01813 |
| 2500 | UK2016 | A-54 | A-104.1 | 0.90487 | 0.01250 |
| 2501 | UK2016 | A-275 | A-104.1 | 0.14969 | 0.00910 |
| 2502 | UK2016 | A-509 | A-104.1 | 0.86626 | 0.01536 |
| 2503 | UK2016 | A-600 | A-104.1 | 0.28818 | 0.01068 |
| 2504 | UK2016 | A-04 | A-104.1 | 0.81979 | 0.01011 |
| 2505 | UK2016 | A-092 | A-104.1 | 1.00000 | 0.00000 |
| 2506 | UK2016 | A-244 | A-104.1 | 0.38806 | 0.01607 |
| 2507 | UK2016 | A-342 | A-104.1 | 0.06526 | 0.00567 |
| 2508 | UK2016 | A-389 | A-104.1 | 0.75194 | 0.00679 |
| 2509 | UK2016 | A-547 | A-104.1 | 0.74969 | 0.01795 |
| 2510 | UK2016 | A-196 | A-104.1 | 0.04186 | 0.00707 |
| 2511 | UK2016 | A-360 | A-104.1 | 1.00000 | 0.00000 |
| 2512 | UK2016 | A-465 | A-104.1 | 0.07013 | 0.00872 |
| 2513 | UK2016 | A-161 | A-104.1 | 0.35122 | 0.02123 |
| 2514 | UK2016 | A-200 | A-104.1 | 0.99196 | 0.00357 |
| 2515 | UK2016 | A-325 | A-104.1 | 0.15376 | 0.01173 |
| 2516 | UK2016 | A-333 | A-104.1 | 0.31989 | 0.01835 |
| 2517 | UK2016 | A-104.2 | A-104.1 | 0.65676 | 0.01575 |
| 2518 | UK2016 | A-122 | A-104.1 | 0.93680 | 0.00881 |
| 2519 | UK2016 | A-175 | A-104.1 | 0.19448 | 0.01940 |
| 2520 | UK2016 | A-300 | A-104.1 | 0.32530 | 0.02765 |
| 2521 | UK2016 | A-733 | A-104.1 | 1.00000 | 0.00000 |
| 2522 | UK2016 | A-0 | A-104.1 | 0.44819 | 0.01118 |
| 2523 | UK2016 | A-1.1 | A-104.1 | 0.76091 | 0.01795 |
| 2524 | UK2016 | A-450 | A-104.1 | 0.44731 | 0.01906 |
| 2525 | UK2016 | A-60 | A-104.1 | 0.96594 | 0.00624 |
| 2526 | UK2016 | A-662 | A-104.1 | 1.00000 | 0.00000 |
| 2527 | UK2016 | A-1.2 | A-149 | 0.47008 | 0.02474 |
| 2528 | UK2016 | A-293 | A-149 | 0.98538 | 0.00374 |
| 2529 | UK2016 | A-322 | A-149 | 0.67858 | 0.00972 |
| 2530 | UK2016 | A-421 | A-149 | 0.15677 | 0.02128 |
| 2531 | UK2016 | A-498 | A-149 | 0.41691 | 0.02612 |
| 2532 | UK2016 | A-54 | A-149 | 0.60523 | 0.02015 |
| 2533 | UK2016 | A-275 | A-149 | 0.84516 | 0.01079 |
| 2534 | UK2016 | A-509 | A-149 | 0.00511 | 0.00243 |
| 2535 | UK2016 | A-600 | A-149 | 0.10420 | 0.00840 |
| 2536 | UK2016 | A-04 | A-149 | 0.17005 | 0.01386 |
| 2537 | UK2016 | A-092 | A-149 | 0.67310 | 0.01069 |
| 2538 | UK2016 | A-244 | A-149 | 0.64311 | 0.02101 |
| 2539 | UK2016 | A-342 | A-149 | 0.79147 | 0.01261 |
| 2540 | UK2016 | A-389 | A-149 | 0.91749 | 0.00501 |
| 2541 | UK2016 | A-547 | A-149 | 0.50878 | 0.02607 |

| | | | | | |
|---|---|---|---|---|---|
| 2542 | UK2016 | A-196 | A-149 | 0.34660 | 0.02465 |
| 2543 | UK2016 | A-360 | A-149 | 0.85575 | 0.01143 |
| 2544 | UK2016 | A-465 | A-149 | 0.23553 | 0.02304 |
| 2545 | UK2016 | A-161 | A-149 | 0.17625 | 0.02074 |
| 2546 | UK2016 | A-200 | A-149 | 0.89958 | 0.01415 |
| 2547 | UK2016 | A-325 | A-149 | 0.20539 | 0.01792 |
| 2548 | UK2016 | A-333 | A-149 | 0.57281 | 0.02471 |
| 2549 | UK2016 | A-104.2 | A-149 | 0.41985 | 0.02226 |
| 2550 | UK2016 | A-122 | A-149 | 0.56268 | 0.02482 |
| 2551 | UK2016 | A-175 | A-149 | 0.73259 | 0.02500 |
| 2552 | UK2016 | A-300 | A-149 | 0.03222 | 0.01153 |
| 2553 | UK2016 | A-733 | A-149 | 0.33765 | 0.01792 |
| 2554 | UK2016 | A-0 | A-149 | 0.99367 | 0.00170 |
| 2555 | UK2016 | A-1.1 | A-149 | 0.33998 | 0.02365 |
| 2556 | UK2016 | A-450 | A-149 | 0.23645 | 0.01802 |
| 2557 | UK2016 | A-60 | A-149 | 0.50747 | 0.02571 |
| 2558 | UK2016 | A-662 | A-149 | 0.69738 | 0.01015 |
| 2559 | UK2016 | A-104.1 | A-149 | 0.44703 | 0.01717 |
| 2560 | UK2016 | A-1.2 | A-224 | 0.00405 | 0.00150 |
| 2561 | UK2016 | A-293 | A-224 | 0.35332 | 0.02256 |
| 2562 | UK2016 | A-322 | A-224 | 0.00000 | 0.00000 |
| 2563 | UK2016 | A-421 | A-224 | 0.11282 | 0.01621 |
| 2564 | UK2016 | A-498 | A-224 | 0.55890 | 0.02588 |
| 2565 | UK2016 | A-54 | A-224 | 0.35420 | 0.01999 |
| 2566 | UK2016 | A-275 | A-224 | 0.00000 | 0.00000 |
| 2567 | UK2016 | A-509 | A-224 | 0.57979 | 0.02881 |
| 2568 | UK2016 | A-600 | A-224 | 0.77716 | 0.01319 |
| 2569 | UK2016 | A-04 | A-224 | 0.39040 | 0.01871 |
| 2570 | UK2016 | A-092 | A-224 | 0.92861 | 0.00392 |
| 2571 | UK2016 | A-244 | A-224 | 0.89646 | 0.01070 |
| 2572 | UK2016 | A-342 | A-224 | 0.62563 | 0.01793 |
| 2573 | UK2016 | A-389 | A-224 | 0.55877 | 0.01274 |
| 2574 | UK2016 | A-547 | A-224 | 0.70505 | 0.02439 |
| 2575 | UK2016 | A-196 | A-224 | 0.59396 | 0.02690 |
| 2576 | UK2016 | A-360 | A-224 | 0.47011 | 0.01733 |
| 2577 | UK2016 | A-465 | A-224 | 0.57583 | 0.02467 |
| 2578 | UK2016 | A-161 | A-224 | 0.55359 | 0.02850 |
| 2579 | UK2016 | A-200 | A-224 | 0.00000 | 0.00000 |
| 2580 | UK2016 | A-325 | A-224 | 0.35050 | 0.02302 |
| 2581 | UK2016 | A-333 | A-224 | 0.56114 | 0.02837 |
| 2582 | UK2016 | A-104.2 | A-224 | 0.93564 | 0.00913 |
| 2583 | UK2016 | A-122 | A-224 | 0.00000 | 0.00000 |
| 2584 | UK2016 | A-175 | A-224 | 0.31313 | 0.02982 |
| 2585 | UK2016 | A-300 | A-224 | 0.45607 | 0.03964 |
| 2586 | UK2016 | A-733 | A-224 | 0.08353 | 0.00924 |
| 2587 | UK2016 | A-0 | A-224 | 0.00000 | 0.00000 |
| 2588 | UK2016 | A-1.1 | A-224 | 0.29899 | 0.02294 |
| 2589 | UK2016 | A-450 | A-224 | 0.42573 | 0.02293 |

| | | | | | |
|---|---|---|---|---|---|
| 2590 | UK2016 | A-60 | A-224 | 0.08342 | 0.01439 |
| 2591 | UK2016 | A-662 | A-224 | 0.00000 | 0.00000 |
| 2592 | UK2016 | A-104.1 | A-224 | 0.97598 | 0.00524 |
| 2593 | UK2016 | A-149 | A-224 | 0.96136 | 0.00738 |
| 2594 | UK2016 | A-1.2 | A-323 | 0.99337 | 0.00452 |
| 2595 | UK2016 | A-293 | A-323 | 0.85116 | 0.01926 |
| 2596 | UK2016 | A-322 | A-323 | 0.48371 | 0.01714 |
| 2597 | UK2016 | A-421 | A-323 | 0.05210 | 0.01402 |
| 2598 | UK2016 | A-498 | A-323 | 0.42977 | 0.03839 |
| 2599 | UK2016 | A-54 | A-323 | 0.87107 | 0.01782 |
| 2600 | UK2016 | A-275 | A-323 | 0.69222 | 0.02057 |
| 2601 | UK2016 | A-509 | A-323 | 0.67404 | 0.03175 |
| 2602 | UK2016 | A-600 | A-323 | 0.69973 | 0.01964 |
| 2603 | UK2016 | A-04 | A-323 | 0.45914 | 0.02418 |
| 2604 | UK2016 | A-092 | A-323 | 0.69858 | 0.01524 |
| 2605 | UK2016 | A-244 | A-323 | 0.67482 | 0.02582 |
| 2606 | UK2016 | A-342 | A-323 | 0.17618 | 0.01790 |
| 2607 | UK2016 | A-389 | A-323 | 0.63166 | 0.01502 |
| 2608 | UK2016 | A-547 | A-323 | 0.17579 | 0.02360 |
| 2609 | UK2016 | A-196 | A-323 | 0.99241 | 0.00479 |
| 2610 | UK2016 | A-360 | A-323 | 0.67193 | 0.01895 |
| 2611 | UK2016 | A-465 | A-323 | 0.03330 | 0.01169 |
| 2612 | UK2016 | A-161 | A-323 | 0.29491 | 0.03381 |
| 2613 | UK2016 | A-200 | A-323 | 0.67952 | 0.03383 |
| 2614 | UK2016 | A-325 | A-323 | 0.44583 | 0.03290 |
| 2615 | UK2016 | A-333 | A-323 | 0.59723 | 0.03463 |
| 2616 | UK2016 | A-104.2 | A-323 | 0.83571 | 0.01667 |
| 2617 | UK2016 | A-122 | A-323 | 0.85855 | 0.02319 |
| 2618 | UK2016 | A-175 | A-323 | 0.85565 | 0.02290 |
| 2619 | UK2016 | A-300 | A-323 | 0.38968 | 0.04508 |
| 2620 | UK2016 | A-733 | A-323 | 0.60950 | 0.02360 |
| 2621 | UK2016 | A-0 | A-323 | 0.20532 | 0.01851 |
| 2622 | UK2016 | A-1.1 | A-323 | 0.39735 | 0.02943 |
| 2623 | UK2016 | A-450 | A-323 | 0.88331 | 0.01674 |
| 2624 | UK2016 | A-60 | A-323 | 0.16408 | 0.02655 |
| 2625 | UK2016 | A-662 | A-323 | 0.46520 | 0.01525 |
| 2626 | UK2016 | A-104.1 | A-323 | 0.38787 | 0.02271 |
| 2627 | UK2016 | A-149 | A-323 | 0.10280 | 0.02054 |
| 2628 | UK2016 | A-224 | A-323 | 0.81714 | 0.02246 |
| 2629 | UK2016 | A-1.2 | A-880 | 0.34661 | 0.01929 |
| 2630 | UK2016 | A-293 | A-880 | 0.61664 | 0.01918 |
| 2631 | UK2016 | A-322 | A-880 | 0.98214 | 0.00157 |
| 2632 | UK2016 | A-421 | A-880 | 0.00699 | 0.00252 |
| 2633 | UK2016 | A-498 | A-880 | 0.49435 | 0.02358 |
| 2634 | UK2016 | A-54 | A-880 | 0.37486 | 0.01992 |
| 2635 | UK2016 | A-275 | A-880 | 0.58343 | 0.01313 |
| 2636 | UK2016 | A-509 | A-880 | 0.22496 | 0.01897 |
| 2637 | UK2016 | A-600 | A-880 | 0.85759 | 0.00829 |

| | | | | | |
|---|---|---|---|---|---|
| 2638 | UK2016 | A-04 | A-880 | 0.05740 | 0.00716 |
| 2639 | UK2016 | A-092 | A-880 | 0.43593 | 0.00867 |
| 2640 | UK2016 | A-244 | A-880 | 0.04155 | 0.00566 |
| 2641 | UK2016 | A-342 | A-880 | 0.94712 | 0.00540 |
| 2642 | UK2016 | A-389 | A-880 | 0.66736 | 0.00997 |
| 2643 | UK2016 | A-547 | A-880 | 0.37760 | 0.02251 |
| 2644 | UK2016 | A-196 | A-880 | 0.35975 | 0.02189 |
| 2645 | UK2016 | A-360 | A-880 | 0.38807 | 0.01327 |
| 2646 | UK2016 | A-465 | A-880 | 0.18663 | 0.01640 |
| 2647 | UK2016 | A-161 | A-880 | 0.62538 | 0.02557 |
| 2648 | UK2016 | A-200 | A-880 | 0.68313 | 0.02005 |
| 2649 | UK2016 | A-325 | A-880 | 0.97566 | 0.00394 |
| 2650 | UK2016 | A-333 | A-880 | 0.26156 | 0.02161 |
| 2651 | UK2016 | A-104.2 | A-880 | 0.88569 | 0.01024 |
| 2652 | UK2016 | A-122 | A-880 | 0.33946 | 0.02121 |
| 2653 | UK2016 | A-175 | A-880 | 0.29715 | 0.02495 |
| 2654 | UK2016 | A-300 | A-880 | 0.77102 | 0.02755 |
| 2655 | UK2016 | A-733 | A-880 | 1.00000 | 0.00000 |
| 2656 | UK2016 | A-0 | A-880 | 0.88766 | 0.00728 |
| 2657 | UK2016 | A-1.1 | A-880 | 0.04179 | 0.00825 |
| 2658 | UK2016 | A-450 | A-880 | 0.37601 | 0.01769 |
| 2659 | UK2016 | A-60 | A-880 | 0.44066 | 0.02510 |
| 2660 | UK2016 | A-662 | A-880 | 0.98611 | 0.00123 |
| 2661 | UK2016 | A-104.1 | A-880 | 0.67580 | 0.01429 |
| 2662 | UK2016 | A-149 | A-880 | 0.96132 | 0.00607 |
| 2663 | UK2016 | A-224 | A-880 | 0.76062 | 0.01601 |
| 2664 | UK2016 | A-323 | A-880 | 0.76000 | 0.02124 |

**Table 2, S3**. Linkage Disequilibrium tests for the panel of 29 loci

| | POP | Locus#1 | Locus#2 | P-Value | S.E. |
|---|---|---|---|---|---|
| 1 | LK2016 | TDM_1.2 | TDM_293 | 0.04083 | 0.00780 |
| 2 | LK2016 | TDM_1.2 | TDM_421 | 0.66944 | 0.02916 |
| 3 | LK2016 | TDM_293 | TDM_421 | 0.12727 | 0.01186 |
| 4 | LK2016 | TDM_1.2 | TDM_498 | 0.76695 | 0.01823 |
| 5 | LK2016 | TDM_293 | TDM_498 | 0.64453 | 0.00984 |
| 6 | LK2016 | TDM_421 | TDM_498 | 0.60759 | 0.01807 |
| 7 | LK2016 | TDM_1.2 | TDM_54 | 0.28478 | 0.01322 |
| 8 | LK2016 | TDM_293 | TDM_54 | 0.84462 | 0.00391 |
| 9 | LK2016 | TDM_421 | TDM_54 | 0.77564 | 0.00792 |
| 10 | LK2016 | TDM_498 | TDM_54 | 0.45726 | 0.00850 |
| 11 | LK2016 | TDM_1.2 | TDM_275 | 0.13662 | 0.01466 |
| 12 | LK2016 | TDM_293 | TDM_275 | 0.79147 | 0.00889 |
| 13 | LK2016 | TDM_421 | TDM_275 | 0.11286 | 0.01160 |
| 14 | LK2016 | TDM_498 | TDM_275 | 0.20837 | 0.01158 |
| 15 | LK2016 | TDM_54 | TDM_275 | 0.74373 | 0.00520 |
| 16 | LK2016 | TDM_1.2 | TDM_509 | 0.11218 | 0.02119 |
| 17 | LK2016 | TDM_293 | TDM_509 | 0.64358 | 0.01960 |
| 18 | LK2016 | TDM_421 | TDM_509 | 0.48831 | 0.02967 |
| 19 | LK2016 | TDM_498 | TDM_509 | 0.30210 | 0.02189 |
| 20 | LK2016 | TDM_54 | TDM_509 | 0.90753 | 0.00655 |
| 21 | LK2016 | TDM_275 | TDM_509 | 0.56530 | 0.02138 |
| 22 | LK2016 | TDM_1.2 | TDM_04 | 0.76452 | 0.00953 |
| 23 | LK2016 | TDM_293 | TDM_04 | 0.50733 | 0.00555 |
| 24 | LK2016 | TDM_421 | TDM_04 | 0.77037 | 0.00701 |
| 25 | LK2016 | TDM_498 | TDM_04 | 0.86251 | 0.00438 |
| 26 | LK2016 | TDM_54 | TDM_04 | 0.38726 | 0.00485 |
| 27 | LK2016 | TDM_275 | TDM_04 | 0.08283 | 0.00385 |
| 28 | LK2016 | TDM_509 | TDM_04 | 0.18345 | 0.01129 |
| 29 | LK2016 | TDM_1.2 | TDM_244 | 0.65608 | 0.02210 |
| 30 | LK2016 | TDM_293 | TDM_244 | 0.62316 | 0.01032 |
| 31 | LK2016 | TDM_421 | TDM_244 | 0.48009 | 0.01829 |
| 32 | LK2016 | TDM_498 | TDM_244 | 0.49799 | 0.01289 |
| 33 | LK2016 | TDM_54 | TDM_244 | 0.00016 | 0.00008 |
| 34 | LK2016 | TDM_275 | TDM_244 | 0.57234 | 0.01254 |
| 35 | LK2016 | TDM_509 | TDM_244 | 0.82331 | 0.01558 |
| 36 | LK2016 | TDM_04 | TDM_244 | 0.64790 | 0.00711 |
| 37 | LK2016 | TDM_1.2 | TDM_342 | 0.49722 | 0.02263 |
| 38 | LK2016 | TDM_293 | TDM_342 | 0.09758 | 0.00574 |
| 39 | LK2016 | TDM_421 | TDM_342 | 0.04336 | 0.00575 |
| 40 | LK2016 | TDM_498 | TDM_342 | 0.40701 | 0.01373 |
| 41 | LK2016 | TDM_54 | TDM_342 | 0.37995 | 0.00714 |
| 42 | LK2016 | TDM_275 | TDM_342 | 0.93822 | 0.00419 |
| 43 | LK2016 | TDM_509 | TDM_342 | 0.43972 | 0.01964 |
| 44 | LK2016 | TDM_04 | TDM_342 | 0.54730 | 0.00784 |
| 45 | LK2016 | TDM_244 | TDM_342 | 0.68842 | 0.01240 |

| | | | | | |
|---|---|---|---|---|---|
| 46 | LK2016 | TDM_1.2 | TDM_389 | 0.94350 | 0.00881 |
| 47 | LK2016 | TDM_293 | TDM_389 | 0.56256 | 0.01208 |
| 48 | LK2016 | TDM_421 | TDM_389 | 0.53957 | 0.01748 |
| 49 | LK2016 | TDM_498 | TDM_389 | 0.27157 | 0.01172 |
| 50 | LK2016 | TDM_54 | TDM_389 | 0.96142 | 0.00206 |
| 51 | LK2016 | TDM_275 | TDM_389 | 0.87524 | 0.00652 |
| 52 | LK2016 | TDM_509 | TDM_389 | 0.96719 | 0.00517 |
| 53 | LK2016 | TDM_04 | TDM_389 | 0.18903 | 0.00627 |
| 54 | LK2016 | TDM_244 | TDM_389 | 0.88164 | 0.00649 |
| 55 | LK2016 | TDM_342 | TDM_389 | 0.38281 | 0.01212 |
| 56 | LK2016 | TDM_1.2 | TDM_547 | 0.64150 | 0.01855 |
| 57 | LK2016 | TDM_293 | TDM_547 | 0.08656 | 0.00554 |
| 58 | LK2016 | TDM_421 | TDM_547 | 0.91300 | 0.00699 |
| 59 | LK2016 | TDM_498 | TDM_547 | 0.23473 | 0.00947 |
| 60 | LK2016 | TDM_54 | TDM_547 | 0.24923 | 0.00550 |
| 61 | LK2016 | TDM_275 | TDM_547 | 0.55842 | 0.00995 |
| 62 | LK2016 | TDM_509 | TDM_547 | 0.96169 | 0.00495 |
| 63 | LK2016 | TDM_04 | TDM_547 | 0.75508 | 0.00583 |
| 64 | LK2016 | TDM_244 | TDM_547 | 0.22818 | 0.00991 |
| 65 | LK2016 | TDM_342 | TDM_547 | 0.69866 | 0.00988 |
| 66 | LK2016 | TDM_389 | TDM_547 | 0.45673 | 0.01052 |
| 67 | LK2016 | TDM_1.2 | TDM_196 | 0.86606 | 0.02180 |
| 68 | LK2016 | TDM_293 | TDM_196 | 0.16872 | 0.01185 |
| 69 | LK2016 | TDM_421 | TDM_196 | 0.44567 | 0.02952 |
| 70 | LK2016 | TDM_498 | TDM_196 | 0.85752 | 0.01400 |
| 71 | LK2016 | TDM_54 | TDM_196 | 0.99857 | 0.00040 |
| 72 | LK2016 | TDM_275 | TDM_196 | 0.89460 | 0.00931 |
| 73 | LK2016 | TDM_509 | TDM_196 | 0.54406 | 0.02929 |
| 74 | LK2016 | TDM_04 | TDM_196 | 0.47246 | 0.01231 |
| 75 | LK2016 | TDM_244 | TDM_196 | 0.59970 | 0.01923 |
| 76 | LK2016 | TDM_342 | TDM_196 | 0.33413 | 0.01764 |
| 77 | LK2016 | TDM_389 | TDM_196 | 0.51841 | 0.01937 |
| 78 | LK2016 | TDM_547 | TDM_196 | 0.69542 | 0.01327 |
| 79 | LK2016 | TDM_1.2 | TDM_360 | 0.50474 | 0.01027 |
| 80 | LK2016 | TDM_293 | TDM_360 | 0.01814 | 0.00142 |
| 81 | LK2016 | TDM_421 | TDM_360 | 0.20583 | 0.00589 |
| 82 | LK2016 | TDM_498 | TDM_360 | 0.76264 | 0.00422 |
| 83 | LK2016 | TDM_54 | TDM_360 | 0.21954 | 0.00319 |
| 84 | LK2016 | TDM_275 | TDM_360 | 0.80951 | 0.00291 |
| 85 | LK2016 | TDM_509 | TDM_360 | 0.40814 | 0.00835 |
| 86 | LK2016 | TDM_04 | TDM_360 | 0.87040 | 0.00181 |
| 87 | LK2016 | TDM_244 | TDM_360 | 0.79627 | 0.00372 |
| 88 | LK2016 | TDM_342 | TDM_360 | 0.03054 | 0.00176 |
| 89 | LK2016 | TDM_389 | TDM_360 | 0.77628 | 0.00476 |
| 90 | LK2016 | TDM_547 | TDM_360 | 0.37528 | 0.00422 |
| 91 | LK2016 | TDM_196 | TDM_360 | 0.08014 | 0.00420 |
| 92 | LK2016 | TDM_1.2 | TDM_465 | 0.87989 | 0.02401 |
| 93 | LK2016 | TDM_293 | TDM_465 | 0.52737 | 0.02089 |

| | | | | | |
|---|---|---|---|---|---|
| 94 | LK2016 | TDM_421 | TDM_465 | 0.04720 | 0.01052 |
| 95 | LK2016 | TDM_498 | TDM_465 | 0.74321 | 0.01933 |
| 96 | LK2016 | TDM_54 | TDM_465 | 0.09833 | 0.00820 |
| 97 | LK2016 | TDM_275 | TDM_465 | 0.01913 | 0.00441 |
| 98 | LK2016 | TDM_509 | TDM_465 | 0.08507 | 0.01974 |
| 99 | LK2016 | TDM_04 | TDM_465 | 0.62968 | 0.01156 |
| 100 | LK2016 | TDM_244 | TDM_465 | 0.94442 | 0.00805 |
| 101 | LK2016 | TDM_342 | TDM_465 | 0.38804 | 0.02264 |
| 102 | LK2016 | TDM_389 | TDM_465 | 0.58331 | 0.02418 |
| 103 | LK2016 | TDM_547 | TDM_465 | 0.21395 | 0.01649 |
| 104 | LK2016 | TDM_196 | TDM_465 | 0.76404 | 0.03059 |
| 105 | LK2016 | TDM_360 | TDM_465 | 0.19085 | 0.00794 |
| 106 | LK2016 | TDM_1.2 | TDM_161 | 0.64322 | 0.02993 |
| 107 | LK2016 | TDM_293 | TDM_161 | 0.70225 | 0.01476 |
| 108 | LK2016 | TDM_421 | TDM_161 | 0.79078 | 0.01670 |
| 109 | LK2016 | TDM_498 | TDM_161 | 0.55122 | 0.01885 |
| 110 | LK2016 | TDM_54 | TDM_161 | 0.25318 | 0.00887 |
| 111 | LK2016 | TDM_275 | TDM_161 | 0.76395 | 0.01256 |
| 112 | LK2016 | TDM_509 | TDM_161 | 0.87404 | 0.01491 |
| 113 | LK2016 | TDM_04 | TDM_161 | 0.42279 | 0.01122 |
| 114 | LK2016 | TDM_244 | TDM_161 | 0.55133 | 0.01659 |
| 115 | LK2016 | TDM_342 | TDM_161 | 0.48330 | 0.01875 |
| 116 | LK2016 | TDM_389 | TDM_161 | 0.69221 | 0.01269 |
| 117 | LK2016 | TDM_547 | TDM_161 | 0.68417 | 0.01235 |
| 118 | LK2016 | TDM_196 | TDM_161 | 0.43878 | 0.02647 |
| 119 | LK2016 | TDM_360 | TDM_161 | 0.76323 | 0.00596 |
| 120 | LK2016 | TDM_465 | TDM_161 | 0.51737 | 0.03168 |
| 121 | LK2016 | TDM_1.2 | TDM_200 | 0.00436 | 0.00222 |
| 122 | LK2016 | TDM_293 | TDM_200 | 0.47768 | 0.01195 |
| 123 | LK2016 | TDM_421 | TDM_200 | 0.01814 | 0.00345 |
| 124 | LK2016 | TDM_498 | TDM_200 | 0.24549 | 0.01301 |
| 125 | LK2016 | TDM_54 | TDM_200 | 0.96923 | 0.00155 |
| 126 | LK2016 | TDM_275 | TDM_200 | 0.00875 | 0.00216 |
| 127 | LK2016 | TDM_509 | TDM_200 | 0.92532 | 0.00876 |
| 128 | LK2016 | TDM_04 | TDM_200 | 0.75537 | 0.00537 |
| 129 | LK2016 | TDM_244 | TDM_200 | 0.04985 | 0.00473 |
| 130 | LK2016 | TDM_342 | TDM_200 | 0.40089 | 0.01199 |
| 131 | LK2016 | TDM_389 | TDM_200 | 0.55310 | 0.01231 |
| 132 | LK2016 | TDM_547 | TDM_200 | 0.34276 | 0.00996 |
| 133 | LK2016 | TDM_196 | TDM_200 | 0.11283 | 0.01150 |
| 134 | LK2016 | TDM_360 | TDM_200 | 0.38283 | 0.00519 |
| 135 | LK2016 | TDM_465 | TDM_200 | 0.35573 | 0.02405 |
| 136 | LK2016 | TDM_161 | TDM_200 | 0.25731 | 0.01654 |
| 137 | LK2016 | TDM_1.2 | TDM_325 | 0.75067 | 0.01939 |
| 138 | LK2016 | TDM_293 | TDM_325 | 0.40892 | 0.01278 |
| 139 | LK2016 | TDM_421 | TDM_325 | 0.94954 | 0.00734 |
| 140 | LK2016 | TDM_498 | TDM_325 | 0.49795 | 0.01367 |
| 141 | LK2016 | TDM_54 | TDM_325 | 0.70144 | 0.00734 |

| | | | | | |
|---|---|---|---|---|---|
| 142 | LK2016 | TDM_275 | TDM_325 | 0.55776 | 0.01228 |
| 143 | LK2016 | TDM_509 | TDM_325 | 0.74383 | 0.02025 |
| 144 | LK2016 | TDM_04 | TDM_325 | 0.01440 | 0.00154 |
| 145 | LK2016 | TDM_244 | TDM_325 | 0.31297 | 0.01440 |
| 146 | LK2016 | TDM_342 | TDM_325 | 0.03944 | 0.00487 |
| 147 | LK2016 | TDM_389 | TDM_325 | 0.50434 | 0.01475 |
| 148 | LK2016 | TDM_547 | TDM_325 | 0.48436 | 0.01152 |
| 149 | LK2016 | TDM_196 | TDM_325 | 0.24473 | 0.01760 |
| 150 | LK2016 | TDM_360 | TDM_325 | 0.16010 | 0.00414 |
| 151 | LK2016 | TDM_465 | TDM_325 | 0.91095 | 0.01338 |
| 152 | LK2016 | TDM_161 | TDM_325 | 0.33874 | 0.01812 |
| 153 | LK2016 | TDM_200 | TDM_325 | 0.95655 | 0.00372 |
| 154 | LK2016 | TDM_1.2 | TDM_333 | 0.03031 | 0.00806 |
| 155 | LK2016 | TDM_293 | TDM_333 | 0.47221 | 0.01406 |
| 156 | LK2016 | TDM_421 | TDM_333 | 0.30327 | 0.02015 |
| 157 | LK2016 | TDM_498 | TDM_333 | 0.93721 | 0.00576 |
| 158 | LK2016 | TDM_54 | TDM_333 | 0.46181 | 0.00913 |
| 159 | LK2016 | TDM_275 | TDM_333 | 0.38918 | 0.01260 |
| 160 | LK2016 | TDM_509 | TDM_333 | 0.25072 | 0.02331 |
| 161 | LK2016 | TDM_04 | TDM_333 | 0.44242 | 0.00934 |
| 162 | LK2016 | TDM_244 | TDM_333 | 0.31621 | 0.01320 |
| 163 | LK2016 | TDM_342 | TDM_333 | 0.35621 | 0.01442 |
| 164 | LK2016 | TDM_389 | TDM_333 | 0.55618 | 0.01417 |
| 165 | LK2016 | TDM_547 | TDM_333 | 0.37754 | 0.01282 |
| 166 | LK2016 | TDM_196 | TDM_333 | 0.16838 | 0.01753 |
| 167 | LK2016 | TDM_360 | TDM_333 | 0.52743 | 0.00632 |
| 168 | LK2016 | TDM_465 | TDM_333 | 0.57929 | 0.02930 |
| 169 | LK2016 | TDM_161 | TDM_333 | 0.47665 | 0.02018 |
| 170 | LK2016 | TDM_200 | TDM_333 | 0.51392 | 0.01678 |
| 171 | LK2016 | TDM_325 | TDM_333 | 0.83309 | 0.01095 |
| 172 | LK2016 | TDM_1.2 | TDM_122 | 0.05663 | 0.01281 |
| 173 | LK2016 | TDM_293 | TDM_122 | 0.34073 | 0.01219 |
| 174 | LK2016 | TDM_421 | TDM_122 | 0.17914 | 0.01460 |
| 175 | LK2016 | TDM_498 | TDM_122 | 0.45341 | 0.01559 |
| 176 | LK2016 | TDM_54 | TDM_122 | 0.87687 | 0.00411 |
| 177 | LK2016 | TDM_275 | TDM_122 | 0.00000 | 0.00000 |
| 178 | LK2016 | TDM_509 | TDM_122 | 0.65977 | 0.02245 |
| 179 | LK2016 | TDM_04 | TDM_122 | 0.25634 | 0.00823 |
| 180 | LK2016 | TDM_244 | TDM_122 | 0.65692 | 0.01157 |
| 181 | LK2016 | TDM_342 | TDM_122 | 0.01364 | 0.00229 |
| 182 | LK2016 | TDM_389 | TDM_122 | 0.87629 | 0.00688 |
| 183 | LK2016 | TDM_547 | TDM_122 | 0.30530 | 0.01106 |
| 184 | LK2016 | TDM_196 | TDM_122 | 0.99527 | 0.00151 |
| 185 | LK2016 | TDM_360 | TDM_122 | 0.63005 | 0.00477 |
| 186 | LK2016 | TDM_465 | TDM_122 | 0.20970 | 0.02025 |
| 187 | LK2016 | TDM_161 | TDM_122 | 0.84897 | 0.01198 |
| 188 | LK2016 | TDM_200 | TDM_122 | 0.03085 | 0.00465 |
| 189 | LK2016 | TDM_325 | TDM_122 | 0.72158 | 0.01205 |

| | | | | | |
|---|---|---|---|---|---|
| 190 | LK2016 | TDM_333 | TDM_122 | 0.09490 | 0.00885 |
| 191 | LK2016 | TDM_1.2 | TDM_175 | 0.70410 | 0.02289 |
| 192 | LK2016 | TDM_293 | TDM_175 | 0.65446 | 0.01113 |
| 193 | LK2016 | TDM_421 | TDM_175 | 0.22114 | 0.01710 |
| 194 | LK2016 | TDM_498 | TDM_175 | 0.46412 | 0.01241 |
| 195 | LK2016 | TDM_54 | TDM_175 | 0.94201 | 0.00245 |
| 196 | LK2016 | TDM_275 | TDM_175 | 0.27121 | 0.01064 |
| 197 | LK2016 | TDM_509 | TDM_175 | 0.76943 | 0.01690 |
| 198 | LK2016 | TDM_04 | TDM_175 | 0.45900 | 0.00733 |
| 199 | LK2016 | TDM_244 | TDM_175 | 0.23916 | 0.00999 |
| 200 | LK2016 | TDM_342 | TDM_175 | 0.01450 | 0.00238 |
| 201 | LK2016 | TDM_389 | TDM_175 | 0.67414 | 0.01087 |
| 202 | LK2016 | TDM_547 | TDM_175 | 0.04447 | 0.00391 |
| 203 | LK2016 | TDM_196 | TDM_175 | 0.91795 | 0.00872 |
| 204 | LK2016 | TDM_360 | TDM_175 | 0.22521 | 0.00433 |
| 205 | LK2016 | TDM_465 | TDM_175 | 0.57177 | 0.02483 |
| 206 | LK2016 | TDM_161 | TDM_175 | 0.03427 | 0.00485 |
| 207 | LK2016 | TDM_200 | TDM_175 | 0.37282 | 0.01291 |
| 208 | LK2016 | TDM_325 | TDM_175 | 0.33403 | 0.01369 |
| 209 | LK2016 | TDM_333 | TDM_175 | 0.11212 | 0.00900 |
| 210 | LK2016 | TDM_122 | TDM_175 | 0.18746 | 0.01182 |
| 211 | LK2016 | TDM_1.2 | TDM_300 | 0.40566 | 0.03330 |
| 212 | LK2016 | TDM_293 | TDM_300 | 0.06878 | 0.00829 |
| 213 | LK2016 | TDM_421 | TDM_300 | 0.39524 | 0.02898 |
| 214 | LK2016 | TDM_498 | TDM_300 | 0.86481 | 0.01373 |
| 215 | LK2016 | TDM_54 | TDM_300 | 0.95367 | 0.00419 |
| 216 | LK2016 | TDM_275 | TDM_300 | 0.71781 | 0.01796 |
| 217 | LK2016 | TDM_509 | TDM_300 | 0.08874 | 0.01896 |
| 218 | LK2016 | TDM_04 | TDM_300 | 0.38534 | 0.01455 |
| 219 | LK2016 | TDM_244 | TDM_300 | 0.89925 | 0.00787 |
| 220 | LK2016 | TDM_342 | TDM_300 | 0.09256 | 0.01339 |
| 221 | LK2016 | TDM_389 | TDM_300 | 0.19315 | 0.01606 |
| 222 | LK2016 | TDM_547 | TDM_300 | 0.10784 | 0.01181 |
| 223 | LK2016 | TDM_196 | TDM_300 | 0.60544 | 0.03048 |
| 224 | LK2016 | TDM_360 | TDM_300 | 0.57317 | 0.00881 |
| 225 | LK2016 | TDM_465 | TDM_300 | 0.87875 | 0.02370 |
| 226 | LK2016 | TDM_161 | TDM_300 | 0.70010 | 0.02783 |
| 227 | LK2016 | TDM_200 | TDM_300 | 0.72421 | 0.01906 |
| 228 | LK2016 | TDM_325 | TDM_300 | 0.00513 | 0.00263 |
| 229 | LK2016 | TDM_333 | TDM_300 | 0.60819 | 0.02642 |
| 230 | LK2016 | TDM_122 | TDM_300 | 0.61598 | 0.02441 |
| 231 | LK2016 | TDM_175 | TDM_300 | 0.00332 | 0.00117 |
| 232 | LK2016 | TDM_1.2 | TDM_1.1 | 0.00000 | 0.00000 |
| 233 | LK2016 | TDM_293 | TDM_1.1 | 0.05320 | 0.00486 |
| 234 | LK2016 | TDM_421 | TDM_1.1 | 0.43291 | 0.02131 |
| 235 | LK2016 | TDM_498 | TDM_1.1 | 0.52002 | 0.01411 |
| 236 | LK2016 | TDM_54 | TDM_1.1 | 0.68340 | 0.00688 |
| 237 | LK2016 | TDM_275 | TDM_1.1 | 0.03325 | 0.00413 |

| | | | | | |
|---|---|---|---|---|---|
| 238 | LK2016 | TDM_509 | TDM_1.1 | 0.73225 | 0.01834 |
| 239 | LK2016 | TDM_04 | TDM_1.1 | 0.04691 | 0.00319 |
| 240 | LK2016 | TDM_244 | TDM_1.1 | 0.26475 | 0.01436 |
| 241 | LK2016 | TDM_342 | TDM_1.1 | 0.74916 | 0.01191 |
| 242 | LK2016 | TDM_389 | TDM_1.1 | 0.25561 | 0.01354 |
| 243 | LK2016 | TDM_547 | TDM_1.1 | 0.34520 | 0.01184 |
| 244 | LK2016 | TDM_196 | TDM_1.1 | 0.76571 | 0.01503 |
| 245 | LK2016 | TDM_360 | TDM_1.1 | 0.03281 | 0.00209 |
| 246 | LK2016 | TDM_465 | TDM_1.1 | 0.21140 | 0.02176 |
| 247 | LK2016 | TDM_161 | TDM_1.1 | 0.61393 | 0.02043 |
| 248 | LK2016 | TDM_200 | TDM_1.1 | 0.00005 | 0.00004 |
| 249 | LK2016 | TDM_325 | TDM_1.1 | 0.53796 | 0.01503 |
| 250 | LK2016 | TDM_333 | TDM_1.1 | 0.18426 | 0.01262 |
| 251 | LK2016 | TDM_122 | TDM_1.1 | 0.60376 | 0.01385 |
| 252 | LK2016 | TDM_175 | TDM_1.1 | 0.61115 | 0.01324 |
| 253 | LK2016 | TDM_300 | TDM_1.1 | 0.36434 | 0.02147 |
| 254 | LK2016 | TDM_1.2 | TDM_450 | 0.23009 | 0.01951 |
| 255 | LK2016 | TDM_293 | TDM_450 | 0.08176 | 0.00516 |
| 256 | LK2016 | TDM_421 | TDM_450 | 0.08389 | 0.00631 |
| 257 | LK2016 | TDM_498 | TDM_450 | 0.16772 | 0.00783 |
| 258 | LK2016 | TDM_54 | TDM_450 | 0.35576 | 0.00563 |
| 259 | LK2016 | TDM_275 | TDM_450 | 0.52291 | 0.00789 |
| 260 | LK2016 | TDM_509 | TDM_450 | 0.57608 | 0.01556 |
| 261 | LK2016 | TDM_04 | TDM_450 | 0.01592 | 0.00138 |
| 262 | LK2016 | TDM_244 | TDM_450 | 0.48452 | 0.01008 |
| 263 | LK2016 | TDM_342 | TDM_450 | 0.00224 | 0.00059 |
| 264 | LK2016 | TDM_389 | TDM_450 | 0.76353 | 0.00744 |
| 265 | LK2016 | TDM_547 | TDM_450 | 0.60243 | 0.00864 |
| 266 | LK2016 | TDM_196 | TDM_450 | 0.60810 | 0.01531 |
| 267 | LK2016 | TDM_360 | TDM_450 | 0.50707 | 0.00391 |
| 268 | LK2016 | TDM_465 | TDM_450 | 0.91833 | 0.00815 |
| 269 | LK2016 | TDM_161 | TDM_450 | 0.24647 | 0.01037 |
| 270 | LK2016 | TDM_200 | TDM_450 | 0.45828 | 0.01061 |
| 271 | LK2016 | TDM_325 | TDM_450 | 0.79486 | 0.00803 |
| 272 | LK2016 | TDM_333 | TDM_450 | 0.71058 | 0.00936 |
| 273 | LK2016 | TDM_122 | TDM_450 | 0.33944 | 0.00998 |
| 274 | LK2016 | TDM_175 | TDM_450 | 0.18690 | 0.00707 |
| 275 | LK2016 | TDM_300 | TDM_450 | 0.29370 | 0.01576 |
| 276 | LK2016 | TDM_1.1 | TDM_450 | 0.97810 | 0.00207 |
| 277 | LK2016 | TDM_1.2 | TDM_60 | 0.64088 | 0.03429 |
| 278 | LK2016 | TDM_293 | TDM_60 | 0.95148 | 0.00682 |
| 279 | LK2016 | TDM_421 | TDM_60 | 0.20680 | 0.02488 |
| 280 | LK2016 | TDM_498 | TDM_60 | 0.98602 | 0.00342 |
| 281 | LK2016 | TDM_54 | TDM_60 | 0.66307 | 0.01004 |
| 282 | LK2016 | TDM_275 | TDM_60 | 0.11166 | 0.01130 |
| 283 | LK2016 | TDM_509 | TDM_60 | 0.99976 | 0.00024 |
| 284 | LK2016 | TDM_04 | TDM_60 | 0.15064 | 0.00855 |
| 285 | LK2016 | TDM_244 | TDM_60 | 0.60295 | 0.02008 |

| | | | | | |
|---|---|---|---|---|---|
| 286 | LK2016 | TDM_342 | TDM_60 | 0.88290 | 0.01076 |
| 287 | LK2016 | TDM_389 | TDM_60 | 0.35086 | 0.02076 |
| 288 | LK2016 | TDM_547 | TDM_60 | 0.97395 | 0.00310 |
| 289 | LK2016 | TDM_196 | TDM_60 | 0.21647 | 0.02577 |
| 290 | LK2016 | TDM_360 | TDM_60 | 0.95937 | 0.00246 |
| 291 | LK2016 | TDM_465 | TDM_60 | 0.56992 | 0.03541 |
| 292 | LK2016 | TDM_161 | TDM_60 | 0.39038 | 0.02879 |
| 293 | LK2016 | TDM_200 | TDM_60 | 0.00127 | 0.00074 |
| 294 | LK2016 | TDM_325 | TDM_60 | 0.07525 | 0.01138 |
| 295 | LK2016 | TDM_333 | TDM_60 | 0.86923 | 0.01455 |
| 296 | LK2016 | TDM_122 | TDM_60 | 0.53552 | 0.02129 |
| 297 | LK2016 | TDM_175 | TDM_60 | 0.65597 | 0.01985 |
| 298 | LK2016 | TDM_300 | TDM_60 | 0.15095 | 0.02305 |
| 299 | LK2016 | TDM_1.1 | TDM_60 | 0.00882 | 0.00321 |
| 300 | LK2016 | TDM_450 | TDM_60 | 0.60362 | 0.01459 |
| 301 | LK2016 | TDM_1.2 | TDM_149 | 0.14532 | 0.01910 |
| 302 | LK2016 | TDM_293 | TDM_149 | 0.01944 | 0.00304 |
| 303 | LK2016 | TDM_421 | TDM_149 | 0.53109 | 0.02034 |
| 304 | LK2016 | TDM_498 | TDM_149 | 0.53881 | 0.01394 |
| 305 | LK2016 | TDM_54 | TDM_149 | 0.84283 | 0.00463 |
| 306 | LK2016 | TDM_275 | TDM_149 | 0.51537 | 0.01326 |
| 307 | LK2016 | TDM_509 | TDM_149 | 0.65825 | 0.02194 |
| 308 | LK2016 | TDM_04 | TDM_149 | 0.88523 | 0.00348 |
| 309 | LK2016 | TDM_244 | TDM_149 | 0.96921 | 0.00379 |
| 310 | LK2016 | TDM_342 | TDM_149 | 0.57766 | 0.01391 |
| 311 | LK2016 | TDM_389 | TDM_149 | 0.04797 | 0.00441 |
| 312 | LK2016 | TDM_547 | TDM_149 | 0.00623 | 0.00147 |
| 313 | LK2016 | TDM_196 | TDM_149 | 0.14459 | 0.01701 |
| 314 | LK2016 | TDM_360 | TDM_149 | 0.30264 | 0.00696 |
| 315 | LK2016 | TDM_465 | TDM_149 | 0.09446 | 0.01296 |
| 316 | LK2016 | TDM_161 | TDM_149 | 0.01272 | 0.00348 |
| 317 | LK2016 | TDM_200 | TDM_149 | 0.18632 | 0.01039 |
| 318 | LK2016 | TDM_325 | TDM_149 | 0.48433 | 0.01524 |
| 319 | LK2016 | TDM_333 | TDM_149 | 0.93828 | 0.00613 |
| 320 | LK2016 | TDM_122 | TDM_149 | 0.30205 | 0.01380 |
| 321 | LK2016 | TDM_175 | TDM_149 | 0.16480 | 0.01061 |
| 322 | LK2016 | TDM_300 | TDM_149 | 0.65400 | 0.02682 |
| 323 | LK2016 | TDM_1.1 | TDM_149 | 0.47572 | 0.01583 |
| 324 | LK2016 | TDM_450 | TDM_149 | 0.42883 | 0.01025 |
| 325 | LK2016 | TDM_60 | TDM_149 | 0.68101 | 0.02114 |
| 326 | LK2016 | TDM_1.2 | TDM_224 | 0.59562 | 0.03207 |
| 327 | LK2016 | TDM_293 | TDM_224 | 0.37136 | 0.01840 |
| 328 | LK2016 | TDM_421 | TDM_224 | 0.62742 | 0.02439 |
| 329 | LK2016 | TDM_498 | TDM_224 | 0.44710 | 0.01919 |
| 330 | LK2016 | TDM_54 | TDM_224 | 0.43479 | 0.01006 |
| 331 | LK2016 | TDM_275 | TDM_224 | 0.00000 | 0.00000 |
| 332 | LK2016 | TDM_509 | TDM_224 | 0.93970 | 0.01219 |
| 333 | LK2016 | TDM_04 | TDM_224 | 0.86727 | 0.00504 |

| | | | | | |
|---|---|---|---|---|---|
| 334 | LK2016 | TDM_244 | TDM_224 | 0.07338 | 0.01003 |
| 335 | LK2016 | TDM_342 | TDM_224 | 0.86742 | 0.00984 |
| 336 | LK2016 | TDM_389 | TDM_224 | 0.59866 | 0.01728 |
| 337 | LK2016 | TDM_547 | TDM_224 | 0.01797 | 0.00335 |
| 338 | LK2016 | TDM_196 | TDM_224 | 0.73453 | 0.02397 |
| 339 | LK2016 | TDM_360 | TDM_224 | 0.76431 | 0.00570 |
| 340 | LK2016 | TDM_465 | TDM_224 | 0.45013 | 0.03230 |
| 341 | LK2016 | TDM_161 | TDM_224 | 0.30325 | 0.02014 |
| 342 | LK2016 | TDM_200 | TDM_224 | 0.00336 | 0.00125 |
| 343 | LK2016 | TDM_325 | TDM_224 | 0.77946 | 0.01419 |
| 344 | LK2016 | TDM_333 | TDM_224 | 0.89973 | 0.00999 |
| 345 | LK2016 | TDM_122 | TDM_224 | 0.00000 | 0.00000 |
| 346 | LK2016 | TDM_175 | TDM_224 | 0.79404 | 0.01473 |
| 347 | LK2016 | TDM_300 | TDM_224 | 0.73540 | 0.02579 |
| 348 | LK2016 | TDM_1.1 | TDM_224 | 0.29593 | 0.01700 |
| 349 | LK2016 | TDM_450 | TDM_224 | 0.56287 | 0.01308 |
| 350 | LK2016 | TDM_60 | TDM_224 | 0.02218 | 0.00578 |
| 351 | LK2016 | TDM_149 | TDM_224 | 0.00834 | 0.00261 |
| 352 | LK2016 | TDM_1.2 | TDM_323 | 0.78774 | 0.03124 |
| 353 | LK2016 | TDM_293 | TDM_323 | 0.41271 | 0.02191 |
| 354 | LK2016 | TDM_421 | TDM_323 | 0.78171 | 0.02606 |
| 355 | LK2016 | TDM_498 | TDM_323 | 0.55699 | 0.02483 |
| 356 | LK2016 | TDM_54 | TDM_323 | 0.13822 | 0.00907 |
| 357 | LK2016 | TDM_275 | TDM_323 | 0.96573 | 0.00681 |
| 358 | LK2016 | TDM_509 | TDM_323 | 0.85011 | 0.02501 |
| 359 | LK2016 | TDM_04 | TDM_323 | 0.85249 | 0.00867 |
| 360 | LK2016 | TDM_244 | TDM_323 | 0.27580 | 0.02049 |
| 361 | LK2016 | TDM_342 | TDM_323 | 0.36313 | 0.02533 |
| 362 | LK2016 | TDM_389 | TDM_323 | 0.39707 | 0.02444 |
| 363 | LK2016 | TDM_547 | TDM_323 | 0.92601 | 0.00965 |
| 364 | LK2016 | TDM_196 | TDM_323 | 0.99873 | 0.00127 |
| 365 | LK2016 | TDM_360 | TDM_323 | 0.23250 | 0.00856 |
| 366 | LK2016 | TDM_465 | TDM_323 | 0.91078 | 0.02147 |
| 367 | LK2016 | TDM_161 | TDM_323 | 0.57703 | 0.03295 |
| 368 | LK2016 | TDM_200 | TDM_323 | 0.89813 | 0.01302 |
| 369 | LK2016 | TDM_325 | TDM_323 | 0.10483 | 0.01896 |
| 370 | LK2016 | TDM_333 | TDM_323 | 0.64882 | 0.02812 |
| 371 | LK2016 | TDM_122 | TDM_323 | 0.51248 | 0.03106 |
| 372 | LK2016 | TDM_175 | TDM_323 | 0.10702 | 0.01521 |
| 373 | LK2016 | TDM_300 | TDM_323 | 0.79664 | 0.02838 |
| 374 | LK2016 | TDM_1.1 | TDM_323 | 0.05215 | 0.01100 |
| 375 | LK2016 | TDM_450 | TDM_323 | 0.99999 | 0.00001 |
| 376 | LK2016 | TDM_60 | TDM_323 | 0.64734 | 0.03277 |
| 377 | LK2016 | TDM_149 | TDM_323 | 0.11515 | 0.01588 |
| 378 | LK2016 | TDM_224 | TDM_323 | 0.36861 | 0.03164 |
| 379 | LK2016 | TDM_1.2 | TDM_880 | 0.55729 | 0.01236 |
| 380 | LK2016 | TDM_293 | TDM_880 | 0.44564 | 0.00642 |
| 381 | LK2016 | TDM_421 | TDM_880 | 0.74430 | 0.00811 |

| | | | | | |
|---|---|---|---|---|---|
| 382 | LK2016 | TDM_498 | TDM_880 | 0.78067 | 0.00682 |
| 383 | LK2016 | TDM_54 | TDM_880 | 0.20337 | 0.00388 |
| 384 | LK2016 | TDM_275 | TDM_880 | 0.35480 | 0.00809 |
| 385 | LK2016 | TDM_509 | TDM_880 | 0.65500 | 0.01208 |
| 386 | LK2016 | TDM_04 | TDM_880 | 0.51319 | 0.00485 |
| 387 | LK2016 | TDM_244 | TDM_880 | 0.01857 | 0.00258 |
| 388 | LK2016 | TDM_342 | TDM_880 | 0.80830 | 0.00661 |
| 389 | LK2016 | TDM_389 | TDM_880 | 0.83916 | 0.00459 |
| 390 | LK2016 | TDM_547 | TDM_880 | 0.81643 | 0.00501 |
| 391 | LK2016 | TDM_196 | TDM_880 | 0.35242 | 0.01299 |
| 392 | LK2016 | TDM_360 | TDM_880 | 0.42733 | 0.00340 |
| 393 | LK2016 | TDM_465 | TDM_880 | 0.12533 | 0.00858 |
| 394 | LK2016 | TDM_161 | TDM_880 | 0.52767 | 0.00983 |
| 395 | LK2016 | TDM_200 | TDM_880 | 0.51440 | 0.00774 |
| 396 | LK2016 | TDM_325 | TDM_880 | 0.00852 | 0.00165 |
| 397 | LK2016 | TDM_333 | TDM_880 | 0.63623 | 0.00902 |
| 398 | LK2016 | TDM_122 | TDM_880 | 0.56856 | 0.00805 |
| 399 | LK2016 | TDM_175 | TDM_880 | 0.49495 | 0.00784 |
| 400 | LK2016 | TDM_300 | TDM_880 | 0.13317 | 0.00928 |
| 401 | LK2016 | TDM_1.1 | TDM_880 | 0.19559 | 0.00681 |
| 402 | LK2016 | TDM_450 | TDM_880 | 0.69935 | 0.00571 |
| 403 | LK2016 | TDM_60 | TDM_880 | 0.87820 | 0.00604 |
| 404 | LK2016 | TDM_149 | TDM_880 | 0.36653 | 0.00817 |
| 405 | LK2016 | TDM_224 | TDM_880 | 0.94134 | 0.00423 |
| 406 | LK2016 | TDM_323 | TDM_880 | 0.72860 | 0.01331 |
| 407 | LK2014 | TDM_1.2 | TDM_293 | 0.99767 | 0.00121 |
| 408 | LK2014 | TDM_1.2 | TDM_421 | 0.65112 | 0.01671 |
| 409 | LK2014 | TDM_293 | TDM_421 | 0.98453 | 0.00291 |
| 410 | LK2014 | TDM_1.2 | TDM_498 | 0.66058 | 0.01556 |
| 411 | LK2014 | TDM_293 | TDM_498 | 0.53898 | 0.01293 |
| 412 | LK2014 | TDM_421 | TDM_498 | 0.18318 | 0.01039 |
| 413 | LK2014 | TDM_1.2 | TDM_54 | 0.21911 | 0.00755 |
| 414 | LK2014 | TDM_293 | TDM_54 | 0.46787 | 0.00616 |
| 415 | LK2014 | TDM_421 | TDM_54 | 0.76806 | 0.00560 |
| 416 | LK2014 | TDM_498 | TDM_54 | 0.19351 | 0.00580 |
| 417 | LK2014 | TDM_1.2 | TDM_275 | 0.21649 | 0.01657 |
| 418 | LK2014 | TDM_293 | TDM_275 | 0.07972 | 0.00769 |
| 419 | LK2014 | TDM_421 | TDM_275 | 0.23015 | 0.01256 |
| 420 | LK2014 | TDM_498 | TDM_275 | 0.57373 | 0.01180 |
| 421 | LK2014 | TDM_54 | TDM_275 | 0.02948 | 0.00227 |
| 422 | LK2014 | TDM_1.2 | TDM_509 | 0.26517 | 0.02369 |
| 423 | LK2014 | TDM_293 | TDM_509 | 0.59215 | 0.02461 |
| 424 | LK2014 | TDM_421 | TDM_509 | 0.35041 | 0.02297 |
| 425 | LK2014 | TDM_498 | TDM_509 | 0.15872 | 0.01353 |
| 426 | LK2014 | TDM_54 | TDM_509 | 0.47206 | 0.01205 |
| 427 | LK2014 | TDM_275 | TDM_509 | 0.06623 | 0.00871 |
| 428 | LK2014 | TDM_1.2 | TDM_04 | 0.48503 | 0.01020 |
| 429 | LK2014 | TDM_293 | TDM_04 | 0.01493 | 0.00196 |

| | | | | | |
|---|---|---|---|---|---|
| 430 | LK2014 | TDM_421 | TDM_04 | 0.03159 | 0.00285 |
| 431 | LK2014 | TDM_498 | TDM_04 | 0.44384 | 0.00672 |
| 432 | LK2014 | TDM_54 | TDM_04 | 0.81374 | 0.00355 |
| 433 | LK2014 | TDM_275 | TDM_04 | 0.41081 | 0.00672 |
| 434 | LK2014 | TDM_509 | TDM_04 | 0.44482 | 0.01117 |
| 435 | LK2014 | TDM_1.2 | TDM_244 | 0.57804 | 0.01159 |
| 436 | LK2014 | TDM_293 | TDM_244 | 0.72723 | 0.00693 |
| 437 | LK2014 | TDM_421 | TDM_244 | 0.07679 | 0.00414 |
| 438 | LK2014 | TDM_498 | TDM_244 | 0.33315 | 0.00787 |
| 439 | LK2014 | TDM_54 | TDM_244 | 0.54949 | 0.00566 |
| 440 | LK2014 | TDM_275 | TDM_244 | 0.63026 | 0.00722 |
| 441 | LK2014 | TDM_509 | TDM_244 | 0.55583 | 0.01358 |
| 442 | LK2014 | TDM_04 | TDM_244 | 0.37835 | 0.00540 |
| 443 | LK2014 | TDM_1.2 | TDM_342 | 0.43787 | 0.02401 |
| 444 | LK2014 | TDM_293 | TDM_342 | 0.06048 | 0.00744 |
| 445 | LK2014 | TDM_421 | TDM_342 | 0.68058 | 0.01528 |
| 446 | LK2014 | TDM_498 | TDM_342 | 0.49583 | 0.01414 |
| 447 | LK2014 | TDM_54 | TDM_342 | 0.74796 | 0.00705 |
| 448 | LK2014 | TDM_275 | TDM_342 | 0.34368 | 0.01364 |
| 449 | LK2014 | TDM_509 | TDM_342 | 0.72221 | 0.02170 |
| 450 | LK2014 | TDM_04 | TDM_342 | 0.69673 | 0.00695 |
| 451 | LK2014 | TDM_244 | TDM_342 | 0.99061 | 0.00102 |
| 452 | LK2014 | TDM_1.2 | TDM_389 | 0.52185 | 0.01532 |
| 453 | LK2014 | TDM_293 | TDM_389 | 0.15796 | 0.01061 |
| 454 | LK2014 | TDM_421 | TDM_389 | 0.65864 | 0.01019 |
| 455 | LK2014 | TDM_498 | TDM_389 | 0.17675 | 0.00784 |
| 456 | LK2014 | TDM_54 | TDM_389 | 0.24500 | 0.00654 |
| 457 | LK2014 | TDM_275 | TDM_389 | 0.92766 | 0.00457 |
| 458 | LK2014 | TDM_509 | TDM_389 | 0.57808 | 0.01648 |
| 459 | LK2014 | TDM_04 | TDM_389 | 0.21466 | 0.00607 |
| 460 | LK2014 | TDM_244 | TDM_389 | 0.01540 | 0.00156 |
| 461 | LK2014 | TDM_342 | TDM_389 | 0.69597 | 0.01301 |
| 462 | LK2014 | TDM_1.2 | TDM_547 | 0.56781 | 0.01610 |
| 463 | LK2014 | TDM_293 | TDM_547 | 0.56593 | 0.01318 |
| 464 | LK2014 | TDM_421 | TDM_547 | 0.74612 | 0.00923 |
| 465 | LK2014 | TDM_498 | TDM_547 | 0.02693 | 0.00297 |
| 466 | LK2014 | TDM_54 | TDM_547 | 0.97633 | 0.00114 |
| 467 | LK2014 | TDM_275 | TDM_547 | 0.30609 | 0.00910 |
| 468 | LK2014 | TDM_509 | TDM_547 | 0.15477 | 0.01246 |
| 469 | LK2014 | TDM_04 | TDM_547 | 0.15459 | 0.00479 |
| 470 | LK2014 | TDM_244 | TDM_547 | 0.53392 | 0.00835 |
| 471 | LK2014 | TDM_342 | TDM_547 | 0.32150 | 0.01446 |
| 472 | LK2014 | TDM_389 | TDM_547 | 0.23212 | 0.00900 |
| 473 | LK2014 | TDM_1.2 | TDM_196 | 0.80898 | 0.02178 |
| 474 | LK2014 | TDM_293 | TDM_196 | 0.54109 | 0.02709 |
| 475 | LK2014 | TDM_421 | TDM_196 | 0.80794 | 0.01772 |
| 476 | LK2014 | TDM_498 | TDM_196 | 0.22745 | 0.01985 |
| 477 | LK2014 | TDM_54 | TDM_196 | 0.87368 | 0.00797 |

| | | | | | |
|---|---|---|---|---|---|
| 478 | LK2014 | TDM_275 | TDM_196 | 0.74397 | 0.01985 |
| 479 | LK2014 | TDM_509 | TDM_196 | 0.50850 | 0.03520 |
| 480 | LK2014 | TDM_04 | TDM_196 | 0.57548 | 0.01189 |
| 481 | LK2014 | TDM_244 | TDM_196 | 0.51504 | 0.01506 |
| 482 | LK2014 | TDM_342 | TDM_196 | 0.23029 | 0.02271 |
| 483 | LK2014 | TDM_389 | TDM_196 | 0.17884 | 0.01585 |
| 484 | LK2014 | TDM_547 | TDM_196 | 0.89171 | 0.01028 |
| 485 | LK2014 | TDM_1.2 | TDM_360 | 0.59623 | 0.01600 |
| 486 | LK2014 | TDM_293 | TDM_360 | 0.13997 | 0.01010 |
| 487 | LK2014 | TDM_421 | TDM_360 | 0.59091 | 0.01203 |
| 488 | LK2014 | TDM_498 | TDM_360 | 0.27679 | 0.01101 |
| 489 | LK2014 | TDM_54 | TDM_360 | 0.72738 | 0.00578 |
| 490 | LK2014 | TDM_275 | TDM_360 | 0.73875 | 0.01114 |
| 491 | LK2014 | TDM_509 | TDM_360 | 0.15249 | 0.01217 |
| 492 | LK2014 | TDM_04 | TDM_360 | 0.29454 | 0.00709 |
| 493 | LK2014 | TDM_244 | TDM_360 | 0.59988 | 0.00746 |
| 494 | LK2014 | TDM_342 | TDM_360 | 0.87984 | 0.00761 |
| 495 | LK2014 | TDM_389 | TDM_360 | 0.61298 | 0.00989 |
| 496 | LK2014 | TDM_547 | TDM_360 | 0.66806 | 0.00855 |
| 497 | LK2014 | TDM_196 | TDM_360 | 0.77489 | 0.01657 |
| 498 | LK2014 | TDM_1.2 | TDM_465 | 0.88211 | 0.01817 |
| 499 | LK2014 | TDM_293 | TDM_465 | 0.06545 | 0.01265 |
| 500 | LK2014 | TDM_421 | TDM_465 | 0.66723 | 0.02143 |
| 501 | LK2014 | TDM_498 | TDM_465 | 0.64117 | 0.02037 |
| 502 | LK2014 | TDM_54 | TDM_465 | 0.38113 | 0.01313 |
| 503 | LK2014 | TDM_275 | TDM_465 | 0.38766 | 0.02407 |
| 504 | LK2014 | TDM_509 | TDM_465 | 0.96096 | 0.00934 |
| 505 | LK2014 | TDM_04 | TDM_465 | 0.16315 | 0.01090 |
| 506 | LK2014 | TDM_244 | TDM_465 | 0.50411 | 0.01499 |
| 507 | LK2014 | TDM_342 | TDM_465 | 0.26369 | 0.02411 |
| 508 | LK2014 | TDM_389 | TDM_465 | 0.52998 | 0.02219 |
| 509 | LK2014 | TDM_547 | TDM_465 | 0.43835 | 0.02061 |
| 510 | LK2014 | TDM_196 | TDM_465 | 1.00000 | 0.00000 |
| 511 | LK2014 | TDM_360 | TDM_465 | 0.49488 | 0.02073 |
| 512 | LK2014 | TDM_1.2 | TDM_161 | 0.65822 | 0.02191 |
| 513 | LK2014 | TDM_293 | TDM_161 | 0.02469 | 0.00485 |
| 514 | LK2014 | TDM_421 | TDM_161 | 0.22026 | 0.01266 |
| 515 | LK2014 | TDM_498 | TDM_161 | 0.19833 | 0.01371 |
| 516 | LK2014 | TDM_54 | TDM_161 | 0.59462 | 0.00991 |
| 517 | LK2014 | TDM_275 | TDM_161 | 0.94515 | 0.00604 |
| 518 | LK2014 | TDM_509 | TDM_161 | 0.56432 | 0.02553 |
| 519 | LK2014 | TDM_04 | TDM_161 | 0.31377 | 0.00912 |
| 520 | LK2014 | TDM_244 | TDM_161 | 0.89016 | 0.00526 |
| 521 | LK2014 | TDM_342 | TDM_161 | 0.40421 | 0.02032 |
| 522 | LK2014 | TDM_389 | TDM_161 | 0.64835 | 0.01373 |
| 523 | LK2014 | TDM_547 | TDM_161 | 0.68608 | 0.01371 |
| 524 | LK2014 | TDM_196 | TDM_161 | 0.70258 | 0.02643 |
| 525 | LK2014 | TDM_360 | TDM_161 | 0.10472 | 0.00895 |

| | | | | | |
|---|---|---|---|---|---|
| 526 | LK2014 | TDM_465 | TDM_161 | 0.23772 | 0.02386 |
| 527 | LK2014 | TDM_1.2 | TDM_200 | 0.01027 | 0.00326 |
| 528 | LK2014 | TDM_293 | TDM_200 | 0.30931 | 0.01789 |
| 529 | LK2014 | TDM_421 | TDM_200 | 0.48045 | 0.01780 |
| 530 | LK2014 | TDM_498 | TDM_200 | 0.07836 | 0.00816 |
| 531 | LK2014 | TDM_54 | TDM_200 | 0.65916 | 0.00924 |
| 532 | LK2014 | TDM_275 | TDM_200 | 0.00055 | 0.00030 |
| 533 | LK2014 | TDM_509 | TDM_200 | 0.63917 | 0.02476 |
| 534 | LK2014 | TDM_04 | TDM_200 | 0.44658 | 0.01028 |
| 535 | LK2014 | TDM_244 | TDM_200 | 0.25245 | 0.00875 |
| 536 | LK2014 | TDM_342 | TDM_200 | 0.57301 | 0.01773 |
| 537 | LK2014 | TDM_389 | TDM_200 | 0.19127 | 0.01318 |
| 538 | LK2014 | TDM_547 | TDM_200 | 0.19305 | 0.01132 |
| 539 | LK2014 | TDM_196 | TDM_200 | 0.00941 | 0.00358 |
| 540 | LK2014 | TDM_360 | TDM_200 | 0.33607 | 0.01284 |
| 541 | LK2014 | TDM_465 | TDM_200 | 0.21570 | 0.02639 |
| 542 | LK2014 | TDM_161 | TDM_200 | 0.02384 | 0.00609 |
| 543 | LK2014 | TDM_1.2 | TDM_325 | 0.09868 | 0.01009 |
| 544 | LK2014 | TDM_293 | TDM_325 | 0.25228 | 0.01191 |
| 545 | LK2014 | TDM_421 | TDM_325 | 0.28282 | 0.01457 |
| 546 | LK2014 | TDM_498 | TDM_325 | 0.02922 | 0.00416 |
| 547 | LK2014 | TDM_54 | TDM_325 | 0.09348 | 0.00522 |
| 548 | LK2014 | TDM_275 | TDM_325 | 0.60289 | 0.01179 |
| 549 | LK2014 | TDM_509 | TDM_325 | 0.24119 | 0.01981 |
| 550 | LK2014 | TDM_04 | TDM_325 | 0.58223 | 0.00753 |
| 551 | LK2014 | TDM_244 | TDM_325 | 0.10948 | 0.00638 |
| 552 | LK2014 | TDM_342 | TDM_325 | 0.33666 | 0.01596 |
| 553 | LK2014 | TDM_389 | TDM_325 | 0.60123 | 0.01212 |
| 554 | LK2014 | TDM_547 | TDM_325 | 0.46795 | 0.01204 |
| 555 | LK2014 | TDM_196 | TDM_325 | 0.31982 | 0.02422 |
| 556 | LK2014 | TDM_360 | TDM_325 | 0.23307 | 0.01149 |
| 557 | LK2014 | TDM_465 | TDM_325 | 0.04547 | 0.00836 |
| 558 | LK2014 | TDM_161 | TDM_325 | 0.49984 | 0.02029 |
| 559 | LK2014 | TDM_200 | TDM_325 | 0.32321 | 0.01831 |
| 560 | LK2014 | TDM_1.2 | TDM_333 | 0.67162 | 0.02227 |
| 561 | LK2014 | TDM_293 | TDM_333 | 0.08946 | 0.01248 |
| 562 | LK2014 | TDM_421 | TDM_333 | 0.14737 | 0.01417 |
| 563 | LK2014 | TDM_498 | TDM_333 | 0.02780 | 0.00443 |
| 564 | LK2014 | TDM_54 | TDM_333 | 0.64555 | 0.00947 |
| 565 | LK2014 | TDM_275 | TDM_333 | 0.04225 | 0.00646 |
| 566 | LK2014 | TDM_509 | TDM_333 | 0.32057 | 0.03084 |
| 567 | LK2014 | TDM_04 | TDM_333 | 0.03648 | 0.00379 |
| 568 | LK2014 | TDM_244 | TDM_333 | 0.26220 | 0.01051 |
| 569 | LK2014 | TDM_342 | TDM_333 | 0.51661 | 0.02385 |
| 570 | LK2014 | TDM_389 | TDM_333 | 0.16496 | 0.01248 |
| 571 | LK2014 | TDM_547 | TDM_333 | 0.88834 | 0.00721 |
| 572 | LK2014 | TDM_196 | TDM_333 | 0.62358 | 0.03264 |
| 573 | LK2014 | TDM_360 | TDM_333 | 0.09980 | 0.00898 |

| | | | | | |
|---|---|---|---|---|---|
| 574 | LK2014 | TDM_465 | TDM_333 | 0.31856 | 0.03326 |
| 575 | LK2014 | TDM_161 | TDM_333 | 0.14668 | 0.01619 |
| 576 | LK2014 | TDM_200 | TDM_333 | 0.48997 | 0.02308 |
| 577 | LK2014 | TDM_325 | TDM_333 | 0.08221 | 0.01059 |
| 578 | LK2014 | TDM_1.2 | TDM_122 | 0.26852 | 0.01370 |
| 579 | LK2014 | TDM_293 | TDM_122 | 0.08521 | 0.00784 |
| 580 | LK2014 | TDM_421 | TDM_122 | 0.15178 | 0.00843 |
| 581 | LK2014 | TDM_498 | TDM_122 | 0.82545 | 0.00840 |
| 582 | LK2014 | TDM_54 | TDM_122 | 0.41061 | 0.00680 |
| 583 | LK2014 | TDM_275 | TDM_122 | 0.00000 | 0.00000 |
| 584 | LK2014 | TDM_509 | TDM_122 | 0.11508 | 0.01168 |
| 585 | LK2014 | TDM_04 | TDM_122 | 0.84494 | 0.00401 |
| 586 | LK2014 | TDM_244 | TDM_122 | 0.94772 | 0.00242 |
| 587 | LK2014 | TDM_342 | TDM_122 | 0.58690 | 0.01335 |
| 588 | LK2014 | TDM_389 | TDM_122 | 0.06517 | 0.00478 |
| 589 | LK2014 | TDM_547 | TDM_122 | 0.15535 | 0.00805 |
| 590 | LK2014 | TDM_196 | TDM_122 | 0.14278 | 0.01450 |
| 591 | LK2014 | TDM_360 | TDM_122 | 0.21964 | 0.00906 |
| 592 | LK2014 | TDM_465 | TDM_122 | 0.46366 | 0.02196 |
| 593 | LK2014 | TDM_161 | TDM_122 | 0.24745 | 0.01388 |
| 594 | LK2014 | TDM_200 | TDM_122 | 0.00722 | 0.00143 |
| 595 | LK2014 | TDM_325 | TDM_122 | 0.51852 | 0.01259 |
| 596 | LK2014 | TDM_333 | TDM_122 | 0.49053 | 0.01666 |
| 597 | LK2014 | TDM_1.2 | TDM_175 | 0.00594 | 0.00159 |
| 598 | LK2014 | TDM_293 | TDM_175 | 0.74933 | 0.00951 |
| 599 | LK2014 | TDM_421 | TDM_175 | 0.18868 | 0.00835 |
| 600 | LK2014 | TDM_498 | TDM_175 | 0.70339 | 0.00890 |
| 601 | LK2014 | TDM_54 | TDM_175 | 0.24092 | 0.00529 |
| 602 | LK2014 | TDM_275 | TDM_175 | 0.38388 | 0.00958 |
| 603 | LK2014 | TDM_509 | TDM_175 | 0.42169 | 0.01443 |
| 604 | LK2014 | TDM_04 | TDM_175 | 0.20349 | 0.00560 |
| 605 | LK2014 | TDM_244 | TDM_175 | 0.45772 | 0.00667 |
| 606 | LK2014 | TDM_342 | TDM_175 | 0.30795 | 0.01123 |
| 607 | LK2014 | TDM_389 | TDM_175 | 0.02691 | 0.00244 |
| 608 | LK2014 | TDM_547 | TDM_175 | 0.47183 | 0.00981 |
| 609 | LK2014 | TDM_196 | TDM_175 | 0.73070 | 0.01776 |
| 610 | LK2014 | TDM_360 | TDM_175 | 0.73403 | 0.00813 |
| 611 | LK2014 | TDM_465 | TDM_175 | 0.46243 | 0.01729 |
| 612 | LK2014 | TDM_161 | TDM_175 | 0.75059 | 0.00911 |
| 613 | LK2014 | TDM_200 | TDM_175 | 0.58597 | 0.01279 |
| 614 | LK2014 | TDM_325 | TDM_175 | 0.04346 | 0.00407 |
| 615 | LK2014 | TDM_333 | TDM_175 | 0.38177 | 0.01484 |
| 616 | LK2014 | TDM_122 | TDM_175 | 0.80697 | 0.00606 |
| 617 | LK2014 | TDM_1.2 | TDM_300 | 0.19226 | 0.02463 |
| 618 | LK2014 | TDM_293 | TDM_300 | 0.82089 | 0.02021 |
| 619 | LK2014 | TDM_421 | TDM_300 | 0.97370 | 0.00553 |
| 620 | LK2014 | TDM_498 | TDM_300 | 0.42600 | 0.02393 |
| 621 | LK2014 | TDM_54 | TDM_300 | 0.93339 | 0.00489 |

| | | | | | |
|---|---|---|---|---|---|
| 622 | LK2014 | TDM_275 | TDM_300 | 0.40734 | 0.02695 |
| 623 | LK2014 | TDM_509 | TDM_300 | 0.19547 | 0.02635 |
| 624 | LK2014 | TDM_04 | TDM_300 | 0.45967 | 0.01276 |
| 625 | LK2014 | TDM_244 | TDM_300 | 0.34888 | 0.01578 |
| 626 | LK2014 | TDM_342 | TDM_300 | 0.38910 | 0.02813 |
| 627 | LK2014 | TDM_389 | TDM_300 | 0.05484 | 0.00846 |
| 628 | LK2014 | TDM_547 | TDM_300 | 0.04745 | 0.01014 |
| 629 | LK2014 | TDM_196 | TDM_300 | 0.40748 | 0.03996 |
| 630 | LK2014 | TDM_360 | TDM_300 | 0.10897 | 0.01279 |
| 631 | LK2014 | TDM_465 | TDM_300 | 0.12223 | 0.02281 |
| 632 | LK2014 | TDM_161 | TDM_300 | 0.46441 | 0.02919 |
| 633 | LK2014 | TDM_200 | TDM_300 | 0.88666 | 0.01580 |
| 634 | LK2014 | TDM_325 | TDM_300 | 0.30985 | 0.02468 |
| 635 | LK2014 | TDM_333 | TDM_300 | 0.39643 | 0.03244 |
| 636 | LK2014 | TDM_122 | TDM_300 | 0.00981 | 0.00292 |
| 637 | LK2014 | TDM_175 | TDM_300 | 0.71323 | 0.01861 |
| 638 | LK2014 | TDM_1.2 | TDM_1.1 | 0.00000 | 0.00000 |
| 639 | LK2014 | TDM_293 | TDM_1.1 | 0.43243 | 0.01296 |
| 640 | LK2014 | TDM_421 | TDM_1.1 | 0.22075 | 0.01061 |
| 641 | LK2014 | TDM_498 | TDM_1.1 | 0.24939 | 0.00874 |
| 642 | LK2014 | TDM_54 | TDM_1.1 | 0.41365 | 0.00749 |
| 643 | LK2014 | TDM_275 | TDM_1.1 | 0.04018 | 0.00351 |
| 644 | LK2014 | TDM_509 | TDM_1.1 | 0.98558 | 0.00321 |
| 645 | LK2014 | TDM_04 | TDM_1.1 | 0.68170 | 0.00718 |
| 646 | LK2014 | TDM_244 | TDM_1.1 | 0.49321 | 0.00901 |
| 647 | LK2014 | TDM_342 | TDM_1.1 | 0.40970 | 0.01319 |
| 648 | LK2014 | TDM_389 | TDM_1.1 | 0.89290 | 0.00511 |
| 649 | LK2014 | TDM_547 | TDM_1.1 | 0.64183 | 0.00992 |
| 650 | LK2014 | TDM_196 | TDM_1.1 | 0.05263 | 0.01007 |
| 651 | LK2014 | TDM_360 | TDM_1.1 | 0.68161 | 0.00943 |
| 652 | LK2014 | TDM_465 | TDM_1.1 | 0.49318 | 0.02243 |
| 653 | LK2014 | TDM_161 | TDM_1.1 | 0.58553 | 0.01415 |
| 654 | LK2014 | TDM_200 | TDM_1.1 | 0.00111 | 0.00066 |
| 655 | LK2014 | TDM_325 | TDM_1.1 | 0.43789 | 0.01291 |
| 656 | LK2014 | TDM_333 | TDM_1.1 | 0.99151 | 0.00188 |
| 657 | LK2014 | TDM_122 | TDM_1.1 | 0.10144 | 0.00683 |
| 658 | LK2014 | TDM_175 | TDM_1.1 | 0.14586 | 0.00687 |
| 659 | LK2014 | TDM_300 | TDM_1.1 | 0.64671 | 0.02289 |
| 660 | LK2014 | TDM_1.2 | TDM_450 | 0.68674 | 0.01816 |
| 661 | LK2014 | TDM_293 | TDM_450 | 0.03498 | 0.00756 |
| 662 | LK2014 | TDM_421 | TDM_450 | 0.24343 | 0.01398 |
| 663 | LK2014 | TDM_498 | TDM_450 | 0.32316 | 0.01264 |
| 664 | LK2014 | TDM_54 | TDM_450 | 0.61931 | 0.00781 |
| 665 | LK2014 | TDM_275 | TDM_450 | 0.67545 | 0.01569 |
| 666 | LK2014 | TDM_509 | TDM_450 | 0.52973 | 0.02418 |
| 667 | LK2014 | TDM_04 | TDM_450 | 0.70161 | 0.00672 |
| 668 | LK2014 | TDM_244 | TDM_450 | 0.05202 | 0.00395 |
| 669 | LK2014 | TDM_342 | TDM_450 | 0.00119 | 0.00050 |

| | | | | | |
|---|---|---|---|---|---|
| 670 | LK2014 | TDM_389 | TDM_450 | 0.68415 | 0.01297 |
| 671 | LK2014 | TDM_547 | TDM_450 | 0.44980 | 0.01313 |
| 672 | LK2014 | TDM_196 | TDM_450 | 0.77539 | 0.01975 |
| 673 | LK2014 | TDM_360 | TDM_450 | 0.18414 | 0.00985 |
| 674 | LK2014 | TDM_465 | TDM_450 | 0.57250 | 0.02565 |
| 675 | LK2014 | TDM_161 | TDM_450 | 0.08068 | 0.00990 |
| 676 | LK2014 | TDM_200 | TDM_450 | 0.05410 | 0.00880 |
| 677 | LK2014 | TDM_325 | TDM_450 | 0.80188 | 0.01145 |
| 678 | LK2014 | TDM_333 | TDM_450 | 0.36445 | 0.02238 |
| 679 | LK2014 | TDM_122 | TDM_450 | 0.14636 | 0.01156 |
| 680 | LK2014 | TDM_175 | TDM_450 | 0.96995 | 0.00279 |
| 681 | LK2014 | TDM_300 | TDM_450 | 0.20269 | 0.02134 |
| 682 | LK2014 | TDM_1.1 | TDM_450 | 0.36220 | 0.01485 |
| 683 | LK2014 | TDM_1.2 | TDM_60 | 0.22042 | 0.02067 |
| 684 | LK2014 | TDM_293 | TDM_60 | 0.48670 | 0.02419 |
| 685 | LK2014 | TDM_421 | TDM_60 | 0.07486 | 0.00953 |
| 686 | LK2014 | TDM_498 | TDM_60 | 0.40726 | 0.01803 |
| 687 | LK2014 | TDM_54 | TDM_60 | 0.10370 | 0.00651 |
| 688 | LK2014 | TDM_275 | TDM_60 | 0.21178 | 0.01601 |
| 689 | LK2014 | TDM_509 | TDM_60 | 0.44470 | 0.02926 |
| 690 | LK2014 | TDM_04 | TDM_60 | 0.81277 | 0.00637 |
| 691 | LK2014 | TDM_244 | TDM_60 | 0.77633 | 0.00829 |
| 692 | LK2014 | TDM_342 | TDM_60 | 0.72698 | 0.01804 |
| 693 | LK2014 | TDM_389 | TDM_60 | 0.25351 | 0.01423 |
| 694 | LK2014 | TDM_547 | TDM_60 | 0.34055 | 0.01309 |
| 695 | LK2014 | TDM_196 | TDM_60 | 0.31861 | 0.02886 |
| 696 | LK2014 | TDM_360 | TDM_60 | 0.07615 | 0.00792 |
| 697 | LK2014 | TDM_465 | TDM_60 | 0.62746 | 0.03050 |
| 698 | LK2014 | TDM_161 | TDM_60 | 0.05639 | 0.01149 |
| 699 | LK2014 | TDM_200 | TDM_60 | 0.02961 | 0.00802 |
| 700 | LK2014 | TDM_325 | TDM_60 | 0.10756 | 0.00988 |
| 701 | LK2014 | TDM_333 | TDM_60 | 0.90145 | 0.01244 |
| 702 | LK2014 | TDM_122 | TDM_60 | 0.00654 | 0.00181 |
| 703 | LK2014 | TDM_175 | TDM_60 | 0.10778 | 0.00821 |
| 704 | LK2014 | TDM_300 | TDM_60 | 0.06683 | 0.01450 |
| 705 | LK2014 | TDM_1.1 | TDM_60 | 0.00864 | 0.00247 |
| 706 | LK2014 | TDM_450 | TDM_60 | 0.65820 | 0.01846 |
| 707 | LK2014 | TDM_1.2 | TDM_149 | 0.00760 | 0.00196 |
| 708 | LK2014 | TDM_293 | TDM_149 | 0.60413 | 0.01468 |
| 709 | LK2014 | TDM_421 | TDM_149 | 0.04837 | 0.00483 |
| 710 | LK2014 | TDM_498 | TDM_149 | 0.42111 | 0.01266 |
| 711 | LK2014 | TDM_54 | TDM_149 | 0.11102 | 0.00438 |
| 712 | LK2014 | TDM_275 | TDM_149 | 0.23570 | 0.00960 |
| 713 | LK2014 | TDM_509 | TDM_149 | 0.19546 | 0.01765 |
| 714 | LK2014 | TDM_04 | TDM_149 | 0.36638 | 0.00794 |
| 715 | LK2014 | TDM_244 | TDM_149 | 0.65833 | 0.00749 |
| 716 | LK2014 | TDM_342 | TDM_149 | 0.99587 | 0.00104 |
| 717 | LK2014 | TDM_389 | TDM_149 | 0.83613 | 0.00723 |

| | | | | | |
|---|---|---|---|---|---|
| 718 | LK2014 | TDM_547 | TDM_149 | 0.57141 | 0.01053 |
| 719 | LK2014 | TDM_196 | TDM_149 | 0.92375 | 0.01145 |
| 720 | LK2014 | TDM_360 | TDM_149 | 0.66030 | 0.01253 |
| 721 | LK2014 | TDM_465 | TDM_149 | 0.06509 | 0.01229 |
| 722 | LK2014 | TDM_161 | TDM_149 | 0.10824 | 0.01011 |
| 723 | LK2014 | TDM_200 | TDM_149 | 0.63012 | 0.01604 |
| 724 | LK2014 | TDM_325 | TDM_149 | 0.00217 | 0.00088 |
| 725 | LK2014 | TDM_333 | TDM_149 | 0.38425 | 0.02119 |
| 726 | LK2014 | TDM_122 | TDM_149 | 0.11333 | 0.00755 |
| 727 | LK2014 | TDM_175 | TDM_149 | 0.06286 | 0.00422 |
| 728 | LK2014 | TDM_300 | TDM_149 | 0.66287 | 0.02330 |
| 729 | LK2014 | TDM_1.1 | TDM_149 | 0.40048 | 0.01082 |
| 730 | LK2014 | TDM_450 | TDM_149 | 0.34161 | 0.01528 |
| 731 | LK2014 | TDM_60 | TDM_149 | 0.72883 | 0.01547 |
| 732 | LK2014 | TDM_1.2 | TDM_224 | 0.52850 | 0.02692 |
| 733 | LK2014 | TDM_293 | TDM_224 | 0.15364 | 0.01614 |
| 734 | LK2014 | TDM_421 | TDM_224 | 0.57916 | 0.01936 |
| 735 | LK2014 | TDM_498 | TDM_224 | 0.05557 | 0.00834 |
| 736 | LK2014 | TDM_54 | TDM_224 | 0.05523 | 0.00459 |
| 737 | LK2014 | TDM_275 | TDM_224 | 0.37372 | 0.01987 |
| 738 | LK2014 | TDM_509 | TDM_224 | 0.07186 | 0.01652 |
| 739 | LK2014 | TDM_04 | TDM_224 | 0.18059 | 0.00882 |
| 740 | LK2014 | TDM_244 | TDM_224 | 0.85976 | 0.00657 |
| 741 | LK2014 | TDM_342 | TDM_224 | 0.39095 | 0.02106 |
| 742 | LK2014 | TDM_389 | TDM_224 | 0.23404 | 0.01448 |
| 743 | LK2014 | TDM_547 | TDM_224 | 0.41478 | 0.01769 |
| 744 | LK2014 | TDM_196 | TDM_224 | 0.39697 | 0.03439 |
| 745 | LK2014 | TDM_360 | TDM_224 | 0.11420 | 0.01142 |
| 746 | LK2014 | TDM_465 | TDM_224 | 0.69368 | 0.02718 |
| 747 | LK2014 | TDM_161 | TDM_224 | 0.22886 | 0.02017 |
| 748 | LK2014 | TDM_200 | TDM_224 | 0.54435 | 0.02411 |
| 749 | LK2014 | TDM_325 | TDM_224 | 0.00056 | 0.00056 |
| 750 | LK2014 | TDM_333 | TDM_224 | 0.27148 | 0.02367 |
| 751 | LK2014 | TDM_122 | TDM_224 | 0.51679 | 0.01826 |
| 752 | LK2014 | TDM_175 | TDM_224 | 0.43267 | 0.01568 |
| 753 | LK2014 | TDM_300 | TDM_224 | 0.60340 | 0.03016 |
| 754 | LK2014 | TDM_1.1 | TDM_224 | 0.78837 | 0.01464 |
| 755 | LK2014 | TDM_450 | TDM_224 | 0.34883 | 0.01986 |
| 756 | LK2014 | TDM_60 | TDM_224 | 0.18081 | 0.01858 |
| 757 | LK2014 | TDM_149 | TDM_224 | 0.26214 | 0.01556 |
| 758 | LK2014 | TDM_1.2 | TDM_323 | 0.64056 | 0.03129 |
| 759 | LK2014 | TDM_293 | TDM_323 | 0.47726 | 0.02516 |
| 760 | LK2014 | TDM_421 | TDM_323 | 0.62059 | 0.02085 |
| 761 | LK2014 | TDM_498 | TDM_323 | 0.34515 | 0.01896 |
| 762 | LK2014 | TDM_54 | TDM_323 | 0.37159 | 0.01339 |
| 763 | LK2014 | TDM_275 | TDM_323 | 0.46080 | 0.02463 |
| 764 | LK2014 | TDM_509 | TDM_323 | 0.32382 | 0.03363 |
| 765 | LK2014 | TDM_04 | TDM_323 | 0.92776 | 0.00558 |

| | | | | | |
|---|---|---|---|---|---|
| 766 | LK2014 | TDM_244 | TDM_323 | 0.16923 | 0.01038 |
| 767 | LK2014 | TDM_342 | TDM_323 | 0.47988 | 0.02540 |
| 768 | LK2014 | TDM_389 | TDM_323 | 0.53648 | 0.02122 |
| 769 | LK2014 | TDM_547 | TDM_323 | 0.36105 | 0.01982 |
| 770 | LK2014 | TDM_196 | TDM_323 | 0.06883 | 0.01809 |
| 771 | LK2014 | TDM_360 | TDM_323 | 0.54143 | 0.02208 |
| 772 | LK2014 | TDM_465 | TDM_323 | 0.93535 | 0.01926 |
| 773 | LK2014 | TDM_161 | TDM_323 | 0.52730 | 0.02674 |
| 774 | LK2014 | TDM_200 | TDM_323 | 0.97311 | 0.00795 |
| 775 | LK2014 | TDM_325 | TDM_323 | 0.26409 | 0.02260 |
| 776 | LK2014 | TDM_333 | TDM_323 | 0.72854 | 0.02761 |
| 777 | LK2014 | TDM_122 | TDM_323 | 0.54160 | 0.02096 |
| 778 | LK2014 | TDM_175 | TDM_323 | 0.34782 | 0.01632 |
| 779 | LK2014 | TDM_300 | TDM_323 | 0.90183 | 0.02071 |
| 780 | LK2014 | TDM_1.1 | TDM_323 | 0.61109 | 0.02158 |
| 781 | LK2014 | TDM_450 | TDM_323 | 0.37634 | 0.02754 |
| 782 | LK2014 | TDM_60 | TDM_323 | 0.36410 | 0.02831 |
| 783 | LK2014 | TDM_149 | TDM_323 | 0.87369 | 0.01241 |
| 784 | LK2014 | TDM_224 | TDM_323 | 0.51633 | 0.03535 |
| 785 | LK2014 | TDM_1.2 | TDM_880 | 0.08140 | 0.00926 |
| 786 | LK2014 | TDM_293 | TDM_880 | 0.96276 | 0.00489 |
| 787 | LK2014 | TDM_421 | TDM_880 | 0.00611 | 0.00163 |
| 788 | LK2014 | TDM_498 | TDM_880 | 0.53478 | 0.01261 |
| 789 | LK2014 | TDM_54 | TDM_880 | 0.56021 | 0.00922 |
| 790 | LK2014 | TDM_275 | TDM_880 | 0.43402 | 0.01235 |
| 791 | LK2014 | TDM_509 | TDM_880 | 0.38481 | 0.02054 |
| 792 | LK2014 | TDM_04 | TDM_880 | 0.07338 | 0.00518 |
| 793 | LK2014 | TDM_244 | TDM_880 | 0.13801 | 0.00534 |
| 794 | LK2014 | TDM_342 | TDM_880 | 0.45063 | 0.01724 |
| 795 | LK2014 | TDM_389 | TDM_880 | 0.37836 | 0.01158 |
| 796 | LK2014 | TDM_547 | TDM_880 | 0.84317 | 0.00717 |
| 797 | LK2014 | TDM_196 | TDM_880 | 0.95428 | 0.00821 |
| 798 | LK2014 | TDM_360 | TDM_880 | 0.44602 | 0.01249 |
| 799 | LK2014 | TDM_465 | TDM_880 | 0.41318 | 0.02458 |
| 800 | LK2014 | TDM_161 | TDM_880 | 0.32021 | 0.01698 |
| 801 | LK2014 | TDM_200 | TDM_880 | 0.82389 | 0.01281 |
| 802 | LK2014 | TDM_325 | TDM_880 | 0.66318 | 0.01300 |
| 803 | LK2014 | TDM_333 | TDM_880 | 0.59622 | 0.01983 |
| 804 | LK2014 | TDM_122 | TDM_880 | 0.82333 | 0.00937 |
| 805 | LK2014 | TDM_175 | TDM_880 | 0.71667 | 0.00920 |
| 806 | LK2014 | TDM_300 | TDM_880 | 0.92595 | 0.01101 |
| 807 | LK2014 | TDM_1.1 | TDM_880 | 0.83409 | 0.00801 |
| 808 | LK2014 | TDM_450 | TDM_880 | 0.27487 | 0.01467 |
| 809 | LK2014 | TDM_60 | TDM_880 | 0.27404 | 0.01624 |
| 810 | LK2014 | TDM_149 | TDM_880 | 0.56538 | 0.01259 |
| 811 | LK2014 | TDM_224 | TDM_880 | 0.40678 | 0.02070 |
| 812 | LK2014 | TDM_323 | TDM_880 | 0.77250 | 0.01917 |
| 813 | UK2016 | TDM_1.2 | TDM_293 | 0.72915 | 0.01992 |

| | | | | | |
|---|---|---|---|---|---|
| 814 | UK2016 | TDM_1.2 | TDM_421 | 0.67379 | 0.02697 |
| 815 | UK2016 | TDM_293 | TDM_421 | 0.81609 | 0.02066 |
| 816 | UK2016 | TDM_1.2 | TDM_498 | 0.59096 | 0.02715 |
| 817 | UK2016 | TDM_293 | TDM_498 | 0.75872 | 0.01997 |
| 818 | UK2016 | TDM_421 | TDM_498 | 0.78111 | 0.02687 |
| 819 | UK2016 | TDM_1.2 | TDM_54 | 0.63066 | 0.02358 |
| 820 | UK2016 | TDM_293 | TDM_54 | 0.89285 | 0.01151 |
| 821 | UK2016 | TDM_421 | TDM_54 | 0.24042 | 0.02380 |
| 822 | UK2016 | TDM_498 | TDM_54 | 0.31405 | 0.02568 |
| 823 | UK2016 | TDM_1.2 | TDM_275 | 0.04851 | 0.00705 |
| 824 | UK2016 | TDM_293 | TDM_275 | 0.16518 | 0.01205 |
| 825 | UK2016 | TDM_421 | TDM_275 | 0.30365 | 0.01776 |
| 826 | UK2016 | TDM_498 | TDM_275 | 0.11457 | 0.01171 |
| 827 | UK2016 | TDM_54 | TDM_275 | 0.36679 | 0.01495 |
| 828 | UK2016 | TDM_1.2 | TDM_509 | 0.05168 | 0.01138 |
| 829 | UK2016 | TDM_293 | TDM_509 | 0.95783 | 0.00879 |
| 830 | UK2016 | TDM_421 | TDM_509 | 0.39367 | 0.03385 |
| 831 | UK2016 | TDM_498 | TDM_509 | 0.14202 | 0.02576 |
| 832 | UK2016 | TDM_54 | TDM_509 | 0.58143 | 0.02828 |
| 833 | UK2016 | TDM_275 | TDM_509 | 0.84633 | 0.01237 |
| 834 | UK2016 | TDM_1.2 | TDM_04 | 0.33283 | 0.02230 |
| 835 | UK2016 | TDM_293 | TDM_04 | 0.07229 | 0.00823 |
| 836 | UK2016 | TDM_421 | TDM_04 | 0.58938 | 0.02330 |
| 837 | UK2016 | TDM_498 | TDM_04 | 0.74979 | 0.01723 |
| 838 | UK2016 | TDM_54 | TDM_04 | 0.31488 | 0.02023 |
| 839 | UK2016 | TDM_275 | TDM_04 | 0.03118 | 0.00460 |
| 840 | UK2016 | TDM_509 | TDM_04 | 0.67931 | 0.01901 |
| 841 | UK2016 | TDM_1.2 | TDM_244 | 0.07136 | 0.01196 |
| 842 | UK2016 | TDM_293 | TDM_244 | 0.66228 | 0.02042 |
| 843 | UK2016 | TDM_421 | TDM_244 | 0.71371 | 0.02105 |
| 844 | UK2016 | TDM_498 | TDM_244 | 0.99088 | 0.00339 |
| 845 | UK2016 | TDM_54 | TDM_244 | 0.44467 | 0.02133 |
| 846 | UK2016 | TDM_275 | TDM_244 | 0.52568 | 0.01577 |
| 847 | UK2016 | TDM_509 | TDM_244 | 0.43824 | 0.02387 |
| 848 | UK2016 | TDM_04 | TDM_244 | 0.00391 | 0.00194 |
| 849 | UK2016 | TDM_1.2 | TDM_342 | 0.47634 | 0.01978 |
| 850 | UK2016 | TDM_293 | TDM_342 | 0.19598 | 0.01560 |
| 851 | UK2016 | TDM_421 | TDM_342 | 0.44453 | 0.02025 |
| 852 | UK2016 | TDM_498 | TDM_342 | 0.50723 | 0.02184 |
| 853 | UK2016 | TDM_54 | TDM_342 | 0.89747 | 0.01005 |
| 854 | UK2016 | TDM_275 | TDM_342 | 0.15755 | 0.01015 |
| 855 | UK2016 | TDM_509 | TDM_342 | 0.90763 | 0.01026 |
| 856 | UK2016 | TDM_04 | TDM_342 | 0.93329 | 0.00614 |
| 857 | UK2016 | TDM_244 | TDM_342 | 0.88657 | 0.00922 |
| 858 | UK2016 | TDM_1.2 | TDM_389 | 0.38951 | 0.01277 |
| 859 | UK2016 | TDM_293 | TDM_389 | 0.83398 | 0.00714 |
| 860 | UK2016 | TDM_421 | TDM_389 | 0.04379 | 0.00595 |
| 861 | UK2016 | TDM_498 | TDM_389 | 0.34955 | 0.01536 |

| | | | | | |
|---|---|---|---|---|---|
| 862 | UK2016 | TDM_54 | TDM_389 | 0.64874 | 0.01253 |
| 863 | UK2016 | TDM_275 | TDM_389 | 0.40833 | 0.00969 |
| 864 | UK2016 | TDM_509 | TDM_389 | 0.77385 | 0.01066 |
| 865 | UK2016 | TDM_04 | TDM_389 | 0.03116 | 0.00334 |
| 866 | UK2016 | TDM_244 | TDM_389 | 0.73372 | 0.00977 |
| 867 | UK2016 | TDM_342 | TDM_389 | 0.09438 | 0.00543 |
| 868 | UK2016 | TDM_1.2 | TDM_547 | 0.30050 | 0.02641 |
| 869 | UK2016 | TDM_293 | TDM_547 | 0.04279 | 0.00652 |
| 870 | UK2016 | TDM_421 | TDM_547 | 0.14791 | 0.02206 |
| 871 | UK2016 | TDM_498 | TDM_547 | 0.95213 | 0.00967 |
| 872 | UK2016 | TDM_54 | TDM_547 | 0.45620 | 0.02677 |
| 873 | UK2016 | TDM_275 | TDM_547 | 0.62660 | 0.01615 |
| 874 | UK2016 | TDM_509 | TDM_547 | 0.94226 | 0.01258 |
| 875 | UK2016 | TDM_04 | TDM_547 | 0.75713 | 0.01718 |
| 876 | UK2016 | TDM_244 | TDM_547 | 0.71216 | 0.02159 |
| 877 | UK2016 | TDM_342 | TDM_547 | 0.60879 | 0.02155 |
| 878 | UK2016 | TDM_389 | TDM_547 | 0.75885 | 0.01158 |
| 879 | UK2016 | TDM_1.2 | TDM_196 | 0.52582 | 0.02797 |
| 880 | UK2016 | TDM_293 | TDM_196 | 0.82744 | 0.01787 |
| 881 | UK2016 | TDM_421 | TDM_196 | 0.73453 | 0.02720 |
| 882 | UK2016 | TDM_498 | TDM_196 | 0.46383 | 0.03164 |
| 883 | UK2016 | TDM_54 | TDM_196 | 0.16378 | 0.01931 |
| 884 | UK2016 | TDM_275 | TDM_196 | 0.55833 | 0.01966 |
| 885 | UK2016 | TDM_509 | TDM_196 | 0.48222 | 0.03213 |
| 886 | UK2016 | TDM_04 | TDM_196 | 0.61054 | 0.02201 |
| 887 | UK2016 | TDM_244 | TDM_196 | 0.84595 | 0.01549 |
| 888 | UK2016 | TDM_342 | TDM_196 | 0.16071 | 0.01420 |
| 889 | UK2016 | TDM_389 | TDM_196 | 0.97149 | 0.00328 |
| 890 | UK2016 | TDM_547 | TDM_196 | 0.72685 | 0.02739 |
| 891 | UK2016 | TDM_1.2 | TDM_360 | 0.24646 | 0.01500 |
| 892 | UK2016 | TDM_293 | TDM_360 | 0.36623 | 0.01705 |
| 893 | UK2016 | TDM_421 | TDM_360 | 0.57918 | 0.02067 |
| 894 | UK2016 | TDM_498 | TDM_360 | 0.59528 | 0.01883 |
| 895 | UK2016 | TDM_54 | TDM_360 | 0.77128 | 0.01114 |
| 896 | UK2016 | TDM_275 | TDM_360 | 0.58175 | 0.01155 |
| 897 | UK2016 | TDM_509 | TDM_360 | 0.57870 | 0.02083 |
| 898 | UK2016 | TDM_04 | TDM_360 | 0.95091 | 0.00432 |
| 899 | UK2016 | TDM_244 | TDM_360 | 0.55895 | 0.01410 |
| 900 | UK2016 | TDM_342 | TDM_360 | 0.61835 | 0.01231 |
| 901 | UK2016 | TDM_389 | TDM_360 | 0.68536 | 0.00633 |
| 902 | UK2016 | TDM_547 | TDM_360 | 0.95692 | 0.00611 |
| 903 | UK2016 | TDM_196 | TDM_360 | 0.21153 | 0.01530 |
| 904 | UK2016 | TDM_1.2 | TDM_465 | 0.14230 | 0.01890 |
| 905 | UK2016 | TDM_293 | TDM_465 | 0.27522 | 0.02192 |
| 906 | UK2016 | TDM_421 | TDM_465 | 0.32340 | 0.02871 |
| 907 | UK2016 | TDM_498 | TDM_465 | 0.43592 | 0.03128 |
| 908 | UK2016 | TDM_54 | TDM_465 | 0.65901 | 0.02242 |
| 909 | UK2016 | TDM_275 | TDM_465 | 0.68201 | 0.01739 |

| | | | | | |
|---|---|---|---|---|---|
| 910 | UK2016 | TDM_509 | TDM_465 | 0.24473 | 0.02706 |
| 911 | UK2016 | TDM_04 | TDM_465 | 0.88741 | 0.01120 |
| 912 | UK2016 | TDM_244 | TDM_465 | 0.80794 | 0.01647 |
| 913 | UK2016 | TDM_342 | TDM_465 | 0.59904 | 0.01951 |
| 914 | UK2016 | TDM_389 | TDM_465 | 0.81044 | 0.00955 |
| 915 | UK2016 | TDM_547 | TDM_465 | 0.06132 | 0.01425 |
| 916 | UK2016 | TDM_196 | TDM_465 | 0.67653 | 0.02866 |
| 917 | UK2016 | TDM_360 | TDM_465 | 0.36046 | 0.01819 |
| 918 | UK2016 | TDM_1.2 | TDM_161 | 0.07134 | 0.01489 |
| 919 | UK2016 | TDM_293 | TDM_161 | 0.20511 | 0.02047 |
| 920 | UK2016 | TDM_421 | TDM_161 | 0.62386 | 0.03332 |
| 921 | UK2016 | TDM_498 | TDM_161 | 0.65907 | 0.02984 |
| 922 | UK2016 | TDM_54 | TDM_161 | 0.00972 | 0.00320 |
| 923 | UK2016 | TDM_275 | TDM_161 | 0.69357 | 0.01731 |
| 924 | UK2016 | TDM_509 | TDM_161 | 0.24056 | 0.02938 |
| 925 | UK2016 | TDM_04 | TDM_161 | 0.23735 | 0.01813 |
| 926 | UK2016 | TDM_244 | TDM_161 | 0.47401 | 0.02448 |
| 927 | UK2016 | TDM_342 | TDM_161 | 0.23698 | 0.01878 |
| 928 | UK2016 | TDM_389 | TDM_161 | 0.16918 | 0.01211 |
| 929 | UK2016 | TDM_547 | TDM_161 | 0.76577 | 0.02532 |
| 930 | UK2016 | TDM_196 | TDM_161 | 0.22553 | 0.02707 |
| 931 | UK2016 | TDM_360 | TDM_161 | 0.21900 | 0.01620 |
| 932 | UK2016 | TDM_465 | TDM_161 | 0.80676 | 0.02263 |
| 933 | UK2016 | TDM_1.2 | TDM_200 | 0.00049 | 0.00049 |
| 934 | UK2016 | TDM_293 | TDM_200 | 0.29994 | 0.02555 |
| 935 | UK2016 | TDM_421 | TDM_200 | 0.36155 | 0.03114 |
| 936 | UK2016 | TDM_498 | TDM_200 | 0.64998 | 0.02819 |
| 937 | UK2016 | TDM_54 | TDM_200 | 0.99212 | 0.00314 |
| 938 | UK2016 | TDM_275 | TDM_200 | 0.00000 | 0.00000 |
| 939 | UK2016 | TDM_509 | TDM_200 | 0.39038 | 0.03234 |
| 940 | UK2016 | TDM_04 | TDM_200 | 0.22073 | 0.02368 |
| 941 | UK2016 | TDM_244 | TDM_200 | 0.25595 | 0.02462 |
| 942 | UK2016 | TDM_342 | TDM_200 | 0.53687 | 0.02326 |
| 943 | UK2016 | TDM_389 | TDM_200 | 0.12550 | 0.01185 |
| 944 | UK2016 | TDM_547 | TDM_200 | 0.59542 | 0.03266 |
| 945 | UK2016 | TDM_196 | TDM_200 | 0.75713 | 0.02820 |
| 946 | UK2016 | TDM_360 | TDM_200 | 0.05442 | 0.00758 |
| 947 | UK2016 | TDM_465 | TDM_200 | 0.97644 | 0.00719 |
| 948 | UK2016 | TDM_161 | TDM_200 | 0.69036 | 0.02992 |
| 949 | UK2016 | TDM_1.2 | TDM_325 | 0.68750 | 0.02424 |
| 950 | UK2016 | TDM_293 | TDM_325 | 0.31276 | 0.02202 |
| 951 | UK2016 | TDM_421 | TDM_325 | 0.63234 | 0.02828 |
| 952 | UK2016 | TDM_498 | TDM_325 | 0.91626 | 0.01261 |
| 953 | UK2016 | TDM_54 | TDM_325 | 0.12698 | 0.01594 |
| 954 | UK2016 | TDM_275 | TDM_325 | 0.52198 | 0.01653 |
| 955 | UK2016 | TDM_509 | TDM_325 | 0.59461 | 0.02731 |
| 956 | UK2016 | TDM_04 | TDM_325 | 0.32719 | 0.01962 |
| 957 | UK2016 | TDM_244 | TDM_325 | 0.45562 | 0.02320 |

| | | | | | |
|---|---|---|---|---|---|
| 958 | UK2016 | TDM_342 | TDM_325 | 0.61290 | 0.01866 |
| 959 | UK2016 | TDM_389 | TDM_325 | 0.31005 | 0.01139 |
| 960 | UK2016 | TDM_547 | TDM_325 | 0.85715 | 0.01501 |
| 961 | UK2016 | TDM_196 | TDM_325 | 0.89152 | 0.01663 |
| 962 | UK2016 | TDM_360 | TDM_325 | 0.36412 | 0.01618 |
| 963 | UK2016 | TDM_465 | TDM_325 | 0.47372 | 0.02681 |
| 964 | UK2016 | TDM_161 | TDM_325 | 0.33078 | 0.02853 |
| 965 | UK2016 | TDM_200 | TDM_325 | 0.53947 | 0.03042 |
| 966 | UK2016 | TDM_1.2 | TDM_333 | 0.81776 | 0.02200 |
| 967 | UK2016 | TDM_293 | TDM_333 | 0.21114 | 0.01771 |
| 968 | UK2016 | TDM_421 | TDM_333 | 0.22845 | 0.02532 |
| 969 | UK2016 | TDM_498 | TDM_333 | 0.96303 | 0.00745 |
| 970 | UK2016 | TDM_54 | TDM_333 | 0.13536 | 0.01931 |
| 971 | UK2016 | TDM_275 | TDM_333 | 0.71724 | 0.01613 |
| 972 | UK2016 | TDM_509 | TDM_333 | 0.94006 | 0.01001 |
| 973 | UK2016 | TDM_04 | TDM_333 | 0.19807 | 0.01986 |
| 974 | UK2016 | TDM_244 | TDM_333 | 0.16828 | 0.01963 |
| 975 | UK2016 | TDM_342 | TDM_333 | 0.17759 | 0.01780 |
| 976 | UK2016 | TDM_389 | TDM_333 | 0.21631 | 0.01351 |
| 977 | UK2016 | TDM_547 | TDM_333 | 0.99605 | 0.00220 |
| 978 | UK2016 | TDM_196 | TDM_333 | 0.92072 | 0.01287 |
| 979 | UK2016 | TDM_360 | TDM_333 | 0.21491 | 0.01362 |
| 980 | UK2016 | TDM_465 | TDM_333 | 0.18579 | 0.02298 |
| 981 | UK2016 | TDM_161 | TDM_333 | 0.66687 | 0.02918 |
| 982 | UK2016 | TDM_200 | TDM_333 | 0.71723 | 0.03074 |
| 983 | UK2016 | TDM_325 | TDM_333 | 0.73203 | 0.02310 |
| 984 | UK2016 | TDM_1.2 | TDM_122 | 0.00432 | 0.00287 |
| 985 | UK2016 | TDM_293 | TDM_122 | 0.84220 | 0.01491 |
| 986 | UK2016 | TDM_421 | TDM_122 | 0.05248 | 0.01505 |
| 987 | UK2016 | TDM_498 | TDM_122 | 0.13106 | 0.01835 |
| 988 | UK2016 | TDM_54 | TDM_122 | 0.33754 | 0.02432 |
| 989 | UK2016 | TDM_275 | TDM_122 | 0.00000 | 0.00000 |
| 990 | UK2016 | TDM_509 | TDM_122 | 0.32969 | 0.03258 |
| 991 | UK2016 | TDM_04 | TDM_122 | 0.02255 | 0.00483 |
| 992 | UK2016 | TDM_244 | TDM_122 | 0.77771 | 0.01970 |
| 993 | UK2016 | TDM_342 | TDM_122 | 0.64015 | 0.02044 |
| 994 | UK2016 | TDM_389 | TDM_122 | 0.78564 | 0.00993 |
| 995 | UK2016 | TDM_547 | TDM_122 | 0.34428 | 0.02915 |
| 996 | UK2016 | TDM_196 | TDM_122 | 0.83094 | 0.01959 |
| 997 | UK2016 | TDM_360 | TDM_122 | 0.95743 | 0.00487 |
| 998 | UK2016 | TDM_465 | TDM_122 | 0.73916 | 0.02588 |
| 999 | UK2016 | TDM_161 | TDM_122 | 0.28093 | 0.02774 |
| 1000 | UK2016 | TDM_200 | TDM_122 | 0.00000 | 0.00000 |
| 1001 | UK2016 | TDM_325 | TDM_122 | 0.71955 | 0.02461 |
| 1002 | UK2016 | TDM_333 | TDM_122 | 0.52370 | 0.03195 |
| 1003 | UK2016 | TDM_1.2 | TDM_175 | 0.02557 | 0.01005 |
| 1004 | UK2016 | TDM_293 | TDM_175 | 0.06368 | 0.01107 |
| 1005 | UK2016 | TDM_421 | TDM_175 | 0.20478 | 0.02818 |

| | | | | | |
|---|---|---|---|---|---|
| 1006 | UK2016 | TDM_498 | TDM_175 | 0.73444 | 0.02800 |
| 1007 | UK2016 | TDM_54 | TDM_175 | 0.17617 | 0.02514 |
| 1008 | UK2016 | TDM_275 | TDM_175 | 0.26379 | 0.01908 |
| 1009 | UK2016 | TDM_509 | TDM_175 | 0.43395 | 0.03325 |
| 1010 | UK2016 | TDM_04 | TDM_175 | 0.77185 | 0.02103 |
| 1011 | UK2016 | TDM_244 | TDM_175 | 0.89219 | 0.01566 |
| 1012 | UK2016 | TDM_342 | TDM_175 | 0.29604 | 0.02304 |
| 1013 | UK2016 | TDM_389 | TDM_175 | 0.24296 | 0.01448 |
| 1014 | UK2016 | TDM_547 | TDM_175 | 0.52985 | 0.03334 |
| 1015 | UK2016 | TDM_196 | TDM_175 | 0.15296 | 0.02343 |
| 1016 | UK2016 | TDM_360 | TDM_175 | 0.37696 | 0.02612 |
| 1017 | UK2016 | TDM_465 | TDM_175 | 0.01204 | 0.00481 |
| 1018 | UK2016 | TDM_161 | TDM_175 | 0.97211 | 0.00957 |
| 1019 | UK2016 | TDM_200 | TDM_175 | 0.47461 | 0.03507 |
| 1020 | UK2016 | TDM_325 | TDM_175 | 0.91051 | 0.01349 |
| 1021 | UK2016 | TDM_333 | TDM_175 | 0.95507 | 0.01148 |
| 1022 | UK2016 | TDM_122 | TDM_175 | 0.54132 | 0.03181 |
| 1023 | UK2016 | TDM_1.2 | TDM_300 | 0.68214 | 0.03436 |
| 1024 | UK2016 | TDM_293 | TDM_300 | 0.16994 | 0.02679 |
| 1025 | UK2016 | TDM_421 | TDM_300 | 0.30474 | 0.03831 |
| 1026 | UK2016 | TDM_498 | TDM_300 | 0.37197 | 0.04181 |
| 1027 | UK2016 | TDM_54 | TDM_300 | 0.64543 | 0.03738 |
| 1028 | UK2016 | TDM_275 | TDM_300 | 0.15782 | 0.01803 |
| 1029 | UK2016 | TDM_509 | TDM_300 | 0.07379 | 0.02149 |
| 1030 | UK2016 | TDM_04 | TDM_300 | 0.68672 | 0.03029 |
| 1031 | UK2016 | TDM_244 | TDM_300 | 0.92248 | 0.01656 |
| 1032 | UK2016 | TDM_342 | TDM_300 | 0.45669 | 0.03138 |
| 1033 | UK2016 | TDM_389 | TDM_300 | 0.87977 | 0.01269 |
| 1034 | UK2016 | TDM_547 | TDM_300 | 0.66292 | 0.03797 |
| 1035 | UK2016 | TDM_196 | TDM_300 | 0.13201 | 0.02628 |
| 1036 | UK2016 | TDM_360 | TDM_300 | 0.53489 | 0.02922 |
| 1037 | UK2016 | TDM_465 | TDM_300 | 0.61857 | 0.03966 |
| 1038 | UK2016 | TDM_161 | TDM_300 | 0.95968 | 0.01522 |
| 1039 | UK2016 | TDM_200 | TDM_300 | 0.42426 | 0.04358 |
| 1040 | UK2016 | TDM_325 | TDM_300 | 0.98963 | 0.00542 |
| 1041 | UK2016 | TDM_333 | TDM_300 | 0.78668 | 0.03295 |
| 1042 | UK2016 | TDM_122 | TDM_300 | 0.18427 | 0.03018 |
| 1043 | UK2016 | TDM_175 | TDM_300 | 0.16307 | 0.02960 |
| 1044 | UK2016 | TDM_1.2 | TDM_1.1 | 0.19871 | 0.01957 |
| 1045 | UK2016 | TDM_293 | TDM_1.1 | 0.45712 | 0.02374 |
| 1046 | UK2016 | TDM_421 | TDM_1.1 | 0.80230 | 0.02057 |
| 1047 | UK2016 | TDM_498 | TDM_1.1 | 0.05221 | 0.01188 |
| 1048 | UK2016 | TDM_54 | TDM_1.1 | 0.61916 | 0.02398 |
| 1049 | UK2016 | TDM_275 | TDM_1.1 | 0.00000 | 0.00000 |
| 1050 | UK2016 | TDM_509 | TDM_1.1 | 0.49695 | 0.02891 |
| 1051 | UK2016 | TDM_04 | TDM_1.1 | 0.27777 | 0.02106 |
| 1052 | UK2016 | TDM_244 | TDM_1.1 | 0.08409 | 0.01038 |
| 1053 | UK2016 | TDM_342 | TDM_1.1 | 0.00912 | 0.00226 |

| | | | | | |
|---|---|---|---|---|---|
| 1054 | UK2016 | TDM_389 | TDM_1.1 | 0.45026 | 0.01313 |
| 1055 | UK2016 | TDM_547 | TDM_1.1 | 0.25528 | 0.02393 |
| 1056 | UK2016 | TDM_196 | TDM_1.1 | 0.74522 | 0.02198 |
| 1057 | UK2016 | TDM_360 | TDM_1.1 | 0.09058 | 0.00869 |
| 1058 | UK2016 | TDM_465 | TDM_1.1 | 0.90300 | 0.01211 |
| 1059 | UK2016 | TDM_161 | TDM_1.1 | 0.35779 | 0.02923 |
| 1060 | UK2016 | TDM_200 | TDM_1.1 | 0.00127 | 0.00096 |
| 1061 | UK2016 | TDM_325 | TDM_1.1 | 0.29683 | 0.02171 |
| 1062 | UK2016 | TDM_333 | TDM_1.1 | 0.96440 | 0.00800 |
| 1063 | UK2016 | TDM_122 | TDM_1.1 | 0.36277 | 0.02617 |
| 1064 | UK2016 | TDM_175 | TDM_1.1 | 0.56203 | 0.03264 |
| 1065 | UK2016 | TDM_300 | TDM_1.1 | 0.60429 | 0.03702 |
| 1066 | UK2016 | TDM_1.2 | TDM_450 | 0.12320 | 0.01715 |
| 1067 | UK2016 | TDM_293 | TDM_450 | 0.30268 | 0.02202 |
| 1068 | UK2016 | TDM_421 | TDM_450 | 0.73308 | 0.02462 |
| 1069 | UK2016 | TDM_498 | TDM_450 | 0.23990 | 0.02505 |
| 1070 | UK2016 | TDM_54 | TDM_450 | 0.88867 | 0.01318 |
| 1071 | UK2016 | TDM_275 | TDM_450 | 0.34953 | 0.01589 |
| 1072 | UK2016 | TDM_509 | TDM_450 | 0.27231 | 0.02332 |
| 1073 | UK2016 | TDM_04 | TDM_450 | 0.21988 | 0.01627 |
| 1074 | UK2016 | TDM_244 | TDM_450 | 0.32332 | 0.02009 |
| 1075 | UK2016 | TDM_342 | TDM_450 | 0.39669 | 0.01882 |
| 1076 | UK2016 | TDM_389 | TDM_450 | 0.33245 | 0.01332 |
| 1077 | UK2016 | TDM_547 | TDM_450 | 0.34058 | 0.02548 |
| 1078 | UK2016 | TDM_196 | TDM_450 | 0.63584 | 0.02543 |
| 1079 | UK2016 | TDM_360 | TDM_450 | 0.02515 | 0.00355 |
| 1080 | UK2016 | TDM_465 | TDM_450 | 0.60792 | 0.02663 |
| 1081 | UK2016 | TDM_161 | TDM_450 | 0.28223 | 0.02620 |
| 1082 | UK2016 | TDM_200 | TDM_450 | 0.04536 | 0.01091 |
| 1083 | UK2016 | TDM_325 | TDM_450 | 0.80396 | 0.01721 |
| 1084 | UK2016 | TDM_333 | TDM_450 | 0.27654 | 0.02330 |
| 1085 | UK2016 | TDM_122 | TDM_450 | 0.19523 | 0.02079 |
| 1086 | UK2016 | TDM_175 | TDM_450 | 0.48898 | 0.03061 |
| 1087 | UK2016 | TDM_300 | TDM_450 | 0.02552 | 0.01094 |
| 1088 | UK2016 | TDM_1.1 | TDM_450 | 0.58003 | 0.02412 |
| 1089 | UK2016 | TDM_1.2 | TDM_60 | 0.01910 | 0.00858 |
| 1090 | UK2016 | TDM_293 | TDM_60 | 0.81226 | 0.02021 |
| 1091 | UK2016 | TDM_421 | TDM_60 | 0.69245 | 0.03186 |
| 1092 | UK2016 | TDM_498 | TDM_60 | 0.87494 | 0.01948 |
| 1093 | UK2016 | TDM_54 | TDM_60 | 0.62537 | 0.02670 |
| 1094 | UK2016 | TDM_275 | TDM_60 | 0.00089 | 0.00052 |
| 1095 | UK2016 | TDM_509 | TDM_60 | 0.13175 | 0.02317 |
| 1096 | UK2016 | TDM_04 | TDM_60 | 0.25753 | 0.02113 |
| 1097 | UK2016 | TDM_244 | TDM_60 | 0.99253 | 0.00219 |
| 1098 | UK2016 | TDM_342 | TDM_60 | 0.35838 | 0.02150 |
| 1099 | UK2016 | TDM_389 | TDM_60 | 0.11519 | 0.01011 |
| 1100 | UK2016 | TDM_547 | TDM_60 | 0.83178 | 0.02160 |
| 1101 | UK2016 | TDM_196 | TDM_60 | 0.00768 | 0.00729 |

| | | | | | |
|---|---|---|---|---|---|
| 1102 | UK2016 | TDM_360 | TDM_60 | 0.19456 | 0.01595 |
| 1103 | UK2016 | TDM_465 | TDM_60 | 0.99757 | 0.00144 |
| 1104 | UK2016 | TDM_161 | TDM_60 | 0.84592 | 0.02206 |
| 1105 | UK2016 | TDM_200 | TDM_60 | 0.00068 | 0.00068 |
| 1106 | UK2016 | TDM_325 | TDM_60 | 0.78562 | 0.02061 |
| 1107 | UK2016 | TDM_333 | TDM_60 | 0.56593 | 0.02968 |
| 1108 | UK2016 | TDM_122 | TDM_60 | 0.00480 | 0.00406 |
| 1109 | UK2016 | TDM_175 | TDM_60 | 0.21379 | 0.02957 |
| 1110 | UK2016 | TDM_300 | TDM_60 | 0.21151 | 0.03557 |
| 1111 | UK2016 | TDM_1.1 | TDM_60 | 0.01935 | 0.00468 |
| 1112 | UK2016 | TDM_450 | TDM_60 | 0.40963 | 0.02890 |
| 1113 | UK2016 | TDM_1.2 | TDM_149 | 0.49782 | 0.02468 |
| 1114 | UK2016 | TDM_293 | TDM_149 | 0.98507 | 0.00312 |
| 1115 | UK2016 | TDM_421 | TDM_149 | 0.16233 | 0.01926 |
| 1116 | UK2016 | TDM_498 | TDM_149 | 0.42328 | 0.02636 |
| 1117 | UK2016 | TDM_54 | TDM_149 | 0.61021 | 0.02146 |
| 1118 | UK2016 | TDM_275 | TDM_149 | 0.86048 | 0.01002 |
| 1119 | UK2016 | TDM_509 | TDM_149 | 0.00496 | 0.00157 |
| 1120 | UK2016 | TDM_04 | TDM_149 | 0.17033 | 0.01474 |
| 1121 | UK2016 | TDM_244 | TDM_149 | 0.64420 | 0.01886 |
| 1122 | UK2016 | TDM_342 | TDM_149 | 0.76205 | 0.01399 |
| 1123 | UK2016 | TDM_389 | TDM_149 | 0.90926 | 0.00590 |
| 1124 | UK2016 | TDM_547 | TDM_149 | 0.53099 | 0.02454 |
| 1125 | UK2016 | TDM_196 | TDM_149 | 0.32616 | 0.02236 |
| 1126 | UK2016 | TDM_360 | TDM_149 | 0.85387 | 0.00984 |
| 1127 | UK2016 | TDM_465 | TDM_149 | 0.22218 | 0.02019 |
| 1128 | UK2016 | TDM_161 | TDM_149 | 0.15935 | 0.02211 |
| 1129 | UK2016 | TDM_200 | TDM_149 | 0.88628 | 0.01561 |
| 1130 | UK2016 | TDM_325 | TDM_149 | 0.20553 | 0.02049 |
| 1131 | UK2016 | TDM_333 | TDM_149 | 0.54393 | 0.02458 |
| 1132 | UK2016 | TDM_122 | TDM_149 | 0.56088 | 0.02483 |
| 1133 | UK2016 | TDM_175 | TDM_149 | 0.69497 | 0.02675 |
| 1134 | UK2016 | TDM_300 | TDM_149 | 0.01372 | 0.00594 |
| 1135 | UK2016 | TDM_1.1 | TDM_149 | 0.30123 | 0.01884 |
| 1136 | UK2016 | TDM_450 | TDM_149 | 0.25003 | 0.02068 |
| 1137 | UK2016 | TDM_60 | TDM_149 | 0.42894 | 0.02765 |
| 1138 | UK2016 | TDM_1.2 | TDM_224 | 0.01075 | 0.00397 |
| 1139 | UK2016 | TDM_293 | TDM_224 | 0.37057 | 0.02010 |
| 1140 | UK2016 | TDM_421 | TDM_224 | 0.13917 | 0.01868 |
| 1141 | UK2016 | TDM_498 | TDM_224 | 0.51048 | 0.02639 |
| 1142 | UK2016 | TDM_54 | TDM_224 | 0.38597 | 0.02265 |
| 1143 | UK2016 | TDM_275 | TDM_224 | 0.00000 | 0.00000 |
| 1144 | UK2016 | TDM_509 | TDM_224 | 0.62814 | 0.02631 |
| 1145 | UK2016 | TDM_04 | TDM_224 | 0.37532 | 0.01718 |
| 1146 | UK2016 | TDM_244 | TDM_224 | 0.89088 | 0.00932 |
| 1147 | UK2016 | TDM_342 | TDM_224 | 0.61750 | 0.01692 |
| 1148 | UK2016 | TDM_389 | TDM_224 | 0.57719 | 0.01231 |
| 1149 | UK2016 | TDM_547 | TDM_224 | 0.70908 | 0.02367 |

| | | | | | |
|---|---|---|---|---|---|
| 1150 | UK2016 | TDM_196 | TDM_224 | 0.60658 | 0.02592 |
| 1151 | UK2016 | TDM_360 | TDM_224 | 0.47462 | 0.01415 |
| 1152 | UK2016 | TDM_465 | TDM_224 | 0.56934 | 0.02438 |
| 1153 | UK2016 | TDM_161 | TDM_224 | 0.57827 | 0.02822 |
| 1154 | UK2016 | TDM_200 | TDM_224 | 0.00000 | 0.00000 |
| 1155 | UK2016 | TDM_325 | TDM_224 | 0.40374 | 0.02574 |
| 1156 | UK2016 | TDM_333 | TDM_224 | 0.60386 | 0.02497 |
| 1157 | UK2016 | TDM_122 | TDM_224 | 0.00000 | 0.00000 |
| 1158 | UK2016 | TDM_175 | TDM_224 | 0.18640 | 0.02295 |
| 1159 | UK2016 | TDM_300 | TDM_224 | 0.47726 | 0.04039 |
| 1160 | UK2016 | TDM_1.1 | TDM_224 | 0.28781 | 0.02252 |
| 1161 | UK2016 | TDM_450 | TDM_224 | 0.48131 | 0.02504 |
| 1162 | UK2016 | TDM_60 | TDM_224 | 0.09649 | 0.01690 |
| 1163 | UK2016 | TDM_149 | TDM_224 | 0.93638 | 0.00914 |
| 1164 | UK2016 | TDM_1.2 | TDM_323 | 0.99437 | 0.00410 |
| 1165 | UK2016 | TDM_293 | TDM_323 | 0.87908 | 0.01653 |
| 1166 | UK2016 | TDM_421 | TDM_323 | 0.02923 | 0.00829 |
| 1167 | UK2016 | TDM_498 | TDM_323 | 0.34011 | 0.03444 |
| 1168 | UK2016 | TDM_54 | TDM_323 | 0.87978 | 0.01902 |
| 1169 | UK2016 | TDM_275 | TDM_323 | 0.71782 | 0.02043 |
| 1170 | UK2016 | TDM_509 | TDM_323 | 0.72643 | 0.03079 |
| 1171 | UK2016 | TDM_04 | TDM_323 | 0.48310 | 0.02397 |
| 1172 | UK2016 | TDM_244 | TDM_323 | 0.62202 | 0.03055 |
| 1173 | UK2016 | TDM_342 | TDM_323 | 0.15133 | 0.01868 |
| 1174 | UK2016 | TDM_389 | TDM_323 | 0.71300 | 0.01547 |
| 1175 | UK2016 | TDM_547 | TDM_323 | 0.11435 | 0.01950 |
| 1176 | UK2016 | TDM_196 | TDM_323 | 0.97082 | 0.00879 |
| 1177 | UK2016 | TDM_360 | TDM_323 | 0.72955 | 0.01914 |
| 1178 | UK2016 | TDM_465 | TDM_323 | 0.04866 | 0.01580 |
| 1179 | UK2016 | TDM_161 | TDM_323 | 0.22115 | 0.03173 |
| 1180 | UK2016 | TDM_200 | TDM_323 | 0.75033 | 0.03230 |
| 1181 | UK2016 | TDM_325 | TDM_323 | 0.50838 | 0.03102 |
| 1182 | UK2016 | TDM_333 | TDM_323 | 0.53624 | 0.03520 |
| 1183 | UK2016 | TDM_122 | TDM_323 | 0.87612 | 0.01953 |
| 1184 | UK2016 | TDM_175 | TDM_323 | 0.77839 | 0.02758 |
| 1185 | UK2016 | TDM_300 | TDM_323 | 0.84211 | 0.03351 |
| 1186 | UK2016 | TDM_1.1 | TDM_323 | 0.40802 | 0.03011 |
| 1187 | UK2016 | TDM_450 | TDM_323 | 0.89734 | 0.01728 |
| 1188 | UK2016 | TDM_60 | TDM_323 | 0.16605 | 0.03006 |
| 1189 | UK2016 | TDM_149 | TDM_323 | 0.11251 | 0.01807 |
| 1190 | UK2016 | TDM_224 | TDM_323 | 0.81802 | 0.02224 |
| 1191 | UK2016 | TDM_1.2 | TDM_880 | 0.35330 | 0.01973 |
| 1192 | UK2016 | TDM_293 | TDM_880 | 0.60610 | 0.01903 |
| 1193 | UK2016 | TDM_421 | TDM_880 | 0.01284 | 0.00430 |
| 1194 | UK2016 | TDM_498 | TDM_880 | 0.53670 | 0.02197 |
| 1195 | UK2016 | TDM_54 | TDM_880 | 0.32784 | 0.01925 |
| 1196 | UK2016 | TDM_275 | TDM_880 | 0.60360 | 0.01407 |
| 1197 | UK2016 | TDM_509 | TDM_880 | 0.24450 | 0.02086 |

| | | | | | |
|---|---|---|---|---|---|
| 1198 | UK2016 | TDM_04 | TDM_880 | 0.06404 | 0.00852 |
| 1199 | UK2016 | TDM_244 | TDM_880 | 0.03172 | 0.00449 |
| 1200 | UK2016 | TDM_342 | TDM_880 | 0.92546 | 0.00528 |
| 1201 | UK2016 | TDM_389 | TDM_880 | 0.67430 | 0.01013 |
| 1202 | UK2016 | TDM_547 | TDM_880 | 0.32841 | 0.01947 |
| 1203 | UK2016 | TDM_196 | TDM_880 | 0.30340 | 0.02244 |
| 1204 | UK2016 | TDM_360 | TDM_880 | 0.34560 | 0.01303 |
| 1205 | UK2016 | TDM_465 | TDM_880 | 0.21155 | 0.02040 |
| 1206 | UK2016 | TDM_161 | TDM_880 | 0.55634 | 0.02664 |
| 1207 | UK2016 | TDM_200 | TDM_880 | 0.65000 | 0.02131 |
| 1208 | UK2016 | TDM_325 | TDM_880 | 0.98328 | 0.00376 |
| 1209 | UK2016 | TDM_333 | TDM_880 | 0.29928 | 0.02096 |
| 1210 | UK2016 | TDM_122 | TDM_880 | 0.30098 | 0.02227 |
| 1211 | UK2016 | TDM_175 | TDM_880 | 0.22854 | 0.02138 |
| 1212 | UK2016 | TDM_300 | TDM_880 | 0.78205 | 0.02599 |
| 1213 | UK2016 | TDM_1.1 | TDM_880 | 0.05297 | 0.00860 |
| 1214 | UK2016 | TDM_450 | TDM_880 | 0.38790 | 0.01961 |
| 1215 | UK2016 | TDM_60 | TDM_880 | 0.41133 | 0.02427 |
| 1216 | UK2016 | TDM_149 | TDM_880 | 0.96377 | 0.00537 |
| 1217 | UK2016 | TDM_224 | TDM_880 | 0.77486 | 0.01523 |
| 1218 | UK2016 | TDM_323 | TDM_880 | 0.76747 | 0.02149 |
| 1219 | UK2014 | TDM_1.2 | TDM_293 | 0.42866 | 0.02529 |
| 1220 | UK2014 | TDM_1.2 | TDM_421 | 0.57023 | 0.02894 |
| 1221 | UK2014 | TDM_293 | TDM_421 | 0.11213 | 0.01768 |
| 1222 | UK2014 | TDM_1.2 | TDM_498 | 0.62172 | 0.02801 |
| 1223 | UK2014 | TDM_293 | TDM_498 | 0.06856 | 0.01369 |
| 1224 | UK2014 | TDM_421 | TDM_498 | 0.10493 | 0.01915 |
| 1225 | UK2014 | TDM_1.2 | TDM_54 | 0.08480 | 0.01387 |
| 1226 | UK2014 | TDM_293 | TDM_54 | 0.13464 | 0.01665 |
| 1227 | UK2014 | TDM_421 | TDM_54 | 0.18895 | 0.01886 |
| 1228 | UK2014 | TDM_498 | TDM_54 | 0.53011 | 0.02734 |
| 1229 | UK2014 | TDM_1.2 | TDM_275 | 0.02799 | 0.00825 |
| 1230 | UK2014 | TDM_293 | TDM_275 | 0.14584 | 0.01533 |
| 1231 | UK2014 | TDM_421 | TDM_275 | 0.48824 | 0.02234 |
| 1232 | UK2014 | TDM_498 | TDM_275 | 0.22881 | 0.02101 |
| 1233 | UK2014 | TDM_54 | TDM_275 | 0.01207 | 0.00366 |
| 1234 | UK2014 | TDM_1.2 | TDM_509 | 0.49982 | 0.03066 |
| 1235 | UK2014 | TDM_293 | TDM_509 | 0.76411 | 0.02279 |
| 1236 | UK2014 | TDM_421 | TDM_509 | 0.81586 | 0.02308 |
| 1237 | UK2014 | TDM_498 | TDM_509 | 0.69474 | 0.03364 |
| 1238 | UK2014 | TDM_54 | TDM_509 | 0.05343 | 0.01155 |
| 1239 | UK2014 | TDM_275 | TDM_509 | 0.61812 | 0.02122 |
| 1240 | UK2014 | TDM_1.2 | TDM_04 | 0.09186 | 0.01305 |
| 1241 | UK2014 | TDM_293 | TDM_04 | 0.84555 | 0.01313 |
| 1242 | UK2014 | TDM_421 | TDM_04 | 0.98049 | 0.00461 |
| 1243 | UK2014 | TDM_498 | TDM_04 | 0.59610 | 0.02295 |
| 1244 | UK2014 | TDM_54 | TDM_04 | 0.21973 | 0.01507 |
| 1245 | UK2014 | TDM_275 | TDM_04 | 0.57474 | 0.01697 |

| | | | | | |
|---|---|---|---|---|---|
| 1246 | UK2014 | TDM_509 | TDM_04 | 0.41715 | 0.02203 |
| 1247 | UK2014 | TDM_1.2 | TDM_244 | 0.31117 | 0.02449 |
| 1248 | UK2014 | TDM_293 | TDM_244 | 0.40991 | 0.02404 |
| 1249 | UK2014 | TDM_421 | TDM_244 | 0.16605 | 0.02044 |
| 1250 | UK2014 | TDM_498 | TDM_244 | 0.03270 | 0.00763 |
| 1251 | UK2014 | TDM_54 | TDM_244 | 0.61323 | 0.02097 |
| 1252 | UK2014 | TDM_275 | TDM_244 | 0.15569 | 0.01426 |
| 1253 | UK2014 | TDM_509 | TDM_244 | 0.48249 | 0.02908 |
| 1254 | UK2014 | TDM_04 | TDM_244 | 0.08107 | 0.00889 |
| 1255 | UK2014 | TDM_1.2 | TDM_342 | 0.90426 | 0.01113 |
| 1256 | UK2014 | TDM_293 | TDM_342 | 0.79281 | 0.01957 |
| 1257 | UK2014 | TDM_421 | TDM_342 | 0.74943 | 0.02145 |
| 1258 | UK2014 | TDM_498 | TDM_342 | 0.27272 | 0.02484 |
| 1259 | UK2014 | TDM_54 | TDM_342 | 0.70636 | 0.02147 |
| 1260 | UK2014 | TDM_275 | TDM_342 | 0.54322 | 0.02033 |
| 1261 | UK2014 | TDM_509 | TDM_342 | 0.50616 | 0.02913 |
| 1262 | UK2014 | TDM_04 | TDM_342 | 0.12963 | 0.01239 |
| 1263 | UK2014 | TDM_244 | TDM_342 | 0.48295 | 0.02261 |
| 1264 | UK2014 | TDM_1.2 | TDM_389 | 0.54623 | 0.01889 |
| 1265 | UK2014 | TDM_293 | TDM_389 | 0.17796 | 0.01734 |
| 1266 | UK2014 | TDM_421 | TDM_389 | 0.19977 | 0.01902 |
| 1267 | UK2014 | TDM_498 | TDM_389 | 0.46449 | 0.02461 |
| 1268 | UK2014 | TDM_54 | TDM_389 | 0.54147 | 0.01867 |
| 1269 | UK2014 | TDM_275 | TDM_389 | 0.93636 | 0.00640 |
| 1270 | UK2014 | TDM_509 | TDM_389 | 0.76099 | 0.01757 |
| 1271 | UK2014 | TDM_04 | TDM_389 | 0.98849 | 0.00198 |
| 1272 | UK2014 | TDM_244 | TDM_389 | 0.14850 | 0.01384 |
| 1273 | UK2014 | TDM_342 | TDM_389 | 0.14637 | 0.01210 |
| 1274 | UK2014 | TDM_1.2 | TDM_547 | 0.29735 | 0.02923 |
| 1275 | UK2014 | TDM_293 | TDM_547 | 0.31350 | 0.02933 |
| 1276 | UK2014 | TDM_421 | TDM_547 | 0.68008 | 0.03007 |
| 1277 | UK2014 | TDM_498 | TDM_547 | 0.39897 | 0.03425 |
| 1278 | UK2014 | TDM_54 | TDM_547 | 0.56070 | 0.02663 |
| 1279 | UK2014 | TDM_275 | TDM_547 | 0.40317 | 0.02348 |
| 1280 | UK2014 | TDM_509 | TDM_547 | 0.11150 | 0.01956 |
| 1281 | UK2014 | TDM_04 | TDM_547 | 0.62164 | 0.02258 |
| 1282 | UK2014 | TDM_244 | TDM_547 | 0.48626 | 0.02776 |
| 1283 | UK2014 | TDM_342 | TDM_547 | 0.14736 | 0.01935 |
| 1284 | UK2014 | TDM_389 | TDM_547 | 0.11361 | 0.01444 |
| 1285 | UK2014 | TDM_1.2 | TDM_196 | 0.10360 | 0.01922 |
| 1286 | UK2014 | TDM_293 | TDM_196 | 0.45313 | 0.03571 |
| 1287 | UK2014 | TDM_421 | TDM_196 | 0.46731 | 0.03422 |
| 1288 | UK2014 | TDM_498 | TDM_196 | 0.18168 | 0.03273 |
| 1289 | UK2014 | TDM_54 | TDM_196 | 0.45149 | 0.03299 |
| 1290 | UK2014 | TDM_275 | TDM_196 | 0.32815 | 0.02908 |
| 1291 | UK2014 | TDM_509 | TDM_196 | 0.51279 | 0.04048 |
| 1292 | UK2014 | TDM_04 | TDM_196 | 0.78506 | 0.02360 |
| 1293 | UK2014 | TDM_244 | TDM_196 | 0.35934 | 0.03158 |

| | | | | | |
|---|---|---|---|---|---|
| 1294 | UK2014 | TDM_342 | TDM_196 | 0.44941 | 0.03457 |
| 1295 | UK2014 | TDM_389 | TDM_196 | 0.37004 | 0.02858 |
| 1296 | UK2014 | TDM_547 | TDM_196 | 0.90176 | 0.02009 |
| 1297 | UK2014 | TDM_1.2 | TDM_360 | 0.88302 | 0.01430 |
| 1298 | UK2014 | TDM_293 | TDM_360 | 0.82152 | 0.01977 |
| 1299 | UK2014 | TDM_421 | TDM_360 | 0.92626 | 0.01157 |
| 1300 | UK2014 | TDM_498 | TDM_360 | 0.60867 | 0.02713 |
| 1301 | UK2014 | TDM_54 | TDM_360 | 0.89190 | 0.01047 |
| 1302 | UK2014 | TDM_275 | TDM_360 | 0.91736 | 0.00955 |
| 1303 | UK2014 | TDM_509 | TDM_360 | 0.71381 | 0.02233 |
| 1304 | UK2014 | TDM_04 | TDM_360 | 0.27052 | 0.01563 |
| 1305 | UK2014 | TDM_244 | TDM_360 | 0.58059 | 0.02222 |
| 1306 | UK2014 | TDM_342 | TDM_360 | 0.43900 | 0.02372 |
| 1307 | UK2014 | TDM_389 | TDM_360 | 0.91378 | 0.00824 |
| 1308 | UK2014 | TDM_547 | TDM_360 | 0.29114 | 0.02489 |
| 1309 | UK2014 | TDM_196 | TDM_360 | 0.10350 | 0.01949 |
| 1310 | UK2014 | TDM_1.2 | TDM_465 | 0.45416 | 0.03418 |
| 1311 | UK2014 | TDM_293 | TDM_465 | 0.11636 | 0.01811 |
| 1312 | UK2014 | TDM_421 | TDM_465 | 0.90706 | 0.02047 |
| 1313 | UK2014 | TDM_498 | TDM_465 | 0.32943 | 0.03372 |
| 1314 | UK2014 | TDM_54 | TDM_465 | 0.89020 | 0.01576 |
| 1315 | UK2014 | TDM_275 | TDM_465 | 0.44643 | 0.02613 |
| 1316 | UK2014 | TDM_509 | TDM_465 | 0.61527 | 0.03103 |
| 1317 | UK2014 | TDM_04 | TDM_465 | 0.43165 | 0.02344 |
| 1318 | UK2014 | TDM_244 | TDM_465 | 0.26655 | 0.02859 |
| 1319 | UK2014 | TDM_342 | TDM_465 | 0.40067 | 0.02829 |
| 1320 | UK2014 | TDM_389 | TDM_465 | 0.51949 | 0.02507 |
| 1321 | UK2014 | TDM_547 | TDM_465 | 0.66214 | 0.03118 |
| 1322 | UK2014 | TDM_196 | TDM_465 | 0.23286 | 0.03326 |
| 1323 | UK2014 | TDM_360 | TDM_465 | 0.75299 | 0.02105 |
| 1324 | UK2014 | TDM_1.2 | TDM_161 | 0.37239 | 0.03318 |
| 1325 | UK2014 | TDM_293 | TDM_161 | 0.86622 | 0.01863 |
| 1326 | UK2014 | TDM_421 | TDM_161 | 0.97941 | 0.00790 |
| 1327 | UK2014 | TDM_498 | TDM_161 | 0.95739 | 0.01109 |
| 1328 | UK2014 | TDM_54 | TDM_161 | 0.61297 | 0.02825 |
| 1329 | UK2014 | TDM_275 | TDM_161 | 0.69263 | 0.02310 |
| 1330 | UK2014 | TDM_509 | TDM_161 | 0.85579 | 0.02357 |
| 1331 | UK2014 | TDM_04 | TDM_161 | 0.66316 | 0.02553 |
| 1332 | UK2014 | TDM_244 | TDM_161 | 0.45103 | 0.03128 |
| 1333 | UK2014 | TDM_342 | TDM_161 | 0.65078 | 0.02462 |
| 1334 | UK2014 | TDM_389 | TDM_161 | 0.32612 | 0.02594 |
| 1335 | UK2014 | TDM_547 | TDM_161 | 0.91196 | 0.02005 |
| 1336 | UK2014 | TDM_196 | TDM_161 | 0.10320 | 0.02496 |
| 1337 | UK2014 | TDM_360 | TDM_161 | 0.79534 | 0.02231 |
| 1338 | UK2014 | TDM_465 | TDM_161 | 0.23903 | 0.03175 |
| 1339 | UK2014 | TDM_1.2 | TDM_200 | 0.85900 | 0.02219 |
| 1340 | UK2014 | TDM_293 | TDM_200 | 0.38597 | 0.03180 |
| 1341 | UK2014 | TDM_421 | TDM_200 | 0.05327 | 0.01685 |

| | | | | | |
|---|---|---|---|---|---|
| 1342 | UK2014 | TDM_498 | TDM_200 | 0.90358 | 0.02053 |
| 1343 | UK2014 | TDM_54 | TDM_200 | 0.56913 | 0.03179 |
| 1344 | UK2014 | TDM_275 | TDM_200 | 0.00000 | 0.00000 |
| 1345 | UK2014 | TDM_509 | TDM_200 | 0.70200 | 0.03102 |
| 1346 | UK2014 | TDM_04 | TDM_200 | 0.30818 | 0.02414 |
| 1347 | UK2014 | TDM_244 | TDM_200 | 0.38932 | 0.03232 |
| 1348 | UK2014 | TDM_342 | TDM_200 | 0.23387 | 0.02697 |
| 1349 | UK2014 | TDM_389 | TDM_200 | 0.59746 | 0.02503 |
| 1350 | UK2014 | TDM_547 | TDM_200 | 0.12847 | 0.02563 |
| 1351 | UK2014 | TDM_196 | TDM_200 | 0.66895 | 0.03667 |
| 1352 | UK2014 | TDM_360 | TDM_200 | 0.04171 | 0.01073 |
| 1353 | UK2014 | TDM_465 | TDM_200 | 0.50921 | 0.03773 |
| 1354 | UK2014 | TDM_161 | TDM_200 | 0.80093 | 0.03142 |
| 1355 | UK2014 | TDM_1.2 | TDM_325 | 0.22189 | 0.02256 |
| 1356 | UK2014 | TDM_293 | TDM_325 | 0.16501 | 0.01859 |
| 1357 | UK2014 | TDM_421 | TDM_325 | 0.83029 | 0.02336 |
| 1358 | UK2014 | TDM_498 | TDM_325 | 0.42207 | 0.03161 |
| 1359 | UK2014 | TDM_54 | TDM_325 | 0.01702 | 0.00515 |
| 1360 | UK2014 | TDM_275 | TDM_325 | 0.28164 | 0.02130 |
| 1361 | UK2014 | TDM_509 | TDM_325 | 0.98442 | 0.00467 |
| 1362 | UK2014 | TDM_04 | TDM_325 | 0.15230 | 0.01362 |
| 1363 | UK2014 | TDM_244 | TDM_325 | 0.87696 | 0.01581 |
| 1364 | UK2014 | TDM_342 | TDM_325 | 0.88865 | 0.01364 |
| 1365 | UK2014 | TDM_389 | TDM_325 | 0.23827 | 0.01783 |
| 1366 | UK2014 | TDM_547 | TDM_325 | 0.77834 | 0.02438 |
| 1367 | UK2014 | TDM_196 | TDM_325 | 0.79234 | 0.02978 |
| 1368 | UK2014 | TDM_360 | TDM_325 | 0.72486 | 0.02121 |
| 1369 | UK2014 | TDM_465 | TDM_325 | 0.45011 | 0.03269 |
| 1370 | UK2014 | TDM_161 | TDM_325 | 0.08021 | 0.01637 |
| 1371 | UK2014 | TDM_200 | TDM_325 | 0.17381 | 0.02672 |
| 1372 | UK2014 | TDM_1.2 | TDM_333 | 0.19987 | 0.02122 |
| 1373 | UK2014 | TDM_293 | TDM_333 | 0.53841 | 0.02398 |
| 1374 | UK2014 | TDM_421 | TDM_333 | 0.84194 | 0.01694 |
| 1375 | UK2014 | TDM_498 | TDM_333 | 0.24456 | 0.02613 |
| 1376 | UK2014 | TDM_54 | TDM_333 | 0.81050 | 0.01564 |
| 1377 | UK2014 | TDM_275 | TDM_333 | 0.38132 | 0.02063 |
| 1378 | UK2014 | TDM_509 | TDM_333 | 0.71031 | 0.02319 |
| 1379 | UK2014 | TDM_04 | TDM_333 | 0.02182 | 0.00497 |
| 1380 | UK2014 | TDM_244 | TDM_333 | 0.45604 | 0.02315 |
| 1381 | UK2014 | TDM_342 | TDM_333 | 0.45322 | 0.02316 |
| 1382 | UK2014 | TDM_389 | TDM_333 | 0.97023 | 0.00430 |
| 1383 | UK2014 | TDM_547 | TDM_333 | 0.77182 | 0.02025 |
| 1384 | UK2014 | TDM_196 | TDM_333 | 0.75175 | 0.02775 |
| 1385 | UK2014 | TDM_360 | TDM_333 | 0.81842 | 0.01412 |
| 1386 | UK2014 | TDM_465 | TDM_333 | 0.63302 | 0.02895 |
| 1387 | UK2014 | TDM_161 | TDM_333 | 0.73450 | 0.02685 |
| 1388 | UK2014 | TDM_200 | TDM_333 | 0.49984 | 0.03255 |
| 1389 | UK2014 | TDM_325 | TDM_333 | 0.64810 | 0.02486 |

| | | | | | |
|---|---|---|---|---|---|
| 1390 | UK2014 | TDM_1.2 | TDM_122 | 0.01588 | 0.00647 |
| 1391 | UK2014 | TDM_293 | TDM_122 | 0.96077 | 0.00957 |
| 1392 | UK2014 | TDM_421 | TDM_122 | 0.41581 | 0.03203 |
| 1393 | UK2014 | TDM_498 | TDM_122 | 0.73619 | 0.02717 |
| 1394 | UK2014 | TDM_54 | TDM_122 | 0.36302 | 0.02604 |
| 1395 | UK2014 | TDM_275 | TDM_122 | 0.00000 | 0.00000 |
| 1396 | UK2014 | TDM_509 | TDM_122 | 0.65969 | 0.03000 |
| 1397 | UK2014 | TDM_04 | TDM_122 | 0.02745 | 0.00725 |
| 1398 | UK2014 | TDM_244 | TDM_122 | 0.49451 | 0.02883 |
| 1399 | UK2014 | TDM_342 | TDM_122 | 0.25135 | 0.02413 |
| 1400 | UK2014 | TDM_389 | TDM_122 | 0.58287 | 0.02370 |
| 1401 | UK2014 | TDM_547 | TDM_122 | 0.68625 | 0.03100 |
| 1402 | UK2014 | TDM_196 | TDM_122 | 0.61629 | 0.03755 |
| 1403 | UK2014 | TDM_360 | TDM_122 | 0.28835 | 0.02678 |
| 1404 | UK2014 | TDM_465 | TDM_122 | 0.30692 | 0.03016 |
| 1405 | UK2014 | TDM_161 | TDM_122 | 0.99122 | 0.00405 |
| 1406 | UK2014 | TDM_200 | TDM_122 | 0.00049 | 0.00049 |
| 1407 | UK2014 | TDM_325 | TDM_122 | 0.36537 | 0.03155 |
| 1408 | UK2014 | TDM_333 | TDM_122 | 0.59214 | 0.02647 |
| 1409 | UK2014 | TDM_1.2 | TDM_175 | 0.82777 | 0.01580 |
| 1410 | UK2014 | TDM_293 | TDM_175 | 0.56528 | 0.02072 |
| 1411 | UK2014 | TDM_421 | TDM_175 | 0.67900 | 0.02049 |
| 1412 | UK2014 | TDM_498 | TDM_175 | 0.45094 | 0.02614 |
| 1413 | UK2014 | TDM_54 | TDM_175 | 0.01298 | 0.00345 |
| 1414 | UK2014 | TDM_275 | TDM_175 | 0.91610 | 0.00736 |
| 1415 | UK2014 | TDM_509 | TDM_175 | 0.03175 | 0.00735 |
| 1416 | UK2014 | TDM_04 | TDM_175 | 0.35978 | 0.01634 |
| 1417 | UK2014 | TDM_244 | TDM_175 | 0.89105 | 0.00891 |
| 1418 | UK2014 | TDM_342 | TDM_175 | 0.57510 | 0.01687 |
| 1419 | UK2014 | TDM_389 | TDM_175 | 0.38281 | 0.01572 |
| 1420 | UK2014 | TDM_547 | TDM_175 | 0.75771 | 0.02050 |
| 1421 | UK2014 | TDM_196 | TDM_175 | 0.35326 | 0.02840 |
| 1422 | UK2014 | TDM_360 | TDM_175 | 0.15395 | 0.01232 |
| 1423 | UK2014 | TDM_465 | TDM_175 | 0.55981 | 0.02576 |
| 1424 | UK2014 | TDM_161 | TDM_175 | 0.60759 | 0.02816 |
| 1425 | UK2014 | TDM_200 | TDM_175 | 0.07758 | 0.01552 |
| 1426 | UK2014 | TDM_325 | TDM_175 | 0.69166 | 0.02011 |
| 1427 | UK2014 | TDM_333 | TDM_175 | 0.63112 | 0.01814 |
| 1428 | UK2014 | TDM_122 | TDM_175 | 0.29550 | 0.02162 |
| 1429 | UK2014 | TDM_1.2 | TDM_300 | 0.85110 | 0.02265 |
| 1430 | UK2014 | TDM_293 | TDM_300 | 0.73932 | 0.02863 |
| 1431 | UK2014 | TDM_421 | TDM_300 | 0.90883 | 0.01849 |
| 1432 | UK2014 | TDM_498 | TDM_300 | 0.00134 | 0.00095 |
| 1433 | UK2014 | TDM_54 | TDM_300 | 0.06559 | 0.01486 |
| 1434 | UK2014 | TDM_275 | TDM_300 | 0.93909 | 0.01127 |
| 1435 | UK2014 | TDM_509 | TDM_300 | 0.54399 | 0.03674 |
| 1436 | UK2014 | TDM_04 | TDM_300 | 0.46130 | 0.02799 |
| 1437 | UK2014 | TDM_244 | TDM_300 | 0.90437 | 0.01887 |

| | | | | | |
|---|---|---|---|---|---|
| 1438 | UK2014 | TDM_342 | TDM_300 | 0.23081 | 0.02725 |
| 1439 | UK2014 | TDM_389 | TDM_300 | 0.67977 | 0.02409 |
| 1440 | UK2014 | TDM_547 | TDM_300 | 0.94583 | 0.01497 |
| 1441 | UK2014 | TDM_196 | TDM_300 | 0.39760 | 0.03881 |
| 1442 | UK2014 | TDM_360 | TDM_300 | 0.43019 | 0.02905 |
| 1443 | UK2014 | TDM_465 | TDM_300 | 0.67471 | 0.03594 |
| 1444 | UK2014 | TDM_161 | TDM_300 | 0.27619 | 0.03429 |
| 1445 | UK2014 | TDM_200 | TDM_300 | 0.81314 | 0.02958 |
| 1446 | UK2014 | TDM_325 | TDM_300 | 0.21593 | 0.03018 |
| 1447 | UK2014 | TDM_333 | TDM_300 | 0.53989 | 0.03411 |
| 1448 | UK2014 | TDM_122 | TDM_300 | 0.20421 | 0.02873 |
| 1449 | UK2014 | TDM_175 | TDM_300 | 0.79881 | 0.02364 |
| 1450 | UK2014 | TDM_1.2 | TDM_1.1 | 0.34470 | 0.02573 |
| 1451 | UK2014 | TDM_293 | TDM_1.1 | 0.22163 | 0.02316 |
| 1452 | UK2014 | TDM_421 | TDM_1.1 | 0.37740 | 0.02572 |
| 1453 | UK2014 | TDM_498 | TDM_1.1 | 0.65765 | 0.02643 |
| 1454 | UK2014 | TDM_54 | TDM_1.1 | 0.44475 | 0.02516 |
| 1455 | UK2014 | TDM_275 | TDM_1.1 | 0.03584 | 0.00679 |
| 1456 | UK2014 | TDM_509 | TDM_1.1 | 0.20454 | 0.02294 |
| 1457 | UK2014 | TDM_04 | TDM_1.1 | 0.36958 | 0.01905 |
| 1458 | UK2014 | TDM_244 | TDM_1.1 | 0.23603 | 0.02112 |
| 1459 | UK2014 | TDM_342 | TDM_1.1 | 0.25911 | 0.01978 |
| 1460 | UK2014 | TDM_389 | TDM_1.1 | 0.03792 | 0.00725 |
| 1461 | UK2014 | TDM_547 | TDM_1.1 | 0.54590 | 0.02816 |
| 1462 | UK2014 | TDM_196 | TDM_1.1 | 0.63301 | 0.03317 |
| 1463 | UK2014 | TDM_360 | TDM_1.1 | 0.50722 | 0.02268 |
| 1464 | UK2014 | TDM_465 | TDM_1.1 | 0.01758 | 0.00575 |
| 1465 | UK2014 | TDM_161 | TDM_1.1 | 0.79614 | 0.02326 |
| 1466 | UK2014 | TDM_200 | TDM_1.1 | 0.64284 | 0.03352 |
| 1467 | UK2014 | TDM_325 | TDM_1.1 | 0.30206 | 0.02397 |
| 1468 | UK2014 | TDM_333 | TDM_1.1 | 0.38167 | 0.02441 |
| 1469 | UK2014 | TDM_122 | TDM_1.1 | 0.89894 | 0.01553 |
| 1470 | UK2014 | TDM_175 | TDM_1.1 | 0.12505 | 0.01224 |
| 1471 | UK2014 | TDM_300 | TDM_1.1 | 0.93246 | 0.01412 |
| 1472 | UK2014 | TDM_1.2 | TDM_450 | 0.90412 | 0.01574 |
| 1473 | UK2014 | TDM_293 | TDM_450 | 0.80337 | 0.02667 |
| 1474 | UK2014 | TDM_421 | TDM_450 | 0.93741 | 0.01569 |
| 1475 | UK2014 | TDM_498 | TDM_450 | 0.25208 | 0.02978 |
| 1476 | UK2014 | TDM_54 | TDM_450 | 0.24368 | 0.02625 |
| 1477 | UK2014 | TDM_275 | TDM_450 | 0.44604 | 0.02497 |
| 1478 | UK2014 | TDM_509 | TDM_450 | 0.57977 | 0.03321 |
| 1479 | UK2014 | TDM_04 | TDM_450 | 0.45275 | 0.02383 |
| 1480 | UK2014 | TDM_244 | TDM_450 | 0.83625 | 0.01924 |
| 1481 | UK2014 | TDM_342 | TDM_450 | 0.79025 | 0.02264 |
| 1482 | UK2014 | TDM_389 | TDM_450 | 0.17624 | 0.01940 |
| 1483 | UK2014 | TDM_547 | TDM_450 | 0.00586 | 0.00321 |
| 1484 | UK2014 | TDM_196 | TDM_450 | 0.65601 | 0.03744 |
| 1485 | UK2014 | TDM_360 | TDM_450 | 0.24426 | 0.02810 |

| | | | | | |
|---|---|---|---|---|---|
| 1486 | UK2014 | TDM_465 | TDM_450 | 0.53562 | 0.03679 |
| 1487 | UK2014 | TDM_161 | TDM_450 | 0.31601 | 0.03540 |
| 1488 | UK2014 | TDM_200 | TDM_450 | 0.53627 | 0.03778 |
| 1489 | UK2014 | TDM_325 | TDM_450 | 0.63368 | 0.03098 |
| 1490 | UK2014 | TDM_333 | TDM_450 | 0.48154 | 0.02800 |
| 1491 | UK2014 | TDM_122 | TDM_450 | 0.05167 | 0.01272 |
| 1492 | UK2014 | TDM_175 | TDM_450 | 0.71807 | 0.02454 |
| 1493 | UK2014 | TDM_300 | TDM_450 | 0.07480 | 0.02078 |
| 1494 | UK2014 | TDM_1.1 | TDM_450 | 0.23444 | 0.02389 |
| 1495 | UK2014 | TDM_1.2 | TDM_60 | 0.74293 | 0.02593 |
| 1496 | UK2014 | TDM_293 | TDM_60 | 0.35408 | 0.03280 |
| 1497 | UK2014 | TDM_421 | TDM_60 | 0.37662 | 0.02972 |
| 1498 | UK2014 | TDM_498 | TDM_60 | 0.82303 | 0.02567 |
| 1499 | UK2014 | TDM_54 | TDM_60 | 0.01169 | 0.00505 |
| 1500 | UK2014 | TDM_275 | TDM_60 | 0.00884 | 0.00287 |
| 1501 | UK2014 | TDM_509 | TDM_60 | 0.99605 | 0.00218 |
| 1502 | UK2014 | TDM_04 | TDM_60 | 0.43824 | 0.02134 |
| 1503 | UK2014 | TDM_244 | TDM_60 | 0.31787 | 0.02788 |
| 1504 | UK2014 | TDM_342 | TDM_60 | 0.15662 | 0.01790 |
| 1505 | UK2014 | TDM_389 | TDM_60 | 0.18687 | 0.01872 |
| 1506 | UK2014 | TDM_547 | TDM_60 | 0.14141 | 0.02241 |
| 1507 | UK2014 | TDM_196 | TDM_60 | 0.33117 | 0.03761 |
| 1508 | UK2014 | TDM_360 | TDM_60 | 0.90587 | 0.01392 |
| 1509 | UK2014 | TDM_465 | TDM_60 | 0.21273 | 0.02853 |
| 1510 | UK2014 | TDM_161 | TDM_60 | 0.72544 | 0.02889 |
| 1511 | UK2014 | TDM_200 | TDM_60 | 0.01014 | 0.00636 |
| 1512 | UK2014 | TDM_325 | TDM_60 | 0.43442 | 0.03190 |
| 1513 | UK2014 | TDM_333 | TDM_60 | 0.32794 | 0.02567 |
| 1514 | UK2014 | TDM_122 | TDM_60 | 0.16840 | 0.02523 |
| 1515 | UK2014 | TDM_175 | TDM_60 | 0.65163 | 0.02419 |
| 1516 | UK2014 | TDM_300 | TDM_60 | 0.77403 | 0.03280 |
| 1517 | UK2014 | TDM_1.1 | TDM_60 | 0.00000 | 0.00000 |
| 1518 | UK2014 | TDM_450 | TDM_60 | 0.96674 | 0.00867 |
| 1519 | UK2014 | TDM_1.2 | TDM_149 | 0.85402 | 0.01764 |
| 1520 | UK2014 | TDM_293 | TDM_149 | 0.76152 | 0.02258 |
| 1521 | UK2014 | TDM_421 | TDM_149 | 0.22388 | 0.02472 |
| 1522 | UK2014 | TDM_498 | TDM_149 | 0.50925 | 0.02867 |
| 1523 | UK2014 | TDM_54 | TDM_149 | 0.73545 | 0.02251 |
| 1524 | UK2014 | TDM_275 | TDM_149 | 0.44027 | 0.02177 |
| 1525 | UK2014 | TDM_509 | TDM_149 | 0.94251 | 0.01121 |
| 1526 | UK2014 | TDM_04 | TDM_149 | 0.18788 | 0.01511 |
| 1527 | UK2014 | TDM_244 | TDM_149 | 0.57924 | 0.02514 |
| 1528 | UK2014 | TDM_342 | TDM_149 | 0.92470 | 0.01071 |
| 1529 | UK2014 | TDM_389 | TDM_149 | 0.67082 | 0.01864 |
| 1530 | UK2014 | TDM_547 | TDM_149 | 0.33559 | 0.02791 |
| 1531 | UK2014 | TDM_196 | TDM_149 | 0.41297 | 0.03488 |
| 1532 | UK2014 | TDM_360 | TDM_149 | 0.28511 | 0.02298 |
| 1533 | UK2014 | TDM_465 | TDM_149 | 0.95921 | 0.01112 |

| | | | | | |
|---|---|---|---|---|---|
| 1534 | UK2014 | TDM_161 | TDM_149 | 0.47780 | 0.03058 |
| 1535 | UK2014 | TDM_200 | TDM_149 | 0.50008 | 0.03279 |
| 1536 | UK2014 | TDM_325 | TDM_149 | 0.15664 | 0.02127 |
| 1537 | UK2014 | TDM_333 | TDM_149 | 0.71971 | 0.02126 |
| 1538 | UK2014 | TDM_122 | TDM_149 | 0.10840 | 0.01713 |
| 1539 | UK2014 | TDM_175 | TDM_149 | 0.31315 | 0.02260 |
| 1540 | UK2014 | TDM_300 | TDM_149 | 0.22523 | 0.02894 |
| 1541 | UK2014 | TDM_1.1 | TDM_149 | 0.10257 | 0.01528 |
| 1542 | UK2014 | TDM_450 | TDM_149 | 0.02204 | 0.00871 |
| 1543 | UK2014 | TDM_60 | TDM_149 | 0.64116 | 0.02917 |
| 1544 | UK2014 | TDM_1.2 | TDM_224 | 0.01011 | 0.00427 |
| 1545 | UK2014 | TDM_293 | TDM_224 | 0.90704 | 0.01700 |
| 1546 | UK2014 | TDM_421 | TDM_224 | 0.98338 | 0.00652 |
| 1547 | UK2014 | TDM_498 | TDM_224 | 0.53258 | 0.03451 |
| 1548 | UK2014 | TDM_54 | TDM_224 | 0.42354 | 0.03085 |
| 1549 | UK2014 | TDM_275 | TDM_224 | 0.00000 | 0.00000 |
| 1550 | UK2014 | TDM_509 | TDM_224 | 0.60865 | 0.03436 |
| 1551 | UK2014 | TDM_04 | TDM_224 | 0.48632 | 0.02418 |
| 1552 | UK2014 | TDM_244 | TDM_224 | 0.43736 | 0.02700 |
| 1553 | UK2014 | TDM_342 | TDM_224 | 0.40762 | 0.03222 |
| 1554 | UK2014 | TDM_389 | TDM_224 | 0.06117 | 0.00985 |
| 1555 | UK2014 | TDM_547 | TDM_224 | 0.41532 | 0.03729 |
| 1556 | UK2014 | TDM_196 | TDM_224 | 0.38237 | 0.03916 |
| 1557 | UK2014 | TDM_360 | TDM_224 | 0.70067 | 0.02740 |
| 1558 | UK2014 | TDM_465 | TDM_224 | 0.73613 | 0.03116 |
| 1559 | UK2014 | TDM_161 | TDM_224 | 0.92737 | 0.01691 |
| 1560 | UK2014 | TDM_200 | TDM_224 | 0.00204 | 0.00176 |
| 1561 | UK2014 | TDM_325 | TDM_224 | 0.36956 | 0.03350 |
| 1562 | UK2014 | TDM_333 | TDM_224 | 0.53389 | 0.02874 |
| 1563 | UK2014 | TDM_122 | TDM_224 | 0.00000 | 0.00000 |
| 1564 | UK2014 | TDM_175 | TDM_224 | 0.95638 | 0.00951 |
| 1565 | UK2014 | TDM_300 | TDM_224 | 0.75172 | 0.03343 |
| 1566 | UK2014 | TDM_1.1 | TDM_224 | 0.41842 | 0.03417 |
| 1567 | UK2014 | TDM_450 | TDM_224 | 0.59818 | 0.03708 |
| 1568 | UK2014 | TDM_60 | TDM_224 | 0.00900 | 0.00451 |
| 1569 | UK2014 | TDM_149 | TDM_224 | 0.37121 | 0.03203 |
| 1570 | UK2014 | TDM_1.2 | TDM_323 | 0.24078 | 0.03143 |
| 1571 | UK2014 | TDM_293 | TDM_323 | 0.42287 | 0.03694 |
| 1572 | UK2014 | TDM_421 | TDM_323 | 0.88538 | 0.02117 |
| 1573 | UK2014 | TDM_498 | TDM_323 | 0.64453 | 0.03743 |
| 1574 | UK2014 | TDM_54 | TDM_323 | 0.15440 | 0.02207 |
| 1575 | UK2014 | TDM_275 | TDM_323 | 0.70125 | 0.02555 |
| 1576 | UK2014 | TDM_509 | TDM_323 | 0.28726 | 0.03514 |
| 1577 | UK2014 | TDM_04 | TDM_323 | 0.22101 | 0.02441 |
| 1578 | UK2014 | TDM_244 | TDM_323 | 0.16853 | 0.02380 |
| 1579 | UK2014 | TDM_342 | TDM_323 | 0.38535 | 0.02892 |
| 1580 | UK2014 | TDM_389 | TDM_323 | 0.92051 | 0.01447 |
| 1581 | UK2014 | TDM_547 | TDM_323 | 0.17193 | 0.02746 |

| | | | | | |
|---|---|---|---|---|---|
| 1582 | UK2014 | TDM_196 | TDM_323 | 0.06623 | 0.01872 |
| 1583 | UK2014 | TDM_360 | TDM_323 | 0.07365 | 0.01637 |
| 1584 | UK2014 | TDM_465 | TDM_323 | 0.67018 | 0.03911 |
| 1585 | UK2014 | TDM_161 | TDM_323 | 0.54522 | 0.04339 |
| 1586 | UK2014 | TDM_200 | TDM_323 | 0.17089 | 0.03225 |
| 1587 | UK2014 | TDM_325 | TDM_323 | 0.25830 | 0.03324 |
| 1588 | UK2014 | TDM_333 | TDM_323 | 0.63955 | 0.03398 |
| 1589 | UK2014 | TDM_122 | TDM_323 | 0.79738 | 0.03140 |
| 1590 | UK2014 | TDM_175 | TDM_323 | 0.19350 | 0.02167 |
| 1591 | UK2014 | TDM_300 | TDM_323 | 0.24688 | 0.03592 |
| 1592 | UK2014 | TDM_1.1 | TDM_323 | 0.73792 | 0.02881 |
| 1593 | UK2014 | TDM_450 | TDM_323 | 0.13175 | 0.02604 |
| 1594 | UK2014 | TDM_60 | TDM_323 | 0.05419 | 0.01772 |
| 1595 | UK2014 | TDM_149 | TDM_323 | 0.23429 | 0.03072 |
| 1596 | UK2014 | TDM_224 | TDM_323 | 0.56353 | 0.03986 |
| 1597 | UK2014 | TDM_1.2 | TDM_880 | 0.33288 | 0.03266 |
| 1598 | UK2014 | TDM_293 | TDM_880 | 0.09517 | 0.01948 |
| 1599 | UK2014 | TDM_421 | TDM_880 | 0.29996 | 0.03260 |
| 1600 | UK2014 | TDM_498 | TDM_880 | 0.65816 | 0.03352 |
| 1601 | UK2014 | TDM_54 | TDM_880 | 0.51308 | 0.02949 |
| 1602 | UK2014 | TDM_275 | TDM_880 | 0.51481 | 0.02681 |
| 1603 | UK2014 | TDM_509 | TDM_880 | 0.10355 | 0.01904 |
| 1604 | UK2014 | TDM_04 | TDM_880 | 0.91418 | 0.01115 |
| 1605 | UK2014 | TDM_244 | TDM_880 | 0.05801 | 0.01157 |
| 1606 | UK2014 | TDM_342 | TDM_880 | 0.15765 | 0.02138 |
| 1607 | UK2014 | TDM_389 | TDM_880 | 0.87127 | 0.01526 |
| 1608 | UK2014 | TDM_547 | TDM_880 | 0.00988 | 0.00567 |
| 1609 | UK2014 | TDM_196 | TDM_880 | 0.77651 | 0.03367 |
| 1610 | UK2014 | TDM_360 | TDM_880 | 0.60056 | 0.02914 |
| 1611 | UK2014 | TDM_465 | TDM_880 | 0.44773 | 0.03767 |
| 1612 | UK2014 | TDM_161 | TDM_880 | 0.36430 | 0.03814 |
| 1613 | UK2014 | TDM_200 | TDM_880 | 0.98186 | 0.00738 |
| 1614 | UK2014 | TDM_325 | TDM_880 | 0.43475 | 0.03195 |
| 1615 | UK2014 | TDM_333 | TDM_880 | 0.02748 | 0.00796 |
| 1616 | UK2014 | TDM_122 | TDM_880 | 0.99424 | 0.00336 |
| 1617 | UK2014 | TDM_175 | TDM_880 | 0.11756 | 0.01762 |
| 1618 | UK2014 | TDM_300 | TDM_880 | 0.92710 | 0.01746 |
| 1619 | UK2014 | TDM_1.1 | TDM_880 | 0.35726 | 0.02995 |
| 1620 | UK2014 | TDM_450 | TDM_880 | 0.91796 | 0.02005 |
| 1621 | UK2014 | TDM_60 | TDM_880 | 0.21626 | 0.02788 |
| 1622 | UK2014 | TDM_149 | TDM_880 | 0.44302 | 0.03338 |
| 1623 | UK2014 | TDM_224 | TDM_880 | 0.72059 | 0.03413 |
| 1624 | UK2014 | TDM_323 | TDM_880 | 0.30471 | 0.03764 |
